# Evidence of adaptive host and vector manipulation by plant viruses revealed through combined meta-analysis and modeling approaches

**DOI:** 10.1101/781690

**Authors:** Quentin Chesnais, Christie A. Bahlai, Angela Peace, David W. Crowder, Nilsa A. Bosque-Pérez, Kerry Mauck

## Abstract

A growing number of studies indicate that plant viruses enhance their own transmission by modifying host phenotypes and vector behavior, leading to the hypothesis that such effects are manipulations resulting from virus adaptations. However, few studies have linked putative manipulations with virus components, and the true frequency and magnitude of host and vector manipulation across virus taxa remains unknown. To address this knowledge gap, we performed a meta-analysis to quantify convergence in virus effects on hosts and vectors across taxonomic groups that share transmission mechanism traits, and thereby stand to benefit from similar sequences of vector behavior. We then combined meta-analysis outputs with an epidemiological model to assess consequences of manipulation for virus spread. Overall, transmission mechanism traits strongly predicted the magnitude and nature of virus effects on vector preferences and performance. Models parameterized with meta-analysis data demonstrate that manipulation effects enhance virus spread, and that viruses with long acquisition times and retention durations are under strong selection pressure to manipulate transmission. By combining meta-analysis with epidemiological modeling, our results confirm that host and vector manipulation are important aspects of plant virus ecology and evolution while emphasizing the need to incorporate more pathosystems and transmission mechanism traits in future studies.

## Introduction

Arthropod-borne plant viruses are ubiquitous, obligate biotrophic parasites. To exploit hosts, plant viruses have evolved adaptations for suppressing host immunity, co-opting host resources for reproduction, and augmenting vascular connections to enable systemic movement of virus particles (Pazhouhandeh *et al*. 2006; Patarroyo *et al*. 2012; Rojas *et al*. 2016; Yang & Li 2018). The host physiological changes that result (symptoms), and the virus genes underlying their expression, are studied primarily because of their economically important effects on plant health. However, symptoms of infection can also affect interactions with arthropod vectors via changes in plant cues mediating host seeking and feeding behaviors, particularly visual characteristics (color, size, shape), odor profiles, palatability, defense chemistry, and nutritional quality (Ngumbi *et al*. 2007; Mauck *et al*. 2014a, b, 2018; Casteel *et al*. 2015; Peñaflor *et al*. 2016); reviewed in (Mauck *et al*. 2018). The importance of vector behavior for virus fitness has led to the hypothesis that viruses might evolve adaptations for eliciting specific symptoms (host phenotypes) that increase transmission-conducive interactions between arthropod vectors and infected hosts.

There are now over 120 published studies that test this hypothesis using combinations of behavioral and biological assays, as well as techniques for plant phenotyping. Many report virus effects on host phenotypes and vector behavior that appear to be cases of adaptive host manipulation – an instance of a parasite evolving to control elements of its host’s phenotype that help maintain or enhance rates of transmission (Poulin 2010). However, despite the growing number of studies on plant virus manipulation of hosts and vectors, we lack a quantitative synthesis of how host phenotypes vary depending on the traits of the viral pathogens under study, including transmission mechanism traits that govern how viruses are acquired, retained, and inoculated by arthropod vectors (Nault 1997; Ng & Falk 2006; Hogenhout *et al*. 2008; Ng & Zhou 2015). This limits our ability to determine whether putative instances of vector manipulation by plant viruses are a result of virus adaptations, or simply by-products of pathology (Thomas *et al*. 2005). For example, if putative manipulations are the product of adaptations, viruses transmitted via the same sequences of vector behavior may exhibit convergence in their effects on plant cues mediating vector-host interactions that lead to efficient transmission (Thomas *et al*. 2005; Mauck *et al*. 2010, 2016). Additionally, a lack of synthesis around virus traits creates a disconnect between the emerging evidence for virus manipulation and other historically rooted fields, including epidemiology, molecular virology, and virus ecology (Malmstrom *et al*. 2011; Alexander *et al*. 2014). As a result, the true frequency and relevance of host manipulation by plant viruses remains unknown.

To address these knowledge gaps, we combined a meta-analysis with mathematical modeling and a review of taxon-specific virus-vector relationships to evaluate the case for plant virus manipulation of hosts and vectors in the context of virus traits underlying the transmission process. We used traits shared by phylogenetically divergent virus lineages, namely infection location in the plant and retention mechanism in the vector, as a framework for evaluating evidence for or against adaptive host manipulation. Within this framework, we quantified the magnitude and direction of virus effects on host plant attractiveness, palatability, and quality to arthropod vectors. We also derived parameter estimates from these data and incorporated them into a model that was explicitly designed to explore virus effects on host-vector relationships in the context of virus traits (Shaw *et al*. 2017). Finally, we interpreted our results in the context of documented virus-vector relationships that influence the ecology of the major virus lineages targeted in our study.

### Predictions based on virus traits

Empirical studies of plant viruses inducing manipulations of hosts and vectors assess vector preferences and performance on infected and non-infected plants as proxies to understand how virus effects on host phenotypes influence transmission. When evaluated across diverse pathosystems, these experiments can serve as an important tool for exploring the adaptive significance of virus effects on host phenotypes (Thomas *et al*. 2005; Mauck *et al*. 2016). Phylogenetically unrelated plant viruses may exhibit convergence in their effects on host phenotypes based on shared virus traits; specifically, requirements for vectors to engage in a narrow suite of behaviors necessary for transmission. Similar convergence in manipulation strategies is apparent across diverse lineages of animal-infecting parasites transmitted by blood-feeding vectors (Thomas *et al*. 2005; Lefèvre & Thomas 2008), supporting the hypothesis that such effects are adaptive. This evidence is essential for understanding manipulation because many of these parasites and their hosts are intractable for functional genomics work to identify genes, and gene targets that may underlie adaptive manipulation (Heil 2016). Convergence of virus effects in the absence of phylogenetic relatedness provides indirect evidence in support of particular effects being the product of virus adaptations rather than by-products of pathology (Thomas *et al*. 2005; Mauck *et al*. 2018).

Here, we used virus traits associated with transmission as a framework for quantifying the adaptive significance of virus effects on host phenotypes using meta-analysis. Viruses can be broadly classified based on the types and durations of vector probing and feeding behaviors required for virion acquisition from, and inoculation to, the host (acquisition/inoculation site) (Brault *et al*. 2010) and the persistence of virions in the vector (retention mechanism) (Fig. 1). Phloem-limited (**PL**) viruses are restricted to the host vascular tissue, whereby acquisition of sufficient virions for transmission depends on vectors engaging in sustained phloem sap ingestion for hours or days (Hogenhout *et al*. 2008; Brault *et al*. 2010). Following acquisition, most PL viruses are retained in the vector for several days (non-circulative, semi-persistent [**NCSP**] viruses) or the duration of the vector’s lifespan (circulative, persistent viruses), and can be inoculated to multiple hosts without re-acquisition. Within the circulative, persistently-transmitted retention mechanism, some viruses traverse the gut barrier and hemolymph to colonize the salivary glands (circulative persistent, non-propagative [**CPNPr**]), while others colonize and replicate in various vector tissues, effectively using the vector as a second host (circulative persistent, propagative [**CPPr**]) (Hogenhout *et al*. 2008). For both CPNPr and CPPr retention mechanisms, as long as the vector survives and periodically moves among hosts, a single acquisition event can lead to multiple new infections.

**Figure 1:**
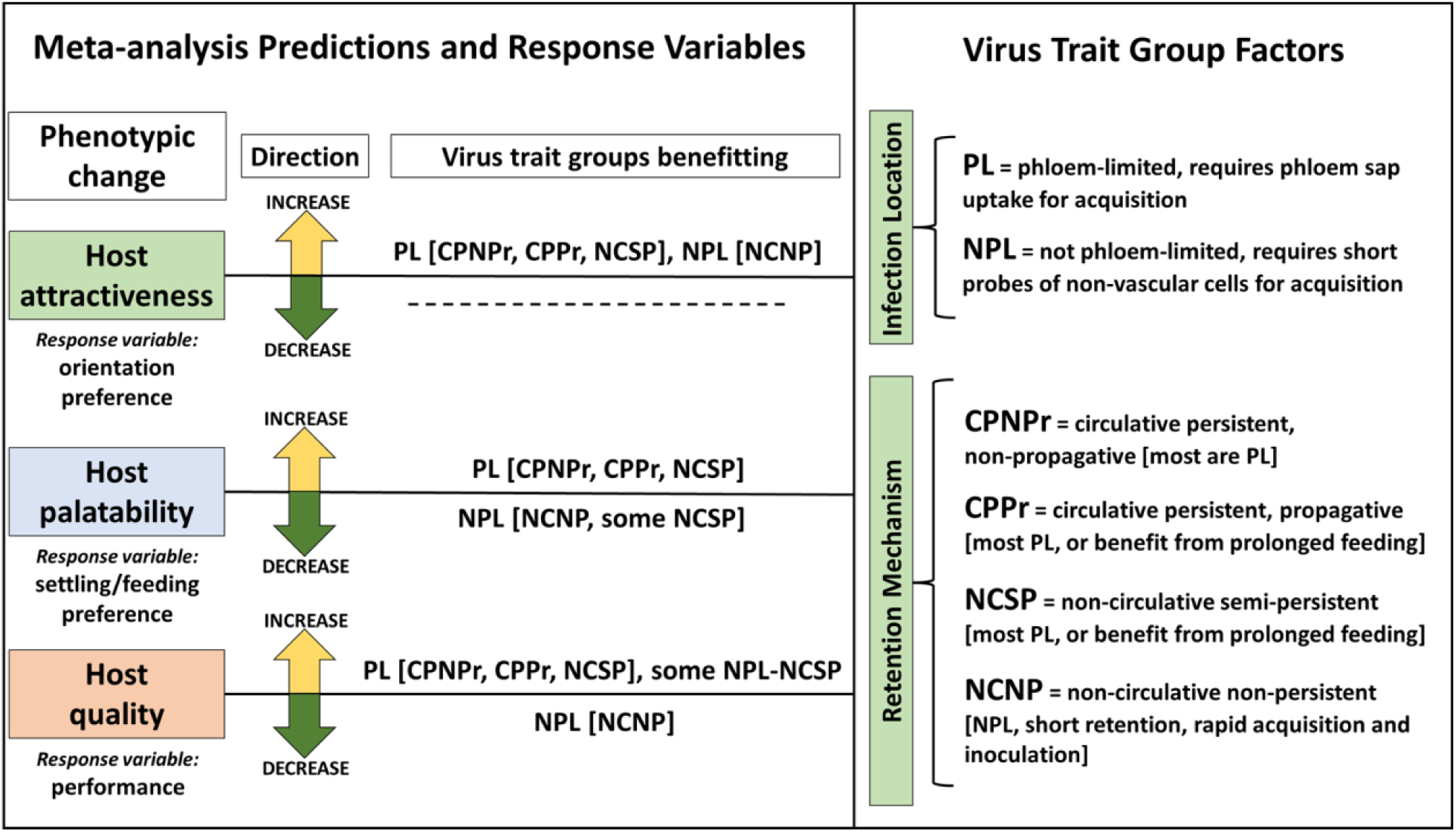
Predictions, response variables, and factors included in the meta-analysis.

In contrast, most non-phloem-limited (**NPL**) viruses are acquired and inoculated following brief probes of non-vascular tissues such as epidermal or mesophyll cells (Martin *et al*. 1997; Ng & Falk 2006). These non-circulative, non-persistent (**NCNP**) viruses account for approximately 40% of all known vector-borne plant viruses, and are retained for very short periods of time following acquisition, which limits inoculation potential for a viruliferous vector (a vector carrying the virus and capable of transmitting) to about 1-2 plants (Nault 1997; Hogenhout *et al*. 2008). NPL-NCNP viruses are also rapidly lost from vector mouthparts. As a result, the spread of most NPL viruses is favored by rapid dispersal of vectors from infected to receptive hosts (Martin *et al*. 1997; Wang & Ghabrial 2002; Nault 1997; Hogenhout *et al*. 2008; Martin *et al*. 1997; Wang & Ghabrial 2002).

The clear delineations of host infection location (PL *vs*. NPL) and retention mechanism in the vector (NCSP, CPNPr, CPPr and NCNP) provide a convenient framework for evaluating the adaptive significance of host and vector manipulation by plant viruses. Here, we applied this framework to quantify the effects of plant virus infection on three responses: (i) vector orientation preferences (host selection), (ii) vector settling/feeding behavior, and (iii) vector performance. We predicted that viruses from all trait groups should induce host phenotypic changes that result in vector orientation preferences for infected hosts over healthy ones because this increases vector contacts (Fig. 1). We further predicted that enhancements to vector settling/feeding and performance would only be apparent for trait groups that stand to benefit from sustained phloem sap ingestion and production of offspring that will remain viruliferous for long periods following virus acquisition (Fig. 1). We tested these predictions using a meta-analysis of 126 published studies covering 59 viruses belonging to 11 families, studied in association with host plants of 15 different families. Results were interpreted in the context of complementary model simulations and additional ecological dimensions known to affect virus transmission by insect vectors.

## Materials and Methods

### Database assembly

To obtain studies related to virus–host–vector interactions, we conducted an extensive literature search in the ISI Web of Knowledge database and Google Scholar following (Mauck *et al*. 2012). We used a combination of broad search terms including “virus-host-vector interactions”, “plant virus”, “insect vector”, “non-persistently transmitted virus”, “persistent-circulative virus”, “persistent-propagative virus”, “plant virus chemical ecology”, “vector behavior”, and “vector performance” along with specific search terms (family and species names of viruses and their vectors) to identify studies that assesses insect vector attraction, settling and feeding, and performance in relation to infected and non-infected plants. We also surveyed references in review articles about virus–host–vector interactions (Fereres & Moreno 2009; Mauck *et al*. 2010, 2012, 2018; Bosque-Pérez & Eigenbrode 2011; Eigenbrode & Bosque-Perez 2016; Eigenbrode *et al*. 2018). Complete criteria for study inclusion and data selection, the assembled database, and a complete list of references are provided in the electronic supplementary material (ESM_meta-analysis.docx). Data were obtained from tables, or extracted from plots using Plot Digitizer (Huwaldt & Steinhorst 2013). To avoid bias and use all the available data, we recorded multiple data points from a single study if it examined more than one relevant response variable or included multiple host plants, viruses, or vectors.

For each non-infected/infected plant comparison, we recorded the mean, standard deviation and sample size of the relevant response variable measuring either vector orientation preference, settling/feeding behavior, or performance. Orientation preference was defined as any vector response to plant cues without physical contact with the plant, such as studies that used olfactometers. Settling/feeding preference was defined as any behavioral response to the plants that occurred following host contact, such as settling preference, retention time on a host, and time to dispersal as well as feeding behaviors (electrical penetration graph technique or other metrics that quantify ease of feeding on preferred tissues). Vector performance was defined as any physiological response to plants known to affect vector reproduction and/or longevity at the individual or colony level (development time, survival, fecundity, weight, population growth).

We also documented variables related to virus traits (Fig. 1). These included traits associated with transmission (infection location in the plant and retention mechanism in the vector). The retention mechanism for persistent viruses also includes circulation and colonization of the vector (CPNPr), as well as propagation within vectors for a subset of these pathogens (CPPr). Thus, we also considered these virus traits by separating these two retention mechanisms in the analysis. For the analysis by virus family, we included families for which there were at least three independent measures of vector behaviors or performance (ESM_meta-analysis.docx for complete analyses, table S2abc). The full database is available as part of the electronic supplementary data (ESM_Meta-analysis database.xlsx).

### Effect size calculation

For each non-infected/infected plant comparison in the database, we calculated the virus infection effect size using the Hedges’ *g* metric and its confidence interval (CI) (Hedges 1981). The metric is calculated as *g* = [(*Xi* − *Xh*)/*s*] *J*, where *Xi* represents the mean of the vector parameter on the infected plant, *Xh* represents the mean of the vector parameter on the non-infected plant, *s* represents the pooled standard deviation, and *J* is a correction factor for small sample size (Koricheva *et al*. 2013). Positive Hedges’ *g* values indicate that the vector preferred or performed better on infected compared to non-infected plants, whereas negative values indicate they preferred or performed better on non-infected plants. When necessary we reversed the sign of the effect size so that a negative value of *g* always indicates a negative effect of virus infection; for example, decreased development time on infected plants represents increased rather than decreased performance. The Hedges’ *g* and its estimated sampling variance were calculated using the *‘escalc’* function in the ‘*metafor’* package in *R* 3.6.0 (Viechtbauer 2010).

### Meta-analysis model construction

We fit multilevel mixed-effects models using the ‘*rma.mv’* function in the R package *metafor* that weighted each effect size by the inverse of its sampling variance plus the amount of residual heterogeneity not explained by moderators (i.e., additional variables that help us understand the relationships between the dependent and independent variables) (Viechtbauer 2010). To account for the non-independence of data derived from the same paper, we assigned each study case a single identifier (Study ID), corresponding to a single published paper retained in our analysis. We included the study ID as a random effect term in all models.

### Hypothesis testing and meta-regression

To address whether virus infection impacted vector orientation preference, settling/feeding behaviors, and vector performance, we fit random-effects models separately to the vector orientation preference data, vector settling/feeding data, and vector performance data using restricted maximum likelihood (REML). We considered model-estimated mean effect sizes with 95% confidence intervals (CIs) that did not cross zero as evidence for a significant effect.

Initially, we calculated a mean effect size across all studies to assess whether there was an overall effect of plant infection on vector parameters. Second, we tested how various moderators influenced the magnitude of the virus infection effect using meta-regression models conducted separately for each moderator. When significant effects were detected for moderators with more than two groups, the meta-analysis was followed by post-hoc comparisons among groups, carried out using the *multcomp* package in *R* (Hothorn *et al*. 2008). To assess whether there was a significant effect of virus infection for each group, we re-fitted models with no intercepts and the model coefficients and their associated CIs were used to determine whether the effect size was different from zero for each group.

### Heterogeneity statistics and bias analysis

For each mixed-effects model, we assessed residual heterogeneity using the QE statistic (Viechtbauer 2010; Koricheva *et al*. 2013). We found significant QE values for all models (*P* < .0001, ESM_meta-analysis.docx, table S1), suggesting there were important moderators that we did not include in analyses. To assess the potential for publication bias to influence our conclusions, we used funnel plots and meta-regression models with “study year” and “plant domestication” as moderators (Koricheva *et al*. 2013). We found a low probability that publication bias affected our results (ESM_meta-analysis.docx, figures S1, S2 and S3), except that the effect size on vector performance is significantly higher on wild plants than cultivated (*P* = .0003, ESM_meta-analysis.docx, table S1, figure S3). The fail-safe numbers for plant virus infection (overall effects) were also calculated for each dataset. Fail-safe numbers indicate the number of nonsignificant unpublished or missing studies that would negate the results, and are considered robust against publication bias if they are > 5*n* + 10 where *n* is number of studies (Rosenthal 1979).

### Modeling implications for virus spread

The data assembled for the meta-analysis provided an opportunity to leverage information on effect sizes to assess how virus-induced manipulation of hosts and vectors may affect the rate of transmission. To accomplish this, we used a published model (Shaw *et al*. 2017) that was constructed to accommodate a comparison of aphid-transmitted viruses with a phloem-limited, CPNPr infection/retention mechanism and a non-phloem-limited NCNP infection/retention mechanism (see electronic supplementary material file ESM_model parameters.docx). These categories capture the characteristics of most pathosystems in our meta-analysis (ESM_Meta-analysis database.xlsx). We modified the model in several ways to complement the meta-analysis outputs and data. Briefly, these modifications included (1) substitution of new parameters for virus effects (vector orientation preference for infected hosts [δ] [attraction], maximum vector departure rate from infected hosts [*a*i] [settling and feeding], and intrinsic vector growth rate on infected hosts [*r*i] [performance]); (2) modification of parameter values for dispersal loss (μ) and the rate at which vectors become viruliferous (βv) based on values in additional published literature; and (3) changing the vector recovery rate term (γ) to more accurately represent PL-CPNPr and NPL-NCNP transmission mechanisms. Table 1 shows a summary of the model parameters, values and confidence intervals used here, and a full description of model and parameter modifications is provided in the supplementary material (ESM_model parameters.docx).

**Table 1.**
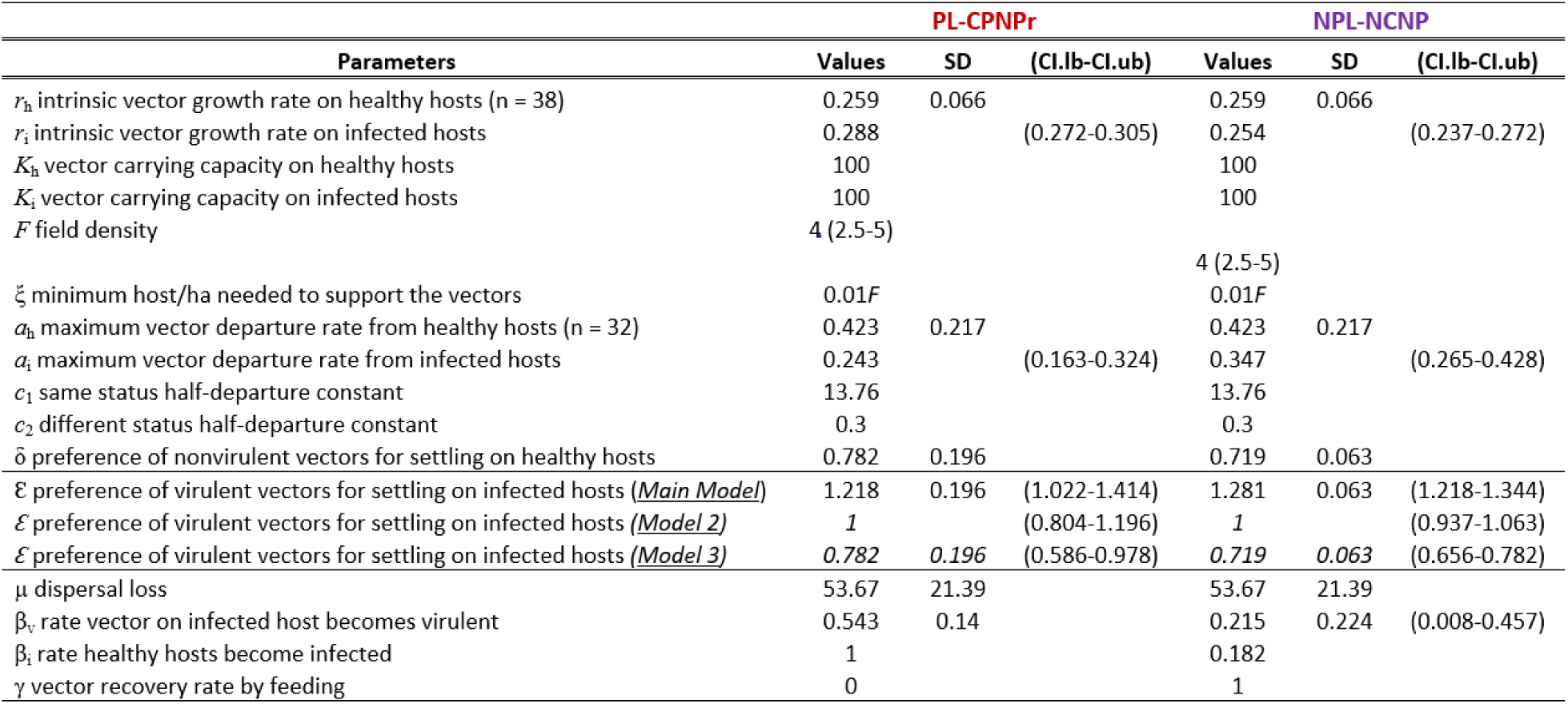
Model parameters (adapted from Shaw *et al*. 2017).

The model explores the preference and performance of vectors before (non-viruliferous) and after acquiring a virus (viruliferous). It tracks the number of viruliferous (V) and non-viruliferous (N) vectors, and the fraction of healthy (H) and infected (I) hosts using a system of ordinary differential equations. To incorporate behavioral preferences, the model tracks the infection status of the host that each vector is on. There are four compartments for the vector population: N_h_ number of non-viruliferous vectors on healthy hosts, N_i_ number of non-viruliferous vectors on infected hosts, V_h_ number of viruliferous vectors on healthy hosts, and V_i_ number of viruliferous vectors on infected hosts. The modified model equations are shown in Table 2.

**Table 2.**
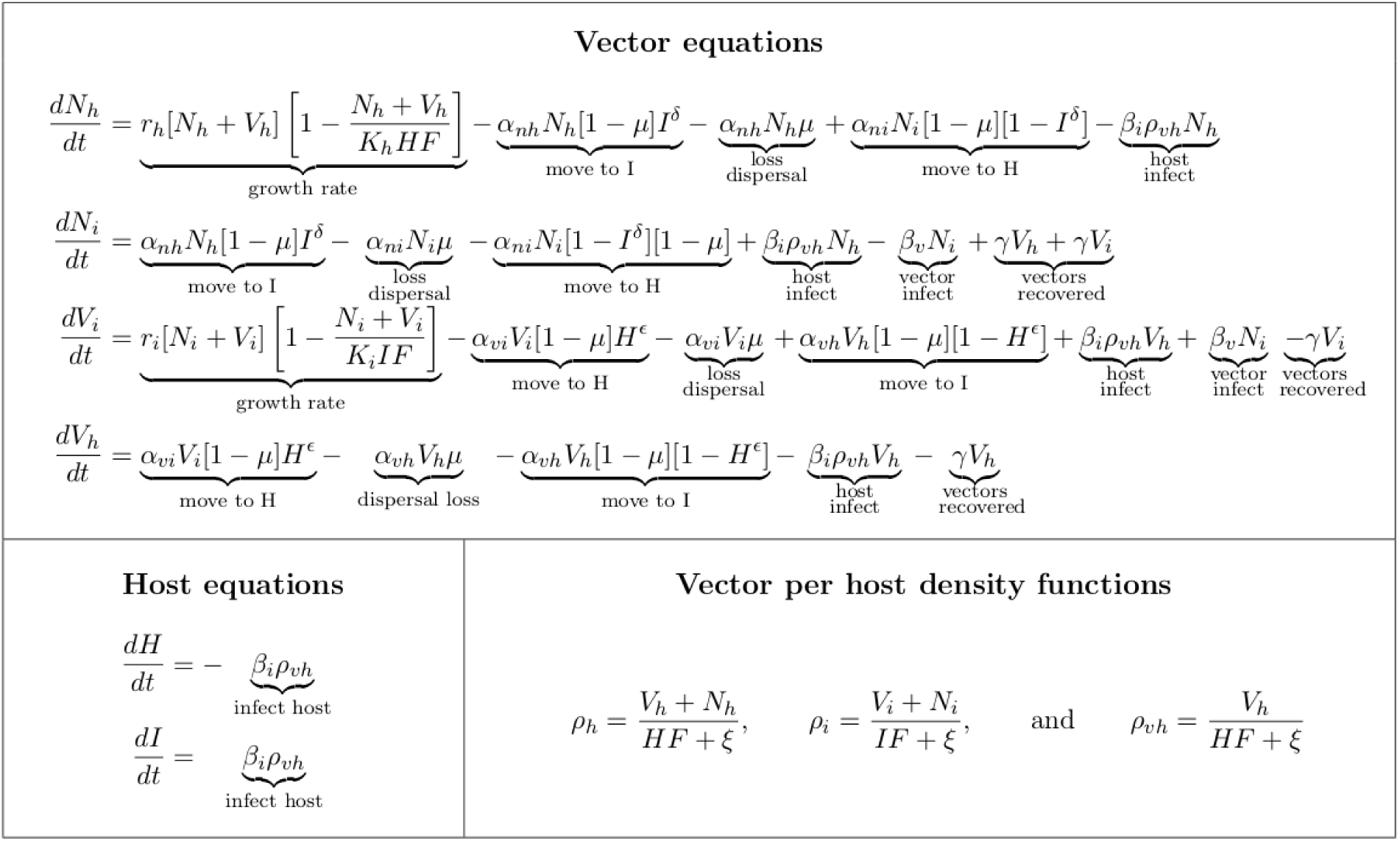
Model equations.

We ran simulations with parameter values as described in Table 1 using Matlab 2018 to explore the impact of virus effects on the time required for 80% of susceptible hosts to become infected (Model 1, discussed in main manuscript). We focused on the behavior (orientation preference [ε] and departure rate [*a*h]) and performance (growth rate [*r*i]) of non-viruliferous vectors in relation to virus-infected and non-infected hosts and assumed that viruliferous vector settling preference (ε) did not change following virus acquisition. We modeled this by setting the viruliferous vector settling preference above one (ε > 1) to simulate maintenance of orientation preference for infected hosts (if present based on CPNPr or NCNP parameter values) even after virus acquisition. The original model also explored the impact of changes in vector preference following acquisition of a virus, so-called *conditional vector preferences* (Roosien et al. 2013; Shaw et al. 2017). Our meta-analysis does not include publications that explore conditional vector preferences due to a lack of studies. However, to consider the possible influence of conditional preferences in the context of the parameter values derived from our database, we ran additional simulations by modifying the degree of viruliferous vector preference for settling on infected vs. healthy hosts. We modified ε to simulate loss of orientation preference after acquisition (chooses equally among infected and healthy hosts, ε = 1) (Model 2) and reversal of orientation preference after acquisition (ε = δ) (Model 3). These additional simulations are included in electronic supplementary file ESM_model outputs.xlsx.

We were also interested in exploring the relative influence of virus effects on vector responses *(r*i, *a*i, and δ) relative to intrinsic virus traits (combinations of βv, βi, and γ values corresponding to CPNPr and NCNP viruses). We combined trait values (βv, βi, and γ) for CPNPr viruses with vector response values (*r*i, *a*i, and δ) for NCNP viruses, and vice versa, then performed simulations as described above for initial and post-acquisition (conditional) vector preferences. Simulations for the main model are presented in the results and simulations for Model 2 and Model 3 in electronic supplementary file ESM_model outputs.xlsx.

## Results

### Plant virus infection effects on vector orientation preference, settling/feeding behavior, and performance

Plant virus infection had significant positive effects on vector orientation preference (figure 1a; ESM_meta-analysis.docx, table S2a), vector settling/feeding behaviors (figure 1b; ESM_meta-analysis.docx, table S2b) and vector performance (figure 1c; ESM_meta-analysis.docx, table S2c). These results were robust to publication bias (fail-safe N, ESM_meta-analysis.docx, Notes S4).

### Analysis based on virus traits: Infection location in plant hosts

The effects of plant virus infection on vector orientation preference vary depending on the location from which the virus must be acquired from and/or transmitted into the host plant (Q_M_ = 31.29, *p* < .0001; ESM_meta-analysis.docx, table S1). Plants infected by phloem-limited (PL) viruses become more attractive to vectors than non-infected plants, whereas plants infected with non-phloem-limited (NPL) viruses do not become more attractive to vectors than non-infected plants (figure 1a; ESM_meta-analysis.docx, table S2a). The effects of plant virus infection on vector settling and feeding behaviors also depend on the virus infection location in the host (Q_M_ = 26.51, *p* < .0001; ESM_meta-analysis.docx, table S1). Plant infection by PL viruses leads to greater rates of vector settling and feeding behaviors associated with host acceptance, whereas NPL virus infections do not alter settling/feeding behaviors (figure 1b; ESM_meta-analysis.docx, table S2b). The stronger effect of PL viruses on vector settling/feeding behaviors is consistent with results for vector performance (Q_M_ = 36.22, *p* < .0001; ESM_meta-analysis.docx, table S1). Plants infected by a PL virus support increased vector performance compared to non-infected plants, whereas plants infected with NPL viruses do not support significantly enhanced vector performance relative to non-infected plants (figure 1c; ESM_meta-analysis.docx, table S2c).

### Analysis based on virus traits: Retention mechanism in the vector

The effects of plant virus infection on vector orientation preference vary depending on how the virus is retained within the vector (Q_M_ = 93.92, *p* < .0001; ESM_meta-analysis.docx, table S1). Plants infected by circulative-persistent non-propagative (CPNPr) viruses were more attractive to vectors than non-infected plants, but enhanced vector attraction was not seen for any other type of virus (figure 2a; ESM_meta-analysis.docx, table S2a). CPNPr virus infection also enhanced host attractiveness to vectors relative to non-infected hosts to a greater extent than both non-circulative semi-persistent (NCSP) viruses and non-circulative non-persistent (NCNP) viruses (figure 2a). The effects of plant virus infection on vector settling and feeding behavior also vary depending on virus retention mechanism (Q_M_ = 37.60, *p* < .0001; ESM_meta-analysis.docx, table S1). Plants infected by circulative-persistent propagative (CPPr) viruses, CPNPr viruses, or NCSP viruses experience phenotypic shifts that encourage greater vector settling/feeding behaviors relative to non-infected plants, but plants infected by NCNP viruses experience similar rates of vector settling and ease of feeding relative to non-infected hosts (figure 2a; ESM_meta-analysis.docx, table S2b). Additionally, vector settling/feeding propensity for infected relative to non-infected hosts is significantly greater for CPNPr and NCSP viruses compared to NCNP viruses (figure 2b). For vector performance, again virus retention mechanism is a significant predictor of variation (Q_M_ = 41.98, *p* < .0001; ESM_meta-analysis.docx, table S1). However, in this case, only infection by CPNPr viruses increases vector performance on infected hosts over non-infected hosts; infections by viruses with the other three retention mechanisms do not increase vector performance on infected hosts relative to non-infected hosts (figure 2c; ESM_meta-analysis.docx, table S2c).

**Figure 2.**
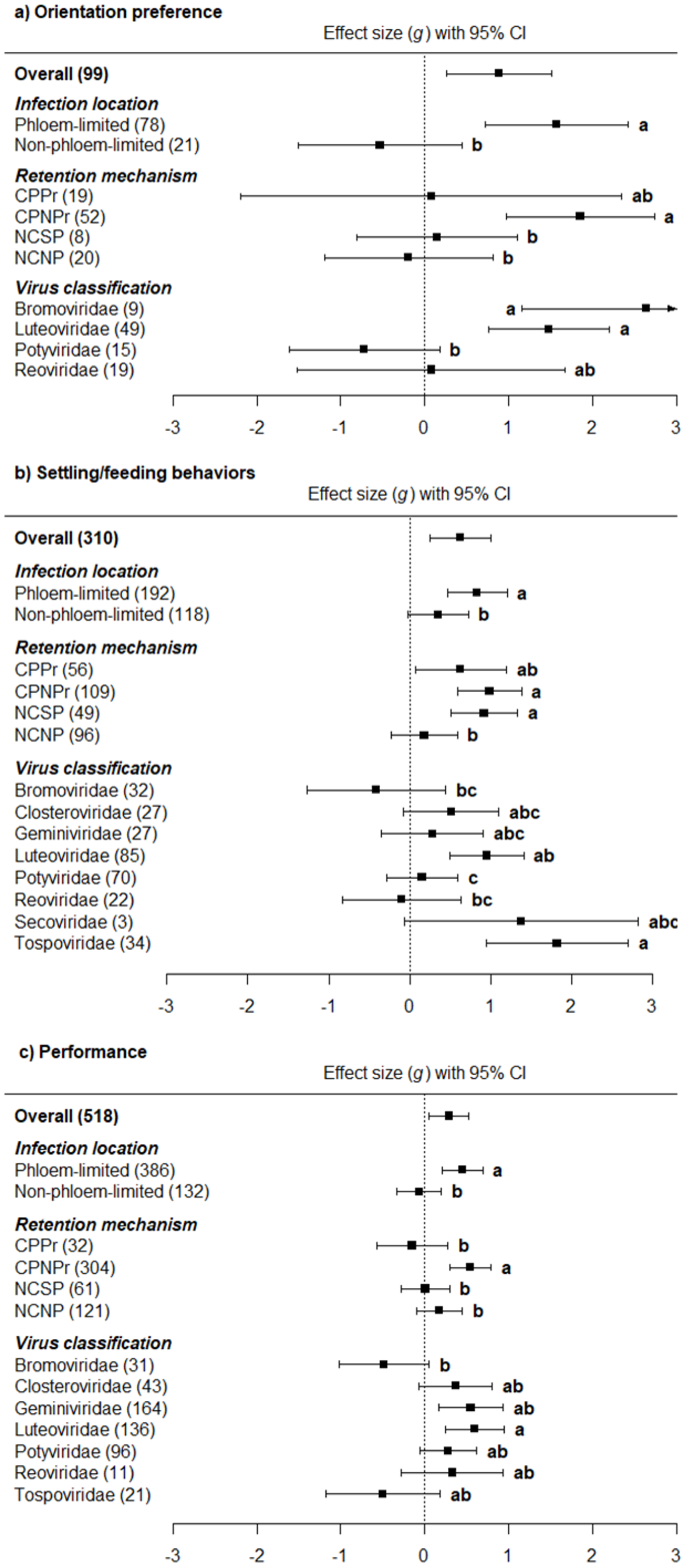
Effect size (Hedges’ *g*) estimates and 95% CIs showing the effects of plant infection on vector a) attraction, b) settling/feeding behaviors and c) performance. Negative values of *d* indicate a negative effect of plant infection on vector behaviors or performance. Arrows drawn at the ends of error bars indicate 95% CIs for Hedges’ *g* that are outside the scale of the plotting region. Numbers in brackets indicate the number of studies used to inform each estimate (see ESM_meta-analysis.docx, table S2a, b and c, for complete sample size information).

### Analysis by phylogeny: Virus classification

The effects of plant infection on vector orientation preference vary depending on virus family (Q_M_ = 112.24, *p* < .0001; ESM_meta-analysis.docx, table S1). Only four virus families had sufficient representation within the literature to be included in the analysis for orientation preference (Figure 2a). Of these families, infections by Bromoviridae and Luteoviridae induce host phenotypes that are more attractive than those of non-infected plants, but plants infected by Potyviridae and Reoviridae are not differentially attractive (figure 2a; ESM_meta-analysis.docx, table S2a). Within studies focusing on vector settling and feeding behavior, eight virus families had sufficient representation for inclusion in the analysis (Figure 2b). The effects of plant infection on vector settling and feeding behaviors vary depending on virus family (Q_M_ = 73.54, *p* < .0001; ESM_meta-analysis.docx, table S1). Infections by Luteoviridae and Tospoviridae increase vector settling and/or ease of feeding on hosts relative to non-infected plants, but infections by Bromoviridae, Closteroviridae, Geminiviridae, Potyviridae, Reoviridae and Secoviridae do not significantly influence vector settling or ease of feeding (figure 2b; ESM_meta-analysis.docx, table S2b). For the vector performance metric, seven virus families were sufficiently well represented for inclusion in the analysis (Figure 2c). As for orientation preference and settling/feeding behaviors, the effects of plant infection status on vector performance varied depending on the virus family (Q_M_ = 70.39, *p* < .0001; ESM_meta-analysis.docx, table S1). Infections by Geminiviridae and Luteoviridae increased vector performance on infected hosts relative to non-infected plants, but infections by representatives of other virus families did not affect vector performance (figure 2c; ESM_meta-analysis.docx, table S2c).

### Modeling implications for virus spread

Figure 3 covers simulations where phloem-limited, circulative-persistent, non-propagative (PL-CPNPr) virus traits (βv and γ) are combined with PL-CPNPr virus parameter values and ranges for (*r, a*, δ and ε), and non-phloem-limited, non-circulative, non-persistent (NPL-NCNP) virus traits are combined with NPL-NCNP virus parameter values and ranges for (*r, a*, δ and ε). Simulation outputs suggest that changes in host phenotype that enhance vector performance, orientation preference, and settling/feeding have little effect on PL-CPNPr virus spread (as time to 80% host infection), but relatively large effects on NPL-NCNP virus spread (Fig. 3, Table 3).

**Table 3.**
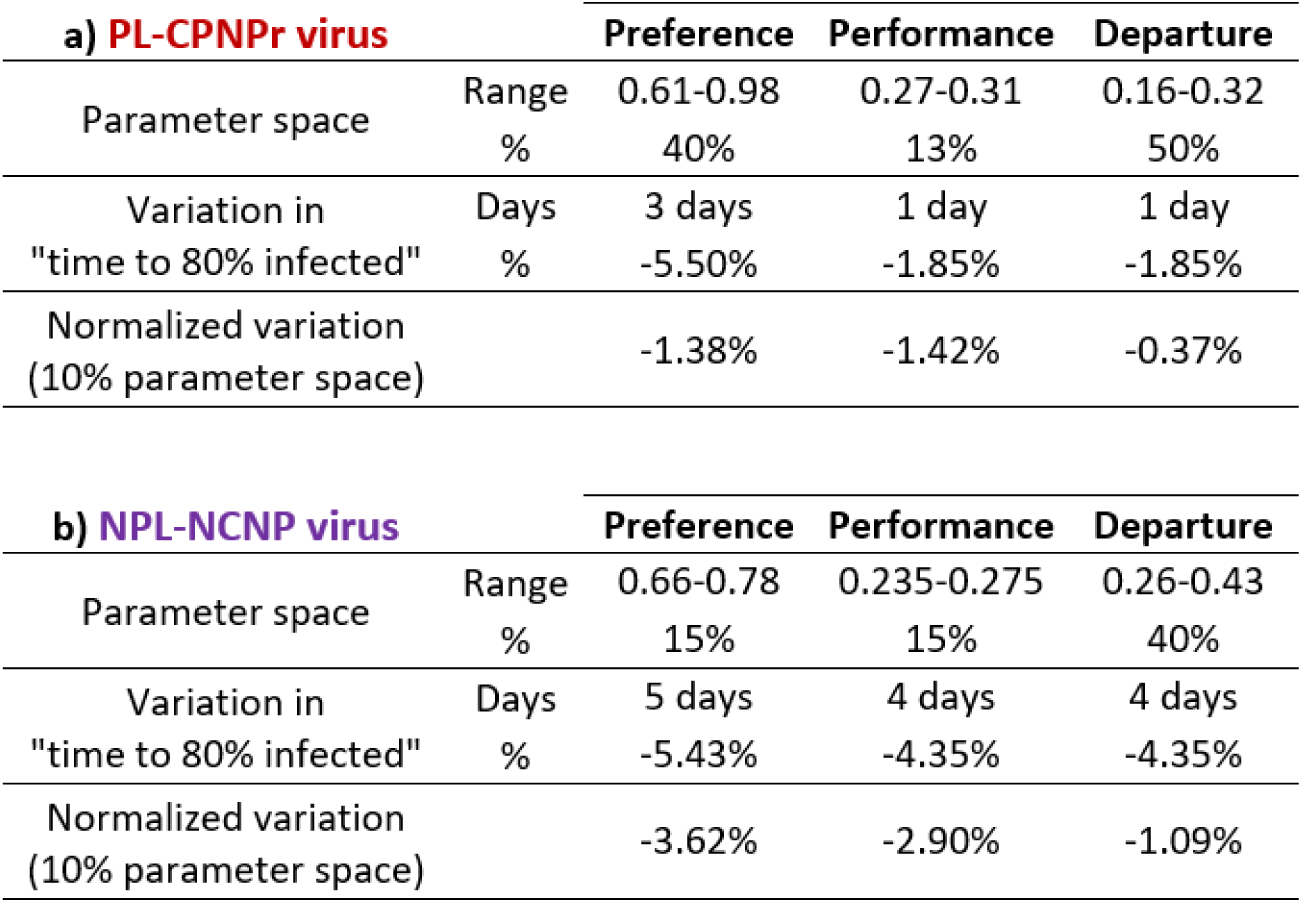
Simulation outputs for a) Phloem-limited, circulative-persistent, non-propagative (PL-CPNPr) virus traits (βv and γ) combined with PL-CPNPr virus parameter values and ranges (*r, a*, δ and ε) and b) Non-phloem-limited, non-circulative, non-persistent (NPL-NCNP) virus traits combined with NPL-NCNP virus parameter values and ranges for (*r, a*, δ and ε).

| a) PL-CPNPr virus |  | Preference | Performance | Departure |
| --- | --- | --- | --- | --- |
| Parameter space | Range | 0.61-0.98 | 0.27-0.31 | 0.16-0.32 |
|  | % | 40% | 13% | 50% |
| Variation in<br>"time to 80% infected" | Days | 3 days | 1 day | 1 day |
|  | % | -5.50% | -1.85% | -1.85% |
| Normalized variation<br>(10% parameter space) |  | -1.38% | -1.42% | -0.37% |

| b) NPL-NCNP virus |  | Preference | Performance | Departure |
| --- | --- | --- | --- | --- |
| Parameter space | Range | 0.66-0.78 | 0.235-0.275 | 0.26-0.43 |
|  | % | 15% | 15% | 40% |
| Variation in<br>"time to 80% infected" | Days | 5 days | 4 days | 4 days |
|  | % | -5.43% | -4.35% | -4.35% |
| Normalized variation<br>(10% parameter space) |  | -3.62% | -2.90% | -1.09% |

**Figure 3:**
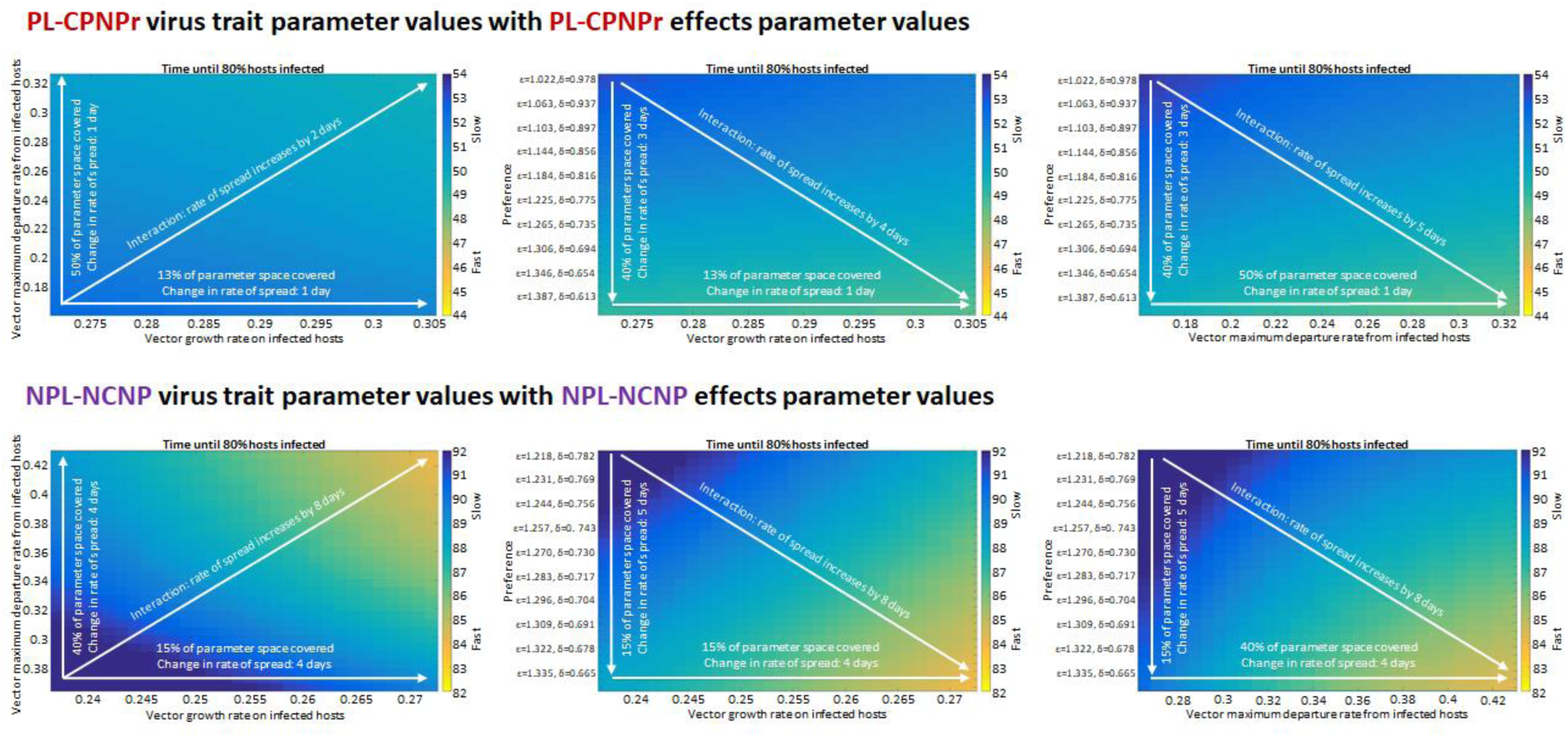
Main model simulation outputs for the two virus trait categories (PL-CPNPr and NPL-NCNP) with their corresponding virus effects values (*r, a*, δ and ε). For this model, virus effects on vector preferences do not change following virion acquisition. Overlays on simulation heatmaps describe the percentage of total available parameter space covered along each axis and the estimated maximum change in virus spread from one end of the axis to the other. Decreases in time to 80% host infection are indicated by arrow directionality. Simulation outputs are also represented quantitatively in Table 3.

In simulations, vector maximum departure rate parameter values for PL-CPNPr viruses covered 50% of the available parameter space (0.16-0.32), but the maximum departure rate within this window only resulted in a 1.85% reduction (approx. 1 day) in time to 80% infection (Fig. 3, Table 3). In contrast, for NPL-NCNP viruses, values for maximum departure rate covered 40% of parameter space (0.26-0.43), with the maximum departure rate within this window producing a 4.35% reduction (approx. 4 days) in time to 80% infection (Fig. 3, Table 3). Parameter values for vector growth rate on infected plants covered 13% of the parameter space for PL-CPNPr viruses (Table 3), and 15% of the parameter space for NPL-NCNP viruses (Table 3). Even though the proportion of space covered is similar, the maximum gain in rate of spread for PL-CPNPr viruses is one day, but NPL-NCNP virus spread occurs four days faster at the largest vector growth rate values (Fig. 3, Table 3). For both PL-CPNPr viruses and NPL-NCNP viruses, vector preference had the greatest effect on time to 80% infection. For a range of values covering 40% of parameter space, PL-CPNPr virus spread occurred three days faster at values corresponding to the maximum preference for infected hosts. However, for NPL-NCNP viruses, a range of values covering just 15% of parameter space reduced time to 80% infection by up to 5.43% (approx. 5 days) at values corresponding to maximum preference for infected hosts. If these results are normalized to the percent change for 10% of parameter space, it is apparent that across all virus effects categories, the same change in parameter values produces between two and three times the effect for NPL-NCNP viruses relative to PL-CPNPr viruses (Table 3).

To explore the relative influence of virus traits *vs*. virus effects on host phenotypes, we ran each simulation a second time using PL-CPNPr virus trait values (β and γ) with NPL-NCNP virus values for virus effects (*r, a*, δ, and ε), and vice versa. Substituting NPL-NCNP virus effects values (*r, a*, δ, and ε) in a model maintaining PL-CPNPr virus trait values (β and γ) had little effect on PL-CPNPr virus spread (Fig. 4). However, for NPL-NCNP trait values (β and γ) paired with PL-CPNPr virus values for virus effects (*r, a*, δ, and ε) this was not the case. First, substituting PL-CPNPr virus values for vector growth rate on infected plants and vector orientation preference resulted in slower overall rates of virus spread relative to use of NPL-NCNP virus parameters (*r, a*, δ, and ε) (Fig. 4). There were also slightly positive effects on virus spread due to substitution of PL-CPNPr virus values for vector maximum departure rate (*a*) (Fig. 4).

**Figure 4:**
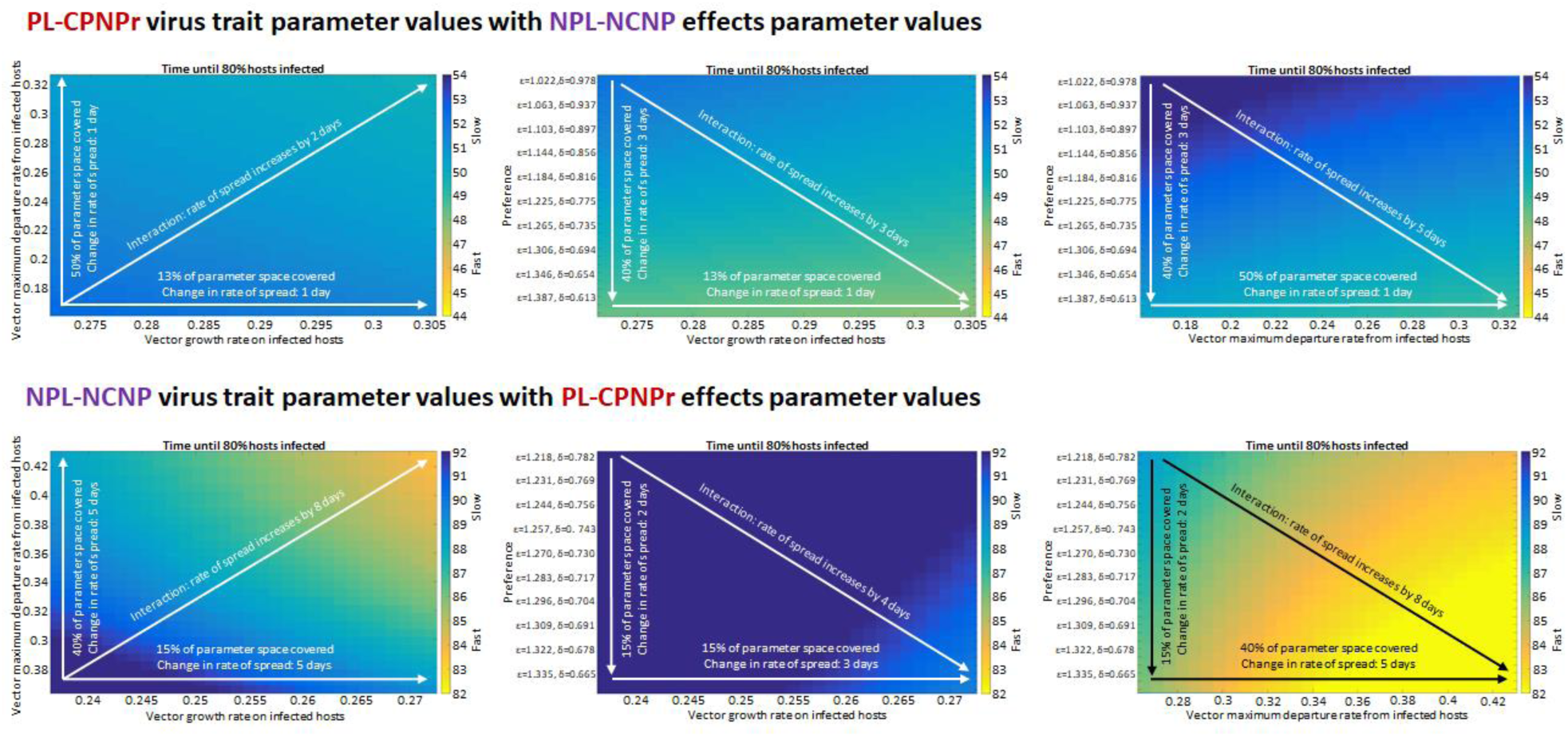
Main model simulation outputs for the two virus trait categories (PL-CPNPr and NPL-NCNP) with virus effects values (*r, a*, δ and ε) corresponding to the opposite virus trait category. For this model, virus effects on vector preferences do not change following virion acquisition. Overlays on simulation heatmaps describe the percentage of total available parameter space covered along each axis and the estimated maximum change in virus spread from one end of the axis to the other. Decreases in time to 80% host infection are indicated by arrow directionality. Simulation outputs are also represented quantitatively in Table 4.

Beyond virus effects parameters, the model simulations reveal the influence of virus traits on the overall rate of virus spread in a host population. In simulations using PL-CPNPr virus traits, with vectors maintaining viruliferous status (γ = 0) and having a dispersal loss (mortality, μ) value of 53.67% +/- 21.39, the minimum time to 80% of hosts becoming infected is 44 days and the maximum is 54 days. In contrast, for the NPL-NCNP virus trait values (same dispersal loss value, but γ = 1 to approximate daily loss of acquired virus) the minimum time to 80% infection is 82 days, with a maximum of 92 days. Thus, according to our simulations, the viruses with NPL-NCNP traits take twice as long to infect 80% of the host population relative to viruses with PL-CPNPr traits in a context that includes an aphid vector capable of colonizing the host.

**Table 4.**
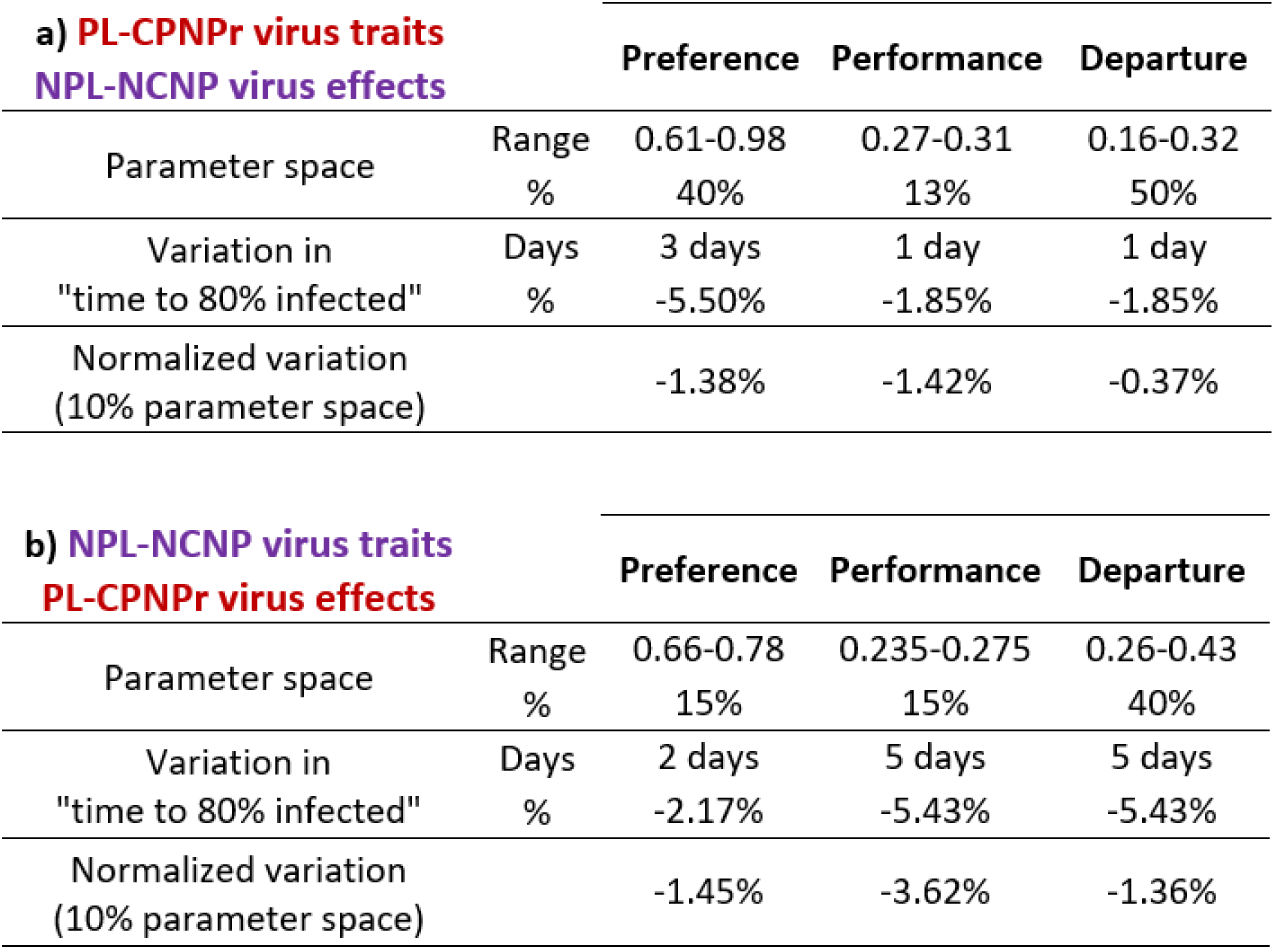
Simulation outputs for a) Phloem-limited, circulative-persistent, non-propagative (PL-CPNPr) virus traits (βv and γ) combined with NPL-NCNP virus parameter values and ranges (*r, a*, δ and ε) and b) Non-phloem-limited, non-circulative, non-persistent (NPL-NCNP) virus traits combined with PL-CPNPr virus parameter values and ranges for (*r, a*, δ and ε).

Inclusion of conditional vector preferences had little effect on the rate of PL-CPNPr virus spread regardless of whether virus effects parameter values were those of PL-CPNPr viruses or NPL-NCNP viruses. NPL-NCNP virus spread was slightly faster when conditional vector preferences were included, either as a loss of preference following virus acquisition (Model 2) or as a switch in preference (Model 3). This effect was only evident when NPL-NCNP virus trait parameter values were paired with NPL-NCNP virus effect parameter values. Simulations and tabular summaries for Model 2 and Model 3 are available in the electronic supplementary material, file ESM_model outputs.xlsx.

## Discussion

### Quantitative synthesis and support for predictions

We predicted that enhanced attractiveness to vectors should be a host phenotypic change common to all virus trait groups, but enhanced palatability and plant quality should be significantly more apparent for trait groups that require long-term feeding for virus acquisition (PL viruses), which includes most representatives of viruses retained in vectors for long periods (CPPr, CPNPr, and NCSP) (Fig. 1). Our synthesis provides support for these predictions, as we saw significant differences based on virus traits for multiple response variables. For example, when considering site of acquisition/inoculation as a factor, PL virus effects conformed to all predictions, eliciting enhancements in vector orientation preferences, settling/feeding preferences, and performance on infected hosts (Fig. 2). NPL viruses also conformed to predictions for settling/feeding and performance response variables; effects on vector preference and performance did not deviate from zero, and were significantly different from preference/performance values for PL viruses in the expected direction (Fig. 2). However, NPL virus effects were not consistent with our prediction that all virus trait groups should enhance plant attractiveness to vectors (Fig. 1, Fig. 2a). On average, NPL viruses have neutral, but not detrimental, effects on vector attraction (Fig. 2a). While this may not be an obvious case of vector manipulation by plant viruses, a neutral result is still consistent with an adaptive explanation for virus effects because we expect selection to disfavor virus genotypes that *reduce* opportunities for transmission, but favor virus genotypes with *neutral to positive* effects on transmission (Anderson *et al*. 1992; Poulin 2010). This hypothesis is also supported by the meta-analysis output: we did not detect a single instance of an effect size confidence interval deviating from zero in a *negative* direction, suggesting that there is selection against virus genotypes that elicit phenotypes with strongly negative effects on vector-host interactions.

The vector settling/feeding response variable is arguably the most important in our study given that it is the stage in the vector-host interaction where the viruliferous status (i.e., virus acquisition) of the vector is determined (Fereres & Moreno 2009). Thus, even if contact with infected hosts is not influenced by infection status (Fig. 2a) and there are no strong effects of infection on vector performance (Fig. 2c), selection should favor virus adaptations that ensure vectors engage in probing and feeding behaviors required for efficient virus acquisition and retention. We detected indirect evidence of adaptations producing positive effects on vector settling/feeding for all of the virus trait categories expected to benefit from them, even when we analyzed the dataset with virus retention mechanism as the factor. Viruses with CPPr, CPNPr, and NCSP retention mechanisms (nearly all of which are PL) significantly enhanced vector settling and feeding preferences for infected hosts (Fig. 2b). We also detected neutral effects for the virus categories expected to experience reductions in transmission when infection enhances palatability, settling, and sustained feeding prior to dispersal (NPL viruses with the NCNP transmission mechanism) (Fig. 2b). Thus, for the settling/feeding response variable, which we argue is the most critical of the three for virus fitness, the prediction of convergence based on shared virus traits is strongly supported.

We hypothesized that virus effects on vector settling/feeding behavior would be congruent with effects on vector performance, but this was not the case (Fig. 2b, c). Instead, viruses have mostly neutral effects. Although PL viruses enhanced performance overall, this appears to be driven by the effects of CPNPr viruses (which had more observations than any other group) (Fig. 2c). CPPr and NCSP viruses, as well as respective virus families containing taxa with these retention mechanisms, do not strongly influence vector performance. This may reflect limitations imposed by other traits inherent to viruses within each retention mechanism category. For example, CPPr viruses also use the vector as a host for replication, so a neutral effect may still be interpreted as evidence of adaptation, as it could indicate that the actively replicating virus does not have strongly pathological effects on the vector, or that phenotypic changes in the host help to counteract slight pathological effects (Belliure *et al*. 2005). Direct effects of viruses are changes in vector behavior or performance that occur as a result of acquiring and retaining virions, and are apparent in some CPPr and CPNPr pathosystems (Stafford *et al*. 2011; Ingwell *et al*. 2012; Moreno-Delafuente *et al*. 2013; Roosien *et al*. 2013; Rajabaskar *et al*. 2014; Mauck *et al*. 2018a, 2019). However, there are not sufficient data to explore the interplay of direct and indirect virus effects on vectors through meta-analysis approaches. Overall, the meta-analysis highlights the need for additional studies across more diverse pathosystems while providing the first quantitative evidence supporting an adaptive explanation for putative instances of vector manipulation by plant viruses.

### Comparing the quantitative synthesis to a model

We leveraged data assembled for the meta-analysis to assess how virus-induced manipulation of hosts and vectors may affect the rate of transmission in theoretical simulations by modifying a published model (Shaw *et al*. 2017) that explores the spread of viruses with divergent traits (aphid-transmitted PL-CPNPr viruses and NPL-NCNP viruses). Simulation outputs indicate that the range of parameter values derived from the meta-analysis database were sufficient to produce positive effects on virus spread for both trait groups (Fig. 3). However, the magnitude of these effects differed depending on the trait group being examined. Across all virus effects categories, the same change in virus effects parameter values produced between two and three times the effect for NPL-NCNP viruses relative to PL-CPNPr viruses (Table 3). This difference suggests that PL-CPNPr viruses (and perhaps PL viruses generally) may be under more intense selection pressure to elicit stronger effects on host phenotypes and vector behavior in order to experience fitness benefits of manipulation. In contrast, our results suggest that NPL-NCNP viruses may experience less intense selection pressure to manipulate hosts and vectors because even small effects can produce significant changes in the rate of spread. These results are strongly congruent with meta-analysis outputs: PL virus effects were significantly different from zero in the positive direction and significantly different from NPL virus effects (which were uniformly neutral) across all three response variables.

Simulations with virus trait parameters swapped with virus effects parameters (Fig. 4, Table 4) reveal that NPL-NCNP viruses may incur costs for eliciting effects that are predicted to be adaptive for PL-CPNPr viruses. Combining NPL-NCNP virus trait parameters with PL-CPNPr effects parameters (Fig. 4) slowed virus spread relative to the original virus trait-virus effects combinations (Fig. 3). Simulations reveal that this is partially due to the influence of parameter values that we predicted to be beneficial for PL-CPNPr virus spread (higher vector growth rate on infected hosts), which is consistent with our initial predictions (Fig. 1). However, NPL-NCNP virus spread was also slowed by enhancing vector orientation preference for infected hosts (Fig. 4), which is at odds with our initial prediction that enhancing vector attraction to infected hosts would be generally beneficial for all virus trait groups (Fig. 1). Thus, the model simulations indicate that neutral effects of NPL viruses on vector attraction to infected hosts (Fig. 2a) may actually lead to faster NPL virus spread (Fig. 3) relative to a situation where vectors are strongly attracted to NPL-infected hosts (Fig. 4). Theoretical work on plant virus manipulation of hosts and vectors (McElhany *et al*. 1995; Sisterson 2008) and parasite manipulation literature generally (Lefèvre *et al*. 2006; Poulin 2010; Lafferty & Shaw 2013; Heil 2016) both support the idea that vector attraction to infected hosts is generally beneficial for parasite fitness. But interpretation of our meta-analysis and model simulations together indicates that relative benefits of vector preferences for infected hosts strongly depend on additional transmission mechanism traits that determine which vector behaviors are required for virus acquisition and transmission.

### Limitations, ecological dimensions, and future directions

The meta-analysis and model both focus on pathosystems consisting of one host species, one virus isolate, and a single colonizing vector species. This simplification is necessary to make empirical and theoretical exercises logistically practical and congruent. However, elimination of additional ecological dimensions will influence interpretations, including those of the present study. A key example is the difference in time to 80% host infection between virus trait groups in our model simulations. PL-CPNPr viruses infected 80% of the host population in roughly half the time it took for NPL-NCNP viruses to reach the same infection level (44-54 days *vs*. 82-92 days) (Figs. 3 and 4). From this result, we might conclude that acquisition from the phloem and indefinite retention in the vector confer large advantages over an NPL-NCNP lifestyle; certainly much larger than virus effects on host phenotype and vector behavior, which, at maximum, increases the rate of virus spread by 9-10 days. However, NPL-NCNP viruses are both common and widely successful; at least 40% of characterized viruses are NPL-NCNP viruses (Hogenhout *et al*. 2008) and globally, NPL viruses are major threats to agricultural production. From this we can conclude that our model (Shaw et al. 2017) and others with similar features (McElhany *et al*. 1995; Sisterson 2008; Roosien *et al*. 2013; Shaw *et al*. 2019) do not fully represent the range and variety of transmission opportunities available for NPL-NCNP viruses.

We explored this possibility by quantifying the number of vector species capable of transmitting each virus included in the meta-analysis. PL viruses have intimate, co-evolved interactions with vectors that are capable of feeding from the phloem of the virus’s host plants (Hogenhout *et al*. 2008). Consistent with this, we found that >80% of the PL viruses in our meta-analysis database are transmitted by three or fewer vector species (Fig. S1, electronic supplementary material, file ESM_vector database.docx). Thus, we hypothesized that PL viruses generate transmission opportunities by interacting with a select few species that colonize hosts long enough to acquire virions. This is the exact scenario represented by most of the empirical studies in our meta-analysis database and our mathematical models, which simulated the transmission of a virus by a single vector capable of feeding and surviving on the host for at least 24 hours (Shaw *et al*. 2017). However, this scenario is the opposite of the ecological reality for NPL-NCNP viruses, which are acquired by rapid probing, but not phloem-feeding, and are therefore transmissible by vectors for which the infected host species is not suitable (Ng & Falk 2006; Fereres 2016). In our models, reliance on a single colonizing vector significantly slowed NPL-NCNP virus spread and seemed to suggest that the NPL-NCNP lifestyle is a major drawback. In reality, nearly 70% of the NPL viruses included in the meta-analysis are transmissible by more than 20 vector species, and all are transmissible by 5 or more vector species (Fig. S1, electronic supplementary material, file ESM_vector database.docx). Thus, while PL viruses rely on a select few colonizing species, most NPL viruses seem to generate transmission opportunities by interacting with a large number of species that need not colonize the host, or feed on the phloem, to acquire virions (Bosque-Pérez & Eigenbrode 2011).

This ecological reality is rarely considered in empirical studies and has not been included in any models exploring vector manipulation to date (McElhany *et al*. 1995; Madden *et al*. 2000; Sisterson 2008; Roosien *et al*. 2013; Shaw *et al*. 2017, 2019; Mauck *et al*. 2018). But we can speculate that the difference in transmission opportunities between PL and NPL viruses will significantly influence the relative fitness benefits of host manipulation for these virus trait groups. Plant viruses have limited coding capacity and fixation of a mutation is strongly dependent on a lack of epistatic interactions with other sites in the genome and a lack of pleiotropic effects (e.g., reduced replication rate in the host or a modified host range) (Bedhomme *et al*. 2012; Betancourt *et al*. 2013; García-Arenal & Fraile 2013; Elena 2016). For NPL viruses, pleiotropic effects may manifest as reduced opportunities for transmission; mutations that facilitate manipulation of one vector species might compromise transmission by several other vector species. However, neutral effects on host phenotypes (as observed in our meta-analysis) would increase the likelihood that most competent vector species will visit some infected plants and engage in behaviors required for efficient NPL virus transmission (rapid probing and dispersal) due to host plant incompatibility, and regardless of infection status (Rydén *et al*. 1983; Sigvald 1989; Angelella *et al*. 2015; Mondal *et al*. 2016). In contrast, PL viruses that rely on a select few vector species for transmission may experience substantial gains by increasing the probability of transmission-conducive contacts with these species (Fig. S1, electronic supplementary material, file ESM_vector database.docx).

Transmission opportunities is just one ecological dimension influencing virus evolution, and there are certainly more factors that could shape selection for or against manipulative traits, including, but not limited to, host diversity, abiotic conditions, vector natural enemies, and effects of host phenotype manipulations on resistance against non-vector pests and other pathogens (Jeger *et al*. 2011; Kersch-Becker & Thaler 2013; Mauck *et al*. 2014b, 2015; Davis *et al*. 2015; Chesnais *et al*. 2019). The present synthesis provides an important milestone by quantifying the magnitude and adaptive significance of virus effects on vectors while also emphasizing the need for additional research in different contexts and pathosystems. For example, PL-CPNPr viruses in the *Luteoviridae* are overrepresented in the meta-analysis (Fig. 2), while some other virus groups consist of only a few observations. Lack of data on certain virus groups may reflect their relative economic importance, quarantine status, or tractability for laboratory studies. Incorporating new pathosystems and ecological dimensions into future empirical and theoretical work is therefore necessary to understand the frequency and relevance of virus manipulation in real-world scenarios. The number of researchers studying this topic is growing (Mauck *et al*. 2018, 2019), as is interest in finding ways to manage manipulative effects of viruses in agricultural contexts (Bak *et al*. 2019) and understanding their importance in wild systems (Alexander *et al*. 2014). Tackling these ambitious research directions requires integrative approaches informed by virology, ecology, entomology, and plant biology. We hope our study will stimulate the research community to develop and test hypotheses that provide a more complete understanding of host and vector manipulation by plant viruses in ecological contexts that include consideration of virus traits.

## Supporting information

Collected supplementary files

