## Supplementary material for "Evidence of adaptive host and vector manipulation by plant viruses revealed through combined meta-analysis and modeling approaches": Collected supplementary files

Start of content for file ESM_meta-analysis.docx

**Electronic Supplementary Materials: “Meta-analysis”**

1. **Additional methods details**
    a. Database assembly

Articles returned in literature searches (see main text for search terms) were first screened for inclusion by scanning the titles and abstracts for relevance. A total of ~300 studies were retained for full-text review and assessed for inclusion using the following criteria: 1) The study must include data on vector attraction/orientation behavior, feeding/settling behavior or performance for both an infected and its healthy relative; 2) The study must report means, sample sizes, and variances of the focal data; 3) The study must report data on a functional vector-virus combination (i.e. non-vector examples were excluded).

Our final database included 126 unique studies that compared vector behaviors and performances on 69 virus*plant combinations (59 viruses belonging to 11 families infecting plants of 15 different families) and their healthy relatives. Results are based on an inconstant number of non-infected/infected plant comparisons made across a diverse group of hosts: vector attraction data included 99 comparisons taken from 25 studies, vector settling/feeding data included 310 comparisons obtained from 67 studies and vector performance data included 518 comparisons taken from 81 studies (see table S2 a, b and c, for complete sample size information).

b. Data selection

We often recorded multiple comparisons and types of response variables from a single study. Common reasons for replication within a study were that data were reported for multiple crop accessions and/or multiple viruses. In these cases, we calculated an effect size for each possible plant x virus x vector combination separately. In a few studies, we also recorded data separately for different vector life stages or sexes that could be responding differently to plant traits. In addition, a few studies reported data for multiple plant parts, and we recorded each of these separately.

Another important source of non-independence of data was that many studies reported multiple types of relevant response variables. We considered three broad categories of data, vector attraction/orientation behavior, feeding/settling behavior and vector performance.

Vector orientation preference data included three types of response variables: emigration and immigration rates, landing preferences and orientation preferences (in a Y-tube olfactometer or close proximity to plant headspace). Vector feeding/settling data also included three types of response variables: settling / arrestment, retention / dispersion, and two selected EPG (i.e. electrical penetration graph technique) variables, “E2” indicating a phloem sap ingestion (usually synonym to a sustained settlement/feeding behavior) and “pd/C” indicating pathway phase and/or intracellular stylet penetrations. Vector performance data included seven types of response variables: population growth rates, individual fecundity, development time, longevity, survival, weight and size.

When studies included data from more than one category of response, we recorded each response separately, e.g. adult survival and pupal weight. If studies included more than one reported measurement within a single type of response category, we used the following criteria for selection of data: 1) For studies that included repeated measures during a single experiment (e.g. retention behavior after 30 minutes, 1 hour, 2 hours and 24 hours), we only reported two measurements corresponding to “initial” and “long-term” decisions; 2) For studies that included virus effects at different points in disease progression, we reported all the measurements (e.g. settling behavior at 15 days post-inoculation [dpi], 20 dpi, 25 dpi, 30 dpi, etc.).

#### 1. Supplementary Tables

**Table S1:** Test statistics and variance components for each model testing the effects of virus infection on a) vector orientation preference, b) vector settling/feeding preference and c) vector performance.

| Moderators | $Q_M$ [a] | Prob ($Q_M$) | $Q_E$ [b] | Prob ($Q_E$) | Variance Study (**$\sigma^2$**) [c] |
|---|---|---|---|---|---|
| **a) Orientation preference** | | | | | |
| None | - | - | 1046 | **<.0001** | 2.41 |
| Infection location | 31.29 | **<.0001** | 1023 | **<.0001** | 4.24 |
| Retention mechanism | 93.92 | **<.0001** | 905 | **<.0001** | 3.97 |
| Virus classification | 112.24 | **<.0001** | 761 | **<.0001** | 1.94 |
| (Domestication) | 0.05 | 0.8179 | 1036 | **<.0001** | 2.48 |
| (Year) | 0.07 | 0.7837 | 1041 | **<.0001** | 2.63 |
| **b) Settling/Feeding preference** | | | | | |
| None | - | - | 3685 | **<.0001** | 2.27 |
| Infection location | 26.51 | **<.0001** | 3567 | **<.0001** | 2.10 |
| Retention mechanism | 37.60 | **<.0001** | 3368 | **<.0001** | 2.11 |
| Virus classification | 73.54 | **<.0001** | 3067 | **<.0001** | 1.50 |
| (Domestication) | 0.29 | 0.5904 | 3685 | **<.0001** | 2.27 |
| (Year) | 0.03 | 0.8581 | 3674 | **<.0001** | 2.32 |
| **c) Performance** | | | | | |
| None | - | - | 5719 | **<.0001** | 1.15 |
| Infection location | 36.22 | **<.0001** | 5717 | **<.0001** | 1.15 |
| Retention mechanism | 41.98 | **<.0001** | 5704 | **<.0001** | 1.04 |
| Virus classification | 70.39 | **<.0001** | 5436 | **<.0001** | 1.02 |
| (Domestication) | 13.21 | **0.0003** | 5718 | **<.0001** | 1.21 |
| (Year) | 0.04 | 0.8400 | 5689 | **<.0001** | 1.16 |

[a] Omnibus test of model coefficients: significant values indicate that the moderator has an overall effect on the magnitude of the virus effect size

[b] Test for residual heterogeneity: significant values suggest the existence of additional explanatory variables not included in the model

[c] indicates the amount of variance attributable to study effect (see text for details on model specification)

**Table S2a:** Model estimated mean effect sizes, 95% confidence intervals, and sample sizes for each level of the various moderators influencing the effect of virus infection on **vector orientation preference**

| **Moderators** | **Estimate** | **CI.lb** | **CI.ub** | **pval** | $N_{total}$[a] | $N_{st}$[b] |
|---|---|---|---|---|---|---|
| None | 0.88 | 0.25 | 1.50 | **0.0060** | 99 | 25 |
| Infection location | | | | | | |
| Phloem-limited | 1.57 | 0.72 | 2.42 | **0.0003** | 78 | 18 |
| Non-phloem-limited | -0.53 | -1.50 | 0.44 | 0.2813 | 21 | 8 |
| Retention mechanism | | | | | | |
| CPP | 0.07 | -2.19 | 2.34 | 0.9482 | 19 | 3 |
| CPNP | 1.85 | 0.97 | 2.73 | **<.0001** | 52 | 14 |
| NCSP | 0.14 | -0.81 | 1.09 | 0.7689 | 8 | 3 |
| NCNP | -0.19 | -1.19 | 0.80 | 0.7048 | 20 | 7 |
| Virus classification | | | | | | |
| Bromoviridae | 2.64 | 1.15 | 4.13 | **0.0005** | 9 | 4 |
| Closteroviridae | -1.14 | -3.89 | 1.60 | 0.4146 | 3 | 1 |
| Geminiviridae | 0.52 | -2.22 | 3.26 | 0.7075 | 3 | 1 |
| Luteoviridae | 1.47 | 0.75 | 2.20 | **<.0001** | 49 | 13 |
| Potyviridae | -0.72 | -1.61 | 0.17 | 0.1135 | 15 | 4 |
| Reoviridae | 0.07 | -1.51 | 1.66 | 0.9270 | 19 | 3 |
| Secoviridae | -3.54 | -6.39 | -0.70 | **0.0146** | 1 | 1 |
| (Domestication) | | | | | | |
| Domesticated | 0.89 | 0.25 | 1.52 | **0.0063** | 97 | 25 |
| Wild | 0.64 | -1.47 | 2.75 | 0.5499 | 2 | 1 |

[a] the total number of infected/healthy comparisons

[b] the number of unique studies

**Table S2b:** Model estimated mean effect sizes, 95% confidence intervals, and sample sizes for each level of the various moderators influencing the effect of virus infection on **vector settling/feeding behavior**

| Moderators | Estimate | CI.lb | CI.ub | pval | $N_{total}$[a] | $N_{st}$[b] |
|---|---|---|---|---|---|---|
| None | 0.63 | 0.25 | 1.00 | **0.0010** | 310 | 67 |
| Infection location | | | | | | |
| Phloem-limited | 0.83 | 0.46 | 1.20 | **<.0001** | 192 | 41 |
| Non-phloem-limited | 0.35 | -0.02 | 0.73 | 0.0661 | 118 | 32 |
| Retention mechanism | | | | | | |
| CPP | 0.63 | 0.07 | 1.19 | **0.0284** | 56 | 13 |
| CPNP | 0.99 | 0.59 | 1.38 | **<.0001** | 109 | 24 |
| NCSP | 0.92 | 0.50 | 1.33 | **<.0001** | 49 | 13 |
| NCNP | 0.18 | -0.23 | 0.59 | 0.3851 | 96 | 26 |
| Virus classification | | | | | | |
| Bromoviridae | -0.41 | -1.26 | 0.44 | 0.3445 | 32 | 9 |
| Caulimoviridae | 0.74 | 0.21 | 1.27 | **0.0065** | 12 | 1 |
| Closteroviridae | 0.51 | -0.07 | 1.10 | 0.0875 | 27 | 7 |
| Geminiviridae | 0.28 | -0.35 | 0.91 | 0.3873 | 23 | 8 |
| Luteoviridae | 0.95 | 0.50 | 1.41 | **<.0001** | 85 | 15 |
| Nanoviridae | 8.01 | 4.44 | 11.57 | **<.0001** | 1 | 1 |
| Potyviridae | 0.15 | -0.28 | 0.59 | 0.4851 | 70 | 19 |
| Reoviridae | -0.09 | -0.83 | 0.64 | 0.7986 | 22 | 4 |
| Secoviridae | 1.37 | -0.07 | 2.82 | 0.0622 | 3 | 3 |
| *Sobemovirus* | 1.60 | 0.11 | 3.08 | **0.0357** | 1 | 1 |
| Tospoviridae | 1.82 | 0.94 | 2.70 | **<.0001** | 34 | 9 |
| (Domestication) | | | | | | |
| Domesticated | 0.63 | 0.26 | 1.01 | **0.0009** | 282 | 65 |
| Wild | 0.57 | 0.13 | 1.00 | **0.0112** | 28 | 11 |

[a] the total number of infected/healthy comparisons

[b] the number of unique studies

**Table S2c:** Model estimated mean effect sizes, 95% confidence intervals, and sample sizes for each level of the various moderators influencing the effect of virus infection on **vector performance**

| Moderators | Estimate | CI.lb | CI.ub | pval | $N_{total}$[a] | $N_{st}$[b] |
|---|---|---|---|---|---|---|
| None | 0.29 | 0.05 | 0.52 | **0.0193** | 518 | 81 |
| Infection location | | | | | | |
| Phloem-limited | 0.44 | 0.20 | 0.69 | **0.0004** | 386 | 58 |
| Non-phloem-limited | -0.07 | -0.34 | 0.20 | 0.6096 | 132 | 27 |
| Retention mechanism | | | | | | |
| CPP | -0.15 | -0.57 | 0.27 | 0.4886 | 32 | 12 |
| CPNP | 0.54 | 0.29 | 0.78 | **<.0001** | 304 | 41 |
| NCSP | 0.01 | -0.28 | 0.29 | 0.9597 | 61 | 10 |
| NCNP | 0.17 | -0.10 | 0.45 | 0.2112 | 121 | 26 |
| Virus classification | | | | | | |
| Bromoviridae | -0.48 | -1.01 | 0.05 | 0.0749 | 31 | 8 |
| Caulimoviridae | -0.35 | -0.81 | 0.10 | 0.1292 | 6 | 1 |
| Closteroviridae | 0.36 | -0.07 | 0.80 | 0.0970 | 43 | 6 |
| Geminiviridae | 0.55 | 0.16 | 0.93 | **0.0052** | 164 | 24 |
| Luteoviridae | 0.59 | 0.24 | 0.95 | **0.0009** | 136 | 17 |
| Nanoviridae | 2.57 | 0.54 | 4.60 | **0.0129** | 4 | 1 |
| Potyviridae | 0.27 | -0.06 | 0.61 | 0.1121 | 96 | 21 |
| Reoviridae | 0.33 | -0.28 | 0.93 | 0.2901 | 11 | 3 |
| Secoviridae | 0.24 | -0.62 | 1.09 | 0.5883 | 4 | 2 |
| Sobemovirus | 0.28 | -0.61 | 1.17 | 0.5372 | 2 | 1 |
| Tospoviridae | -0.49 | -1.17 | 0.18 | 0.1528 | 21 | 9 |
| (Domestication) | | | | | | |
| Domesticated | 0.23 | -0.01 | 0.48 | 0.0624 | 493 | 75 |
| Wild | 0.82 | 0.44 | 1.20 | **<.0001** | 25 | 8 |

[a] the total number of infected/healthy comparisons

[b] the number of unique studies

3. **Supplementary Figures**

**a) Orientation preference**

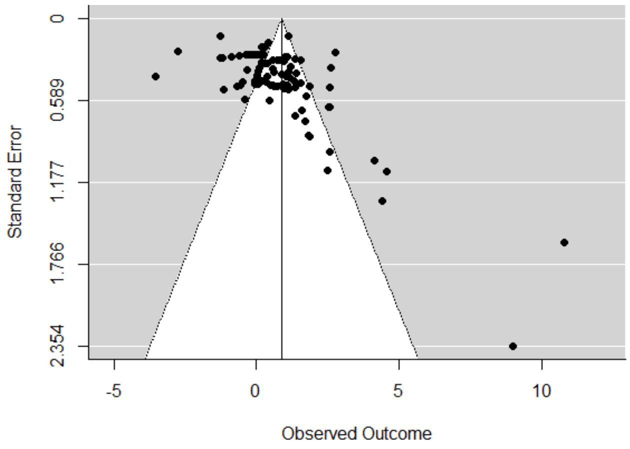


**b) Settling/feeding behaviors**

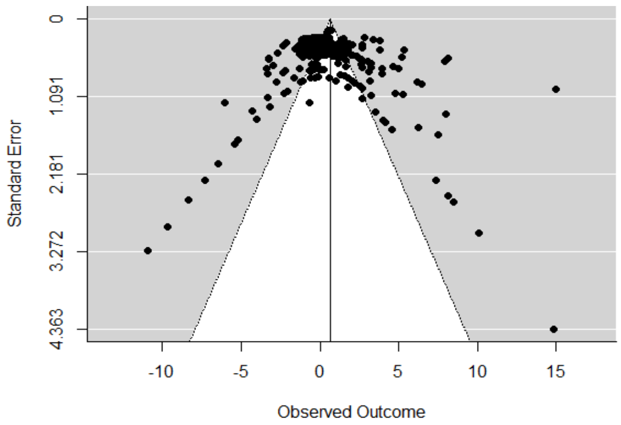


**c) Performance**

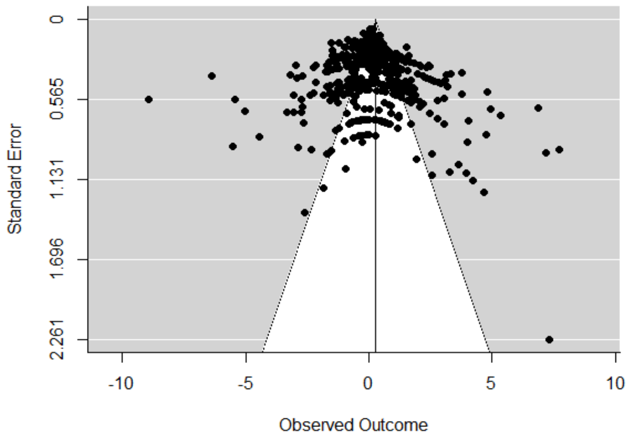


**Fig. S1:** Funnel plots of the effect sizes (Hedges' *d*) against their standard errors for a) vector orientation preference data, b) vector settling/feeding behaviors data and c) vector performance data.

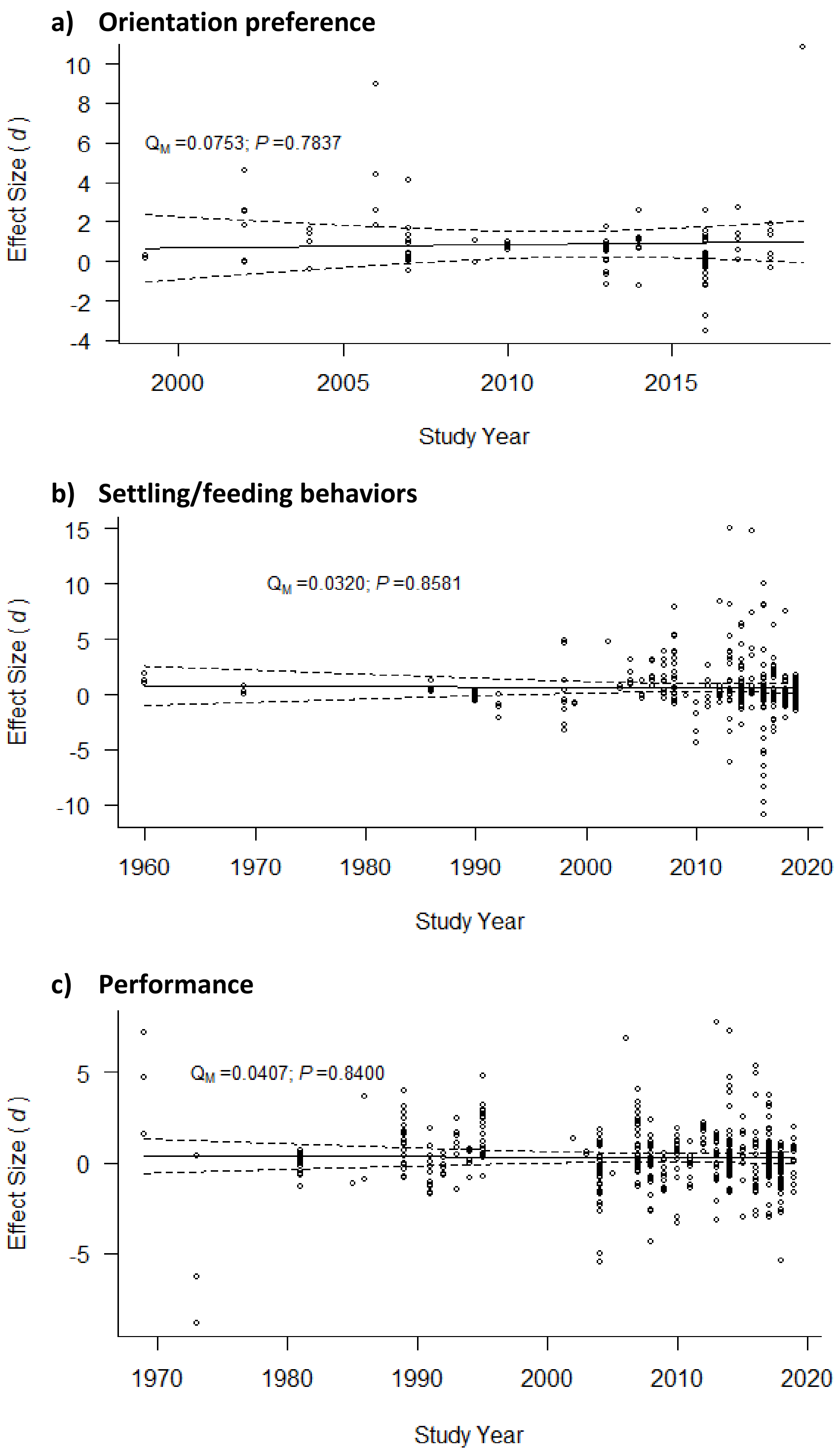


**Fig. S2:** Plots showing the plant infection effect sizes (Hedges' *d*) for a) vector orientation preference data, b) vector settling/feeding behavior data and c) vector performance data as a function of study year. Model statistics test the significance of study year as a moderator in mixed effects meta-regression models. Lines show the model predicted mean effect sizes and bounds of the corresponding 95% confidence intervals.

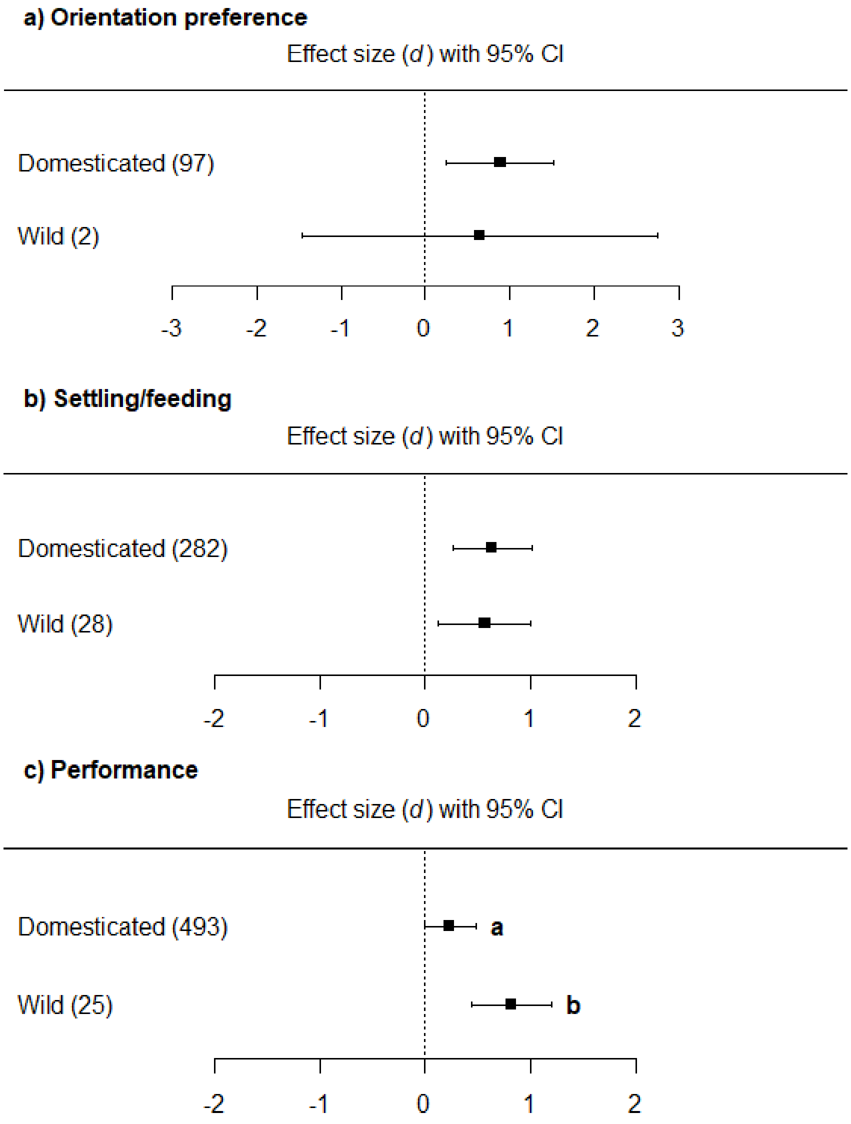


**Fig. S3:** Effect sizes (Hedges' *d*) and 95% confidence intervals comparing the magnitude of differences in a) vector orientation preference data, b) vector settling/feeding behaviors data and c) vector performance data, in domesticated-wild plant comparison. Numbers in brackets indicate the number of studies used to inform each estimate (see electronic supplementary material, table S2a, b and c, for complete sample size information).

#### 4. Assessing publication bias

Funnel plots revealed a slight asymmetry in effect size for plant infection effects on vector attraction (Fig. S1a). However, the fail-safe number (5979) was much higher than the threshold (5*99+10 = 505), indicating that these results are robust despite potential publication bias. For the effects of plant viruses on vector settling/feeding behaviors and performance (Fig. S1 a and b), the fail-safe numbers (settling/feeding: 27661 and performance: 28426) were also much higher than the threshold (settling/feeding: 5*310+10 = 1560 and performance: 5*518+10 = 2600).

#### 5. Complete list of studies included in the meta-analysis

### Start of content for file ESM_Meta-analysis database.xlsx

#### Orientation preference database

| Reference | Study | Year | Virus | Family | circulative | persistent | phloem_acq | Plant_family | wild | Vector | Healthy_mea | Healthy_SE | Healthy_SD | Healthy_n | Virus_mean | Virus_SE | Virus_SD | Virus_n | Parameter |
|---|---|---|---|---|---|---|---|---|---|---|---|---|---|---|---|---|---|---|---|
| Alvarez | 1 | 2007 | PLRV-Wagen | Luteoviridae | yes | CPNP | yes | Solanaceae | no | aphid | 2.0652 | 0.471 | 1.153709668 | 6 | 7.844 | 0.579899999 | 1.420459101 | 6 | headspace |
| Alvarez | 1 | 2007 | PLRV-Wagen | Luteoviridae | yes | CPNP | yes | Solanaceae | no | aphid | 2.8985 | 0.5979 | 1.464549917 | 6 | 3.8225 | 0.90576 | 2.218649829 | 6 | headspace |
| Claudel | 2 | 2018 | TuYV | Luteoviridae | yes | CPNP | yes | Brassicaceae | no | aphid | 26.53 | 3.09 | 9.771437969 | 10 | 44.89 | 4.92 | 15.55840608 | 10 | headspace |
| Claudel | 2 | 2018 | TuYV | Luteoviridae | yes | CPNP | yes | Brassicaceae | no | aphid | 27.72 | 3.4 | 13.16814337 | 15 | 23.91 | 2.26 | 8.752942362 | 15 | headspace |
| dosSantos | 3 | 2016 | BYDV-PAV | Luteoviridae | yes | CPNP | yes | Poaceae | no | aphid | 16.4 | 6.85 | 23.72909606 | 12 | 66.24 | 10.83 | 37.51622049 | 12 | headspace |
| dosSantos | 3 | 2016 | BYDV-PAV | Luteoviridae | yes | CPNP | yes | Poaceae | no | aphid | 43.79 | 11.3 | 39.14434825 | 12 | 45.06 | 11.95 | 41.39601430 | 12 | headspace |
| Eigenbrode | 4 | 2002 | PLRV | Luteoviridae | yes | CPNP | yes | Solanaceae | no | aphid | 32.5986 | 4.3653 | 7.560921390 | 3 | 60.9333 | 5.9983 | 10.38936035 | 3 | headspace |
| Eigenbrode | 4 | 2002 | PLRV | Luteoviridae | yes | CPNP | yes | Solanaceae | no | aphid | 33.6 | 2.7 | 6.613622305 | 6 | 66.4 | 2.7 | 6.613622305 | 6 | headspace |
| Eigenbrode | 4 | 2002 | PLRV | Luteoviridae | yes | CPNP | yes | Solanaceae | no | aphid | 10.7 | 1.9 | 3.8 | 4 | 18.6 | 1.9 | 3.8 | 4 | imigration |
| Eigenbrode | 4 | 2002 | PLRV | Luteoviridae | yes | CPNP | yes | Solanaceae | no | aphid | 9.40074 | 0.441550000 | 1.32465 | 9 | 4.66987 | 0.69373 | 2.08119 | 9 | emigration |
| Eigenbrode | 4 | 2002 | PLRV | Luteoviridae | yes | CPNP | yes | Solanaceae | no | aphid | 24.7905 | 1.1041 | 3.3123 | 9 | 15.4547 | 1.2299 | 3.6897 | 9 | emigration |
| Eigenbrode | 4 | 2002 | PVY | Potyviridae | no | NCNP | no | Solanaceae | no | aphid | 9.40074 | 0.441550000 | 1.32465 | 9 | 10.8147 | 11.3403 | 34.0209 | 9 | emigration |
| Eigenbrode | 4 | 2002 | PVY | Potyviridae | no | NCNP | no | Solanaceae | no | aphid | 15.8266 | 0.851699999 | 2.5551 | 9 | 16.33 | 17.04 | 51.12 | 9 | emigration |
| Eigenbrode | 4 | 2002 | PVY | Potyviridae | no | NCNP | no | Solanaceae | no | aphid | 20.1704 | 0.6938 | 2.0814 | 9 | 19.26 | 19.96 | 59.88 | 9 | emigration |
| Eigenbrode | 4 | 2002 | PVY | Potyviridae | no | NCNP | no | Solanaceae | no | aphid | 23.2531 | 0.694199999 | 2.0826 | 9 | 21.2025 | 21.66 | 64.98 | 9 | emigration |
| Eigenbrode | 4 | 2002 | PVY | Potyviridae | no | NCNP | no | Solanaceae | no | aphid | 24.7905 | 1.1041 | 3.3123 | 9 | 22.4295 | 22.9339 | 68.8017 | 9 | emigration |
| Fereres | 5 | 2016 | ToSRV | Geminivirida | yes | CPNP | yes | Solanaceae | no | Wf | 0.8 | 0.27 | 1.207476707 | 20 | 3.05 | 0.52 | 2.325510696 | 20 | headspace |
| Fereres | 5 | 2016 | ToSRV | Geminivirida | yes | CPNP | yes | Solanaceae | no | Wf | 0.3 | 0.1 | 0.447213595 | 20 | 0.55 | 0.22 | 0.983869910 | 20 | headspace |
| Fereres | 5 | 2016 | ToSRV | Geminivirida | yes | CPNP | yes | Solanaceae | no | Wf | 47.55 | 11.65 | 11.65 | 70 | 52.45 | 11.65 | 11.65 | 70 | Y |
| Fereres | 5 | 2016 | ToCV | Closterovirid | no | NCSP | yes | Solanaceae | no | Wf | 2.6 | 0.61 | 2.728002932 | 20 | 4 | 0.92 | 4.114365078 | 20 | headspace |
| Fereres | 5 | 2016 | ToCV | Closterovirid | no | NCSP | yes | Solanaceae | no | Wf | 0.05 | 0.05 | 0.223606797 | 20 | 0.15 | 0.11 | 0.491934955 | 20 | headspace |
| Fereres | 5 | 2016 | ToCV | Closterovirid | no | NCSP | yes | Solanaceae | no | Wf | 65.48 | 11.18 | 11.18 | 70 | 34.52 | 11.18 | 11.18 | 70 | Y |
| Fereres | 6 | 1999 | SMV-unknov | Potyviridae | no | NCNP | no | Fabaceae | no | aphid | 4.37 | 0.63 | 4.364768035 | 48 | 5.31 | 0.92 | 6.373946971 | 48 | landing |
| Fereres | 6 | 1999 | SMV-unknov | Potyviridae | no | NCNP | no | Fabaceae | no | aphid | 1.87 | 0.55 | 3.810511776 | 48 | 3.33 | 0.81 | 5.611844616 | 48 | landing |
| Jimenez | 7 | 2004 | BYDV | Luteoviridae | yes | CPNP | yes | Poaceae | no | aphid | 6.3 | 1.06 | 2.596459127 | 6 | 11.8 | 1.52 | 3.723224409 | 6 | headspace |
| Jimenez | 7 | 2004 | BYDV | Luteoviridae | yes | CPNP | yes | Poaceae | no | aphid | 7.7 | 1.1 | 2.694438717 | 6 | 6.72 | 0.74 | 1.812622409 | 6 | headspace |
| Jimenez | 7 | 2004 | BYDV | Luteoviridae | yes | CPNP | yes | Poaceae | no | aphid | 10 | 0.65 | 2.055480479 | 10 | 14.1 | 1.59 | 5.028021479 | 10 | headspace |
| Jimenez | 7 | 2004 | BYDV | Luteoviridae | yes | CPNP | yes | Poaceae | no | aphid | 10.5 | 0.71 | 2.245217138 | 10 | 14 | 0.83 | 2.624690457 | 10 | headspace |
| Lu | 8 | 2016 | SRBSDV | Reoviridae | yes | CPP | yes | Poaceae | no | PH | 53.16 | 3.71 | 20.32050688 | 30 | 46.04 | 3.44 | 18.84165597 | 30 | Y |
| Lu | 8 | 2016 | SRBSDV | Reoviridae | yes | CPP | yes | Poaceae | no | PH | 53.38 | 11.63 | 63.70013343 | 30 | 46.62 | 11.37 | 62.27605478 | 30 | Y |
| Lu | 8 | 2016 | SRBSDV | Reoviridae | yes | CPP | yes | Poaceae | no | PH | 69.72 | 5.56 | 30.45337419 | 30 | 29.7456 | 5.8244 | 31.90155263 | 30 | Y |
| Lu | 8 | 2016 | SRBSDV | Reoviridae | yes | CPP | yes | Poaceae | no | PH | 49.3 | 6.080000000 | 33.30153149 | 30 | 49.38 | 6.349999999 | 34.78038240 | 30 | Y |
| Lu | 8 | 2016 | SRBSDV | Reoviridae | yes | CPP | yes | Poaceae | no | PH | 23.32 | 9.52 | 52.14318747 | 30 | 76.15 | 9.789999999 | 53.62203837 | 30 | Y |
| Lu | 8 | 2016 | SRBSDV | Reoviridae | yes | CPP | yes | Poaceae | no | PH | 20.09 | 9.79 | 53.62203837 | 30 | 79.37 | 10.06 | 55.10088928 | 30 | Y |
| Lu | 8 | 2016 | SRBSDV | Reoviridae | yes | CPP | yes | Poaceae | no | PH | 30.09 | 4.5 | 24.64751508 | 30 | 69.64 | 4.760000000 | 26.07159373 | 30 | Y |
| Lu | 8 | 2016 | SRBSDV | Reoviridae | yes | CPP | yes | Poaceae | no | PH | 55.97 | 0.700000000 | 3.834057902 | 30 | 42.98 | 3.69 | 20.21096237 | 30 | Y |
| Lu | 8 | 2016 | RRSV | Reoviridae | yes | CPP | yes | Poaceae | no | PH | 56.91 | 12.24 | 67.04124103 | 30 | 43.35 | 12.24 | 67.04124103 | 30 | Y |
| Lu | 8 | 2016 | RRSV | Reoviridae | yes | CPP | yes | Poaceae | no | PH | 35.9 | 19.42 | 106.3677206 | 30 | 63.56 | 19.42 | 106.3677206 | 30 | Y |
| Lu | 8 | 2016 | RRSV | Reoviridae | yes | CPP | yes | Poaceae | no | PH | 44.15 | 12.76 | 69.88939833 | 30 | 55.59 | 13.56 | 74.27117879 | 30 | Y |
| Lu | 8 | 2016 | RRSV | Reoviridae | yes | CPP | yes | Poaceae | no | PH | 58.51 | 12.5 | 68.46531968 | 30 | 41.75 | 13.04 | 71.42302149 | 30 | Y |
| Lu | 8 | 2016 | RRSV | Reoviridae | yes | CPP | yes | Poaceae | no | PH | 60.1 | 5.59 | 30.61769096 | 30 | 39.36 | 6.65 | 36.42355007 | 30 | Y |
| Lu | 8 | 2016 | RRSV | Reoviridae | yes | CPP | yes | Poaceae | no | PH | 26.86 | 3.72 | 20.37527913 | 30 | 73.67 | 2.66 | 14.56942002 | 30 | Y |
| Lu | 8 | 2016 | RRSV | Reoviridae | yes | CPP | yes | Poaceae | no | PH | 16.75 | 9.31 | 50.99297010 | 30 | 83.51 | 8.239999999 | 45.13233873 | 30 | Y |
| Lu | 8 | 2016 | RRSV | Reoviridae | yes | CPP | yes | Poaceae | no | PH | 66.75 | 5.06 | 27.71476140 | 30 | 33.24 | 5.05 | 27.65998915 | 30 | Y |
| Mauck | 9 | 2010 | CMV-FNY | Bromovirida | no | NCNP | no | Cucurbitacea | no | aphid | 1.62 | 0.41 | 2.008581589 | 24 | 3.21 | 0.35 | 1.714642819 | 24 | headspace |
| Mauck | 9 | 2010 | CMV-FNY | Bromovirida | no | NCNP | no | Cucurbitacea | no | aphid | 1.91 | 0.11 | 0.538887743 | 24 | 3.66 | 0.66 | 3.233326460 | 24 | headspace |
| Mauck | 9 | 2010 | CMV-FNY | Bromovirida | no | NCNP | no | Cucurbitacea | no | aphid | 1.09 | 0.31 | 1.518683640 | 24 | 1.94 | 0.31 | 1.518683640 | 24 | headspace |
| Mauck | 9 | 2010 | CMV-FNY | Bromovirida | no | NCNP | no | Cucurbitacea | no | aphid | 2.22 | 0.27 | 1.322724461 | 24 | 3.88 | 0.41 | 2.008581589 | 24 | headspace |
| Mauck | 10 | 2014 | CMV-KVPG2 | Bromovirida | no | NCNP | no | Solanaceae | no | aphid | 1.244 | 0.155 | 0.465 | 9 | 1.8052 | 0.31726 | 0.95178 | 9 | headspace |
| Mauck | 10 | 2014 | CMV-KVPG2 | Bromovirida | no | NCNP | no | Cucurbitacea | no | aphid | 1.1121 | 0.12693 | 0.38079 | 9 | 2.43798 | 0.51431 | 1.54293 | 9 | headspace |
| Mauck | 10 | 2014 | CMV-P1 | Bromovirida | no | NCNP | no | Solanaceae | no | aphid | 1.15836 | 0.27726 | 0.83178 | 9 | 1.70113 | 0.27255 | 0.81765 | 9 | headspace |
| Medina | 11 | 2009 | BYDV-PAV | Luteoviridae | yes | CPNP | yes | Poaceae | no | aphid | 4.1 | 0.97 | 3.067409330 | 10 | 7.9 | 1.19 | 3.763110415 | 10 | headspace |
| Medina | 11 | 2009 | BYDV-PAV | Luteoviridae | yes | CPNP | yes | Poaceae | no | aphid | 5.1 | 0.67 | 2.118726032 | 10 | 4.9 | 0.95 | 3.004163777 | 10 | headspace |
| Mwando | 12 | 2018 | MCMV | Potyviridae | no | NCNP | no | Poaceae | no | thrips | 2.44685 | 0.2939 | 1.018099464 | 12 | 3.8686 | 0.31352 | 1.086065138 | 12 | Y |
| Mwando | 12 | 2018 | MCMV | Potyviridae | no | NCNP | no | Poaceae | no | thrips | 2.74 | 0.33307 | 1.153788324 | 12 | 3.16266 | 0.32816 | 1.136779586 | 12 | Y |
| Mwando | 12 | 2018 | MCMV | Potyviridae | no | NCNP | no | Poaceae | no | thrips | 2.5227 | 0.24723 | 0.856429842 | 12 | 4.22301 | 0.262359999 | 0.908841699 | 12 | Y |
| Mwando | 12 | 2018 | MCMV | Potyviridae | no | NCNP | no | Poaceae | no | thrips | 2.94652 | 0.25238 | 0.874269965 | 12 | 4.32896 | 0.24723 | 0.856429842 | 12 | Y |
| Ngumbi | 13 | 2007 | PLRV-Idaho | Luteoviridae | yes | CPNP | yes | Solanaceae | no | aphid | 12.4629 | 1.543 | 3.450252889 | 5 | 7.15134 | 0.86053 | 1.924203576 | 5 | emigration |
| Ngumbi | 13 | 2007 | PLRV-Idaho | Luteoviridae | yes | CPNP | yes | Solanaceae | no | aphid | 21.632 | 1.3354 | 2.986045177 | 5 | 16.2315 | 1.8397 | 4.113694258 | 5 | emigration |
| Penaflor | 14 | 2016 | BPMV | Secoviridae | no | NCSP | no | Fabaceae | no | beetle | 77.03 | 15.07 | 15.07 | 30 | 22.97 | 15.03 | 15.03 | 30 | Y |
| Rajabaskar | 15 | 2014 | PLRV | Luteoviridae | yes | CPNP | yes | Solanaceae | no | aphid | 36.86 | 2.76 | 10.68943403 | 15 | 63.21 | 2.4 | 9.295160030 | 15 | headspace |
| Rajabaskar | 24 | 2013 | PLRV | Luteoviridae | yes | CPNP | yes | Solanaceae | no | aphid | 17.0724 | 0.700900000 | 2.622527662 | 14 | 16.8185 | 1.7095 | 6.396363302 | 14 | emigration |
| Rajabaskar | 24 | 2013 | PLRV | Luteoviridae | yes | CPNP | yes | Solanaceae | no | aphid | 19.7991 | 0.897000000 | 3.356266675 | 14 | 16.0979 | 1.2612 | 4.718978296 | 14 | emigration |
| Rajabaskar | 24 | 2013 | PLRV | Luteoviridae | yes | CPNP | yes | Solanaceae | no | aphid | 16.5279 | 0.925000000 | 3.461033082 | 14 | 16.1618 | 1.7377 | 6.501878040 | 14 | emigration |
| Rajabaskar | 24 | 2013 | PLRV | Luteoviridae | yes | CPNP | yes | Solanaceae | no | aphid | 19.5348 | 0.784800000 | 2.936452717 | 14 | 15.8054 | 2.0461 | 7.655805179 | 14 | emigration |
| Rajabaskar | 16 | 2013 | PLRV | Luteoviridae | yes | CPNP | yes | Solanaceae | no | aphid | 11.7827 | 1.0526 | 3.1578 | 9 | 13.854 | 0.798 | 2.394 | 9 | arrestment |
| Rajabaskar | 16 | 2013 | PLRV | Luteoviridae | yes | CPNP | yes | Solanaceae | no | aphid | 13.5993 | 1.0187 | 3.0561 | 9 | 16.7572 | 1.0357 | 3.1071 | 9 | arrestment |
| Rajabaskar | 16 | 2013 | PLRV | Luteoviridae | yes | CPNP | yes | Solanaceae | no | aphid | 13.6163 | 1.2903 | 3.8709 | 9 | 15.8913 | 1.2225 | 3.6675 | 9 | arrestment |
| Rajabaskar | 16 | 2013 | PLRV | Luteoviridae | yes | CPNP | yes | Solanaceae | no | aphid | 10.7357 | 1.1044 | 3.3132 | 9 | 12.8762 | 1.5123 | 4.5369 | 9 | arrestment |
| Rajabaskar | 16 | 2013 | PLRV | Luteoviridae | yes | CPNP | yes | Solanaceae | no | aphid | 12.7566 | 1.1966 | 3.5898 | 9 | 14.5028 | 1.065 | 3.195 | 9 | arrestment |
| Rajabaskar | 16 | 2013 | PLRV | Luteoviridae | yes | CPNP | yes | Solanaceae | no | aphid | 11.7927 | 1.0651 | 3.1953 | 9 | 16.905 | 0.7758 | 2.3274 | 9 | arrestment |
| Rajabaskar | 16 | 2013 | PLRV | Luteoviridae | yes | CPNP | yes | Solanaceae | no | aphid | 14.8132 | 1.5082 | 4.5246 | 9 | 9.82382 | 1.18398 | 3.55194 | 9 | arrestment |
| Rajabaskar | 16 | 2013 | PLRV | Luteoviridae | yes | CPNP | yes | Solanaceae | no | aphid | 14.7851 | 1.3107 | 3.9321 | 9 | 11.9803 | 1.3389 | 4.0167 | 9 | arrestment |
| Rajabaskar | 16 | 2013 | PLRV | Luteoviridae | yes | CPNP | yes | Solanaceae | no | aphid | 15.6025 | 1.48 | 4.44 | 9 | 13.3897 | 0.9161 | 2.7483 | 9 | arrestment |
| Safari | 25 | 2019 | CMV | Bromovirida | no | NCNP | no | Brassicaceae | no | aphid | 36.62 | 0.970000000 | 3.360178566 | 12 | 64.24 | 0.270000000 | 0.935307436 | 12 | headspace |
| Shresta | 17 | 2017 | SVYV | Potyviridae | no | NCSP | yes | Cucurbitacea | no | Wf | 0.489583333 | 0.244416666 | 0.244416666 | 16 | 0.510416666 | 0.244583333 | 0.244583333 | 16 | landing |
| Shresta | 17 | 2017 | SVYV | Potyviridae | no | NCSP | yes | Cucurbitacea | no | Wf | 0.427083333 | 0.241916666 | 0.241916666 | 16 | 0.572916666 | 0.242083333 | 0.242083333 | 16 | landing |
| Shresta | 17 | 2017 | SVYV | Potyviridae | no | NCSP | yes | Cucurbitacea | no | Wf | 0.364583333 | 0.235416666 | 0.235416666 | 16 | 0.635416666 | 0.235583333 | 0.235583333 | 16 | landing |
| Shresta | 17 | 2017 | SVYV | Potyviridae | no | NCSP | yes | Cucurbitacea | no | Wf | 0.333333333 | 0.230666666 | 0.230666666 | 16 | 0.666666666 | 0.231333333 | 0.231333333 | 16 | landing |
| Srinivasan | 18 | 2006 | PLRV-Idaho | Luteoviridae | yes | CPNP | yes | Solanaceae | yes | aphid | 7.788 | 1.43538 | 2.87076 | 4 | 3.2712 | 0.42225 | 0.8445 | 4 | emigration |
| Srinivasan | 18 | 2006 | PLRV-Idaho | Luteoviridae | yes | CPNP | yes | Solanaceae | yes | aphid | 21.0227 | 0.380099999 | 0.760199999 | 4 | 13.5507 | 0.969299999 | 1.9386 | 4 | emigration |
| Srinivasan | 18 | 2006 | PLRV-Idaho | Luteoviridae | yes | CPNP | yes | Solanaceae | no | aphid | 10.6163 | 0.971 | 1.942 | 4 | 6.26838 | 0.379989999 | 0.759979999 | 4 | emigration |
| Srinivasan | 18 | 2006 | PLRV-Idaho | Luteoviridae | yes | CPNP | yes | Solanaceae | no | aphid | 25.0331 | 0.4221 | 0.8442 | 4 | 11.2291 | 0.8444 | 1.6888 | 4 | emigration |
| Wang | 19 | 2014 | SRBSDV | Reoviridae | yes | CPP | yes | Poaceae | no | PH | 8 | 0.89 | 10.90022935 | 150 | 18 | 0.55 | 6.736096792 | 150 | Y |
| Wang | 19 | 2014 | RRSV | Reoviridae | yes | CPP | yes | Poaceae | no | PH | 16.8 | 0.58 | 7.103520254 | 150 | 7.8 | 0.58 | 7.103520254 | 150 | Y |
| Wang | 20 | 2018 | RDV | Reoviridae | yes | CPP | yes | Poaceae | no | LH | 636.062 | 777.982 | 3479.241274 | 20 | 2005.91 | 2314.38 | 10350.22201 | 20 | Y |
| Werner | 21 | 2007 | PLRV-Idaho | Luteoviridae | yes | CPNP | yes | Solanaceae | no | aphid | 12.2158 | 2.1048 | 6.655962019 | 10 | 12.1278 | 2.22217 | 7.027118548 | 10 | emigration |
| Werner | 21 | 2007 | PLRV-Idaho | Luteoviridae | yes | CPNP | yes | Solanaceae | no | aphid | 15.6529 | 2.5433 | 8.042620773 | 10 | 9.36601 | 1.49119 | 4.715556824 | 10 | emigration |
| Werner | 21 | 2007 | PLRV-Idaho | Luteoviridae | yes | CPNP | yes | Solanaceae | no | aphid | 12.4236 | 1.3448 | 4.252630997 | 10 | 8.32969 | 0.876940000 | 2.773127771 | 10 | emigration |
| Werner | 21 | 2007 | PLRV-Idaho | Luteoviridae | yes | CPNP | yes | Solanaceae | no | aphid | 12.2938 | 2.3093 | 7.302647800 | 10 | 11.855 | 2.74901 | 8.693132910 | 10 | emigration |
| Werner | 21 | 2007 | PLRV-Idaho | Luteoviridae | yes | CPNP | yes | Solanaceae | no | aphid | 11.959 | 1.7536 | 5.545370104 | 10 | 9.38571 | 2.57348 | 8.138058312 | 10 | emigration |
| Werner | 21 | 2007 | PLRV-Idaho | Luteoviridae | yes | CPNP | yes | Solanaceae | no | aphid | 11.4352 | 2.7303 | 8.633966695 | 10 | 9.35109 | 2.14242 | 6.774926904 | 10 | emigration |
| Werner | 21 | 2007 | PLRV-Idaho | Luteoviridae | yes | CPNP | yes | Solanaceae | no | aphid | 11.3015 | 2.0844 | 6.591451554 | 10 | 10.7145 | 1.3794 | 4.362045804 | 10 | emigration |
| Werner | 21 | 2007 | PLRV-Idaho | Luteoviridae | yes | CPNP | yes | Solanaceae | no | aphid | 8.84887 | 2.46633 | 7.799220261 | 10 | 8.87835 | 1.14469 | 3.619827614 | 10 | emigration |
| Werner | 21 | 2007 | PLRV-Idaho | Luteoviridae | yes | CPNP | yes | Solanaceae | no | aphid | 11.4749 | 2.4363 | 7.704257063 | 10 | 10.5353 | 1.11521 | 3.526603669 | 10 | emigration |
| Werner | 21 | 2007 | PLRV-Idaho | Luteoviridae | yes | CPNP | yes | Solanaceae | no | aphid | 7.26035 | 1.90778 | 6.032930074 | 10 | 10.2549 | 1.7319 | 5.476748679 | 10 | emigration |
| Wu | 22 | 2014 | PEMV | Luteoviridae | yes | CPNP | yes | Fabaceae | no | aphid | 39.73 | 5.18 | 18.67675560 | 13 | 60.43 | 5.52 | 19.90264304 | 13 | Y |
| Wu | 22 | 2014 | BLRV | Luteoviridae | yes | CPNP | yes | Fabaceae | no | aphid | 40.54 | 4.01 | 14.45826061 | 13 | 59.62 | 4.69 | 16.91003548 | 13 | Y |
| Wu | 23 | 2017 | CMV | Bromovirida | no | NCNP | no | Brassicaceae | no | aphid | 34.0707 | 11.53 | 11.53 | 65 | 65.9285 | 11.47 | 11.47 | 65 | Y |

### Settling/feeding preference database

| Reference | Study | Year | Virus | Family | circulative | persistent | phloem_acq | Plant_family | wild | Vector | Healthy_mea | Healthy_SE | Healthy_SD | Healthy_n | Virus_mean | Virus_SE | Virus_SD | Virus_n | Parameter |
|---|---|---|---|---|---|---|---|---|---|---|---|---|---|---|---|---|---|---|---|
| Abe | 1 | 2012 | TSWV | Tospoviridae | yes | CPP | yes | Brassicaceae | no | thrips | 10.264 | 1.525 | 1.525 | 3 | 31.267 | 2.3458 | 2.3458 | 3 | settling |
| Alvarez | 2 | 2007 | PLRV | Luteoviridae | yes | CPNP | yes | Solanaceae | no | aphid | 10 | 8 | 35.77708763 | 20 | 1 | 1 | 4.472135954 | 20 | EPG_E2 |
| Alvarez | 2 | 2007 | PLRV | Luteoviridae | yes | CPNP | yes | Solanaceae | no | aphid | 28 | 15 | 58.09475019 | 15 | 62 | 26 | 107.2007462 | 17 | EPG_E2 |
| Alvarez | 2 | 2007 | PLRV | Luteoviridae | yes | CPNP | yes | Solanaceae | no | aphid | 70 | 4 | 17.88854381 | 20 | 62 | 5 | 22.36067977 | 20 | EPG |
| Alvarez | 2 | 2007 | PLRV | Luteoviridae | yes | CPNP | yes | Solanaceae | no | aphid | 52 | 6 | 23.2379000 | 15 | 51 | 5 | 20.61552812 | 17 | EPG |
| Angelela | 66 | 2018 | WMV | Potyviridae | no | NCNP | no | Fabaceae | yes | aphid | 6.08 | 4.44 | 4.44 | 12 | 10.57 | 3.87 | 3.87 | 7 | EPG |
| Angelela | 66 | 2018 | WMV | Potyviridae | no | NCNP | no | Fabaceae | no | aphid | 2.73 | 2.8 | 2.8 | 11 | 4.44 | 4.77 | 4.77 | 9 | EPG |
| Baker | 3 | 1960 | BYV | Closteroviric | no | NCSP | yes | Amaranthac | no | aphid | 29 | 19.4 | 19.4 | 21 | 54 | 21.3 | 21.3 | 21 | settling |
| Baker | 3 | 1960 | BYV | Closteroviric | no | NCSP | yes | Amaranthac | no | aphid | 22 | 23.4 | 23.4 | 12 | 50 | 27.2 | 27.2 | 13 | settling |
| Baker | 3 | 1960 | BYV | Closteroviric | no | NCSP | yes | Amaranthac | no | aphid | 21 | 15.4 | 15.4 | 27 | 55 | 21.3 | 21.3 | 21 | settling |
| Blua | 5 | 1992 | ZYMV | Potyviridae | no | NCNP | no | Cucurbitacea | no | aphid | 8.16 | 0.57 | 2.906441122 | 26 | 8.23 | 0.6 | 3.117691453 | 27 | EPG |
| Blua | 5 | 1992 | ZYMV | Potyviridae | no | NCNP | no | Cucurbitacea | no | aphid | 8.16 | 0.57 | 2.906441122 | 26 | 12.68 | 0.870000000 | 4.685093382 | 29 | EPG |
| Blua | 5 | 1992 | ZYMV | Potyviridae | no | NCNP | no | Cucurbitacea | no | aphid | 1.74 | 0.13 | 0.662872536 | 26 | 0.99 | 0.14 | 0.727461339 | 27 | EPG_E2 |
| Blua | 5 | 1992 | ZYMV | Potyviridae | no | NCNP | no | Cucurbitacea | no | aphid | 1.74 | 0.13 | 0.662872536 | 26 | 0.32 | 0.12 | 0.646219776 | 29 | EPG_E2 |
| Blua | 5 | 1992 | ZYMV | Potyviridae | no | NCNP | no | Cucurbitacea | no | aphid | 2.33 | 0.23 | 0.796743371 | 12 | 2.33 | 0.24 | 0.831384387 | 12 | retention |
| Blua | 5 | 1992 | ZYMV | Potyviridae | no | NCNP | no | Cucurbitacea | no | aphid | 2.33 | 0.23 | 0.796743371 | 12 | 1.6 | 0.23 | 0.796743371 | 12 | retention |
| Boquel | 6 | 2011 | PVY | Potyviridae | no | NCNP | no | Solanaceae | no | aphid | 68.36 | 13.64 | 60.99993442 | 20 | 207.9 | 32.1 | 150.5623458 | 22 | EPG |
| Boquel | 6 | 2011 | PVY | Potyviridae | no | NCNP | no | Solanaceae | no | aphid | 96.55 | 25.25 | 112.9214328 | 20 | 26.9 | 14.28 | 66.97913705 | 22 | EPG_E2 |
| Boquel | 6 | 2011 | PVY | Potyviridae | no | NCNP | no | Solanaceae | no | aphid | 259 | 38 | 165.6381598 | 19 | 306.9 | 39.4 | 176.2021566 | 20 | EPG |
| Boquel | 6 | 2011 | PVY | Potyviridae | no | NCNP | no | Solanaceae | no | aphid | 34.88 | 16.29 | 71.00646379 | 19 | 186 | 34.4 | 153.8414768 | 20 | EPG_E2 |
| Boquel | 7 | 2012 | PVY | Potyviridae | no | NCNP | no | Solanaceae | no | aphid | 37.4 | 3.6 | 16.09968943 | 20 | 38.1 | 3 | 13.41640786 | 20 | EPG |
| Boquel | 7 | 2012 | PVY | Potyviridae | no | NCNP | no | Solanaceae | no | aphid | 97.3 | 30 | 134.1640786 | 20 | 140.9 | 24.2 | 108.2256901 | 20 | EPG_E2 |
| Boquel | 7 | 2012 | PVY | Potyviridae | no | NCNP | no | Solanaceae | no | aphid | 19.7 | 3.2 | 14.31083505 | 20 | 29.7 | 2.9 | 12.96919426 | 20 | EPG |
| Boquel | 7 | 2012 | PVY | Potyviridae | no | NCNP | no | Solanaceae | no | aphid | 0.001 | 0.001 | 0.004472135 | 20 | 13.5 | 5.9 | 26.38560213 | 20 | EPG_E2 |
| Boquel | 7 | 2012 | PVY | Potyviridae | no | NCNP | no | Solanaceae | no | aphid | 15.5 | 3.2 | 14.31083505 | 20 | 17.6 | 3.2 | 14.31083505 | 20 | EPG |
| Boquel | 7 | 2012 | PVY | Potyviridae | no | NCNP | no | Solanaceae | no | aphid | 3.5 | 3.5 | 15.65247584 | 20 | 16.3 | 14.5 | 64.84597134 | 20 | EPG_E2 |
| Boquel | 7 | 2012 | PVY | Potyviridae | no | NCNP | no | Solanaceae | no | aphid | 52.93 | 18.17 | 18.17 | 29 | 47.49 | 18.21 | 18.21 | 29 | settling |
| Boquel | 7 | 2012 | PVY | Potyviridae | no | NCNP | no | Solanaceae | no | aphid | 57.33 | 20.67 | 20.67 | 22 | 42.84 | 20.66 | 20.66 | 22 | settling |
| Boquel | 7 | 2012 | PVY | Potyviridae | no | NCNP | no | Solanaceae | no | aphid | 49.69 | 23.11 | 23.11 | 18 | 50.21 | 23.09 | 23.09 | 18 | settling |
| Camo-sousa | 8 | 2014 | CMV | Bromovirida | no | NCNP | no | Cucurbitacea | no | aphid | 0.43 | 0.45 | 1.8 | 16 | 1.01 | 0.58 | 2.32 | 16 | dispersion |
| Camo-sousa | 8 | 2014 | CMV | Bromovirida | no | NCNP | no | Cucurbitacea | no | aphid | 2.73 | 0.96 | 3.84 | 16 | 10.77 | 2.53 | 10.12 | 16 | dispersion |
| Camo-sousa | 8 | 2014 | CMV | Bromovirida | no | NCNP | no | Cucurbitacea | no | aphid | 6.93 | 3.13 | 5.421319027 | 3 | 5.31 | 1.09 | 1.887935380 | 3 | settling |
| Camo-sousa | 8 | 2014 | CMV | Bromovirida | no | NCNP | no | Cucurbitacea | no | aphid | 11.57 | 1.81 | 3.135011961 | 3 | 7.47 | 1.62 | 2.805922308 | 3 | settling |
| Camo-sousa | 8 | 2014 | CMV | Bromovirida | no | NCNP | no | Cucurbitacea | no | aphid | 2.93 | 0.47 | 2.574296020 | 30 | 5.64 | 0.79 | 4.180287071 | 28 | EPG |
| Camo-sousa | 8 | 2014 | CMV | Bromovirida | no | NCNP | no | Cucurbitacea | no | aphid | 18.99 | 3.4 | 18.62256695 | 30 | 35.91 | 4.55 | 24.07633693 | 28 | EPG |
| Camo-sousa | 8 | 2014 | CMV | Bromovirida | no | NCNP | no | Cucurbitacea | no | aphid | 26.28 | 49.24 | 269.6985873 | 30 | 0.57 | 0.67 | 3.545306756 | 28 | EPG_E2 |
| Carmo-sousa | 9 | 2016 | CABYV | Luteoviridae | yes | CPNP | yes | Cucurbitacea | no | aphid | 6.459 | 1.655 | 7.021570337 | 18 | 7.28 | 2.55 | 10.81873375 | 18 | settling |
| Carmo-sousa | 9 | 2016 | CABYV | Luteoviridae | yes | CPNP | yes | Cucurbitacea | no | aphid | 10.73 | 2.699 | 11.45088721 | 18 | 8.235 | 0.941999999 | 3.996567527 | 18 | settling |
| Carmo-sousa | 9 | 2016 | CABYV | Luteoviridae | yes | CPNP | yes | Cucurbitacea | no | aphid | 5.15 | 0.14 | 0.56 | 16 | 5.01487 | 0.1487 | 0.5948 | 16 | EPG |
| Carmo-sousa | 9 | 2016 | CABYV | Luteoviridae | yes | CPNP | yes | Cucurbitacea | no | aphid | 52.52 | 29.0535 | 116.214 | 16 | 105.04 | 26.02 | 104.08 | 16 | EPG_E2 |
| Carmo-sousa | 9 | 2016 | CABYV | Luteoviridae | yes | CPNP | yes | Cucurbitacea | no | aphid | 5.12 | 1.49 | 5.770745185 | 15 | 7.64 | 2.2 | 8.520563361 | 15 | dispersion |
| Carmo-sousa | 9 | 2016 | CABYV | Luteoviridae | yes | CPNP | yes | Cucurbitacea | no | aphid | 14.45 | 2.24 | 8.675482695 | 15 | 15.71 | 2.2 | 8.520563361 | 15 | dispersion |
| Casteel | 10 | 2014 | TuMV | Potyviridae | no | NCNP | no | Solanaceae | no | aphid | 31 | 7.97 | 27.60888987 | 12 | 69.77 | 7.53 | 26.08468516 | 12 | settling |
| Casteel | 10 | 2014 | TuMV | Potyviridae | no | NCNP | no | Solanaceae | no | aphid | 29.12 | 4.65 | 16.10807251 | 12 | 71.44 | 5.52 | 19.12184091 | 12 | settling |
| Casteel | 10 | 2014 | TuMV | Potyviridae | no | NCNP | no | Solanaceae | no | aphid | 24.52 | 5.48 | 18.98327685 | 12 | 75.28 | 5.849999999 | 20.26499444 | 12 | settling |
| Casteel | 10 | 2014 | TuMV | Potyviridae | no | NCNP | no | Solanaceae | no | aphid | 51.51 | 16.79 | 58.16226611 | 12 | 47.92 | 17.17 | 59.47862473 | 12 | settling |
| Castle | 11 | 1998 | PLRV | Luteoviridae | yes | CPNP | yes | Solanaceae | yes | aphid | 7.16279 | 0.96457 | 0.96457 | 8 | 7.52859 | 0.96464 | 0.96464 | 8 | settling |
| Castle | 11 | 1998 | PLRV | Luteoviridae | yes | CPNP | yes | Solanaceae | yes | aphid | 9.55226 | 1.08114 | 1.08114 | 8 | 9.05332 | 1.01458 | 1.01458 | 8 | settling |
| Castle | 11 | 1998 | PLRV | Luteoviridae | yes | CPNP | yes | Solanaceae | no | aphid | 7.15942 | 0.844790000 | 0.844790000 | 17 | 11.0849 | 0.6957 | 0.6957 | 17 | settling |
| Castle | 11 | 1998 | PLRV | Luteoviridae | yes | CPNP | yes | Solanaceae | no | aphid | 7.89236 | 1.25881 | 1.25881 | 17 | 13.7391 | 1.1927 | 1.1927 | 17 | settling |
| Castle | 11 | 1998 | PLRV | Luteoviridae | yes | CPNP | yes | Solanaceae | yes | aphid | 12.2677 | 0.9557 | 0.9557 | 15 | 9.20281 | 0.856829999 | 0.856829999 | 15 | settling |
| Castle | 11 | 1998 | PLRV | Luteoviridae | yes | CPNP | yes | Solanaceae | yes | aphid | 14.9289 | 0.9887 | 0.9887 | 15 | 11.6168 | 0.988900000 | 0.988900000 | 15 | settling |
| Castle | 11 | 1998 | PVY | Potyviridae | no | NCNP | no | Solanaceae | yes | aphid | 7.16279 | 0.96457 | 0.96457 | 8 | 8.60976 | 1.24756 | 1.24756 | 8 | settling |
| Castle | 11 | 1998 | PVY | Potyviridae | no | NCNP | no | Solanaceae | yes | aphid | 9.55226 | 1.08114 | 1.08114 | 8 | 8.72071 | 1.16422 | 1.16422 | 8 | settling |
| Castle | 11 | 1998 | PVY | Potyviridae | no | NCNP | no | Solanaceae | no | aphid | 7.15942 | 0.844790000 | 0.844790000 | 17 | 6.06627 | 0.74535 | 0.74535 | 17 | settling |
| Castle | 11 | 1998 | PVY | Potyviridae | no | NCNP | no | Solanaceae | no | aphid | 7.89236 | 1.25881 | 1.25881 | 17 | 5.01034 | 0.77846 | 0.77846 | 17 | settling |
| Chaisuekul | 12 | 2005 | TSWV | Tospoviridae | yes | CPP | yes | Solanaceae | no | thrips | 4.89 | 0.81 | 1.62 | 4 | 4.18 | 0.7 | 1.4 | 4 | eggs |
| Chaisuekul | 12 | 2005 | TSWV | Tospoviridae | yes | CPP | yes | Solanaceae | no | thrips | 2.25 | 0.8 | 1.6 | 4 | 2.12 | 0.78 | 1.56 | 4 | eggs |
| Chaisuekul | 12 | 2005 | TSWV | Tospoviridae | yes | CPP | yes | Caryophyllac | no | thrips | 4.86 | 0.35 | 0.7 | 4 | 6.4 | 0.63 | 1.26 | 4 | eggs |
| Chaisuekul | 12 | 2005 | TSWV | Tospoviridae | yes | CPP | yes | Caryophyllac | no | thrips | 0.88 | 0.19 | 0.38 | 4 | 0.88 | 0.36 | 0.72 | 4 | eggs |
| Chen | 13 | 2013 | TYLCV | Geminivirida | yes | CPNP | yes | Solanaceae | yes | Wf | 31.53 | 3.24 | 11.22368923 | 12 | 68.3234 | 3.160599999 | 10.94863956 | 12 | settling |
| Chen | 13 | 2013 | TYLCV | Geminivirida | yes | CPNP | yes | Solanaceae | yes | Wf | 33 | 2.43 | 8.417766924 | 12 | 66.8736 | 2.3464 | 8.128168029 | 12 | settling |
| Chen | 14 | 2014 | WSMoV | Tospoviridae | yes | CPP | yes | Cucurbitacea | no | thrips | 0.001 | 0.001 | 0.001732050 | 3 | 2.32 | 0.33 | 0.571576766 | 3 | settling |
| Chen | 14 | 2014 | WSMoV | Tospoviridae | yes | CPP | yes | Cucurbitacea | no | thrips | 1.34 | 0.65 | 1.125833024 | 3 | 3.37 | 1.18 | 2.043819952 | 3 | settling |
| Chen | 14 | 2014 | WSMoV | Tospoviridae | yes | CPP | yes | Cucurbitacea | no | thrips | 1.37 | 0.64 | 1.108512516 | 3 | 2.38 | 0.92 | 1.593486742 | 3 | settling |
| Chen | 14 | 2014 | WSMoV | Tospoviridae | yes | CPP | yes | Cucurbitacea | no | thrips | 2.75 | 1.19 | 2.061140461 | 3 | 2.4 | 0.89 | 1.541525218 | 3 | settling |
| Chesnais | 15 | 2018 | TuYV | Luteoviridae | yes | CPNP | yes | Brassicaceae | no | aphid | 62.3 | 4.9 | 18.97761839 | 15 | 60.4 | 6.4 | 24.78709341 | 15 | retention |
| Chesnais | 15 | 2018 | TuYV | Luteoviridae | yes | CPNP | yes | Brassicaceae | no | aphid | 16.1 | 3.4 | 13.16814337 | 15 | 19.8 | 3.4 | 13.16814337 | 15 | arrested |
| Chesnais | 15 | 2018 | TuYV | Luteoviridae | yes | CPNP | yes | Brassicaceae | no | aphid | 42.8 | 2.8 | 10.84435336 | 15 | 69 | 5.4 | 20.91411006 | 15 | retention |
| Chesnais | 15 | 2018 | TuYV | Luteoviridae | yes | CPNP | yes | Brassicaceae | no | aphid | 16.4 | 3.5 | 13.55544171 | 15 | 35.3 | 2.1 | 8.133265027 | 15 | arrested |
| Chesnais | 15 | 2018 | TuYV | Luteoviridae | yes | CPNP | yes | Brassicaceae | no | aphid | 50 | 6 | 23.2379000 | 15 | 31.5 | 6.6 | 25.56169008 | 15 | retention |
| Chesnais | 15 | 2018 | TuYV | Luteoviridae | yes | CPNP | yes | Brassicaceae | no | aphid | 12.3 | 3.6 | 13.94274004 | 15 | 3.9 | 1.8 | 6.971370023 | 15 | arrested |
| Chesnais | 15 | 2018 | TuYV | Luteoviridae | yes | CPNP | yes | Brassicaceae | no | aphid | 32.4 | 7 | 27.11088342 | 15 | 21.5 | 5.4 | 20.91411006 | 15 | retention |
| Chesnais | 15 | 2018 | TuYV | Luteoviridae | yes | CPNP | yes | Brassicaceae | no | aphid | 11.7 | 3.8 | 14.71733671 | 15 | 8.1 | 2 | 7.745966692 | 15 | arrested |
| Chesnais | 15 | 2018 | TuYV | Luteoviridae | yes | CPNP | yes | Brassicaceae | no | aphid | 58.35 | 4.731 | 21.15767520 | 20 | 62.4 | 7.314 | 32.70920237 | 20 | EPG |
| Chesnais | 15 | 2018 | TuYV | Luteoviridae | yes | CPNP | yes | Brassicaceae | no | aphid | 51.605 | 6.452 | 28.85422118 | 20 | 117.195 | 14.135 | 63.21364172 | 20 | EPG_E2 |
| Chesnais | 15 | 2018 | TuYV | Luteoviridae | yes | CPNP | yes | Brassicaceae | no | aphid | 103.15 | 10.405 | 46.53257461 | 20 | 127.85 | 10.387 | 46.45207616 | 20 | EPG |
| Chesnais | 15 | 2018 | TuYV | Luteoviridae | yes | CPNP | yes | Brassicaceae | no | aphid | 113.2 | 25.318 | 113.2255381 | 20 | 37.059 | 10.126 | 45.28484868 | 20 | EPG_E2 |
| Chesnais | 15 | 2018 | CaMV | Caulimovirid | no | NCSP | no | Brassicaceae | no | aphid | 69.7 | 4.1 | 15.87923171 | 15 | 66.8 | 4.3 | 16.65382838 | 15 | retention |
| Chesnais | 15 | 2018 | CaMV | Caulimovirid | no | NCSP | no | Brassicaceae | no | aphid | 26.1 | 4.6 | 17.81572339 | 15 | 19.7 | 4 | 15.49193338 | 15 | arrested |
| Chesnais | 15 | 2018 | CaMV | Caulimovirid | no | NCSP | no | Brassicaceae | no | aphid | 50.3 | 5.7 | 22.07600507 | 15 | 50.3 | 5.9 | 22.85060174 | 15 | retention |
| Chesnais | 15 | 2018 | CaMV | Caulimovirid | no | NCSP | no | Brassicaceae | no | aphid | 45.8 | 6.1 | 23.62519841 | 15 | 48.3 | 5.8 | 22.46330340 | 15 | arrested |
| Chesnais | 15 | 2018 | CaMV | Caulimovirid | no | NCSP | no | Brassicaceae | no | aphid | 31.7 | 3.5 | 13.55544171 | 15 | 39.2 | 4.5 | 17.42842505 | 15 | retention |
| Chesnais | 15 | 2018 | CaMV | Caulimovirid | no | NCSP | no | Brassicaceae | no | aphid | 12.1 | 3.6 | 13.94274004 | 15 | 11.2 | 1.7 | 6.584071688 | 15 | arrested |
| Chesnais | 15 | 2018 | CaMV | Caulimovirid | no | NCSP | no | Brassicaceae | no | aphid | 32 | 4.9 | 18.97761839 | 15 | 30.9 | 4 | 15.49193338 | 15 | retention |
| Chesnais | 15 | 2018 | CaMV | Caulimovirid | no | NCSP | no | Brassicaceae | no | aphid | 11.1 | 3.2 | 12.39354670 | 15 | 4.5 | 1.7 | 6.584071688 | 15 | arrested |
| Chesnais | 15 | 2018 | CaMV | Caulimovirid | no | NCSP | no | Brassicaceae | no | aphid | 58.35 | 4.731 | 21.15767520 | 20 | 98.5 | 10.548 | 47.17209005 | 20 | EPG |
| Chesnais | 15 | 2018 | CaMV | Caulimovirid | no | NCSP | no | Brassicaceae | no | aphid | 51.605 | 6.452 | 28.85422118 | 20 | 41.457 | 19.107 | 85.44910169 | 20 | EPG_E2 |
| Chesnais | 15 | 2018 | CaMV | Caulimovirid | no | NCSP | no | Brassicaceae | no | aphid | 103.15 | 10.405 | 46.53257461 | 20 | 111 | 14.167 | 63.35675007 | 20 | EPG |
| Chesnais | 15 | 2018 | CaMV | Caulimovirid | no | NCSP | no | Brassicaceae | no | aphid | 113.2 | 25.318 | 113.2255381 | 20 | 86.767 | 29.741 | 133.0057954 | 20 | EPG_E2 |
| Chesnais | 65 | 2019 | TuYV | Luteoviridae | yes | CPNP | yes | Brassicaceae | no | aphid | 53.59 | 4.359 | 16.88233440 | 15 | 50.5 | 2.915 | 11.28974645 | 15 | retention |
| Chesnais | 65 | 2019 | TuYV | Luteoviridae | yes | CPNP | yes | Brassicaceae | no | aphid | 31.83 | 2.812 | 10.89082916 | 15 | 27.07 | 3.89 | 15.06590521 | 15 | arrested |
| Chesnais | 65 | 2019 | TuYV | Luteoviridae | yes | CPNP | yes | Brassicaceae | no | aphid | 57.19 | 8.164 | 31.61903603 | 15 | 79.31 | 5.012 | 19.41139253 | 15 | retention |
| Chesnais | 65 | 2019 | TuYV | Luteoviridae | yes | CPNP | yes | Brassicaceae | no | aphid | 14.96 | 3.671 | 14.21772186 | 15 | 22.65 | 5.587 | 21.63835795 | 15 | arrested |
| Chesnais | 65 | 2019 | TuYV | Luteoviridae | yes | CPNP | yes | Brassicaceae | yes | aphid | 39.67 | 4.88 | 18.90015872 | 15 | 49.3 | 4.6 | 17.81572339 | 15 | retention |
| Chesnais | 65 | 2019 | TuYV | Luteoviridae | yes | CPNP | yes | Brassicaceae | yes | aphid | 17.31 | 3.38 | 13.09068371 | 15 | 19.83 | 3.83 | 14.83352621 | 15 | arrested |
| Chesnais | 65 | 2019 | TuYV | Luteoviridae | yes | CPNP | yes | Brassicaceae | no | aphid | 37.907 | 4.1757 | 16.17241655 | 15 | 64.97 | 3.636 | 14.08216744 | 15 | retention |
| Chesnais | 65 | 2019 | TuYV | Luteoviridae | yes | CPNP | yes | Brassicaceae | no | aphid | 28.6 | 3.75 | 14.52368754 | 15 | 50.87 | 4.949 | 19.16739458 | 15 | arrested |
| Chesnais | 65 | 2019 | TuYV | Luteoviridae | yes | CPNP | yes | Brassicaceae | no | aphid | 48.7 | 5.688 | 22.02952927 | 15 | 57.69 | 5.918 | 22.92031544 | 15 | retention |
| Chesnais | 65 | 2019 | TuYV | Luteoviridae | yes | CPNP | yes | Brassicaceae | no | aphid | 34.17 | 4.668 | 18.07908626 | 15 | 42.96 | 6.51 | 25.21312158 | 15 | arrested |
| Chesnais | 65 | 2019 | TuYV | Luteoviridae | yes | CPNP | yes | Brassicaceae | yes | aphid | 27 | 4.2 | 16.26653005 | 15 | 50 | 6.02 | 23.31535974 | 15 | retention |
| Chesnais | 65 | 2019 | TuYV | Luteoviridae | yes | CPNP | yes | Brassicaceae | yes | aphid | 22 | 3.5 | 13.55544171 | 15 | 43 | 4.2 | 16.26653005 | 15 | arrested |

### Settling/feeding preference database continued

| | | | | | | | | | | | | | | | | | | | |
|---|---|---|---|---|---|---|---|---|---|---|---|---|---|---|---|---|---|---|---|
| Daimei | 16 | 2017 | GBNV | Tospoviridae | yes | CPP | yes | Fabaceae | no | thrips | 4.239 | 1.041 | 2.082 | 4 | 6.99 | 0.37 | 0.74 | 4 | settling |
| Daimei | 16 | 2017 | GBNV | Tospoviridae | yes | CPP | yes | Fabaceae | no | thrips | 6.256 | 0.564 | 1.128 | 4 | 10 | 0.872 | 1.744 | 4 | settling |
| Daimei | 16 | 2017 | GBNV | Tospoviridae | yes | CPP | yes | Fabaceae | no | thrips | 2.97 | 0.94 | 1.88 | 4 | 8.273 | 0.375999999 | 0.751999999 | 4 | settling |
| Daimei | 16 | 2017 | GBNV | Tospoviridae | yes | CPP | yes | Fabaceae | no | thrips | 4.7 | 1.5 | 3 | 4 | 11.28 | 1.25 | 2.5 | 4 | settling |
| Daimei | 16 | 2017 | GBNV | Tospoviridae | yes | CPP | yes | Fabaceae | no | thrips | 3.99 | 0.8 | 1.6 | 4 | 8.22 | 0.639999999 | 1.28 | 4 | settling |
| Daimei | 16 | 2017 | GBNV | Tospoviridae | yes | CPP | yes | Fabaceae | no | thrips | 6.26 | 1.03 | 2.06 | 4 | 9.73 | 0.65 | 1.3 | 4 | settling |
| Daimei | 16 | 2017 | GBNV | Tospoviridae | yes | CPP | yes | Solanaceae | no | thrips | 3.97 | 0.57 | 1.14 | 4 | 8.22 | 0.789999999 | 1.58 | 4 | settling |
| Daimei | 16 | 2017 | GBNV | Tospoviridae | yes | CPP | yes | Solanaceae | no | thrips | 6.29 | 0.61 | 1.22 | 4 | 9.77 | 0.83 | 1.66 | 4 | settling |
| Eigenbrode | 17 | 2002 | PLRV | Luteoviridae | yes | CPNP | yes | Solanaceae | no | aphid | 30.3131 | 3.2474 | 8.591812807 | 7 | 69.6869 | 2.582399999 | 6.832388185 | 7 | settling |
| Fang | 18 | 2013 | TYLCV | Geminivirida | yes | CPNP | yes | Solanaceae | no | Wf | 71.68 | 4.559999999 | 12.89762768 | 8 | 28.17 | 4.15 | 11.73797256 | 8 | settling |
| Fang | 18 | 2013 | TYLCV | Geminivirida | yes | CPNP | yes | Solanaceae | no | Wf | 80.9392 | 3.5912 | 10.15744749 | 8 | 19.337 | 3.1768 | 8.985347289 | 8 | settling |
| Fang | 18 | 2013 | TYLCV | Geminivirida | yes | CPNP | yes | Solanaceae | no | Wf | 35.4167 | 7.9133 | 22.38219236 | 8 | 64.3056 | 7.7744 | 21.98932383 | 8 | settling |
| Fang | 18 | 2013 | TYLCV | Geminivirida | yes | CPNP | yes | Solanaceae | no | Wf | 26.5278 | 3.1922 | 9.028905067 | 8 | 73.75 | 2.7778 | 7.856804867 | 8 | settling |
| Fereres | 19 | 1990 | BYDV | Luteoviridae | yes | CPNP | yes | Poaceae | no | aphid | 67.2 | 13.06 | 45.24116709 | 12 | 65.7 | 11.01 | 38.13975878 | 12 | EPG_E2 |
| Fereres | 19 | 1990 | BYDV | Luteoviridae | yes | CPNP | yes | Poaceae | no | aphid | 83.5 | 10.92 | 37.82798963 | 12 | 61.9 | 11.48 | 39.76788654 | 12 | EPG_E2 |
| Fereres | 19 | 1990 | BYDV | Luteoviridae | yes | CPNP | yes | Poaceae | no | aphid | 78.9 | 10.86 | 37.62014354 | 12 | 93.1 | 12.82 | 44.40978270 | 12 | EPG_E2 |
| Fereres | 19 | 1990 | BYDV | Luteoviridae | yes | CPNP | yes | Poaceae | no | aphid | 67.2 | 13.06 | 45.24116709 | 12 | 77.2 | 11.83 | 40.98032210 | 12 | EPG_E2 |
| Fereres | 19 | 1990 | BYDV | Luteoviridae | yes | CPNP | yes | Poaceae | no | aphid | 83.5 | 10.92 | 37.82798963 | 12 | 72.3 | 13.87 | 48.04708940 | 12 | EPG_E2 |
| Fereres | 19 | 1990 | BYDV | Luteoviridae | yes | CPNP | yes | Poaceae | no | aphid | 78.9 | 10.86 | 37.62014354 | 12 | 60.8 | 16.01 | 55.46026685 | 12 | EPG_E2 |
| Fereres | 19 | 1990 | BYDV | Luteoviridae | yes | CPNP | yes | Poaceae | no | aphid | 7.2 | 1.19 | 4.122280922 | 12 | 9.4 | 1 | 3.464101615 | 12 | EPG |
| Fereres | 19 | 1990 | BYDV | Luteoviridae | yes | CPNP | yes | Poaceae | no | aphid | 8.2 | 2.15 | 7.447818472 | 12 | 7.1 | 0.57 | 1.974537920 | 12 | EPG |
| Fereres | 19 | 1990 | BYDV | Luteoviridae | yes | CPNP | yes | Poaceae | no | aphid | 6.1 | 0.99 | 3.429460598 | 12 | 7.3 | 1.14 | 3.949075841 | 12 | EPG |
| Fereres | 19 | 1990 | BYDV | Luteoviridae | yes | CPNP | yes | Poaceae | no | aphid | 7.2 | 1.19 | 4.122280922 | 12 | 9.2 | 0.89 | 3.083050437 | 12 | EPG |
| Fereres | 19 | 1990 | BYDV | Luteoviridae | yes | CPNP | yes | Poaceae | no | aphid | 8.2 | 2.15 | 7.447818472 | 12 | 7.3 | 1.22 | 4.226203970 | 12 | EPG |
| Fereres | 19 | 1990 | BYDV | Luteoviridae | yes | CPNP | yes | Poaceae | no | aphid | 6.1 | 0.99 | 3.429460598 | 12 | 8.2 | 1.21 | 4.191562954 | 12 | EPG |
| Fereres | 20 | 2016 | ToSRV | Geminivirida | yes | CPNP | yes | Solanaceae | no | Wf | 3.05 | 0.52 | 2.325510696 | 20 | 0.8 | 0.27 | 1.207476707 | 20 | settling |
| Fereres | 20 | 2016 | ToCV | Closteroviric | no | NCSP | yes | Solanaceae | no | Wf | 2.6 | 0.61 | 2.728002932 | 20 | 4 | 0.92 | 4.114365078 | 20 | settling |
| Fereres | 21 | 1999 | SMV | Potyviridae | no | NCNP | no | Fabaceae | no | aphid | 9.37 | 0.24 | 0.96 | 16 | 8.38 | 0.289999999 | 1.16 | 16 | retention |
| Fereres | 21 | 1999 | SMV | Potyviridae | no | NCNP | no | Fabaceae | no | aphid | 7.49 | 0.48 | 1.92 | 16 | 6.31 | 0.33 | 1.32 | 16 | retention |
| Gadhave | 70 | 2019 | PRSV | Potyviridae | no | NCNP | no | Cucurbitacea | no | aphid | 1.88567 | 0.45221 | 1.430013580 | 10 | 0.204778 | 0.132253 | 0.418220707 | 10 | emigration |
| Gadhave | 70 | 2019 | PRSV | Potyviridae | no | NCNP | no | Cucurbitacea | no | aphid | 11.6625 | 2.0095 | 6.354596958 | 10 | 9.62298 | 1.86715 | 5.904446733 | 10 | arrested |
| Ghosh | 22 | 2016 | CBDV | Nanoviridae | yes | CPNP | yes | Zingiberacea | no | aphid | 5.5 | 0.4 | 1.264911064 | 10 | 32 | 1.36 | 4.300697617 | 10 | settling |
| Guo | 68 | 2019 | CMV | Bromovirida | no | NCNP | no | Cucurbitacea | no | aphid | 3005 | 615 | 3368.493728 | 30 | 5390 | 542 | 2968.656261 | 30 | EPG |
| Guo | 68 | 2019 | CMV | Bromovirida | no | NCNP | no | Cucurbitacea | no | aphid | 1667 | 388 | 2125.163523 | 30 | 62.9 | 57.68 | 315.9263711 | 30 | EPG_E2 |
| Hily | 23 | 2014 | CMV | Bromovirida | no | NCNP | no | Brassicaceae | no | aphid | 5.33 | 1.22 | 5.456005865 | 20 | 3.42 | 0.8 | 3.577708763 | 20 | settling |
| Hily | 23 | 2014 | CMV | Bromovirida | no | NCNP | no | Brassicaceae | no | aphid | 5.48 | 0.93 | 4.159086438 | 20 | 3.27 | 76 | 339.8823325 | 20 | settling |
| Hodge | 24 | 2008 | PEMV | Luteoviridae | yes | CPNP | yes | Fabaceae | no | aphid | 60 | 30.4 | 30.4 | 10 | 40 | 30.4 | 30.4 | 10 | settling |
| Hodge | 24 | 2008 | PEMV | Luteoviridae | yes | CPNP | yes | Fabaceae | no | aphid | 50.4 | 31 | 31 | 10 | 49.6 | 31 | 31 | 10 | settling |
| Hodge | 24 | 2008 | PEMV | Luteoviridae | yes | CPNP | yes | Fabaceae | no | aphid | 36.2 | 18.8 | 18.8 | 25 | 63.8 | 18.8 | 18.8 | 25 | settling |
| Hodge | 24 | 2008 | PEMV | Luteoviridae | yes | CPNP | yes | Fabaceae | no | aphid | 44.7 | 17.8 | 17.8 | 30 | 55.3 | 17.8 | 17.8 | 30 | settling |
| Hodge | 24 | 2008 | PEMV | Luteoviridae | yes | CPNP | yes | Fabaceae | no | aphid | 38.1 | 17.4 | 17.4 | 30 | 61.9 | 17.4 | 17.4 | 30 | settling |
| Hodge | 24 | 2008 | PEMV | Luteoviridae | yes | CPNP | yes | Fabaceae | no | aphid | 10.6 | 19.1 | 19.1 | 10 | 89.4 | 19.1 | 19.1 | 10 | settling |
| Hodge | 24 | 2008 | PEMV | Luteoviridae | yes | CPNP | yes | Fabaceae | no | aphid | 25.9 | 17.5 | 17.5 | 24 | 74.1 | 17.5 | 17.5 | 24 | settling |
| Hodge | 24 | 2008 | PEMV | Luteoviridae | yes | CPNP | yes | Fabaceae | no | aphid | 20 | 11.1 | 11.1 | 50 | 80 | 11.1 | 11.1 | 50 | settling |
| Hodge | 24 | 2008 | PEMV | Luteoviridae | yes | CPNP | yes | Fabaceae | no | aphid | 15.6 | 13 | 13 | 30 | 84.4 | 13 | 13 | 30 | settling |
| Hodge | 24 | 2008 | BYMV | Potyviridae | no | NCNP | no | Fabaceae | no | aphid | 62.7 | 30 | 30 | 10 | 37.3 | 30 | 30 | 10 | settling |
| Hodge | 24 | 2008 | BYMV | Potyviridae | no | NCNP | no | Fabaceae | no | aphid | 48.3 | 31 | 31 | 10 | 51.7 | 31 | 31 | 10 | settling |
| Hodge | 24 | 2008 | BYMV | Potyviridae | no | NCNP | no | Fabaceae | no | aphid | 34.9 | 18.7 | 18.7 | 25 | 65.1 | 18.7 | 18.7 | 25 | settling |
| Hodge | 24 | 2008 | BYMV | Potyviridae | no | NCNP | no | Fabaceae | no | aphid | 45.6 | 17.8 | 17.8 | 30 | 54.4 | 17.8 | 17.8 | 30 | settling |
| Hodge | 24 | 2008 | BYMV | Potyviridae | no | NCNP | no | Fabaceae | no | aphid | 53.1 | 17.9 | 17.9 | 30 | 46.9 | 17.9 | 17.9 | 30 | settling |
| Hodge | 24 | 2008 | BYMV | Potyviridae | no | NCNP | no | Fabaceae | no | aphid | 13.9 | 21.4 | 21.4 | 10 | 86.1 | 21.4 | 21.4 | 10 | settling |
| Hodge | 24 | 2008 | BYMV | Potyviridae | no | NCNP | no | Fabaceae | no | aphid | 30.9 | 18.5 | 18.5 | 24 | 69.1 | 18.5 | 18.5 | 24 | settling |
| Hodge | 24 | 2008 | BYMV | Potyviridae | no | NCNP | no | Fabaceae | no | aphid | 13 | 9.3 | 9.3 | 50 | 87 | 9.3 | 9.3 | 50 | settling |
| Hodge | 24 | 2008 | BYMV | Potyviridae | no | NCNP | no | Fabaceae | no | aphid | 21.7 | 14.8 | 14.8 | 30 | 78.3 | 14.8 | 14.8 | 30 | settling |
| Hodge | 25 | 2011 | BYMV | Potyviridae | no | NCNP | no | Fabaceae | no | aphid | 23.222 | 3.842 | 11.526 | 9 | 54.296 | 3.675 | 11.025 | 9 | settling |
| Jimenez | 26 | 2004 | BYDV | Luteoviridae | yes | CPNP | yes | Poaceae | no | aphid | 11 | 1.17 | 3.699864862 | 10 | 15.2 | 1.45 | 4.585302607 | 10 | settling |
| Jimenez | 26 | 2004 | BYDV | Luteoviridae | yes | CPNP | yes | Poaceae | no | aphid | 10.8 | 1.1 | 3.478505426 | 10 | 15 | 1.54 | 4.869907596 | 10 | settling |
| Keough | 27 | 2016 | SVNV | Tospoviridae | yes | CPP | yes | Fabaceae | no | thrips | 1.774 | 0.362 | 1.618913215 | 20 | 1.84977 | 0.37923 | 1.695968118 | 20 | settling |
| Keough | 27 | 2016 | SVNV | Tospoviridae | yes | CPP | yes | Fabaceae | no | thrips | 3.236 | 0.362 | 1.618913215 | 20 | 1.394 | 0.344 | 1.538414768 | 20 | settling |
| Kliot | 69 | 2019 | TYLCV | Geminivirida | yes | CPNP | yes | Solanaceae | no | Wf | 66.66 | 7.91 | 25.01361629 | 10 | 33.34 | 7.81 | 24.69738852 | 10 | settling |
| Kliot | 69 | 2019 | TYLCV | Geminivirida | yes | CPNP | yes | Solanaceae | no | Wf | 40.26 | 5.760000000 | 18.21471932 | 10 | 59.43 | 5.55 | 17.55064101 | 10 | settling |
| Legarrea | 28 | 2015 | TYLCV | Geminivirida | yes | CPNP | yes | Solanaceae | no | Wf | 47 | 6.26 | 24.24487574 | 15 | 53.349 | 6.4952 | 25.15580143 | 15 | settling |
| Legarrea | 28 | 2015 | TYLCV | Geminivirida | yes | CPNP | yes | Solanaceae | no | Wf | 49.95 | 6.9849 | 27.05240137 | 15 | 50.4 | 6.989 | 27.06828060 | 15 | settling |
| Legarrea | 28 | 2015 | TYLCV | Geminivirida | yes | CPNP | yes | Solanaceae | no | Wf | 42.5948 | 2.641 | 10.22854901 | 15 | 57.4052 | 2.5248 | 9.778508352 | 15 | settling |
| Legarrea | 28 | 2015 | TYLCV | Geminivirida | yes | CPNP | yes | Solanaceae | no | Wf | 45.9532 | 2.885 | 11.17355695 | 15 | 54.0254 | 2.538 | 9.829631732 | 15 | settling |
| Legarrea | 28 | 2015 | TYLCV | Geminivirida | yes | CPNP | yes | Solanaceae | no | Wf | 47.88 | 4.5988 | 17.81107581 | 15 | 52.7207 | 5.4413 | 21.07406428 | 15 | settling |
| Legarrea | 28 | 2015 | TYLCV | Geminivirida | yes | CPNP | yes | Solanaceae | no | Wf | 49.4559 | 4.474 | 17.32772749 | 15 | 51.5115 | 5.3185 | 20.59846192 | 15 | settling |
| Lei | 29 | 2016 | SRBSDV | Reoviridae | yes | CPP | yes | Poaceae | no | PH | 133.26 | 6.36 | 37.08485405 | 34 | 182.52 | 4.82 | 28.10518813 | 34 | EPG_E2 |
| Lei | 29 | 2016 | SRBSDV | Reoviridae | yes | CPP | yes | Poaceae | no | PH | 12.25 | 1.57 | 9.154594474 | 34 | 8.78 | 0.67 | 3.906737769 | 34 | EPG |
| Lightle | 30 | 2014 | RpLV | Reoviridae | yes | CPP | yes | Rosaceae | no | aphid | 51.72 | 5.38 | 21.52 | 16 | 47.17 | 4.97 | 19.88 | 16 | settling |
| Lightle | 30 | 2014 | RLMV | Closteroviric | no | NCSP | yes | Rosaceae | no | aphid | 43.72 | 5.24 | 20.96 | 16 | 53.79 | 5.52 | 22.08 | 16 | settling |
| Lightle | 30 | 2014 | RpLV | Reoviridae | yes | CPP | yes | Rosaceae | no | aphid | 60.6679 | 5.658599999 | 22.6344 | 16 | 38.961 | 5.5659 | 22.2636 | 16 | settling |
| Lightle | 30 | 2014 | RLMV | Closteroviric | no | NCSP | yes | Rosaceae | no | aphid | 41.1874 | 5.2876 | 21.1504 | 16 | 58.7199 | 5.5658 | 22.2632 | 16 | settling |
| Lightle | 30 | 2014 | RLMV | Closteroviric | no | NCSP | yes | Rosaceae | no | aphid | 872.1 | 467.5 | 1927.551879 | 17 | 665.3 | 316.9 | 1381.335075 | 19 | EPG_E2 |
| Lightle | 30 | 2014 | RLMV | Closteroviric | no | NCSP | yes | Rosaceae | no | aphid | 257.7 | 27.7 | 114.2100258 | 17 | 282.6 | 26.2 | 114.2031523 | 19 | EPG |
| Lightle | 30 | 2014 | RLMV | Closteroviric | no | NCSP | yes | Rosaceae | no | aphid | 284.2 | 30.6 | 126.1670321 | 17 | 352.7 | 28.9 | 125.9721794 | 19 | EPG |
| Liu | 31 | 2013 | TYLCV | Geminivirida | yes | CPNP | yes | Solanaceae | no | Wf | 27.96 | 9.4268 | 50.76487160 | 29 | 28.8477 | 4.4589 | 24.01191135 | 29 | EPG_E2 |
| Liu | 31 | 2013 | TYLCV | Geminivirida | yes | CPNP | yes | Solanaceae | no | Wf | 63.359 | 9.8055 | 53.70693537 | 30 | 75.3305 | 11.2051 | 60.34131018 | 29 | EPG_E2 |
| Liu | 31 | 2013 | TYLCV | Geminivirida | yes | CPNP | yes | Solanaceae | no | Wf | 76.79 | 11.41 | 61.44473044 | 29 | 110.37 | 12.54 | 67.52996668 | 29 | EPG |
| Liu | 31 | 2013 | TYLCV | Geminivirida | yes | CPNP | yes | Solanaceae | no | Wf | 50.37 | 8.06 | 44.14643813 | 30 | 53.51 | 7.6 | 40.92725253 | 29 | EPG |
| Lu | 32 | 2016 | SRBSDV | Reoviridae | yes | CPP | yes | Poaceae | no | PH | 68.98 | 2.386399999 | 4.133366047 | 3 | 31.0193 | 2.3859 | 4.132500021 | 3 | settling |
| Lu | 32 | 2016 | SRBSDV | Reoviridae | yes | CPP | yes | Poaceae | no | PH | 72.135 | 1.988199999 | 3.443663415 | 3 | 28.0633 | 2.1871 | 3.788168321 | 3 | settling |
| Lu | 32 | 2016 | SRBSDV | Reoviridae | yes | CPP | yes | Poaceae | no | PH | 65.9461 | 3.179599999 | 5.507228747 | 3 | 33.8545 | 2.1875 | 3.788861141 | 3 | settling |
| Lu | 32 | 2016 | SRBSDV | Reoviridae | yes | CPP | yes | Poaceae | no | PH | 52.99 | 2.39 | 4.139601430 | 3 | 46.8 | 2.19 | 3.793191268 | 3 | settling |
| Lu | 32 | 2016 | SRBSDV | Reoviridae | yes | CPP | yes | Poaceae | no | PH | 30.5057 | 6.9543 | 12.04520093 | 3 | 68.699 | 6.161 | 10.67116502 | 3 | settling |
| Lu | 32 | 2016 | SRBSDV | Reoviridae | yes | CPP | yes | Poaceae | no | PH | 26.1055 | 2.1845 | 3.783664989 | 3 | 73.89 | 2.19 | 3.793191268 | 3 | settling |
| Lu | 32 | 2016 | SRBSDV | Reoviridae | yes | CPP | yes | Poaceae | no | PH | 54.906 | 7.554 | 13.08391180 | 3 | 45.09 | 7.558699999 | 13.09205243 | 3 | settling |
| Lu | 32 | 2016 | SRBSDV | Reoviridae | yes | CPP | yes | Poaceae | no | PH | 75.9534 | 4.566599999 | 7.909583217 | 3 | 23.45 | 4.77 | 8.261882352 | 3 | settling |
| Lu | 32 | 2016 | RRSV | Reoviridae | yes | CPP | yes | Poaceae | no | PH | 69.69 | 1.8 | 3.117691453 | 3 | 31.6 | 3.3898 | 5.871305827 | 3 | settling |
| Lu | 32 | 2016 | RRSV | Reoviridae | yes | CPP | yes | Poaceae | no | PH | 64.7361 | 0.993900000 | 1.721485297 | 3 | 35.96 | 1.398 | 2.421407028 | 3 | settling |
| Lu | 32 | 2016 | RRSV | Reoviridae | yes | CPP | yes | Poaceae | no | PH | 68.948 | 4.582000000 | 7.936256800 | 3 | 31.55 | 3.98 | 6.893562214 | 3 | settling |
| Lu | 32 | 2016 | RRSV | Reoviridae | yes | CPP | yes | Poaceae | no | PH | 63.58 | 3.989999999 | 6.910882722 | 3 | 36.71 | 3.7881 | 6.561181664 | 3 | settling |
| Lu | 32 | 2016 | RRSV | Reoviridae | yes | CPP | yes | Poaceae | no | PH | 44.07 | 3.19 | 5.525242076 | 3 | 56.026 | 2.991899999 | 5.182122811 | 3 | settling |
| Lu | 32 | 2016 | RRSV | Reoviridae | yes | CPP | yes | Poaceae | no | PH | 28.345 | 2.785 | 4.823761499 | 3 | 71.156 | 1.994 | 3.453709310 | 3 | settling |
| Lu | 32 | 2016 | RRSV | Reoviridae | yes | CPP | yes | Poaceae | no | PH | 37.74 | 3.19 | 5.525242076 | 3 | 63.15 | 2.5977 | 4.499348382 | 3 | settling |
| Lu | 32 | 2016 | RRSV | Reoviridae | yes | CPP | yes | Poaceae | no | PH | 77.049 | 2.991 | 5.180563965 | 3 | 22.85 | 2.99 | 5.178831914 | 3 | settling |
| Macias | 33 | 1969 | BYV | Closteroviric | no | NCSP | no | Amaranthace | yes | aphid | 24 | 29.6 | 29.6 | 8 | 50 | 34.6 | 34.6 | 8 | settling |
| Macias | 33 | 1969 | BYV | Closteroviric | no | NCSP | no | Amaranthace | yes | aphid | 17 | 26 | 26 | 8 | 43 | 34.3 | 34.3 | 8 | settling |
| Macias | 33 | 1969 | SBMV | Potyviridae | no | NCNP | no | Amaranthace | no | aphid | 24 | 29.6 | 29.6 | 8 | 26 | 30.4 | 30.4 | 8 | settling |
| Macias | 33 | 1969 | SBMV | Potyviridae | no | NCNP | no | Amaranthace | no | aphid | 17 | 26 | 26 | 8 | 25 | 30 | 30 | 8 | settling |
| Maluta | 34 | 2017 | ToCV | Closteroviric | no | NCSP | no | Solanaceae | yes | Wf | 32.96 | 3.15 | 9.961174629 | 10 | 7.309 | 0.855 | 2.703747399 | 10 | settling |
| Maluta | 34 | 2017 | ToCV | Closteroviric | no | NCSP | no | Solanaceae | yes | Wf | 32.059 | 2.721 | 8.604557513 | 10 | 11.164 | 1.336 | 4.224802953 | 10 | settling |
| Maluta | 34 | 2017 | ToSRV | Geminivirida | yes | CPNP | yes | Solanaceae | no | Wf | 26.237 | 3.093 | 9.780924802 | 10 | 19.1429 | 3.2261 | 10.20182395 | 10 | settling |
| Maluta | 34 | 2017 | ToSRV | Geminivirida | yes | CPNP | yes | Solanaceae | no | Wf | 42.638 | 4.412 | 13.95196903 | 10 | 27.64 | 3.42 | 10.81498959 | 10 | settling |
| Maluta | 35 | 2017 | ToCV | Closteroviric | no | NCSP | no | Solanaceae | yes | Wf | 16.42 | 8.51 | 40.81252626 | 23 | 22.45 | 12.9 | 61.86622665 | 23 | EPG_E2 |
| Maluta | 35 | 2017 | ToCV | Closteroviric | no | NCSP | no | Solanaceae | yes | Wf | 2.49 | 0.67 | 3.213207120 | 23 | 3.27 | 1.14 | 5.467247936 | 23 | EPG |
| Maluta | 35 | 2017 | ToSRV | Geminivirida | yes | CPNP | yes | Solanaceae | no | Wf | 23.93 | 10.89 | 51.07862762 | 22 | 6.18 | 3.25 | 15.24385121 | 22 | EPG_E2 |
| Maluta | 35 | 2017 | ToSRV | Geminivirida | yes | CPNP | yes | Solanaceae | no | Wf | 3.44 | 1.12 | 5.253265651 | 22 | 5.92 | 2.35 | 11.02247703 | 22 | EPG |

### Settling/feeding preference database continued

| | | | | | | | | | | | | | | | | | | | |
|---|---|---|---|---|---|---|---|---|---|---|---|---|---|---|---|---|---|---|---|
| Maris | 36 | 2004 | TSWV | Tospoviridae | yes | CPP | yes | Solanaceae | no | thrips | 0.2 | 0.2 | 0.692820323 | 12 | 4 | 1.2 | 4.156921938 | 12 | settling |
| Maris | 36 | 2004 | TSWV | Tospoviridae | yes | CPP | yes | Solanaceae | no | thrips | 0.5 | 0.1 | 0.346410161 | 12 | 5.2 | 0.9 | 3.117691453 | 12 | settling |
| Maris | 36 | 2004 | TSWV | Tospoviridae | yes | CPP | yes | Solanaceae | no | thrips | 0.3 | 0.2 | 0.692820323 | 12 | 4.5 | 0.8 | 2.771281292 | 12 | settling |
| Maris | 36 | 2004 | TSWV | Tospoviridae | yes | CPP | yes | Solanaceae | yes | thrips | 1.3 | 0.4 | 1.385640646 | 12 | 6.5 | 0.5 | 1.732050807 | 12 | settling |
| Mauck | 37 | 2010 | CMV | Bromovirida | no | NCNP | no | Cucurbitacea | no | aphid | 38.5 | 2.59 | 5.18 | 4 | 28.15 | 2.75 | 5.5 | 4 | retention |
| Mauck | 37 | 2010 | CMV | Bromovirida | no | NCNP | no | Cucurbitacea | no | aphid | 42.49 | 2.82 | 5.64 | 4 | 17.61 | 2.13 | 4.26 | 4 | retention |
| Mauck | 37 | 2010 | CMV | Bromovirida | no | NCNP | no | Cucurbitacea | no | aphid | 0.61 | 0.19 | 0.38 | 4 | 0.26 | 0.25 | 0.5 | 4 | arrested |
| Mauck | 37 | 2010 | CMV | Bromovirida | no | NCNP | no | Cucurbitacea | no | aphid | 18.2 | 1.64 | 3.28 | 4 | 4.15 | 1.98 | 3.96 | 4 | arrested |
| Mauck | 38 | 2014 | CMV | Bromovirida | no | NCNP | no | Cucurbitacea | no | aphid | 25.96 | 1.09 | 2.437314095 | 5 | 21.44 | 1.64 | 3.667151483 | 5 | retention |
| Mauck | 38 | 2014 | CMV | Bromovirida | no | NCNP | no | Cucurbitacea | no | aphid | 27.17 | 1 | 2.236067977 | 5 | 16.01 | 2.09 | 4.673382072 | 5 | retention |
| Mauck | 38 | 2014 | CMV | Bromovirida | no | NCNP | no | Cucurbitacea | no | aphid | 17.29 | 2.42 | 7.26 | 9 | 15.07 | 0.57 | 1.71 | 9 | retention |
| Mauck | 38 | 2014 | CMV | Bromovirida | no | NCNP | no | Cucurbitacea | no | aphid | 4.58 | 0.91 | 2.73 | 9 | 4.7 | 1.12 | 3.36 | 9 | retention |
| Mauck | 38 | 2014 | CMV | Bromovirida | no | NCNP | no | Solanaceae | no | aphid | 24.84 | 1.24 | 3.72 | 9 | 28.47 | 0.790000000 | 2.370000000 | 9 | retention |
| Mauck | 38 | 2014 | CMV | Bromovirida | no | NCNP | no | Solanaceae | no | aphid | 26.46 | 0.919999999 | 2.759999999 | 9 | 29.49 | 0.790000000 | 2.370000000 | 9 | retention |
| Mauck | 38 | 2014 | CMV | Bromovirida | no | NCNP | no | Solanaceae | no | aphid | 19.02 | 1.94 | 3.88 | 4 | 17.02 | 1.24 | 2.48 | 4 | retention |
| Mauck | 38 | 2014 | CMV | Bromovirida | no | NCNP | no | Solanaceae | no | aphid | 11.02 | 1.53 | 3.06 | 4 | 11.98 | 2.48 | 4.96 | 4 | retention |
| Mauck | 38 | 2014 | CMV | Bromovirida | no | NCNP | no | Solanaceae | no | aphid | 28.73 | 1.59 | 3.555348084 | 5 | 27.91 | 1.72 | 3.846036921 | 5 | retention |
| Mauck | 38 | 2014 | CMV | Bromovirida | no | NCNP | no | Solanaceae | no | aphid | 28.97 | 1.21 | 2.705642252 | 5 | 26.83 | 2.02 | 4.516857314 | 5 | retention |
| Medina | 39 | 2009 | BYDV | Luteoviridae | yes | CPNP | yes | Poaceae | no | aphid | 1.17 | 0.42 | 1.454922678 | 12 | 1.08 | 0.29 | 1.004589468 | 12 | settling |
| Montlor | 41 | 1986 | BYDV_RPV | Luteoviridae | yes | CPNP | yes | Poaceae | no | aphid | 11.39 | 1.12 | 5.132484778 | 21 | 9.75 | 0.890000000 | 4.268290055 | 23 | EPG |
| Montlor | 41 | 1986 | BYDV_RPV | Luteoviridae | yes | CPNP | yes | Poaceae | no | aphid | 106.71 | 16.782 | 76.90478531 | 21 | 206.471 | 14.904 | 71.47707302 | 23 | EPG_E2 |
| Montlor | 41 | 1986 | BYDV_RPV | Luteoviridae | yes | CPNP | yes | Poaceae | no | aphid | 6.05 | 0.91 | 4.268278341 | 22 | 5.07 | 0.51 | 2.392112037 | 22 | EPG |
| Montlor | 41 | 1986 | BYDV_RPV | Luteoviridae | yes | CPNP | yes | Poaceae | no | aphid | 280.669 | 24.283 | 113.8973658 | 22 | 324.75 | 12.95 | 60.74088408 | 22 | EPG_E2 |
| Murphy | 42 | 2018 | PVY | Potyviridae | no | NCNP | no | Amaranthac | yes | aphid | 24.75 | 3.1 | 5.369357503 | 3 | 23.47 | 5.2 | 9.006664199 | 3 | settling |
| Murphy | 42 | 2018 | PVY | Potyviridae | no | NCNP | no | Amaranthac | yes | aphid | 21.99 | 0.9 | 1.558845726 | 3 | 12.59 | 2.8 | 4.849742261 | 3 | settling |
| Musser | 43 | 2003 | BPMV | Secoviridae | no | NCSP | no | Fabaceae | no | beetle | 32.2638 | 5.251099999 | 46.96726622 | 80 | 54.9587 | 5.2511 | 46.96726622 | 80 | feeding |
| Musser | 43 | 2003 | SBMV | Sobemoviru | no | NCSP | no | Fabaceae | no | beetle | 32.2638 | 5.251099999 | 46.96726622 | 80 | 67.1234 | 5.5599 | 49.72925739 | 80 | feeding |
| Ogada | 44 | 2013 | TSWV | Tospoviridae | yes | CPP | yes | Solanaceae | no | thrips | 5.67 | 1.53 | 1.53 | 60 | 12.67 | 2.08 | 2.08 | 60 | settling |
| Ogada | 44 | 2013 | TSWV | Tospoviridae | yes | CPP | yes | Solanaceae | no | thrips | 6.33 | 0.58 | 0.58 | 60 | 13 | 1 | 1 | 60 | settling |
| Ogada | 44 | 2013 | TSWV | Tospoviridae | yes | CPP | yes | Solanaceae | no | thrips | 3.67 | 0.58 | 0.58 | 60 | 16 | 1 | 1 | 60 | settling |
| Ogada | 44 | 2013 | TSWV | Tospoviridae | yes | CPP | yes | Solanaceae | no | thrips | 7 | 0.01 | 0.01 | 60 | 11 | 2 | 2 | 60 | settling |
| Ogada | 44 | 2013 | TSWV | Tospoviridae | yes | CPP | yes | Solanaceae | no | thrips | 5 | 2.65 | 2.65 | 60 | 14 | 2.65 | 2.65 | 60 | settling |
| Ogada | 44 | 2013 | TSWV | Tospoviridae | yes | CPP | yes | Solanaceae | no | thrips | 3.67 | 0.58 | 0.58 | 60 | 16 | 1 | 1 | 60 | settling |
| Pereira | 67 | 2019 | ToCV | Closteroviric | no | NCSP | yes | Solanaceae | no | Wf | 20.33 | 4.52 | 17.50588472 | 15 | 16.3 | 4.63 | 17.93191289 | 15 | settling |
| Pereira | 67 | 2019 | ToCV | Closteroviric | no | NCSP | yes | Solanaceae | no | Wf | 13.09 | 5.35 | 20.72046090 | 15 | 23.2 | 7.01 | 27.14961325 | 15 | settling |
| Pereira | 67 | 2019 | ToCV | Closteroviric | no | NCSP | yes | Solanaceae | no | Wf | 35.88 | 8.79 | 34.04352361 | 15 | 44.44 | 16.29 | 63.09089870 | 15 | settling |
| Pereira | 67 | 2019 | ToCV | Closteroviric | no | NCSP | yes | Solanaceae | no | Wf | 29.23 | 13.79 | 53.40844034 | 15 | 57.41 | 16.41 | 63.55565671 | 15 | settling |
| Pereira | 67 | 2019 | ToCV | Closteroviric | no | NCSP | yes | Solanaceae | no | Wf | 23.5 | 3.3 | 12.78084504 | 15 | 24.2 | 6.4 | 24.78709341 | 15 | eggs |
| Pereira | 67 | 2019 | ToCV | Closteroviric | no | NCSP | yes | Solanaceae | no | Wf | 23.3 | 6.1 | 23.62519841 | 15 | 28.8 | 3.7 | 14.33003838 | 15 | eggs |
| Pereira | 67 | 2019 | ToCV | Closteroviric | no | NCSP | yes | Solanaceae | no | Wf | 9.4 | 0.9 | 3.485685011 | 15 | 9.3 | 0.7 | 2.711088342 | 15 | eggs |
| Pereira | 67 | 2019 | ToCV | Closteroviric | no | NCSP | yes | Solanaceae | no | Wf | 10.2 | 0.8 | 3.098386676 | 15 | 9.8 | 0.6 | 2.323790007 | 15 | eggs |
| Pereira | 67 | 2019 | ToCV | Closteroviric | no | NCSP | yes | Solanaceae | no | Wf | 1.08 | 0.33 | 1.278084504 | 15 | 2.86 | 0.46 | 1.781572339 | 15 | dispersion |
| Pereira | 67 | 2019 | ToCV | Closteroviric | no | NCSP | yes | Solanaceae | no | Wf | 1.65 | 0.34 | 1.316814337 | 15 | 5.04 | 0.7 | 2.711088342 | 15 | dispersion |
| Pereira | 67 | 2019 | ToCV | Closteroviric | no | NCSP | yes | Solanaceae | no | Wf | 0.61 | 0.25 | 0.968245836 | 15 | 3.47 | 0.86 | 3.330765677 | 15 | dispersion |
| Pereira | 67 | 2019 | ToCV | Closteroviric | no | NCSP | yes | Solanaceae | no | Wf | 0.39 | 0.16 | 0.619677335 | 15 | 1.04 | 0.46 | 1.781572339 | 15 | dispersion |
| Penaflor | 45 | 2016 | SMV | Potyviridae | no | NCNP | no | Fabaceae | no | aphid | 2.915 | 0.352 | 1.113121736 | 10 | 3.85 | 0.389 | 1.230126009 | 10 | settling |
| Penaflor | 45 | 2016 | SMV | Potyviridae | no | NCNP | no | Fabaceae | no | aphid | 3 | 0.303 | 0.958170131 | 10 | 4.409 | 0.303 | 0.958170131 | 10 | settling |
| Penaflor | 46 | 2016 | BPMV | Secoviridae | no | NCSP | no | Fabaceae | no | beetle | 0.75257 | 0.3 | 0.9 | 9 | 1.931 | 0.449 | 1.347 | 9 | feeding |
| Porras | 47 | 2018 | BYDV | Luteoviridae | yes | CPNP | yes | Poaceae | no | aphid | 222.11 | 7.035999999 | 17.23460983 | 6 | 371.86 | 8.029999999 | 19.66940263 | 6 | EPG_E2 |
| Porras | 47 | 2018 | BYDV | Luteoviridae | yes | CPNP | yes | Poaceae | no | aphid | 367.31 | 9.610000000 | 23.53959642 | 6 | 393.27 | 7.69 | 18.83657612 | 6 | EPG_E2 |
| Rajabaskar | 48 | 2014 | PLRV | Luteoviridae | yes | CPNP | yes | Solanaceae | no | aphid | 28.8179 | 2.2434 | 8.688650838 | 15 | 78.1708 | 1.7257 | 6.683607360 | 15 | retention |
| Rajabaskar | 48 | 2014 | PLRV | Luteoviridae | yes | CPNP | yes | Solanaceae | no | aphid | 71.5272 | 1.984400000 | 7.685548152 | 15 | 21.5703 | 1.8982 | 7.351696987 | 15 | dispersion |
| Rajabaskar | 48 | 2014 | PLRV | Luteoviridae | yes | CPNP | yes | Solanaceae | no | aphid | 36.2268 | 2.9784 | 8.424187348 | 8 | 63.9658 | 2.977999999 | 8.423055977 | 8 | settling |
| Rajabaskar | 48 | 2014 | PLRV | Luteoviridae | yes | CPNP | yes | Solanaceae | no | aphid | 38.169 | 2.9784 | 8.424187348 | 8 | 61.5038 | 2.849100000 | 8.058471721 | 8 | settling |
| Ren | 49 | 2015 | PVY | Potyviridae | no | NCNP | no | Solanaceae | no | aphid | 30 | 1.1 | 4.260281680 | 15 | 28 | 1.3 | 5.034878350 | 15 | EPG |
| Ren | 49 | 2015 | PVY | Potyviridae | no | NCNP | no | Solanaceae | no | aphid | 6 | 0.9 | 3.485685011 | 15 | 11 | 1.1 | 4.260281680 | 15 | EPG |
| Salvaudon | 50 | 2013 | ZYMV | Potyviridae | no | NCNP | no | Cucurbitacea | no | aphid | 4.74 | 1.08 | 3.415259872 | 10 | 6.3 | 0.889 | 2.811264839 | 10 | settling |
| Salvaudon | 50 | 2013 | WMV | Potyviridae | no | NCNP | no | Cucurbitacea | no | aphid | 3.04 | 0.61 | 1.928989372 | 10 | 5.61 | 0.779999999 | 2.466576574 | 10 | settling |
| Shalileh | 52 | 2016 | TSWV | Tospoviridae | yes | CPP | yes | Solanaceae | no | thrips | 6.653 | 0.463 | 3.586382578 | 60 | 12.009 | 2.127 | 16.47567115 | 60 | settling |
| Shresta | 53 | 2017 | SVYV | Potyviridae | no | NCSP | yes | Cucurbitacea | no | Wf | 47.33 | 3.71 | 18.17521389 | 24 | 40.42 | 3.8 | 18.61612204 | 24 | settling |
| Shresta | 53 | 2017 | SVYV | Potyviridae | no | NCSP | yes | Cucurbitacea | no | Wf | 61.92 | 3.89 | 19.05703019 | 24 | 29.94 | 0.369999999 | 1.812622409 | 24 | settling |
| Shresta | 53 | 2017 | SVYV | Potyviridae | no | NCSP | yes | Cucurbitacea | no | Wf | 40.82 | 4.14 | 20.28177507 | 24 | 50.59 | 4.37 | 21.40854035 | 24 | settling |
| Shresta | 53 | 2017 | SVYV | Potyviridae | no | NCSP | yes | Cucurbitacea | no | Wf | 42.5 | 3.45 | 16.90147922 | 24 | 49.97 | 4.26 | 20.86965260 | 24 | settling |
| Shresta | 54 | 2016 | SVYV | Potyviridae | no | NCSP | yes | Cucurbitacea | no | Wf | 14.85 | 1.29 | 7.065620991 | 30 | 17.1 | 1.55 | 8.489699641 | 30 | settling |
| Shresta | 54 | 2016 | SVYV | Potyviridae | no | NCSP | yes | Cucurbitacea | no | Wf | 20.056 | 1.374 | 7.525707940 | 30 | 16.34 | 1.41 | 7.722888060 | 30 | settling |
| Smith | 55 | 2017 | BPMV | Secoviridae | no | NCSP | no | Fabaceae | no | beetle | 0.689 | 0.072 | 0.394360241 | 30 | 2.196 | 0.123 | 0.673698745 | 30 | feeding |
| Srinivasan | 56 | 2007 | PLRV | Luteoviridae | yes | CPNP | yes | Solanaceae | no | aphid | 10.1656 | 1.6051 | 5.560229502 | 12 | 27.6688 | 2.0637 | 7.148866503 | 12 | settling |
| Srinivasan | 56 | 2007 | PLRV | Luteoviridae | yes | CPNP | yes | Solanaceae | no | aphid | 8.1351 | 1.5946 | 3.905956343 | 6 | 16.6486 | 2.2703 | 5.561076563 | 6 | settling |
| Srinivasan | 56 | 2007 | PLRV | Luteoviridae | yes | CPNP | yes | Solanaceae | no | aphid | 7.3683 | 1.5399 | 5.334370077 | 12 | 28.5748 | 1.4832 | 5.137955515 | 12 | settling |
| Srinivasan | 56 | 2007 | PLRV | Luteoviridae | yes | CPNP | yes | Solanaceae | no | aphid | 7.61388 | 1.07376 | 2.630164106 | 6 | 16.0412 | 0.911100000 | 2.231730104 | 6 | settling |
| Srinivasan | 57 | 2006 | PLRV | Luteoviridae | yes | CPNP | yes | Solanaceae | no | aphid | 0.3067 | 0.15654 | 0.442761982 | 8 | 1.92261 | 0.44639 | 1.262581584 | 8 | settling |
| Srinivasan | 57 | 2006 | PLRV | Luteoviridae | yes | CPNP | yes | Solanaceae | yes | aphid | 3.24862 | 0.2377 | 0.672317127 | 8 | 4.8587 | 0.56237 | 1.590622562 | 8 | settling |
| Srinivasan | 57 | 2006 | PLRV | Luteoviridae | yes | CPNP | yes | Solanaceae | no | aphid | 6.84 | 1.22667 | 4.249309528 | 12 | 22.36 | 1.5467 | 5.357925968 | 12 | settling |
| Srinivasan | 57 | 2006 | PLRV | Luteoviridae | yes | CPNP | yes | Solanaceae | yes | aphid | 29.4 | 2.08 | 7.205331359 | 12 | 44.33 | 3.15 | 10.91192008 | 12 | settling |
| Srinivasan | 57 | 2006 | PLRV | Luteoviridae | yes | CPNP | yes | Solanaceae | no | aphid | 9.6744 | 1.3122 | 4.545594139 | 12 | 27.4435 | 1.8953 | 6.565511791 | 12 | settling |
| Srinivasan | 57 | 2006 | PLRV | Luteoviridae | yes | CPNP | yes | Solanaceae | yes | aphid | 32.1413 | 1.3121 | 4.545247729 | 12 | 37.0824 | 1.9926 | 6.902568878 | 12 | settling |
| Srinivasan | 56 | 2007 | PVY | Potyviridae | no | NCNP | no | Solanaceae | no | aphid | 10.1656 | 1.6051 | 5.560229502 | 12 | 17.5796 | 1.758 | 6.089890639 | 12 | settling |
| Srinivasan | 56 | 2007 | PVY | Potyviridae | no | NCNP | no | Solanaceae | no | aphid | 8.1351 | 1.5946 | 3.905956343 | 6 | 15.7297 | 1.9189 | 4.700325867 | 6 | settling |
| Srinivasan | 56 | 2007 | PVY | Potyviridae | no | NCNP | no | Solanaceae | no | aphid | 7.3683 | 1.5399 | 5.334370077 | 12 | 8.308 | 3.1932 | 11.06156927 | 12 | settling |
| Srinivasan | 56 | 2007 | PVY | Potyviridae | no | NCNP | no | Solanaceae | no | aphid | 7.61388 | 1.07376 | 2.630164106 | 6 | 11.0304 | 1.1713 | 2.869087335 | 6 | settling |
| Tomitaka | 58 | 2015 | TSWV | Tospoviridae | yes | CPP | yes | Brassicaceae | no | thrips | 0.508666666 | 0.194666666 | 0.194666666 | 3 | 4.168333333 | 0.198666666 | 0.198666666 | 3 | feeding |
| Tomitaka | 58 | 2015 | TSWV | Tospoviridae | yes | CPP | yes | Brassicaceae | no | thrips | 0.508666666 | 0.194666666 | 0.194666666 | 3 | 2.359666666 | 0.206666666 | 0.206666666 | 3 | feeding |
| Tomitaka | 58 | 2015 | TSWV | Tospoviridae | yes | CPP | yes | Brassicaceae | no | thrips | 2.977 | 0.907 | 0.907 | 3 | 8.92 | 1.326 | 1.326 | 3 | settling |
| Tomitaka | 58 | 2015 | TSWV | Tospoviridae | yes | CPP | yes | Brassicaceae | no | thrips | 2.681 | 0.938 | 0.938 | 3 | 8.268 | 1.52 | 1.52 | 3 | settling |
| Tungadi | 59 | 2017 | CMV | Bromovirida | no | NCNP | no | Solanaceae | no | aphid | 0.561 | 0.051 | 0.051 | 6 | 0.439 | 0.051 | 0.051 | 6 | settling |
| Tungadi | 59 | 2017 | CMV | Bromovirida | no | NCNP | no | Solanaceae | no | aphid | 0.551 | 0.04 | 0.04 | 6 | 0.449 | 0.04 | 0.04 | 6 | settling |
| Wang | 60 | 2018 | RDV | Reoviridae | yes | CPP | yes | Poaceae | no | LH | 19.4 | 5.5 | 19.83053201 | 13 | 7.6 | 2.2 | 8.520563361 | 15 | EPG_E2 |
| Wang | 60 | 2018 | RDV | Reoviridae | yes | CPP | yes | Poaceae | no | LH | 13.2 | 4 | 14.42220510 | 13 | 9 | 1.9 | 7.358668357 | 15 | EPG |
| Westwood | 61 | 2013 | CMV | Bromovirida | no | NCNP | no | Brassicaceae | no | aphid | 10 | 0.001 | 0.001732050 | 3 | 5.97 | 1.148 | 1.988394327 | 3 | retention |
| Westwood | 61 | 2013 | CMV | Bromovirida | no | NCNP | no | Brassicaceae | no | aphid | 22.15 | 3.377 | 16.54385372 | 24 | 14.63 | 1.72 | 8.426244715 | 24 | EPG_E2 |
| Wosula | 62 | 2014 | SPFMV | Potyviridae | no | NCNP | no | Convolvulac | no | aphid | 5.1 | 0.7 | 1.979898987 | 8 | 9 | 1.5 | 4.242640687 | 8 | EPG |
| Wosula | 62 | 2014 | SPFMV | Potyviridae | no | NCNP | no | Convolvulac | no | aphid | 44.5 | 14.6 | 29.2 | 4 | 32.4 | 11.5 | 32.52691193 | 8 | EPG_E2 |
| Wosula | 62 | 2014 | SPFMV | Potyviridae | no | NCNP | no | Convolvulac | no | aphid | 6.6 | 1 | 2.828427124 | 8 | 5.7 | 1.6 | 4.525483399 | 8 | EPG |
| Wosula | 62 | 2014 | SPFMV | Potyviridae | no | NCNP | no | Convolvulac | no | aphid | 20.1 | 3.6 | 6.235382907 | 3 | 12.6 | 12.6 | 12.6 | 1 | EPG_E2 |
| Wosula | 62 | 2014 | SPFMV | Potyviridae | no | NCNP | no | Convolvulac | yes | aphid | 5.4 | 1.1 | 3.111269837 | 8 | 3.1 | 0.8 | 2.262741699 | 8 | EPG |
| Wosula | 62 | 2014 | SPFMV | Potyviridae | no | NCNP | no | Convolvulac | yes | aphid | 14.7 | 8.1 | 16.2 | 4 | 0.01 | 0.01 | 0.001 | 0 | EPG_E2 |
| Wosula | 62 | 2014 | SPFMV | Potyviridae | no | NCNP | no | Convolvulac | yes | aphid | 6.9 | 1.1 | 3.111269837 | 8 | 3.6 | 0.9 | 2.545584412 | 8 | EPG |
| Wosula | 62 | 2014 | SPFMV | Potyviridae | no | NCNP | no | Convolvulac | yes | aphid | 33 | 10 | 26.45751311 | 7 | 27.3 | 14.4 | 35.27265229 | 6 | EPG_E2 |
| Wu | 63 | 2017 | CMV | Bromovirida | no | NCNP | no | Brassicaceae | no | aphid | 41.3405 | 1.0295 | 2.302031982 | 5 | 58.72 | 1.21 | 2.705642252 | 5 | settling |
| Ziebel | 64 | 2011 | CMV | Bromovirida | no | NCNP | no | Solanaceae | no | aphid | 3.96 | 0.48 | 1.795995545 | 14 | 7.54 | 1.72 | 6.435650705 | 14 | EPG_E2 |
| Ziebel | 64 | 2011 | CMV | Bromovirida | no | NCNP | no | Solanaceae | no | aphid | 212.545 | 22.91 | 85.72137073 | 14 | 231.636 | 30.364 | 113.6116848 | 14 | EPG |

### Performance database

| Reference | Study | Year | Virus | Family | circulative | persistent | phloem_acq | Plant_family | wild | Vector | Healthy_mea | Healthy_SE | Healthy_SD | Healthy_n | Virus_mean | Virus_SE | Virus_SD | Virus_n | Parameter |
|---|---|---|---|---|---|---|---|---|---|---|---|---|---|---|---|---|---|---|---|
| Adachi | 1 | 2018 | TuMV | Potyviridae | no | NCNP | no | Brassicaceae | no | aphid | 25.97 | 6.47 | 19.41 | 9 | 22.88 | 5.34 | 16.02 | 9 | Population |
| Adachi | 1 | 2018 | TuMV | Potyviridae | no | NCNP | no | Brassicaceae | no | aphid | 11.83 | 1.72 | 5.16 | 9 | 19.47 | 2.55 | 7.65 | 9 | Population |
| Adachi | 1 | 2018 | TuMV | Potyviridae | no | NCNP | no | Brassicaceae | no | aphid | 9.91 | 1.17 | 3.51 | 9 | 7.76 | 0.99 | 2.97 | 9 | Population |
| Adachi | 1 | 2018 | TuMV | Potyviridae | no | NCNP | no | Brassicaceae | no | aphid | 3.74 | 0.7 | 2.1 | 9 | 6.04 | 0.67 | 2.01 | 9 | Population |
| Bak | 4 | 2017 | PVY | Potyviridae | no | NCNP | no | Solanaceae | no | aphid | 4.344 | 0.6316 | 3.094195443 | 24 | 9.8059 | 1.1947 | 6.207843299 | 27 | Fecundity |
| Bak | 4 | 2017 | TuMV | Potyviridae | no | NCNP | no | Solanaceae | no | aphid | 1.835 | 0.5038 | 1.745214393 | 12 | 3.7961 | 0.3478 | 1.555408885 | 20 | Fecundity |
| Bak | 4 | 2017 | PVY | Potyviridae | no | NCNP | no | Solanaceae | no | aphid | 1.3939 | 0.3434 | 0.971281874 | 8 | 1.0505 | 0.2222 | 0.628476507 | 8 | Fecundity |
| Bak | 4 | 2017 | TuMV | Potyviridae | no | NCNP | no | Solanaceae | no | aphid | 2.8687 | 0.5454 | 1.542624153 | 8 | 2.707 | 0.8889 | 2.514188871 | 8 | Fecundity |
| Belliure | 5 | 2005 | TSWV | Tospoviridae | yes | CPP | yes | Solanaceae | no | thrips | 10.8879 | 0.5166 | 1.862627788 | 13 | 12.3632 | 1.6529 | 3.3058 | 4 | Dev_time |
| Belliure | 6 | 2008 | TSWV | Tospoviridae | yes | CPP | yes | Solanaceae | no | thrips | 0.384 | 0.019 | 0.164544826 | 75 | 0.662 | 0.025 | 0.201556443 | 65 | Weight |
| Blua | 7 | 1992 | ZYMV | Potyviridae | no | NCNP | no | Cucurbitacea | no | aphid | 334.634 | 91.336 | 408.4670095 | 20 | 542.556 | 73.47 | 328.5678286 | 20 | Population |
| Blua | 7 | 1992 | ZYMV | Potyviridae | no | NCNP | no | Cucurbitacea | no | aphid | 7.42967 | 0.58043 | 2.595761872 | 20 | 6.82646 | 0.25284 | 1.130734854 | 20 | Dev_time |
| Blua | 7 | 1992 | ZYMV | Potyviridae | no | NCNP | no | Cucurbitacea | no | aphid | 519.072 | 167.767 | 750.2768327 | 20 | 330.675 | 85.808 | 383.7450420 | 20 | Population |
| Blua | 7 | 1992 | ZYMV | Potyviridae | no | NCNP | no | Cucurbitacea | no | aphid | 7.02577 | 0.39363 | 1.760366875 | 20 | 7.17255 | 0.24141 | 1.079618340 | 20 | Dev_time |
| Blua | 7 | 1992 | ZYMV | Potyviridae | no | NCNP | no | Cucurbitacea | no | aphid | 383.846 | 150 | 670.8203932 | 20 | 79.23 | 50.77 | 227.0503424 | 20 | Population |
| Blua | 7 | 1992 | ZYMV | Potyviridae | no | NCNP | no | Cucurbitacea | no | aphid | 6.5218 | 0.42017 | 1.879057364 | 20 | 8.09645 | 0.53535 | 2.394157983 | 20 | Dev_time |
| Blua | 8 | 1994 | ZYMV | Potyviridae | no | NCNP | no | Cucurbitacea | no | aphid | 6.3 | 0.2 | 0.894427190 | 20 | 7.1 | 0.2 | 0.894427190 | 20 | Dev_time |
| Blua | 8 | 1994 | ZYMV | Potyviridae | no | NCNP | no | Cucurbitacea | no | aphid | 13 | 0.8 | 3.577708763 | 20 | 15.7 | 0.8 | 3.577708763 | 20 | Longevity |
| Blua | 8 | 1994 | ZYMV | Potyviridae | no | NCNP | no | Cucurbitacea | no | aphid | 13 | 0.8 | 3.577708763 | 20 | 15.2 | 0.8 | 3.577708763 | 20 | Longevity |
| Blua | 8 | 1994 | ZYMV | Potyviridae | no | NCNP | no | Cucurbitacea | no | aphid | 13 | 0.8 | 3.577708763 | 20 | 13 | 0.8 | 3.577708763 | 20 | Longevity |
| Blua | 8 | 1994 | ZYMV | Potyviridae | no | NCNP | no | Cucurbitacea | no | aphid | 30.4 | 3.5 | 15.65247584 | 20 | 44.4 | 5.4 | 24.14953415 | 20 | Fecundity |
| Blua | 8 | 1994 | ZYMV | Potyviridae | no | NCNP | no | Cucurbitacea | no | aphid | 30.4 | 3.5 | 15.65247584 | 20 | 42.4 | 5.2 | 23.25510696 | 20 | Fecundity |
| Blua | 8 | 1994 | ZYMV | Potyviridae | no | NCNP | no | Cucurbitacea | no | aphid | 30.4 | 3.5 | 15.65247584 | 20 | 32.8 | 4.9 | 21.91346617 | 20 | Fecundity |
| Boni | 9 | 2017 | EACMV | Geminivirida | yes | CPNP | yes | Euphorbiace | no | Wf | 25.8 | 0.3 | 0.6 | 4 | 25.3 | 0.4 | 0.8 | 4 | Dev_time |
| Boni | 9 | 2017 | EACMV_UG | Geminivirida | yes | CPNP | yes | Euphorbiace | no | Wf | 25.8 | 0.3 | 0.6 | 4 | 25.6 | 0.3 | 0.6 | 4 | Dev_time |
| Boni | 9 | 2017 | EACMV | Geminivirida | yes | CPNP | yes | Euphorbiace | no | Wf | 25.5 | 0.2 | 0.4 | 4 | 25.3 | 0.2 | 0.4 | 4 | Dev_time |
| Boni | 9 | 2017 | EACMV_UG | Geminivirida | yes | CPNP | yes | Euphorbiace | no | Wf | 25.5 | 0.2 | 0.4 | 4 | 25.8 | 0.1 | 0.2 | 4 | Dev_time |
| Boni | 9 | 2017 | EACMV | Geminivirida | yes | CPNP | yes | Euphorbiace | no | Wf | 27.1 | 0.3 | 0.6 | 4 | 25.6 | 0.2 | 0.4 | 4 | Dev_time |
| Boni | 9 | 2017 | EACMV_UG | Geminivirida | yes | CPNP | yes | Euphorbiace | no | Wf | 27.1 | 0.3 | 0.6 | 4 | 26.4 | 0.2 | 0.4 | 4 | Dev_time |
| Boni | 9 | 2017 | EACMV | Geminivirida | yes | CPNP | yes | Euphorbiace | no | Wf | 7.3 | 1.7 | 3.4 | 4 | 12.8 | 5.9 | 11.8 | 4 | Fecundity |
| Boni | 9 | 2017 | EACMV_UG | Geminivirida | yes | CPNP | yes | Euphorbiace | no | Wf | 7.3 | 1.7 | 3.4 | 4 | 8.3 | 1.7 | 3.4 | 4 | Fecundity |
| Boni | 9 | 2017 | EACMV | Geminivirida | yes | CPNP | yes | Euphorbiace | no | Wf | 12.3 | 2.1 | 4.2 | 4 | 14 | 3.8 | 7.6 | 4 | Fecundity |
| Boni | 9 | 2017 | EACMV_UG | Geminivirida | yes | CPNP | yes | Euphorbiace | no | Wf | 12.3 | 2.1 | 4.2 | 4 | 20.3 | 10.1 | 20.2 | 4 | Fecundity |
| Boni | 9 | 2017 | EACMV | Geminivirida | yes | CPNP | yes | Euphorbiace | no | Wf | 8.5 | 5.6 | 11.2 | 4 | 20.8 | 8.6 | 17.2 | 4 | Fecundity |
| Boni | 9 | 2017 | EACMV_UG | Geminivirida | yes | CPNP | yes | Euphorbiace | no | Wf | 8.5 | 5.6 | 11.2 | 4 | 17.5 | 9.1 | 18.2 | 4 | Fecundity |
| Boni | 9 | 2017 | EACMV | Geminivirida | yes | CPNP | yes | Euphorbiace | no | Wf | 56.3 | 19.1 | 38.2 | 4 | 45.8 | 10.3 | 20.6 | 4 | Survival |
| Boni | 9 | 2017 | EACMV_UG | Geminivirida | yes | CPNP | yes | Euphorbiace | no | Wf | 56.3 | 19.1 | 38.2 | 4 | 54.6 | 5.5 | 11 | 4 | Survival |
| Boni | 9 | 2017 | EACMV | Geminivirida | yes | CPNP | yes | Euphorbiace | no | Wf | 68 | 3.6 | 7.2 | 4 | 50.3 | 14.2 | 28.4 | 4 | Survival |
| Boni | 9 | 2017 | EACMV_UG | Geminivirida | yes | CPNP | yes | Euphorbiace | no | Wf | 68 | 3.6 | 7.2 | 4 | 62.5 | 8.7 | 17.4 | 4 | Survival |
| Boni | 9 | 2017 | EACMV | Geminivirida | yes | CPNP | yes | Euphorbiace | no | Wf | 68.7 | 15.2 | 30.4 | 4 | 61.5 | 18.4 | 36.8 | 4 | Survival |
| Boni | 9 | 2017 | EACMV_UG | Geminivirida | yes | CPNP | yes | Euphorbiace | no | Wf | 68.7 | 15.2 | 30.4 | 4 | 41.3 | 17.3 | 34.6 | 4 | Survival |
| Boni | 9 | 2017 | EACMV | Geminivirida | yes | CPNP | yes | Euphorbiace | no | Wf | 22.4 | 0.3 | 0.6 | 4 | 24.3 | 0.4 | 0.8 | 4 | Dev_time |
| Boni | 9 | 2017 | EACMV | Geminivirida | yes | CPNP | yes | Euphorbiace | no | Wf | 22.4 | 0.7 | 1.4 | 4 | 23.9 | 0.3 | 0.6 | 4 | Dev_time |
| Boni | 9 | 2017 | EACMV | Geminivirida | yes | CPNP | yes | Euphorbiace | no | Wf | 24.6 | 0.5 | 1 | 4 | 21.2 | 0.4 | 0.8 | 4 | Dev_time |
| Boni | 9 | 2017 | EACMV | Geminivirida | yes | CPNP | yes | Euphorbiace | no | Wf | 10 | 3.6 | 7.2 | 4 | 17.3 | 5.5 | 11 | 4 | Fecundity |
| Boni | 9 | 2017 | EACMV | Geminivirida | yes | CPNP | yes | Euphorbiace | no | Wf | 6.8 | 1.3 | 2.6 | 4 | 21.3 | 10.1 | 20.2 | 4 | Fecundity |
| Boni | 9 | 2017 | EACMV | Geminivirida | yes | CPNP | yes | Euphorbiace | no | Wf | 17.8 | 7.8 | 15.6 | 4 | 17.8 | 9.3 | 18.6 | 4 | Fecundity |
| Boni | 9 | 2017 | EACMV | Geminivirida | yes | CPNP | yes | Euphorbiace | no | Wf | 50.3 | 7.4 | 14.8 | 4 | 32.4 | 14.7 | 29.4 | 4 | Survival |
| Boni | 9 | 2017 | EACMV | Geminivirida | yes | CPNP | yes | Euphorbiace | no | Wf | 22.4 | 4.7 | 9.4 | 4 | 35.3 | 8.3 | 16.6 | 4 | Survival |
| Boni | 9 | 2017 | EACMV | Geminivirida | yes | CPNP | yes | Euphorbiace | no | Wf | 28.7 | 3.7 | 7.4 | 4 | 20 | 10.6 | 21.2 | 4 | Survival |
| Cassone | 10 | 2014 | SMV | Potyviridae | no | NCNP | no | Fabaceae | no | aphid | 8.48361 | 1.22016 | 1.22016 | 30 | 8.61222 | 1.05919 | 1.05919 | 30 | Survival |
| Cassone | 10 | 2014 | SMV | Potyviridae | no | NCNP | no | Fabaceae | no | aphid | 27.4345 | 3.8216 | 3.8216 | 30 | 29.2873 | 3.5235 | 3.5235 | 30 | Fecundity |
| Casteel | 11 | 2014 | TuMV | Potyviridae | no | NCNP | no | Solanaceae | no | aphid | 2.94872 | 0.68138 | 2.890850511 | 18 | 13.3591 | 1.1652 | 3.864531205 | 11 | Fecundity |
| Casteel | 11 | 2014 | CMV | Bromovirida | no | NCNP | no | Solanaceae | no | aphid | 2.94872 | 0.68138 | 2.890850511 | 18 | 1.15422 | 0.61531 | 2.302279206 | 14 | Fecundity |
| Casteel | 11 | 2014 | TuMV | Potyviridae | no | NCNP | no | Brassicaceae | no | aphid | 9.5159 | 0.8733 | 2.470065408 | 8 | 13.4334 | 0.597 | 2.602262669 | 19 | Fecundity |
| Casteel | 11 | 2014 | CMV | Bromovirida | no | NCNP | no | Solanaceae | no | aphid | 6.8239 | 1.3522 | 5.736898737 | 18 | 14.0566 | 1.4151 | 5.834606770 | 17 | Fecundity |
| Casteel | 11 | 2014 | TuMV | Potyviridae | no | NCNP | no | Solanaceae | no | aphid | 43.5399 | 12.4851 | 41.40839217 | 11 | 91.3928 | 5.6338 | 22.5352 | 16 | Survival |
| Casteel | 11 | 2014 | CMV | Bromovirida | no | NCNP | no | Solanaceae | no | aphid | 1.18702 | 0.12055 | 0.2411 | 4 | 2.24884 | 0.36631 | 0.969165162 | 7 | Weight |
| Castle | 12 | 1993 | PLRV | Luteoviridae | yes | CPNP | yes | Solanaceae | no | aphid | 235.736 | 12.012 | 69.00368651 | 33 | 430.931 | 15.015 | 87.55174270 | 34 | Weight |
| Castle | 12 | 1993 | PVY | Potyviridae | no | NCNP | no | Solanaceae | no | aphid | 235.736 | 12.012 | 69.00368651 | 33 | 308.559 | 12.762 | 66.31329721 | 27 | Weight |
| Castle | 12 | 1993 | PLRV | Luteoviridae | yes | CPNP | yes | Solanaceae | no | aphid | 2.4804 | 0.32617 | 1.873703998 | 33 | 5.6131 | 0.4779 | 2.341222296 | 24 | Fecundity |
| Castle | 12 | 1993 | PVY | Potyviridae | no | NCNP | no | Solanaceae | no | aphid | 2.4804 | 0.32617 | 1.873703998 | 33 | 3.55752 | 0.72061 | 2.790910529 | 15 | Fecundity |
| Castle | 12 | 1993 | PLRV | Luteoviridae | yes | CPNP | yes | Solanaceae | no | aphid | 1.2037 | 0.258811 | 1.636864487 | 40 | 4.356 | 0.34274 | 2.167678090 | 40 | Fecundity |
| Castle | 12 | 1993 | PVY | Potyviridae | no | NCNP | no | Solanaceae | no | aphid | 1.2037 | 0.258811 | 1.636864487 | 40 | 2.5515 | 0.2437 | 1.541294131 | 40 | Fecundity |
| Chen | 13 | 2014 | WSMoV | Tospoviridae | yes | CPP | yes | Cucurbitacea | no | thrips | 136.1 | 6.89 | 38.97572577 | 32 | 132.474 | 4.714 | 18.856 | 16 | Dev_time |
| Chen | 13 | 2014 | WSMoV | Tospoviridae | yes | CPP | yes | Cucurbitacea | no | thrips | 1.38 | 0.15 | 0.807774721 | 29 | 1.32 | 0.14 | 0.779487010 | 31 | Size |
| Chen | 14 | 2013 | TYLCV | Geminivirida | yes | CPNP | yes | Solanaceae | yes | Wf | 51.4118 | 2.5882 | 14.17615523 | 30 | 90.1176 | 5.2942 | 28.99752763 | 30 | Fecundity |
| Chen | 14 | 2013 | TYLCV | Geminivirida | yes | CPNP | yes | Solanaceae | yes | Wf | 752.885 | 8.653 | 61.18594977 | 50 | 788.452 | 4.817 | 34.06133364 | 50 | Size |
| Chen | 14 | 2013 | TYLCV | Geminivirida | yes | CPNP | yes | Solanaceae | yes | Wf | 674.853 | 5.317 | 37.59686755 | 50 | 698.104 | 6.505 | 45.99729611 | 50 | Size |
| Chen | 14 | 2013 | TYLCV | Geminivirida | yes | CPNP | yes | Solanaceae | yes | Wf | 20.0966 | 0.1486 | 0.514765500 | 12 | 20.2637 | 0.1486 | 0.514765500 | 12 | Dev_time |
| Chen | 14 | 2013 | TYLCV | Geminivirida | yes | CPNP | yes | Solanaceae | yes | Wf | 53.9885 | 4.3731 | 15.14886277 | 12 | 69.5071 | 3.7066 | 12.84003904 | 12 | Survival |
| Chen | 14 | 2013 | TYLCV | Geminivirida | yes | CPNP | yes | Solanaceae | yes | Wf | 18.3464 | 0.7345 | 4.023022184 | 30 | 24.8965 | 0.8457 | 4.632089668 | 30 | Longevity |
| Chesnais | 15 | 2018 | TuYV | Luteoviridae | yes | CPNP | yes | Brassicaceae | no | aphid | 8.897 | 0.159 | 0.992954681 | 39 | 7.923 | 0.113 | 0.705684773 | 39 | Dev_time |
| Chesnais | 15 | 2018 | CaMV | Caulimovirid | no | NCSP | no | Brassicaceae | no | aphid | 8.897 | 0.159 | 0.992954681 | 39 | 9.235 | 0.164 | 0.956276110 | 34 | Dev_time |
| Chesnais | 15 | 2018 | TuYV | Luteoviridae | yes | CPNP | yes | Brassicaceae | no | aphid | 4.104 | 0.119 | 0.743154761 | 39 | 4.284 | 0.223 | 1.392634553 | 39 | Fecundity |
| Chesnais | 15 | 2018 | CaMV | Caulimovirid | no | NCSP | no | Brassicaceae | no | aphid | 4.104 | 0.119 | 0.743154761 | 39 | 3.258 | 0.146 | 0.851318976 | 34 | Fecundity |
| Chesnais | 15 | 2018 | TuYV | Luteoviridae | yes | CPNP | yes | Brassicaceae | no | aphid | 0.269 | 0.005 | 0.031224989 | 39 | 0.297 | 0.006 | 0.037469987 | 39 | R |
| Chesnais | 15 | 2018 | CaMV | Caulimovirid | no | NCSP | no | Brassicaceae | no | aphid | 0.269 | 0.005 | 0.031224989 | 39 | 0.242 | 0.005 | 0.029154759 | 34 | R |
| Chesnais | 15 | 2018 | TuYV | Luteoviridae | yes | CPNP | yes | Brassicaceae | no | aphid | 10 | 0.384 | 2.103254620 | 30 | 11.522 | 0.266 | 1.275691185 | 23 | Dev_time |
| Chesnais | 15 | 2018 | CaMV | Caulimovirid | no | NCSP | no | Brassicaceae | no | aphid | 10 | 0.384 | 2.103254620 | 30 | 10.962 | 0.225 | 1.147279390 | 26 | Dev_time |
| Chesnais | 15 | 2018 | TuYV | Luteoviridae | yes | CPNP | yes | Brassicaceae | no | aphid | 2.215 | 0.142 | 0.777766031 | 30 | 1.515 | 0.146 | 0.700191402 | 23 | Fecundity |
| Chesnais | 15 | 2018 | CaMV | Caulimovirid | no | NCSP | no | Brassicaceae | no | aphid | 2.215 | 0.142 | 0.777766031 | 30 | 1.285 | 0.101 | 0.515000970 | 26 | Fecundity |
| Chesnais | 15 | 2018 | TuYV | Luteoviridae | yes | CPNP | yes | Brassicaceae | no | aphid | 0.221 | 0.009 | 0.049295030 | 30 | 0.144 | 0.008 | 0.038366652 | 23 | R |
| Chesnais | 15 | 2018 | CaMV | Caulimovirid | no | NCSP | no | Brassicaceae | no | aphid | 0.221 | 0.009 | 0.049295030 | 30 | 0.117 | 0.008 | 0.040792156 | 26 | R |
| Chesnais | 81 | 2019 | TuYV | Luteoviridae | yes | CPNP | yes | Brassicaceae | no | aphid | 0.2749 | 0.0067 | 0.037900923 | 32 | 0.2542 | 0.0049 | 0.028148356 | 33 | R |
| Chesnais | 81 | 2019 | TuYV | Luteoviridae | yes | CPNP | yes | Brassicaceae | no | aphid | 0.2822 | 0.0086 | 0.050878286 | 35 | 0.2231 | 0.0074 | 0.043149044 | 34 | R |
| Chesnais | 81 | 2019 | TuYV | Luteoviridae | yes | CPNP | yes | Brassicaceae | yes | aphid | 0.2399 | 0.0069 | 0.039032294 | 32 | 0.2457 | 0.008 | 0.044542114 | 31 | R |
| Chesnais | 81 | 2019 | TuYV | Luteoviridae | yes | CPNP | yes | Brassicaceae | no | aphid | 347.2667 | 20.2781 | 111.0677279 | 30 | 260.7 | 15.2418 | 83.48277676 | 30 | weight |
| Chesnais | 81 | 2019 | TuYV | Luteoviridae | yes | CPNP | yes | Brassicaceae | no | aphid | 191.333 | 13.7967 | 75.56763809 | 30 | 93.4333 | 6.8601 | 37.57431516 | 30 | weight |
| Chesnais | 81 | 2019 | TuYV | Luteoviridae | yes | CPNP | yes | Brassicaceae | yes | aphid | 247.7667 | 15.0147 | 82.23889884 | 30 | 251.8667 | 17.6709 | 96.78750541 | 30 | weight |
| Colvin | 16 | 2006 | EACMV_UG | Geminivirida | yes | CPNP | yes | Euphorbiace | no | Wf | 214.988 | 76.781 | 76.781 | 35 | 1670.76 | 285.63 | 285.63 | 35 | Population |
| Costa | 17 | 1991 | CdTV | Geminivirida | yes | CPNP | yes | Solanaceae | no | Wf | 0.17 | 0.02 | 0.052915026 | 7 | 0.08 | 0.03 | 0.079372539 | 7 | Survival |
| Costa | 17 | 1991 | CLCV | Geminivirida | yes | CPNP | yes | Malvaceae | no | Wf | 0.55 | 0.05 | 0.141421356 | 8 | 0.35 | 0.05 | 0.141421356 | 8 | Survival |
| Costa | 17 | 1991 | LIYV | Closteroviric | no | NCSP | yes | Cucurbitacea | no | Wf | 0.59 | 0.04 | 0.089442719 | 5 | 0.6 | 0.07 | 0.156524758 | 5 | Survival |
| Costa | 17 | 1991 | LIYV | Closteroviric | no | NCSP | yes | Asteraceae | no | Wf | 0.26 | 0.05 | 0.111803398 | 5 | 0.32 | 0.07 | 0.156524758 | 5 | Survival |
| Costa | 17 | 1991 | WCMoV | Geminivirida | yes | CPNP | yes | Cucurbitacea | no | Wf | 0.52 | 0.06 | 0.103923048 | 3 | 0.75 | 0.05 | 0.086602540 | 3 | Survival |
| Costa | 17 | 1991 | WCMoV | Geminivirida | yes | CPNP | yes | Cucurbitacea | no | Wf | 0.83 | 0.03 | 0.051961524 | 3 | 0.72 | 0.03 | 0.051961524 | 3 | Survival |
| Costa | 17 | 1991 | CdTV | Geminivirida | yes | CPNP | yes | Solanaceae | no | Wf | 64.12 | 5.33 | 15.99 | 9 | 35.94 | 8.08 | 24.24 | 9 | Fecundity |
| Costa | 17 | 1991 | CLCV | Geminivirida | yes | CPNP | yes | Malvaceae | no | Wf | 78.55 | 11.73 | 33.17745017 | 8 | 47.92 | 6.8 | 20.4 | 9 | Fecundity |
| Costa | 17 | 1991 | LIYV | Closteroviric | no | NCSP | yes | Cucurbitacea | no | Wf | 41.33 | 4.93 | 12.07598443 | 6 | 56.82 | 7.46 | 18.27319348 | 6 | Fecundity |
| Costa | 17 | 1991 | LIYV | Closteroviric | no | NCSP | yes | Asteraceae | no | Wf | 76.62 | 10.45 | 23.36691036 | 5 | 72.6 | 6.97 | 15.58539380 | 5 | Fecundity |
| Costa | 17 | 1991 | WCMoV | Geminivirida | yes | CPNP | yes | Cucurbitacea | no | Wf | 59.94 | 6.93 | 16.97496391 | 6 | 51.38 | 5.89 | 13.17044038 | 5 | Fecundity |
| Costa | 17 | 1991 | WCMoV | Geminivirida | yes | CPNP | yes | Cucurbitacea | no | Wf | 128.29 | 12.14 | 21.02709680 | 3 | 85.8 | 12.09 | 20.94049426 | 3 | Fecundity |
| Cui | 18 | 2016 | TYLCV | Geminivirida | yes | CPNP | yes | Solanaceae | no | Wf | 59.4215 | 3.3884 | 16.59970208 | 24 | 36.9421 | 1.9009 | 9.312470104 | 24 | Fecundity |
| Cui | 18 | 2016 | TYLCV | Geminivirida | yes | CPNP | yes | Solanaceae | no | Wf | 80.6798 | 5.5076 | 26.98161941 | 24 | 49.3463 | 4.019 | 19.68899855 | 24 | Fecundity |
| Cui | 18 | 2016 | TYLCV | Geminivirida | yes | CPNP | yes | Solanaceae | no | Wf | 274.763 | 9.953 | 48.75954281 | 24 | 149.645 | 7.109 | 34.82684516 | 24 | Fecundity |
| Cui | 18 | 2016 | TYLCV | Geminivirida | yes | CPNP | yes | Solanaceae | no | Wf | 502.754 | 23.016 | 112.7549118 | 24 | 280.891 | 4.286 | 20.99702607 | 24 | Fecundity |

### Performance database continued

| | | | | | | | | | | | | | | | | | | | |
|---|---|---|---|---|---|---|---|---|---|---|---|---|---|---|---|---|---|---|---|
| Daimei | 19 | 2017 | GBNV | Tospoviridae | yes | CPP | yes | Fabaceae | no | thrips | 58.39 | 0.649999999 | 4.110960958 | 40 | 61.72 | 3.55 | 22.4521713 | 40 | Survival |
| Daimei | 19 | 2017 | GBNV | Tospoviridae | yes | CPP | yes | Fabaceae | no | thrips | 82.26 | 2.58 | 16.3173527 | 40 | 83.44 | 1.4 | 8.85437744 | 40 | Survival |
| Daimei | 19 | 2017 | GBNV | Tospoviridae | yes | CPP | yes | Fabaceae | no | thrips | 61.5 | 0.969999999 | 2.16898593 | 5 | 54.19 | 1.08000000 | 2.41495341 | 5 | Fecundity |
| Daimei | 19 | 2017 | GBNV | Tospoviridae | yes | CPP | yes | Fabaceae | no | thrips | 30.29 | 1.2 | 3.79473319 | 10 | 28.28 | 0.33 | 1.04355162 | 10 | Longevity |
| Daimei | 19 | 2017 | GBNV | Tospoviridae | yes | CPP | yes | Fabaceae | no | thrips | 3.95 | 0.12 | 0.75894663 | 40 | 3.55 | 0.05 | 0.31622776 | 40 | Dev_time |
| Daimei | 19 | 2017 | GBNV | Tospoviridae | yes | CPP | yes | Fabaceae | no | thrips | 14.78 | 0.17 | 1.07517440 | 40 | 13.6 | 0.17 | 1.07517440 | 40 | Dev_time |
| Davis | 20 | 2015 | BYDV_PAV | Luteoviridae | yes | CPNP | yes | Poaceae | no | aphid | 20.35 | 8.28 | 32.0683021 | 15 | 38.62 | 4.67 | 18.0868322 | 15 | Fecundity |
| Davis | 20 | 2015 | BYDV_PAV | Luteoviridae | yes | CPNP | yes | Poaceae | no | aphid | 41.38 | 7.93 | 30.7127579 | 15 | 53.96 | 8.02 | 31.0613264 | 15 | Fecundity |
| Davis | 21 | 2017 | BLRV | Luteoviridae | yes | CPNP | yes | Fabaceae | no | aphid | 28 | 3 | 9.48683298 | 10 | 46 | 3 | 9.48683298 | 10 | Survival |
| Davis | 21 | 2017 | BLRV | Luteoviridae | yes | CPNP | yes | Fabaceae | no | aphid | 33 | 2 | 6.32455532 | 10 | 48 | 5 | 15.8113883 | 10 | Survival |
| Davis | 21 | 2017 | BLRV | Luteoviridae | yes | CPNP | yes | Fabaceae | no | aphid | 42 | 3 | 9.48683298 | 10 | 55 | 6 | 18.9736659 | 10 | Survival |
| Davis | 21 | 2017 | BLRV | Luteoviridae | yes | CPNP | yes | Fabaceae | no | aphid | 46 | 2 | 6.32455532 | 10 | 53 | 5 | 15.8113883 | 10 | Survival |
| Davis | 21 | 2017 | BLRV | Luteoviridae | yes | CPNP | yes | Fabaceae | no | aphid | 14 | 1 | 3.16227766 | 10 | 32 | 4 | 12.6491106 | 10 | Survival |
| Davis | 21 | 2017 | BLRV | Luteoviridae | yes | CPNP | yes | Fabaceae | no | aphid | 52 | 4 | 12.6491106 | 10 | 65 | 5 | 15.8113883 | 10 | Survival |
| Davis | 21 | 2017 | BLRV | Luteoviridae | yes | CPNP | yes | Fabaceae | no | aphid | 8.8 | 0.4 | 1.26491106 | 10 | 9 | 0.4 | 1.26491106 | 10 | Dev_time |
| Davis | 21 | 2017 | BLRV | Luteoviridae | yes | CPNP | yes | Fabaceae | no | aphid | 8.8 | 0.5 | 1.58113883 | 10 | 8.5 | 0.4 | 1.26491106 | 10 | Dev_time |
| Davis | 21 | 2017 | BLRV | Luteoviridae | yes | CPNP | yes | Fabaceae | no | aphid | 9.7 | 0.6 | 1.89736659 | 10 | 10.1 | 0.5 | 1.58113883 | 10 | Dev_time |
| Davis | 21 | 2017 | BLRV | Luteoviridae | yes | CPNP | yes | Fabaceae | no | aphid | 10.3 | 0.2 | 0.63245553 | 10 | 8.9 | 0.5 | 1.58113883 | 10 | Dev_time |
| Davis | 21 | 2017 | BLRV | Luteoviridae | yes | CPNP | yes | Fabaceae | no | aphid | 9.1 | 0.5 | 1.58113883 | 10 | 9.1 | 0.5 | 1.58113883 | 10 | Dev_time |
| Davis | 21 | 2017 | BLRV | Luteoviridae | yes | CPNP | yes | Fabaceae | no | aphid | 7.5 | 0.3 | 0.94868329 | 10 | 7.4 | 0.3 | 0.94868329 | 10 | Dev_time |
| Davis | 21 | 2017 | BLRV | Luteoviridae | yes | CPNP | yes | Fabaceae | no | aphid | 0.195 | 0.018 | 0.05692099 | 10 | 0.226 | 0.013 | 0.04110960 | 10 | R |
| Davis | 21 | 2017 | BLRV | Luteoviridae | yes | CPNP | yes | Fabaceae | no | aphid | 0.247 | 0.023 | 0.07273238 | 10 | 0.27 | 0.034 | 0.10751744 | 10 | R |
| Davis | 21 | 2017 | BLRV | Luteoviridae | yes | CPNP | yes | Fabaceae | no | aphid | 0.208 | 0.006 | 0.01897366 | 10 | 0.203 | 0.025 | 0.07905694 | 10 | R |
| Davis | 21 | 2017 | BLRV | Luteoviridae | yes | CPNP | yes | Fabaceae | no | aphid | 0.144 | 0.014 | 0.04427188 | 10 | 0.255 | 0.027 | 0.08538149 | 10 | R |
| Davis | 21 | 2017 | BLRV | Luteoviridae | yes | CPNP | yes | Fabaceae | no | aphid | 0.157 | 0.024 | 0.07589466 | 10 | 0.193 | 0.3 | 0.94868329 | 10 | R |
| Davis | 21 | 2017 | BLRV | Luteoviridae | yes | CPNP | yes | Fabaceae | no | aphid | 0.359 | 0.016 | 0.05059644 | 10 | 0.364 | 0.011 | 0.03478505 | 10 | R |
| DeAngelis | 22 | 1993 | INSV | Tospoviridae | yes | CPP | yes | Campanulac | yes | thrips | 7.6 | 0.1 | 0.74161984 | 55 | 8.77 | 0.11 | 0.80833161 | 54 | Dev_time |
| Donaldson | 23 | 2007 | AMV | Bromovirida | no | NCNP | no | Fabaceae | no | aphid | 0.16 | 0.02 | 0.06928203 | 12 | 0.05 | 0.03 | 0.10392304 | 12 | Population |
| Donaldson | 23 | 2007 | AMV | Bromovirida | no | NCNP | no | Fabaceae | no | aphid | 0.07 | 0.02 | 0.06928203 | 12 | 0.04 | 0.01 | 0.03464101 | 12 | Population |
| Donaldson | 23 | 2007 | AMV | Bromovirida | no | NCNP | no | Fabaceae | no | aphid | 0.15 | 0.04 | 0.12649110 | 10 | 0.08 | 0.06 | 0.18973665 | 10 | Population |
| Donaldson | 23 | 2007 | AMV | Bromovirida | no | NCNP | no | Fabaceae | no | aphid | 0.253 | 0.001 | 0.00316227 | 10 | 0.196 | 0.001 | 0.00316227 | 10 | R |
| Donaldson | 23 | 2007 | AMV | Bromovirida | no | NCNP | no | Fabaceae | no | aphid | 0.359 | 0.002 | 0.00692820 | 12 | 0.396 | 0.003 | 0.01039230 | 12 | R |
| Donaldson | 23 | 2007 | SMV | Potyviridae | no | NCNP | no | Fabaceae | no | aphid | 500 | 89.27 | 178.54 | 4 | 270 | 57.18 | 114.36 | 4 | Population |
| dosSantos | 24 | 2016 | BYDV_PAV | Luteoviridae | yes | CPNP | yes | Poaceae | no | aphid | 298.191 | 23.884 | 82.7366029 | 12 | 395.175 | 23.884 | 82.7366029 | 12 | Population |
| Ellsbury | 25 | 1985 | BYMV | Potyviridae | no | NCNP | no | Fabaceae | no | aphid | 326.2 | 30.6 | 61.2 | 4 | 249 | 25.7 | 51.4 | 4 | Fecundity |
| Fereres | 26 | 1989 | BYDV_PAV | Luteoviridae | yes | CPNP | yes | Poaceae | no | aphid | 0.29 | 0.01 | 0.02828427 | 8 | 0.31 | 0.01 | 0.02828427 | 8 | R |
| Fereres | 26 | 1989 | BYDV_RPV | Luteoviridae | yes | CPNP | yes | Poaceae | no | aphid | 0.29 | 0.01 | 0.02828427 | 8 | 0.3 | 0.01 | 0.02828427 | 8 | R |
| Fereres | 26 | 1989 | BYDV_PAV | Luteoviridae | yes | CPNP | yes | Poaceae | no | aphid | 0.29 | 0.01 | 0.02828427 | 8 | 0.31 | 0.01 | 0.02828427 | 8 | R |
| Fereres | 26 | 1989 | BYDV_RPV | Luteoviridae | yes | CPNP | yes | Poaceae | no | aphid | 0.29 | 0.01 | 0.02828427 | 8 | 0.32 | 0.01 | 0.02828427 | 8 | R |
| Fereres | 26 | 1989 | BYDV_PAV | Luteoviridae | yes | CPNP | yes | Poaceae | no | aphid | 0.29 | 0.01 | 0.02828427 | 8 | 0.32 | 0.01 | 0.02828427 | 8 | R |
| Fereres | 26 | 1989 | BYDV_RPV | Luteoviridae | yes | CPNP | yes | Poaceae | no | aphid | 0.29 | 0.01 | 0.02828427 | 8 | 0.29 | 0.01 | 0.02828427 | 8 | R |
| Fereres | 26 | 1989 | BYDV_PAV | Luteoviridae | yes | CPNP | yes | Poaceae | no | aphid | 9.3 | 0.2 | 0.56568542 | 8 | 8 | 0.3 | 0.84852813 | 8 | Dev_time |
| Fereres | 26 | 1989 | BYDV_RPV | Luteoviridae | yes | CPNP | yes | Poaceae | no | aphid | 9.3 | 0.2 | 0.56568542 | 8 | 8 | 0.01 | 0.02828427 | 8 | Dev_time |
| Fereres | 26 | 1989 | BYDV_PAV | Luteoviridae | yes | CPNP | yes | Poaceae | no | aphid | 9.3 | 0.2 | 0.56568542 | 8 | 8.1 | 0.3 | 0.84852813 | 8 | Dev_time |
| Fereres | 26 | 1989 | BYDV_RPV | Luteoviridae | yes | CPNP | yes | Poaceae | no | aphid | 9.3 | 0.2 | 0.56568542 | 8 | 8 | 0.01 | 0.02828427 | 8 | Dev_time |
| Fereres | 26 | 1989 | BYDV_PAV | Luteoviridae | yes | CPNP | yes | Poaceae | no | aphid | 9.1 | 0.1 | 0.28284271 | 8 | 8 | 0.3 | 0.84852813 | 8 | Dev_time |
| Fereres | 26 | 1989 | BYDV_RPV | Luteoviridae | yes | CPNP | yes | Poaceae | no | aphid | 9.1 | 0.1 | 0.28284271 | 8 | 7.9 | 0.1 | 0.28284271 | 8 | Dev_time |
| Fereres | 26 | 1989 | BYDV_PAV | Luteoviridae | yes | CPNP | yes | Poaceae | no | aphid | 45 | 1.7 | 4.80832611 | 8 | 44.4 | 1.7 | 4.80832611 | 8 | Longevity |
| Fereres | 26 | 1989 | BYDV_RPV | Luteoviridae | yes | CPNP | yes | Poaceae | no | aphid | 45 | 1.7 | 4.80832611 | 8 | 43.3 | 1.6 | 4.52548339 | 8 | Longevity |
| Fereres | 26 | 1989 | BYDV_PAV | Luteoviridae | yes | CPNP | yes | Poaceae | no | aphid | 44.2 | 0.8 | 2.26274169 | 8 | 41.4 | 1.5 | 4.24264068 | 8 | Longevity |
| Fereres | 26 | 1989 | BYDV_RPV | Luteoviridae | yes | CPNP | yes | Poaceae | no | aphid | 44.2 | 0.8 | 2.26274169 | 8 | 41 | 1.6 | 4.52548339 | 8 | Longevity |
| Fereres | 26 | 1989 | BYDV_PAV | Luteoviridae | yes | CPNP | yes | Poaceae | no | aphid | 44.7 | 1.2 | 3.39411254 | 8 | 43.2 | 1.6 | 4.52548339 | 8 | Longevity |
| Fereres | 26 | 1989 | BYDV_RPV | Luteoviridae | yes | CPNP | yes | Poaceae | no | aphid | 44.7 | 1.2 | 3.39411254 | 8 | 44.6 | 1.2 | 3.39411254 | 8 | Longevity |
| Fereres | 26 | 1989 | BYDV_PAV | Luteoviridae | yes | CPNP | yes | Poaceae | no | aphid | 33.2 | 1.5 | 4.24264068 | 8 | 43.7 | 1.9 | 5.37401153 | 8 | Fecundity |
| Fereres | 26 | 1989 | BYDV_RPV | Luteoviridae | yes | CPNP | yes | Poaceae | no | aphid | 33.2 | 1.5 | 4.24264068 | 8 | 38.9 | 2 | 5.65685424 | 8 | Fecundity |
| Fereres | 26 | 1989 | BYDV_PAV | Luteoviridae | yes | CPNP | yes | Poaceae | no | aphid | 35.4 | 2.6 | 7.35391052 | 8 | 44 | 1.8 | 5.09116882 | 8 | Fecundity |
| Fereres | 26 | 1989 | BYDV_RPV | Luteoviridae | yes | CPNP | yes | Poaceae | no | aphid | 35.4 | 2.6 | 7.35391052 | 8 | 47.6 | 2.1 | 5.93969696 | 8 | Fecundity |
| Fereres | 26 | 1989 | BYDV_PAV | Luteoviridae | yes | CPNP | yes | Poaceae | no | aphid | 33.4 | 1.8 | 5.09116882 | 8 | 49 | 2.4 | 6.78822509 | 8 | Fecundity |
| Fereres | 26 | 1989 | BYDV_RPV | Luteoviridae | yes | CPNP | yes | Poaceae | no | aphid | 33.4 | 1.8 | 5.09116882 | 8 | 50.7 | 2.3 | 6.50538238 | 8 | Fecundity |
| Fiebig | 27 | 2004 | BYDV_MAV | Luteoviridae | yes | CPNP | yes | Poaceae | no | aphid | 0.2 | 0.005 | 0.05172040 | 107 | 0.16 | 0.005 | 0.04743416 | 90 | R |
| Fiebig | 27 | 2004 | BYDV_PAV | Luteoviridae | yes | CPNP | yes | Poaceae | no | aphid | 0.2 | 0.005 | 0.05172040 | 107 | 0.16 | 0.007 | 0.05814636 | 69 | R |
| Fiebig | 27 | 2004 | BYDV_MAV | Luteoviridae | yes | CPNP | yes | Poaceae | no | aphid | 0.16 | 0.007 | 0.05511805 | 62 | 0.15 | 0.005 | 0.04213074 | 71 | R |
| Fiebig | 27 | 2004 | BYDV_PAV | Luteoviridae | yes | CPNP | yes | Poaceae | no | aphid | 0.16 | 0.007 | 0.05511805 | 62 | 0.13 | 0.005 | 0.04582575 | 84 | R |
| Fiebig | 27 | 2004 | BYDV_MAV | Luteoviridae | yes | CPNP | yes | Poaceae | no | aphid | 0.87 | 0.03 | 0.31032241 | 107 | 0.701 | 0.03 | 0.28460498 | 90 | Weight |
| Fiebig | 27 | 2004 | BYDV_PAV | Luteoviridae | yes | CPNP | yes | Poaceae | no | aphid | 0.87 | 0.03 | 0.31032241 | 107 | 0.73 | 0.03 | 0.24919871 | 69 | Weight |
| Fiebig | 27 | 2004 | BYDV_MAV | Luteoviridae | yes | CPNP | yes | Poaceae | no | aphid | 0.713 | 0.03 | 0.23622023 | 62 | 0.799 | 0.03 | 0.25278449 | 71 | Weight |
| Fiebig | 27 | 2004 | BYDV_PAV | Luteoviridae | yes | CPNP | yes | Poaceae | no | aphid | 0.713 | 0.03 | 0.23622023 | 62 | 0.732 | 0.02 | 0.18330302 | 84 | Weight |
| Fiebig | 27 | 2004 | BYDV_MAV | Luteoviridae | yes | CPNP | yes | Poaceae | no | aphid | 10.6 | 0.1 | 1.03440804 | 107 | 12.9 | 0.2 | 1.89736659 | 90 | Dev_time |
| Fiebig | 27 | 2004 | BYDV_PAV | Luteoviridae | yes | CPNP | yes | Poaceae | no | aphid | 10.6 | 0.1 | 1.03440804 | 107 | 12.7 | 0.3 | 2.49198715 | 69 | Dev_time |
| Fiebig | 27 | 2004 | BYDV_MAV | Luteoviridae | yes | CPNP | yes | Poaceae | no | aphid | 11.9 | 0.2 | 1.57480157 | 62 | 12.1 | 0.3 | 2.52784493 | 71 | Dev_time |
| Fiebig | 27 | 2004 | BYDV_PAV | Luteoviridae | yes | CPNP | yes | Poaceae | no | aphid | 11.9 | 0.2 | 1.57480157 | 62 | 12.9 | 0.3 | 2.74954541 | 84 | Dev_time |
| Fiebig | 27 | 2004 | BYDV_MAV | Luteoviridae | yes | CPNP | yes | Poaceae | no | aphid | 19.6 | 0.6 | 6.20644825 | 107 | 18.9 | 0.7 | 6.64078308 | 90 | Fecundity |
| Fiebig | 27 | 2004 | BYDV_PAV | Luteoviridae | yes | CPNP | yes | Poaceae | no | aphid | 19.6 | 0.6 | 6.20644825 | 107 | 17.6 | 0.9 | 7.47596147 | 69 | Fecundity |
| Fiebig | 27 | 2004 | BYDV_MAV | Luteoviridae | yes | CPNP | yes | Poaceae | no | aphid | 16.7 | 0.8 | 6.29920629 | 62 | 15.4 | 0.8 | 6.74091981 | 71 | Fecundity |
| Fiebig | 27 | 2004 | BYDV_PAV | Luteoviridae | yes | CPNP | yes | Poaceae | no | aphid | 16.7 | 0.8 | 6.29920629 | 62 | 14.6 | 0.8 | 7.33212111 | 84 | Fecundity |
| Gadhave | 84 | 2019 | PRSV | Potyviridae | no | NCNP | no | Cucurbitacea | no | aphid | 0.3474 | 0.0185 | 0.08273451 | 20 | 0.4738 | 0.0219 | 0.09793977 | 20 | R |
| Gadhave | 84 | 2019 | PRSV | Potyviridae | no | NCNP | no | Cucurbitacea | no | aphid | 17.76 | 1.547 | 6.91839432 | 20 | 38.313 | 2.827 | 12.6427283 | 20 | Fecundity |
| Ghosh | 28 | 2016 | CBDV | Nanoviridae | yes | CPNP | yes | Zingiberacea | no | aphid | 14.5 | 0.2788 | 1.24683150 | 20 | 8.8 | 0.374 | 1.67257884 | 20 | Dev_time |
| Ghosh | 28 | 2016 | CBDV | Nanoviridae | yes | CPNP | yes | Zingiberacea | no | aphid | 10.2 | 0.331 | 1.48027700 | 20 | 17.5 | 0.3171 | 1.41811431 | 20 | Longevity |
| Ghosh | 28 | 2016 | CBDV | Nanoviridae | yes | CPNP | yes | Zingiberacea | no | aphid | 24.7 | 0.4032 | 1.80316521 | 20 | 26.3 | 0.4552 | 2.03571628 | 20 | Longevity |
| Ghosh | 28 | 2016 | CBDV | Nanoviridae | yes | CPNP | yes | Zingiberacea | no | aphid | 4.3 | 0.5673 | 2.53704272 | 20 | 15.6 | 0.3292 | 1.47222715 | 20 | Fecundity |
| Guo | 29 | 2010 | TYLCCNV | Geminivirida | yes | CPNP | yes | Solanaceae | no | Wf | 7.6996 | 1.5733 | 4.97521144 | 10 | 8.4863 | 1.0727 | 3.39217524 | 10 | Fecundity |
| Guo | 29 | 2010 | TYLCCNV | Geminivirida | yes | CPNP | yes | Solanaceae | no | Wf | 54.6841 | 2.6461 | 8.36770291 | 10 | 62.4791 | 3.5042 | 11.0812533 | 10 | Fecundity |
| Hily | 30 | 2014 | CMV | Bromovirida | no | NCNP | no | Brassicaceae | no | aphid | 7.55 | 0.09 | 0.45 | 25 | 7.5 | 0.09 | 0.45 | 25 | Dev_time |
| Hily | 30 | 2014 | CMV | Bromovirida | no | NCNP | no | Brassicaceae | no | aphid | 41.51 | 1.36 | 6.8 | 25 | 45.08 | 1.15 | 5.75 | 25 | Fecundity |
| Hily | 30 | 2014 | CMV | Bromovirida | no | NCNP | no | Brassicaceae | no | aphid | 5.52 | 0.19 | 0.95 | 25 | 6.03 | 0.15 | 0.75 | 25 | Fecundity |
| Hily | 30 | 2014 | CMV | Bromovirida | no | NCNP | no | Brassicaceae | no | aphid | 0.3642 | 0.0056 | 0.028 | 25 | 0.3752 | 0.0046 | 0.023 | 25 | R |
| Hily | 30 | 2014 | CMV | Bromovirida | no | NCNP | no | Brassicaceae | no | aphid | 7.72 | 0.09 | 0.45 | 25 | 7.57 | 0.1 | 0.5 | 25 | Dev_time |
| Hily | 30 | 2014 | CMV | Bromovirida | no | NCNP | no | Brassicaceae | no | aphid | 40.38 | 1.54 | 7.7 | 25 | 42.85 | 1.44 | 7.2 | 25 | Fecundity |
| Hily | 30 | 2014 | CMV | Bromovirida | no | NCNP | no | Brassicaceae | no | aphid | 5.24 | 0.2 | 1 | 25 | 5.66 | 0.17 | 0.85 | 25 | Fecundity |
| Hily | 30 | 2014 | CMV | Bromovirida | no | NCNP | no | Brassicaceae | no | aphid | 0.3531 | 0.0055 | 0.0275 | 25 | 0.366 | 0.00485 | 0.02425 | 25 | R |
| Hodge | 31 | 2008 | PEMV | Luteoviridae | yes | CPNP | yes | Fabaceae | no | aphid | 1.43 | 0.01 | 0.09165151 | 84 | 1.43 | 0.01 | 0.04795831 | 23 | Weight |
| Hodge | 31 | 2008 | BYMV | Potyviridae | no | NCNP | no | Fabaceae | no | aphid | 1.43 | 0.01 | 0.09165151 | 84 | 1.41 | 0.01 | 0.03741657 | 14 | Weight |
| Hodge | 31 | 2008 | PEMV | Luteoviridae | yes | CPNP | yes | Fabaceae | no | aphid | 0.28 | 0.01 | 0.11958260 | 143 | 0.28 | 0.01 | 0.05830951 | 34 | R |
| Hodge | 31 | 2008 | BYMV | Potyviridae | no | NCNP | no | Fabaceae | no | aphid | 0.28 | 0.01 | 0.11958260 | 143 | 0.28 | 0.01 | 0.05656854 | 32 | R |
| Hodge | 31 | 2008 | PEMV | Luteoviridae | yes | CPNP | yes | Fabaceae | no | aphid | 1.68 | 0.05 | 0.45825756 | 84 | 1.79 | 0.11 | 0.52754146 | 23 | Weight |
| Hodge | 31 | 2008 | BYMV | Potyviridae | no | NCNP | no | Fabaceae | no | aphid | 1.68 | 0.05 | 0.45825756 | 84 | 1.63 | 0.13 | 0.48641546 | 14 | Weight |
| Hodge | 31 | 2008 | PEMV | Luteoviridae | yes | CPNP | yes | Fabaceae | no | aphid | 7.9 | 0.2 | 2.48997991 | 155 | 7.8 | 0.4 | 2.49799919 | 39 | Fecundity |
| Hodge | 31 | 2008 | BYMV | Potyviridae | no | NCNP | no | Fabaceae | no | aphid | 7.9 | 0.2 | 2.48997991 | 155 | 7.8 | 0.4 | 2.49799919 | 39 | Fecundity |
| Hodge | 31 | 2008 | PEMV | Luteoviridae | yes | CPNP | yes | Fabaceae | no | aphid | 10.9 | 0.1 | 1.19582607 | 143 | 10.9 | 0.2 | 1.16619037 | 34 | Dev_time |
| Hodge | 31 | 2008 | BYMV | Potyviridae | no | NCNP | no | Fabaceae | no | aphid | 10.9 | 0.1 | 1.19582607 | 143 | 10.6 | 0.1 | 0.56568542 | 32 | Dev_time |

### Performance database continued

| | | | | | | | | | | | | | | | | | | | |
|---|---|---|---|---|---|---|---|---|---|---|---|---|---|---|---|---|---|---|---|
| Hodgson | 32 | 1981 | TuMV | Potyviridae | no | NCNP | no | Brassicaceae | no | aphid | 12.6 | 0.8 | 3.57708763 | 20 | 12.8 | 0.7 | 3.13049516 | 20 | Population |
| Hodgson | 32 | 1981 | TuMV | Potyviridae | no | NCNP | no | Brassicaceae | no | aphid | 10.2 | 0.8 | 3.57708763 | 20 | 11.5 | 0.9 | 4.02492235 | 20 | Population |
| Hodgson | 32 | 1981 | TuMV | Potyviridae | no | NCNP | no | Brassicaceae | no | aphid | 10.1 | 0.8 | 3.57708763 | 20 | 11.6 | 0.9 | 4.02492235 | 20 | Population |
| Hodgson | 32 | 1981 | TuMV | Potyviridae | no | NCNP | no | Brassicaceae | no | aphid | 11.7 | 0.9 | 4.02492235 | 20 | 13.9 | 1.4 | 6.26099033 | 20 | Population |
| Hodgson | 32 | 1981 | TuMV | Potyviridae | no | NCNP | no | Brassicaceae | no | aphid | 19.9 | 2.9 | 12.9691942 | 20 | 25.5 | 2.9 | 12.9691942 | 20 | Population |
| Hodgson | 32 | 1981 | TuMV | Potyviridae | no | NCNP | no | Brassicaceae | no | aphid | 53.7 | 6 | 26.8328157 | 20 | 78.1 | 8.9 | 39.8020099 | 20 | Population |
| Hodgson | 32 | 1981 | TuMV | Potyviridae | no | NCNP | no | Brassicaceae | no | aphid | 11.4 | 0.7 | 3.13049516 | 20 | 11.8 | 0.6 | 2.68328157 | 20 | Population |
| Hodgson | 32 | 1981 | TuMV | Potyviridae | no | NCNP | no | Brassicaceae | no | aphid | 11 | 0.6 | 2.68328157 | 20 | 11.5 | 0.5 | 2.23606797 | 20 | Population |
| Hodgson | 32 | 1981 | TuMV | Potyviridae | no | NCNP | no | Brassicaceae | no | aphid | 10.5 | 0.6 | 2.68328157 | 20 | 10.4 | 0.7 | 3.13049516 | 20 | Population |
| Hodgson | 32 | 1981 | TuMV | Potyviridae | no | NCNP | no | Brassicaceae | no | aphid | 21.9 | 2 | 8.94427190 | 20 | 18 | 1.6 | 7.15541752 | 20 | Population |
| Hodgson | 32 | 1981 | TuMV | Potyviridae | no | NCNP | no | Brassicaceae | no | aphid | 36.6 | 2.5 | 11.1803398 | 20 | 29.4 | 2.8 | 12.5219806 | 20 | Population |
| Hodgson | 32 | 1981 | TuMV | Potyviridae | no | NCNP | no | Brassicaceae | no | aphid | 85.3 | 5.1 | 22.8078933 | 20 | 52.3 | 6.1 | 27.2800293 | 20 | Population |
| Hodgson | 32 | 1981 | TuMV | Potyviridae | no | NCNP | no | Brassicaceae | no | aphid | 815.7 | 17.9 | 126.572113 | 50 | 907.1 | 17.9 | 126.572113 | 50 | Weight |
| Hodgson | 32 | 1981 | TuMV | Potyviridae | no | NCNP | no | Brassicaceae | no | aphid | 1.14 | 0.01 | 0.07071067 | 50 | 1.18 | 0.01 | 0.07071067 | 50 | Size |
| Hodgson | 32 | 1981 | TuMV | Potyviridae | no | NCNP | no | Brassicaceae | no | aphid | 873.5 | 13.2 | 93.3380951 | 50 | 941.5 | 15.5 | 109.601551 | 50 | Weight |
| Hodgson | 32 | 1981 | TuMV | Potyviridae | no | NCNP | no | Brassicaceae | no | aphid | 1.17 | 0.01 | 0.07071067 | 50 | 1.18 | 0.01 | 0.07071067 | 50 | Size |
| Hodgson | 32 | 1981 | TuMV | Potyviridae | no | NCNP | no | Brassicaceae | no | aphid | 968 | 30.2 | 213.546247 | 50 | 937 | 23.8 | 168.291413 | 50 | Weight |
| Hodgson | 32 | 1981 | TuMV | Potyviridae | no | NCNP | no | Brassicaceae | no | aphid | 0.86 | 0.01 | 0.07071067 | 50 | 0.88 | 0.01 | 0.07071067 | 50 | Size |
| Hodgson | 32 | 1981 | TuMV | Potyviridae | no | NCNP | no | Brassicaceae | no | aphid | 1038 | 18.1 | 127.986327 | 50 | 956 | 17 | 120.208152 | 50 | Weight |
| Hodgson | 32 | 1981 | TuMV | Potyviridae | no | NCNP | no | Brassicaceae | no | aphid | 0.89 | 0.01 | 0.07071067 | 50 | 0.89 | 0.01 | 0.07071067 | 50 | Size |
| Jimenez | 33 | 2004 | BYDV_PAV | Luteoviridae | yes | CPNP | yes | Poaceae | no | aphid | 0.239 | 0.004 | 0.01385640 | 12 | 0.263 | 0.008 | 0.02771281 | 12 | R |
| Jimenez | 33 | 2004 | BYDV_PAV | Luteoviridae | yes | CPNP | yes | Poaceae | no | aphid | 0.173 | 0.012 | 0.04156921 | 12 | 0.244 | 0.01 | 0.03464101 | 12 | R |
| Jimenez | 33 | 2004 | BYDV_PAV | Luteoviridae | yes | CPNP | yes | Poaceae | no | aphid | 0.263 | 0.014 | 0.04849742 | 12 | 0.229 | 0.017 | 0.05888972 | 12 | R |
| Jimenez | 33 | 2004 | BYDV_PAV | Luteoviridae | yes | CPNP | yes | Poaceae | no | aphid | 0.272 | 0.01 | 0.03464101 | 12 | 0.234 | 0.006 | 0.02078460 | 12 | R |
| Jimenez | 33 | 2004 | BYDV_PAV | Luteoviridae | yes | CPNP | yes | Poaceae | no | aphid | 28.1 | 0.12 | 0.41569219 | 12 | 25.4 | 0.15 | 0.51961524 | 12 | Longevity |
| Jimenez | 33 | 2004 | BYDV_PAV | Luteoviridae | yes | CPNP | yes | Poaceae | no | aphid | 27 | 0.13 | 0.45033320 | 12 | 25.4 | 0.25 | 0.86602540 | 12 | Longevity |
| Jimenez | 33 | 2004 | BYDV_PAV | Luteoviridae | yes | CPNP | yes | Poaceae | no | aphid | 27 | 0.13 | 0.45033320 | 12 | 25.8 | 0.15 | 0.51961524 | 12 | Longevity |
| Jimenez | 33 | 2004 | BYDV_PAV | Luteoviridae | yes | CPNP | yes | Poaceae | no | aphid | 27.2 | 0.26 | 0.90066641 | 12 | 25.2 | 0.13 | 0.45033320 | 12 | Longevity |
| Jimenez | 33 | 2004 | BYDV_PAV | Luteoviridae | yes | CPNP | yes | Poaceae | no | aphid | 83.7 | 0.98 | 3.39481958 | 12 | 89.6 | 2.01 | 6.96284424 | 12 | Fecundity |
| Jimenez | 33 | 2004 | BYDV_PAV | Luteoviridae | yes | CPNP | yes | Poaceae | no | aphid | 79.8 | 1.3 | 4.50333209 | 12 | 83.8 | 1.05 | 3.63730669 | 12 | Fecundity |
| Jimenez | 33 | 2004 | BYDV_PAV | Luteoviridae | yes | CPNP | yes | Poaceae | no | aphid | 81.6 | 1.72 | 5.95825477 | 12 | 65.2 | 3.72 | 12.8864580 | 12 | Fecundity |
| Jimenez | 33 | 2004 | BYDV_PAV | Luteoviridae | yes | CPNP | yes | Poaceae | no | aphid | 87.6 | 1.91 | 6.61643408 | 12 | 76.8 | 2.88 | 9.97661265 | 12 | Fecundity |
| Jimenez | 33 | 2004 | BYDV_PAV | Luteoviridae | yes | CPNP | yes | Poaceae | no | aphid | 5.75 | 0.14 | 0.48497422 | 12 | 5.18 | 0.01 | 0.03464101 | 12 | Dev_time |
| Jimenez | 33 | 2004 | BYDV_PAV | Luteoviridae | yes | CPNP | yes | Poaceae | no | aphid | 6.45 | 0.25 | 0.86602540 | 12 | 5.82 | 0.1 | 0.34641016 | 12 | Dev_time |
| Jimenez | 33 | 2004 | BYDV_PAV | Luteoviridae | yes | CPNP | yes | Poaceae | no | aphid | 5.55 | 0.12 | 0.41569219 | 12 | 5.88 | 0.23 | 0.79674337 | 12 | Dev_time |
| Jimenez | 33 | 2004 | BYDV_PAV | Luteoviridae | yes | CPNP | yes | Poaceae | no | aphid | 5.45 | 0.18 | 0.62353829 | 12 | 6 | 0.2 | 0.69282032 | 12 | Dev_time |
| Jimenez | 33 | 2004 | BYDV_PAV | Luteoviridae | yes | CPNP | yes | Poaceae | no | aphid | 14 | 0.58 | 2.00917893 | 12 | 16.92 | 0.74 | 2.56343519 | 12 | Reprod_peri |
| Jimenez | 33 | 2004 | BYDV_PAV | Luteoviridae | yes | CPNP | yes | Poaceae | no | aphid | 14.75 | 0.48 | 1.66276877 | 12 | 15.58 | 1.05 | 3.63730669 | 12 | Reprod_peri |
| Jimenez | 33 | 2004 | BYDV_PAV | Luteoviridae | yes | CPNP | yes | Poaceae | no | aphid | 16.33 | 0.8 | 2.77128129 | 12 | 14.17 | 0.62 | 2.14774300 | 12 | Reprod_peri |
| Jimenez | 33 | 2004 | BYDV_PAV | Luteoviridae | yes | CPNP | yes | Poaceae | no | aphid | 15.67 | 1.13 | 3.91443482 | 12 | 13.72 | 1.13 | 3.91443482 | 12 | Reprod_peri |
| Jiu | 34 | 2007 | TbCSV | Geminivirida | yes | CPNP | yes | Solanaceae | no | Wf | 29.6 | 2.1 | 10.9119200 | 27 | 66.4 | 3.9 | 21.3611797 | 30 | Fecundity |
| Jiu | 34 | 2007 | TYLCCNV | Geminivirida | yes | CPNP | yes | Solanaceae | no | Wf | 29.6 | 2.1 | 10.9119200 | 27 | 66.2 | 3.2 | 18.3826004 | 33 | Fecundity |
| Jiu | 34 | 2007 | TbCSV | Geminivirida | yes | CPNP | yes | Solanaceae | no | Wf | 8.3 | 0.5 | 2.64575131 | 28 | 9.6 | 0.6 | 3.39411254 | 32 | Fecundity |
| Jiu | 34 | 2007 | TYLCCNV | Geminivirida | yes | CPNP | yes | Solanaceae | no | Wf | 8.3 | 0.5 | 2.64575131 | 28 | 9.6 | 0.7 | 3.83405790 | 30 | Fecundity |
| Jiu | 34 | 2007 | TbCSV | Geminivirida | yes | CPNP | yes | Solanaceae | no | Wf | 4.1 | 0.4 | 2.07846096 | 27 | 7.2 | 0.7 | 3.83405790 | 30 | Longevity |
| Jiu | 34 | 2007 | TYLCCNV | Geminivirida | yes | CPNP | yes | Solanaceae | no | Wf | 4.1 | 0.4 | 2.07846096 | 27 | 13.2 | 1.1 | 6.31901891 | 33 | Longevity |
| Jiu | 34 | 2007 | TbCSV | Geminivirida | yes | CPNP | yes | Solanaceae | no | Wf | 1.6 | 0.1 | 0.52915026 | 28 | 2.1 | 0.1 | 0.56568542 | 32 | Longevity |
| Jiu | 34 | 2007 | TYLCCNV | Geminivirida | yes | CPNP | yes | Solanaceae | no | Wf | 1.6 | 0.1 | 0.52915026 | 28 | 1.7 | 0.1 | 0.54772255 | 30 | Longevity |
| Jiu | 34 | 2007 | TbCSV | Geminivirida | yes | CPNP | yes | Solanaceae | no | Wf | 72.1 | 4.60000000 | 88.3631144 | 369 | 75.7 | 3.59999999 | 83.1903840 | 534 | Survival |
| Jiu | 34 | 2007 | TYLCCNV | Geminivirida | yes | CPNP | yes | Solanaceae | no | Wf | 72.1 | 4.60000000 | 88.3631144 | 369 | 74.6 | 4.2 | 85.7664269 | 417 | Survival |
| Jiu | 34 | 2007 | TbCSV | Geminivirida | yes | CPNP | yes | Solanaceae | no | Wf | 2.1 | 2 | 27.9284800 | 195 | 7.5 | 3.9 | 51.2964911 | 173 | Survival |
| Jiu | 34 | 2007 | TYLCCNV | Geminivirida | yes | CPNP | yes | Solanaceae | no | Wf | 2.1 | 2 | 27.9284800 | 195 | 4.8 | 2.9 | 42.0249925 | 210 | Survival |
| Jiu | 34 | 2007 | TbCSV | Geminivirida | yes | CPNP | yes | Solanaceae | no | Wf | 23 | 0.2 | 3.84187454 | 369 | 22.5 | 0.1 | 2.31084400 | 534 | Dev_time |
| Jiu | 34 | 2007 | TYLCCNV | Geminivirida | yes | CPNP | yes | Solanaceae | no | Wf | 23 | 0.2 | 3.84187454 | 369 | 22.8 | 0.2 | 4.08411557 | 417 | Dev_time |
| Jiu | 34 | 2007 | TbCSV | Geminivirida | yes | CPNP | yes | Solanaceae | no | Wf | 21.8 | 0.5 | 6.98212002 | 195 | 20.3 | 0.6 | 7.89176786 | 173 | Dev_time |
| Jiu | 34 | 2007 | TYLCCNV | Geminivirida | yes | CPNP | yes | Solanaceae | no | Wf | 21.8 | 0.5 | 6.98212002 | 195 | 23.6 | 0.6 | 8.69482604 | 210 | Dev_time |
| Jiu | 34 | 2007 | TbCSV | Geminivirida | yes | CPNP | yes | Solanaceae | no | Wf | 7.7 | 2.1 | 12.0635815 | 33 | 92 | 11.7 | 59.6585283 | 26 | Fecundity |
| Jiu | 34 | 2007 | TYLCCNV | Geminivirida | yes | CPNP | yes | Solanaceae | no | Wf | 7.7 | 2.1 | 12.0635815 | 33 | 138.4 | 13.3 | 74.0512660 | 31 | Fecundity |
| Jiu | 34 | 2007 | TbCSV | Geminivirida | yes | CPNP | yes | Solanaceae | no | Wf | 4.5 | 2.5 | 3.53553390 | 2 | 4.2 | 0.9 | 2.20454076 | 6 | Fecundity |
| Jiu | 34 | 2007 | TYLCCNV | Geminivirida | yes | CPNP | yes | Solanaceae | no | Wf | 4.5 | 2.5 | 3.53553390 | 2 | 3.5 | 1.3 | 2.6 | 4 | Fecundity |
| Jiu | 34 | 2007 | TbCSV | Geminivirida | yes | CPNP | yes | Solanaceae | no | Wf | 4.1 | 0.3 | 1.72336879 | 33 | 24.9 | 1.9 | 9.68813707 | 26 | Longevity |
| Jiu | 34 | 2007 | TYLCCNV | Geminivirida | yes | CPNP | yes | Solanaceae | no | Wf | 4.1 | 0.3 | 1.72336879 | 33 | 29.1 | 1.9 | 10.5787522 | 31 | Longevity |
| Jiu | 34 | 2007 | TbCSV | Geminivirida | yes | CPNP | yes | Solanaceae | no | Wf | 1.5 | 0.5 | 0.70710678 | 2 | 1.5 | 0.2 | 0.48989794 | 6 | Longevity |
| Jiu | 34 | 2007 | TYLCCNV | Geminivirida | yes | CPNP | yes | Solanaceae | no | Wf | 1.5 | 0.5 | 0.70710678 | 2 | 1.3 | 0.3 | 0.6 | 4 | Longevity |
| Kersch-Beck | 35 | 2014 | PVY_NTN | Potyviridae | no | NCNP | no | Solanaceae | no | aphid | 63.9387 | 10.6207 | 57.1942198 | 29 | 107.816 | 20.598 | 98.7845377 | 23 | Population |
| Kersch-Beck | 35 | 2014 | PVY_N0 | Potyviridae | no | NCNP | no | Solanaceae | no | aphid | 63.9387 | 10.6207 | 57.1942198 | 29 | 99.6628 | 13.7322 | 65.8573176 | 23 | Population |
| Kersch-Beck | 35 | 2014 | PVY_0 | Potyviridae | no | NCNP | no | Solanaceae | no | aphid | 63.9387 | 10.6207 | 57.1942198 | 29 | 87.3257 | 11.4789 | 56.2348956 | 24 | Population |
| Kersch-Beck | 35 | 2014 | PVY_NTN | Potyviridae | no | NCNP | no | Solanaceae | no | aphid | 8.04756 | 1.1964 | 7.66069783 | 41 | 5.50751 | 1.0217 | 6.62137277 | 42 | Fecundity |
| Kersch-Beck | 35 | 2014 | PVY_N0 | Potyviridae | no | NCNP | no | Solanaceae | no | aphid | 8.04756 | 1.1964 | 7.66069783 | 41 | 10.0536 | 1.1873 | 7.69458343 | 42 | Fecundity |
| Kersch-Beck | 35 | 2014 | PVY_0 | Potyviridae | no | NCNP | no | Solanaceae | no | aphid | 8.04756 | 1.1964 | 7.66069783 | 41 | 7.58725 | 1.2426 | 8.24247592 | 44 | Fecundity |
| Lei | 37 | 2014 | SRBSDV | Reoviridae | yes | CPP | yes | Poaceae | no | planthopper | 162.249 | 3.426 | 19.0751607 | 31 | 153.841 | 3.737 | 20.1243608 | 29 | Fecundity |
| Lei | 37 | 2014 | SRBSDV | Reoviridae | yes | CPP | yes | Poaceae | no | planthopper | 13.7 | 0.2 | 1.03923048 | 27 | 13.9 | 0.2 | 1.01980390 | 26 | Dev_time |
| Lei | 37 | 2014 | SRBSDV | Reoviridae | yes | CPP | yes | Poaceae | no | planthopper | 12.9675 | 0.2217 | 1.31159488 | 35 | 14.4961 | 0.2452 | 1.34301571 | 30 | Longevity |
| Li | 38 | 2011 | TYLCV | Geminivirida | yes | CPNP | yes | Solanaceae | no | Wf | 102.3 | 7.6 | 44.3152344 | 34 | 120.3 | 9.8 | 57.1433285 | 34 | Fecundity |
| Li | 38 | 2011 | TYLCV | Geminivirida | yes | CPNP | yes | Solanaceae | no | Wf | 61.6 | 9.3 | 55.8 | 36 | 52.6 | 6.9 | 43.6394317 | 40 | Fecundity |
| Li | 38 | 2011 | TYLCV | Geminivirida | yes | CPNP | yes | Solanaceae | no | Wf | 30.5 | 1.6 | 9.32952303 | 34 | 31.6 | 1.9 | 11.0788086 | 34 | Longevity |
| Li | 38 | 2011 | TYLCV | Geminivirida | yes | CPNP | yes | Solanaceae | no | Wf | 19.6 | 1.6 | 9.6 | 36 | 18.9 | 1.4 | 8.85437744 | 40 | Longevity |
| Li | 38 | 2011 | TYLCV | Geminivirida | yes | CPNP | yes | Solanaceae | no | Wf | 78.7 | 2.9 | 8.7 | 9 | 77.2 | 2.1 | 6.3 | 9 | Survival |
| Li | 38 | 2011 | TYLCV | Geminivirida | yes | CPNP | yes | Solanaceae | no | Wf | 35 | 1.8 | 5.4 | 9 | 40.9 | 1.4 | 4.2 | 9 | Survival |
| Li | 38 | 2011 | TYLCV | Geminivirida | yes | CPNP | yes | Solanaceae | no | Wf | 22.2 | 0.1 | 0.3 | 9 | 22.6 | 0.2 | 0.6 | 9 | Dev_time |
| Li | 38 | 2011 | TYLCV | Geminivirida | yes | CPNP | yes | Solanaceae | no | Wf | 23.2 | 0.3 | 0.9 | 9 | 24.5 | 0.3 | 0.9 | 9 | Dev_time |
| Li | 85 | 2019 | TYLCV | Geminivirida | yes | CPNP | yes | Solanaceae | no | Wf | 53.2258 | 6.4516 | 35.3368685 | 30 | 80.1075 | 3.49469999 | 19.1412602 | 30 | Survival |
| Li | 85 | 2019 | TYLCV | Geminivirida | yes | CPNP | yes | Solanaceae | no | Wf | 4.26309 | 0.70872 | 3.88181930 | 30 | 6.57181 | 0.32215 | 1.76448821 | 30 | Fecundity |
| Li | 39 | 2018 | ToCV | Closteroviric | no | NCSP | yes | Solanaceae | no | Wf | 23.65 | 0.31 | 3.50724963 | 128 | 26.66 | 0.37 | 3.66281312 | 98 | Dev_time |
| Li | 39 | 2018 | ToCV | Closteroviric | no | NCSP | yes | Solanaceae | no | Wf | 82.58 | 3.04 | 37.8476947 | 155 | 78.4 | 3.7 | 41.3672575 | 125 | Survival |
| Li | 39 | 2018 | ToCV | Closteroviric | no | NCSP | yes | Solanaceae | no | Wf | 23.84 | 0.81 | 7.24486024 | 80 | 20.6 | 0.94 | 6.44431532 | 47 | Longevity |
| Li | 39 | 2018 | ToCV | Closteroviric | no | NCSP | yes | Solanaceae | no | Wf | 20.51 | 1.37 | 9.49163842 | 48 | 19.94 | 1.02 | 7.28425699 | 51 | Longevity |
| Li | 39 | 2018 | ToCV | Closteroviric | no | NCSP | yes | Solanaceae | no | Wf | 22.06 | 0.72 | 6.43987577 | 80 | 18.87 | 0.91 | 6.23864568 | 47 | Reprod_peri |
| Li | 39 | 2018 | ToCV | Closteroviric | no | NCSP | yes | Solanaceae | no | Wf | 130.37 | 4.34 | 38.8181400 | 80 | 92.1 | 6.55 | 44.9045376 | 47 | Fecundity |
| Li | 39 | 2018 | ToCV | Closteroviric | no | NCSP | yes | Solanaceae | no | Wf | 0.1334 | 0.0033 | 0.02951609 | 80 | 0.103 | 0.0046 | 0.03153601 | 47 | R |
| Li | 82 | 2018 | SMV | Potyviridae | no | NCNP | no | Fabaceae | no | aphid | 1256.2 | 16.2 | 98.5407529 | 37 | 753.4 | 16.2 | 77.6924706 | 23 | Size |
| Li | 82 | 2018 | SMV | Potyviridae | no | NCNP | no | Fabaceae | no | aphid | 5.1 | 0.19 | 0.87068938 | 21 | 6.2 | 0.19 | 0.87068938 | 21 | Dev_time |
| Li | 82 | 2018 | SMV | Potyviridae | no | NCNP | no | Fabaceae | no | aphid | 5.7 | 0.2 | 0.89442719 | 20 | 7 | 0.19 | 0.84970583 | 20 | Dev_time |
| Li | 82 | 2018 | SMV | Potyviridae | no | NCNP | no | Fabaceae | no | aphid | 20.8 | 0.96 | 6.22151107 | 42 | 18.7 | 0.62 | 4.01805923 | 42 | Reprod_peri |
| Li | 82 | 2018 | SMV | Potyviridae | no | NCNP | no | Fabaceae | no | aphid | 50.5 | 3.09 | 12.36 | 16 | 28.6 | 1.59 | 6.36 | 16 | Fecundity |
| Lightle | 40 | 2014 | RpLV | Reoviridae | yes | CPP | yes | Rosaceae | no | aphid | 16.6071 | 4.2188 | 19.3329703 | 21 | 24.5759 | 6.2946 | 28.8454809 | 21 | Fecundity |
| Lightle | 40 | 2014 | RLMV | Closteroviric | no | NCSP | yes | Rosaceae | no | aphid | 16.6071 | 4.2188 | 19.3329703 | 21 | 27.7902 | 7.3661 | 33.7557108 | 21 | Fecundity |
| Lightle | 40 | 2014 | RpLV | Reoviridae | yes | CPP | yes | Rosaceae | no | aphid | 18.4466 | 0.8215 | 3.76458593 | 21 | 17.7371 | 0.5975 | 2.73808897 | 21 | Dev_time |
| Lightle | 40 | 2014 | RLMV | Closteroviric | no | NCSP | yes | Rosaceae | no | aphid | 18.4466 | 0.8215 | 3.76458593 | 21 | 17.7371 | 0.5975 | 2.73808897 | 21 | Dev_time |
| Lightle | 40 | 2014 | RpLV | Reoviridae | yes | CPP | yes | Rosaceae | no | aphid | 27.0351 | 2.6886 | 12.3207130 | 21 | 42.5691 | 3.3607 | 15.4006621 | 21 | Longevity |
| Lightle | 40 | 2014 | RLMV | Closteroviric | no | NCSP | yes | Rosaceae | no | aphid | 27.0351 | 2.6886 | 12.3207130 | 21 | 41.3368 | 3.5474 | 16.2562290 | 21 | Longevity |

### Performance database continued

| | | | | | | | | | | | | | | | | | | | |
|---|---|---|---|---|---|---|---|---|---|---|---|---|---|---|---|---|---|---|---|
| Liu | 41 | 2014 | BYDV_GAV | Luteoviridae | yes | CPNP | yes | Poaceae | no | aphid | 146.454 | 39.889 | 39.889 | 5 | 189.664 | 19.906 | 19.906 | 5 | Population |
| Liu | 41 | 2014 | BYDV_GAV | Luteoviridae | yes | CPNP | yes | Poaceae | no | aphid | 387.262 | 31.28 | 31.28 | 5 | 578.52 | 48.421 | 48.421 | 5 | Population |
| Liu | 41 | 2014 | BYDV_GAV | Luteoviridae | yes | CPNP | yes | Poaceae | no | aphid | 326.185 | 42.731 | 42.731 | 5 | 491.806 | 14.289 | 14.289 | 5 | Population |
| Liu | 41 | 2014 | BYDV_GAV | Luteoviridae | yes | CPNP | yes | Poaceae | no | aphid | 826.336 | 113.859 | 113.859 | 5 | 1191.27 | 244.81 | 244.81 | 5 | Population |
| Liu | 41 | 2014 | BYDV_GAV | Luteoviridae | yes | CPNP | yes | Poaceae | no | aphid | 1867.16 | 150.91 | 150.91 | 5 | 2021.37 | 156.64 | 156.64 | 5 | Population |
| Liu | 41 | 2014 | BYDV_GAV | Luteoviridae | yes | CPNP | yes | Poaceae | no | aphid | 1464.42 | 59.8 | 59.8 | 5 | 1823.69 | 99.64 | 99.64 | 5 | Population |
| Liu | 42 | 2009 | TYLCCNV | Geminivirida | yes | CPNP | yes | Solanaceae | no | Wf | 67.5 | 8.2 | 49.87865274 | 37 | 25.7 | 4.6 | 25.1952376 | 30 | Fecundity |
| Liu | 42 | 2009 | TYLCV | Geminivirida | yes | CPNP | yes | Solanaceae | no | Wf | 67.5 | 8.2 | 49.87865274 | 37 | 12.1 | 3 | 18.7349939 | 39 | Fecundity |
| Liu | 42 | 2009 | TYLCCNV | Geminivirida | yes | CPNP | yes | Solanaceae | no | Wf | 88.1 | 10.8 | 60.13185511 | 31 | 96.1 | 11.4 | 63.4725137 | 31 | Fecundity |
| Liu | 42 | 2009 | TYLCV | Geminivirida | yes | CPNP | yes | Solanaceae | no | Wf | 88.1 | 10.8 | 60.13185511 | 31 | 68.6 | 11.4 | 71.1929771 | 39 | Fecundity |
| Liu | 42 | 2009 | TYLCCNV | Geminivirida | yes | CPNP | yes | Solanaceae | no | Wf | 16.9 | 1.6 | 9.732420048 | 37 | 8.6 | 1.2 | 6.5726706900 | 30 | Longevity |
| Liu | 42 | 2009 | TYLCV | Geminivirida | yes | CPNP | yes | Solanaceae | no | Wf | 16.9 | 1.6 | 9.732420048 | 37 | 5.8 | 0.9 | 5.620498198 | 39 | Longevity |
| Liu | 42 | 2009 | TYLCCNV | Geminivirida | yes | CPNP | yes | Solanaceae | no | Wf | 16.1 | 1.9 | 10.57875228 | 31 | 21.9 | 2.3 | 12.8058580 | 31 | Longevity |
| Liu | 42 | 2009 | TYLCV | Geminivirida | yes | CPNP | yes | Solanaceae | no | Wf | 16.1 | 1.9 | 10.57875228 | 31 | 18.7 | 2.2 | 13.7389955 | 39 | Longevity |
| Liu | 42 | 2009 | TYLCCNV | Geminivirida | yes | CPNP | yes | Solanaceae | no | Wf | 4.8 | 1.3 | 4.110960958 | 10 | 0.01 | 0.01 | 0.031622776 | 10 | Survival |
| Liu | 42 | 2009 | TYLCV | Geminivirida | yes | CPNP | yes | Solanaceae | no | Wf | 4.8 | 1.3 | 4.110960958 | 10 | 2.5 | 1 | 3.16227766 | 10 | Survival |
| Liu | 42 | 2009 | TYLCCNV | Geminivirida | yes | CPNP | yes | Solanaceae | no | Wf | 84 | 3.9 | 12.33288287 | 10 | 86 | 3.8 | 12.0166551 | 10 | Survival |
| Liu | 42 | 2009 | TYLCV | Geminivirida | yes | CPNP | yes | Solanaceae | no | Wf | 84 | 3.9 | 12.33288287 | 10 | 75.4 | 2.6 | 8.22192191 | 10 | Survival |
| Liu | 43 | 2010 | TYLCCNV | Geminivirida | yes | CPNP | yes | Solanaceae | no | Wf | 83.9 | 15.3 | 94.31553424 | 38 | 144.8 | 19.6 | 112.5934278 | 33 | Fecundity |
| Liu | 43 | 2010 | TYLCCNV | Geminivirida | yes | CPNP | yes | Solanaceae | no | Wf | 16.6 | 2 | 12.32882800 | 38 | 31 | 2.6 | 14.9358628 | 33 | Longevity |
| Liu | 43 | 2010 | TYLCCNV | Geminivirida | yes | CPNP | yes | Solanaceae | no | Wf | 90.8 | 1.5 | 4.743416490 | 10 | 83 | 3.5 | 11.0679718 | 10 | Survival |
| Liu | 43 | 2010 | TYLCCNV | Geminivirida | yes | CPNP | yes | Solanaceae | no | Wf | 18.4 | 0.1 | 0.316227766 | 10 | 19.4 | 0.1 | 0.316227766 | 10 | Dev_time |
| Liu | 43 | 2010 | TYLCCNV | Geminivirida | yes | CPNP | yes | Solanaceae | no | Wf | 323.976 | 114.362 | 255.7212060 | 5 | 1705.81 | 667.1 | 1491.680947 | 5 | Population |
| Liu | 43 | 2010 | TYLCCNV | Geminivirida | yes | CPNP | yes | Solanaceae | no | Wf | 10.3 | 3 | 18.73499399 | 39 | 23.3 | 5.4 | 33.2878356 | 38 | Fecundity |
| Liu | 43 | 2010 | TYLCCNV | Geminivirida | yes | CPNP | yes | Solanaceae | no | Wf | 4.9 | 1 | 6.244997998 | 39 | 9.2 | 2 | 12.3288280 | 38 | Longevity |
| Liu | 43 | 2010 | TYLCCNV | Geminivirida | yes | CPNP | yes | Solanaceae | no | Wf | 8.1 | 2.5 | 8.660254037 | 12 | 5.4 | 1.7 | 5.37587202 | 10 | Survival |
| Liu | 43 | 2010 | TYLCCNV | Geminivirida | yes | CPNP | yes | Solanaceae | no | Wf | 22.8 | 0.2 | 0.692820323 | 12 | 25.1 | 0.2 | 0.632455532 | 10 | Dev_time |
| Luan | 45 | 2013 | TYLCCNV | Geminivirida | yes | CPNP | yes | Solanaceae | no | Wf | 46.7105 | 8.9913 | 31.14677685 | 12 | 81.7982 | 3.9474 | 13.6741947 | 12 | Survival |
| Luan | 45 | 2013 | TYLCCNV | Geminivirida | yes | CPNP | yes | Solanaceae | no | Wf | 22.3214 | 3.5715 | 12.37203891 | 12 | 39.1741 | 3.0134 | 10.4387238 | 12 | Fecundity |
| Maluta | 46 | 2014 | TYLCV | Geminivirida | yes | CPNP | yes | Solanaceae | no | Wf | 33.2 | 5.5 | 28.04460732 | 26 | 52.8 | 6.23 | 31.7668915 | 26 | Fecundity |
| Maluta | 46 | 2014 | TYLCV | Geminivirida | yes | CPNP | yes | Solanaceae | no | Wf | 16.5 | 0.06 | 0.795989949 | 176 | 16.7 | 0.19 | 2.707083301 | 203 | Dev_time |
| Maluta | 46 | 2014 | TYLCV | Geminivirida | yes | CPNP | yes | Solanaceae | no | Wf | 84.3 | 2.77 | 8.31 | 9 | 82.2 | 2.58 | 7.74 | 9 | Survival |
| Maluta | 47 | 2018 | ToCV | Closterovirid | no | NCSP | yes | Solanaceae | no | Wf | 16.5 | 0.2 | 2.332380757 | 136 | 17.9 | 0.5 | 3.905124837 | 61 | Dev_time |
| Maluta | 47 | 2018 | ToCV | Closterovirid | no | NCSP | yes | Solanaceae | no | Wf | 17.8 | 2.6 | 11.91469680 | 21 | 22.7 | 4.3 | 18.7432654 | 19 | Dev_time |
| Maluta | 47 | 2018 | ToCV | Closterovirid | no | NCSP | yes | Solanaceae | no | Wf | 12 | 2.1 | 8.133265027 | 15 | 18.1 | 3.2 | 12.3935467 | 15 | Dev_time |
| Maluta | 47 | 2018 | ToCV | Closterovirid | no | NCSP | yes | Solanaceae | no | Wf | 77.7 | 5.8 | 15.34535760 | 7 | 32.6 | 6.3 | 16.6682332 | 7 | Survival |
| Maluta | 47 | 2018 | ToCV | Closterovirid | no | NCSP | yes | Solanaceae | no | Wf | 198.3 | 30 | 140.7124727 | 22 | 129.5 | 15.8 | 74.1085690 | 22 | Fecundity |
| Maluta | 47 | 2018 | ToCV | Closterovirid | no | NCSP | yes | Solanaceae | no | Wf | 89.07 | 2.3 | 10.78795624 | 22 | 94.24 | 1.6 | 7.50466521 | 22 | Survival |
| Maluta | 47 | 2018 | ToSRV | Geminivirida | yes | CPNP | yes | Solanaceae | no | Wf | 14.6 | 0.1 | 0.938083151 | 88 | 14.7 | 0.2 | 1.93907194 | 94 | Dev_time |
| Maluta | 47 | 2018 | ToSRV | Geminivirida | yes | CPNP | yes | Solanaceae | no | Wf | 23.3 | 1.8 | 8.818163074 | 24 | 23.1 | 1.9 | 8.70689382 | 21 | Dev_time |
| Maluta | 47 | 2018 | ToSRV | Geminivirida | yes | CPNP | yes | Solanaceae | no | Wf | 26.4 | 2 | 9.591663046 | 23 | 17.7 | 1.9 | 8.49705831 | 20 | Dev_time |
| Maluta | 47 | 2018 | ToSRV | Geminivirida | yes | CPNP | yes | Solanaceae | no | Wf | 86.1 | 3.5 | 10.5 | 9 | 52.1 | 4.3 | 12.9 | 9 | Survival |
| Maluta | 47 | 2018 | ToSRV | Geminivirida | yes | CPNP | yes | Solanaceae | no | Wf | 220.9 | 17.9 | 87.69173279 | 24 | 203.4 | 20.2 | 94.7463983 | 22 | Fecundity |
| Maluta | 47 | 2018 | ToSRV | Geminivirida | yes | CPNP | yes | Solanaceae | no | Wf | 89.9 | 2.9 | 14.20704050 | 24 | 92.54 | 2.2 | 10.3189146 | 22 | Survival |
| Mann | 48 | 2008 | CLCuV | Geminivirida | yes | CPNP | yes | Malvaceae | no | Wf | 51.93 | 1.55 | 2.192031021 | 2 | 49.17 | 0.58 | 0.82024386 | 2 | Fecundity |
| Mann | 48 | 2008 | CLCuV | Geminivirida | yes | CPNP | yes | Malvaceae | no | Wf | 50 | 0.63 | 0.890954544 | 2 | 45.6 | 0.71 | 1.00409162 | 2 | Fecundity |
| Mann | 48 | 2008 | CLCuV | Geminivirida | yes | CPNP | yes | Malvaceae | no | Wf | 53.2 | 1.09 | 1.541492782 | 2 | 44.83 | 2.36 | 3.33754400 | 2 | Fecundity |
| Mann | 48 | 2008 | CLCuV | Geminivirida | yes | CPNP | yes | Malvaceae | no | Wf | 3.88 | 0.1 | 0.447213595 | 20 | 3.58 | 0.13 | 0.58137767 | 20 | Dev_time |
| Mann | 48 | 2008 | CLCuV | Geminivirida | yes | CPNP | yes | Malvaceae | no | Wf | 3.96 | 0.1 | 0.447213595 | 20 | 3.97 | 0.07 | 0.31304951 | 20 | Dev_time |
| Mann | 48 | 2008 | CLCuV | Geminivirida | yes | CPNP | yes | Malvaceae | no | Wf | 4.3 | 0.07 | 0.313049516 | 20 | 4.26 | 0.14 | 0.62609903 | 20 | Dev_time |
| Mann | 48 | 2008 | CLCuV | Geminivirida | yes | CPNP | yes | Malvaceae | no | Wf | 15.38 | 0.42 | 1.878297101 | 20 | 13.74 | 0.24 | 1.07331262 | 20 | Dev_time |
| Mann | 48 | 2008 | CLCuV | Geminivirida | yes | CPNP | yes | Malvaceae | no | Wf | 16.17 | 0.39 | 1.744133022 | 20 | 13.91 | 0.76 | 3.39882332 | 20 | Dev_time |
| Mann | 48 | 2008 | CLCuV | Geminivirida | yes | CPNP | yes | Malvaceae | no | Wf | 16.16 | 0.68 | 3.041052449 | 20 | 14.66 | 0.66 | 2.95160973 | 20 | Dev_time |
| Mann | 48 | 2008 | CLCuV | Geminivirida | yes | CPNP | yes | Malvaceae | no | Wf | 5.24 | 0.16 | 0.715541752 | 20 | 4.57 | 0.16 | 0.71554175 | 20 | Dev_time |
| Mann | 48 | 2008 | CLCuV | Geminivirida | yes | CPNP | yes | Malvaceae | no | Wf | 5.55 | 0.25 | 1.118033988 | 20 | 4.47 | 0.14 | 0.62609903 | 20 | Dev_time |
| Mann | 48 | 2008 | CLCuV | Geminivirida | yes | CPNP | yes | Malvaceae | no | Wf | 5.33 | 0.16 | 0.715541752 | 20 | 4.09 | 0.03 | 0.13416407 | 20 | Dev_time |
| Mann | 48 | 2008 | CLCuV | Geminivirida | yes | CPNP | yes | Malvaceae | no | Wf | 9.46 | 0.03 | 0.094868329 | 10 | 8.43 | 0.16 | 0.50596442 | 10 | Longevity |
| Mann | 48 | 2008 | CLCuV | Geminivirida | yes | CPNP | yes | Malvaceae | no | Wf | 10.66 | 0.36 | 1.138419957 | 10 | 9.03 | 0.72 | 2.27683991 | 10 | Longevity |
| Mann | 48 | 2008 | CLCuV | Geminivirida | yes | CPNP | yes | Malvaceae | no | Wf | 12.18 | 0.38 | 1.201665510 | 10 | 10.87 | 0.61 | 1.92898937 | 10 | Longevity |
| Mann | 48 | 2008 | CLCuV | Geminivirida | yes | CPNP | yes | Malvaceae | no | Wf | 7.99 | 0.2 | 0.632455532 | 10 | 6.58 | 0.26 | 0.82219219 | 10 | Longevity |
| Mann | 48 | 2008 | CLCuV | Geminivirida | yes | CPNP | yes | Malvaceae | no | Wf | 8.27 | 0.4 | 1.264911064 | 10 | 6.73 | 0.45 | 1.42302494 | 10 | Longevity |
| Mann | 48 | 2008 | CLCuV | Geminivirida | yes | CPNP | yes | Malvaceae | no | Wf | 11.57 | 0.11 | 0.347850542 | 10 | 9.48 | 0.17 | 0.53758720 | 10 | Longevity |
| Maris | 49 | 2004 | TSWV | Tospoviridae | yes | CPP | yes | Solanaceae | no | thrips | 10.9 | 0.2 | 0.894427190 | 20 | 12.1 | 0.1 | 0.44721359 | 20 | Dev_time |
| Maris | 49 | 2004 | TSWV | Tospoviridae | yes | CPP | yes | Solanaceae | no | thrips | 10.6 | 0.1 | 0.447213595 | 20 | 12.9 | 0.1 | 0.44721359 | 20 | Dev_time |
| Maris | 49 | 2004 | TSWV | Tospoviridae | yes | CPP | yes | Solanaceae | no | thrips | 11.1 | 0.2 | 0.894427190 | 20 | 12.7 | 0.2 | 0.89442719 | 20 | Dev_time |
| Matsuura | 50 | 2009 | TYLCV | Geminivirida | yes | CPNP | yes | Solanaceae | no | Wf | 4.5 | 0.9 | 3.117691453 | 12 | 4.9 | 0.9 | 3.11769145 | 12 | Fecundity |
| Mauck | 51 | 2010 | CMV_FNY | Bromovirida | no | NCNP | no | Cucurbitacea | no | aphid | 149.861 | 9.698 | 33.59485746 | 12 | 264.788 | 22.365 | 77.4746326 | 12 | Population |
| Mauck | 51 | 2010 | CMV_FNY | Bromovirida | no | NCNP | no | Cucurbitacea | no | aphid | 1626.46 | 168.27 | 582.9043787 | 12 | 2484.32 | 152.4 | 527.929086 | 12 | Population |
| Mauck | 51 | 2010 | CMV_FNY | Bromovirida | no | NCNP | no | Cucurbitacea | no | aphid | 46.0504 | 5.3276 | 23.82575151 | 20 | 85.1672 | 7.9934 | 35.7475715 | 20 | Population |
| Mauck | 51 | 2010 | CMV_FNY | Bromovirida | no | NCNP | no | Cucurbitacea | no | aphid | 96.3875 | 13.3245 | 59.58897553 | 20 | 181.882 | 23.983 | 107.255236 | 20 | Population |
| Mauck | 52 | 2014 | CMV_KVPG2 | Bromovirida | no | NCNP | no | Cucurbitacea | no | aphid | 100.925 | 11.329 | 39.24480719 | 12 | 49.5954 | 3.9306 | 13.6159978 | 12 | Population |
| Mauck | 52 | 2014 | CMV_KVPG2 | Bromovirida | no | NCNP | no | Cucurbitacea | no | aphid | 60.5202 | 9.8844 | 29.6532 | 9 | 26.0116 | 2.7745 | 8.3235 | 9 | Population |
| Mauck | 52 | 2014 | CMV_KVPG2 | Bromovirida | no | NCNP | no | Solanaceae | no | aphid | 119.935 | 4.873 | 15.40977903 | 10 | 141.358 | 5.305 | 18.3770590 | 12 | Population |
| Mauck | 52 | 2014 | CMV_P1 | Bromovirida | no | NCNP | no | Solanaceae | no | aphid | 109.967 | 8.329 | 26.33861063 | 10 | 92.165 | 6.1049 | 19.3053888 | 10 | Population |
| Mayer | 53 | 2002 | ToMoV | Geminivirida | yes | CPNP | yes | Solanaceae | no | Wf | 18.6 | 2.9 | 17.15663137 | 35 | 46.8 | 4.2 | 24.8475350 | 35 | Fecundity |
| Moiroux | 54 | 2018 | TuYV | Luteoviridae | yes | CPNP | yes | Brassicaceae | no | aphid | 1.5 | 0.04 | 0.178885438 | 20 | 1.41 | 0.05 | 0.22360679 | 20 | Size |
| Moiroux | 54 | 2018 | TuYV | Luteoviridae | yes | CPNP | yes | Brassicaceae | no | aphid | 0.152 | 0.009 | 0.049295030 | 30 | 0.099 | 0.005 | 0.02738612 | 30 | Weight |
| Moiroux | 54 | 2018 | TuYV | Luteoviridae | yes | CPNP | yes | Brassicaceae | no | aphid | 0.91 | 0.0253 | 0.253 | 100 | 1 | 0.01 | 0.1 | 100 | Survival |
| Moiroux | 54 | 2018 | TuYV | Luteoviridae | yes | CPNP | yes | Brassicaceae | no | aphid | 8.97 | 0.17 | 0.931128347 | 30 | 8 | 0.14 | 0.76681158 | 30 | Dev_time |
| Moiroux | 54 | 2018 | TuYV | Luteoviridae | yes | CPNP | yes | Brassicaceae | no | aphid | 4.17 | 0.12 | 0.657267069 | 30 | 4.54 | 0.24 | 1.31453413 | 30 | Fecundity |
| Moiroux | 54 | 2018 | TuYV | Luteoviridae | yes | CPNP | yes | Brassicaceae | no | aphid | 0.27 | 0.004 | 0.021908902 | 30 | 0.301 | 0.006 | 0.03286335 | 30 | R |
| Montlor | 55 | 1986 | BYDV_RPV | Luteoviridae | yes | CPNP | yes | Poaceae | no | aphid | 32.8 | 14.7 | 32.87019926 | 5 | 251 | 31 | 69.3181073 | 5 | Population |
| Montlor | 55 | 1986 | BYDV_RPV | Luteoviridae | yes | CPNP | yes | Poaceae | no | aphid | 741 | 84.3 | 188.5005305 | 5 | 597.4 | 36.9 | 82.5109083 | 5 | Population |
| Murphy | 56 | 2018 | PVY_N | Potyviridae | no | NCNP | no | Solanaceae | yes | aphid | 5 | 1.9 | 5.026927491 | 7 | 2.4 | 1.4 | 3.70405183 | 7 | Fecundity |
| Murphy | 56 | 2018 | PVY_O | Potyviridae | no | NCNP | no | Solanaceae | yes | aphid | 5 | 1.9 | 5.026927491 | 7 | 2.6 | 0.9 | 2.38117617 | 7 | Fecundity |
| Musser | 57 | 2003 | BPMV | Secoviridae | no | NCSP | no | Fabaceae | no | beetle | 7.5779 | 0.3696 | 3.092295458 | 70 | 8.7982 | 0.3044 | 2.54679312 | 70 | Weight |
| Musser | 57 | 2003 | SBMV | Sobemovirus | no | NCSP | no | Fabaceae | no | beetle | 7.5779 | 0.3696 | 3.092295458 | 70 | 9.1119 | 0.25 | 2.09165006 | 70 | Weight |
| Musser | 57 | 2003 | BPMV | Secoviridae | no | NCSP | no | Fabaceae | no | beetle | 25.2459 | 1.2519 | 15.83542161 | 160 | 32.3994 | 1.3413 | 16.9662521 | 160 | Weight |
| Musser | 57 | 2003 | SBMV | Sobemovirus | no | NCSP | no | Fabaceae | no | beetle | 25.2459 | 1.2519 | 15.83542161 | 160 | 33.5171 | 1.7437 | 22.0562542 | 160 | Weight |
| Pan | 58 | 2013 | TYLCV | Geminivirida | yes | CPNP | yes | Solanaceae | no | Wf | 766.944 | 8.324 | 45.59242568 | 30 | 789.536 | 10.702 | 58.6172681 | 30 | Size |
| Pan | 58 | 2013 | TYLCV | Geminivirida | yes | CPNP | yes | Solanaceae | no | Wf | 847.8 | 5.946 | 32.56758326 | 30 | 789.536 | 3.567 | 19.5372636 | 30 | Size |
| Pan | 58 | 2013 | TYLCV | Geminivirida | yes | CPNP | yes | Solanaceae | no | Wf | 54.4621 | 6.4931 | 35.56417338 | 30 | 94.9728 | 5.6592 | 30.9967149 | 30 | Fecundity |
| Pan | 58 | 2013 | TYLCV | Geminivirida | yes | CPNP | yes | Solanaceae | no | Wf | 114.505 | 9.324 | 51.06965126 | 30 | 101.412 | 7.825 | 42.8592901 | 30 | Fecundity |
| Pan | 58 | 2013 | TYLCV | Geminivirida | yes | CPNP | yes | Solanaceae | no | Wf | 19.6594 | 0.0661 | 0.362044610 | 30 | 18.7169 | 0.1654 | 0.90593311 | 30 | Dev_time |
| Pan | 58 | 2013 | TYLCV | Geminivirida | yes | CPNP | yes | Solanaceae | no | Wf | 18.4689 | 0.0827 | 0.452966555 | 30 | 20.0728 | 0.0992 | 0.54334077 | 30 | Dev_time |
| Pan | 58 | 2013 | TYLCV | Geminivirida | yes | CPNP | yes | Solanaceae | no | Wf | 23.3333 | 1.4121 | 7.734390234 | 30 | 26.5972 | 1.713 | 9.38248741 | 30 | Longevity |
| Pan | 58 | 2013 | TYLCV | Geminivirida | yes | CPNP | yes | Solanaceae | no | Wf | 29.4444 | 1.7825 | 9.763154587 | 30 | 26.6204 | 1.7129 | 9.38193968 | 30 | Longevity |
| Pan | 58 | 2013 | TYLCV | Geminivirida | yes | CPNP | yes | Solanaceae | no | Wf | 88.5809 | 0.396 | 2.168981327 | 30 | 92.0132 | 0.6601 | 3.61551660 | 30 | Survival |
| Pan | 58 | 2013 | TYLCV | Geminivirida | yes | CPNP | yes | Solanaceae | no | Wf | 91.9472 | 0.8581 | 4.700007265 | 30 | 86.2706 | 1.1221 | 6.14599481 | 30 | Survival |

### Performance database continued

| | | | | | | | | | | | | | | | | | | | |
|---|---|---|---|---|---|---|---|---|---|---|---|---|---|---|---|---|---|---|---|
| Pereira | 83 | 2019 | ToCV | Closteroviric | no | NCSP | yes | Solanaceae | no | Wf | 28.33 | 0.65 | 1.592168332 | 6 | 26.67 | 0.87 | 2.131056076 | 6 | Dev_time |
| Pereira | 83 | 2019 | ToCV | Closteroviric | no | NCSP | yes | Solanaceae | no | Wf | 27.33 | 0.4 | 0.979795897 | 6 | 26.67 | 0.4 | 0.979795897 | 6 | Dev_time |
| Penaflor | 59 | 2016 | SMV | Potyviridae | no | NCNP | no | Fabaceae | no | aphid | 17.0575 | 0.9942 | 3.143936449 | 10 | 23.0246 | 1.7486 | 5.529558716 | 10 | Weight |
| Penaflor | 59 | 2016 | SMV | Potyviridae | no | NCNP | no | Fabaceae | no | aphid | 146.555 | 9.154 | 28.94748970 | 10 | 106.691 | 10.299 | 32.56829762 | 10 | Population |
| Penaflor | 59 | 2016 | BPMV | Secoviridae | no | NCSP | no | Fabaceae | no | beetle | 17.0575 | 0.9942 | 3.143936449 | 10 | 18.4639 | 1.0629 | 3.361184924 | 10 | Weight |
| Penaflor | 59 | 2016 | BPMV | Secoviridae | no | NCSP | no | Fabaceae | no | beetle | 146.555 | 9.154 | 28.94748970 | 10 | 131.657 | 8.924 | 28.22016583 | 10 | Population |
| Ponsen | 60 | 1969 | PLRV | Luteoviridae | yes | CPNP | yes | Solanaceae | yes | aphid | 13 | 0.5044 | 2.139987962 | 18 | 36 | 0.985 | 3.94 | 16 | Fecundity |
| Ponsen | 60 | 1969 | PLRV | Luteoviridae | yes | CPNP | yes | Solanaceae | yes | aphid | 11 | 0.3642 | 1.030113158 | 8 | 10 | 0.0525 | 0.21 | 16 | Dev_time |
| Ponsen | 60 | 1969 | PLRV | Luteoviridae | yes | CPNP | yes | Solanaceae | yes | aphid | 7 | 0.4525 | 1.279863273 | 8 | 19 | 0.705 | 2.82 | 16 | Longevity |
| Porras | 61 | 2018 | BYDV | Luteoviridae | yes | CPNP | yes | Poaceae | no | aphid | 22.62 | 26.52 | 102.7115183 | 15 | 42.37 | 46.01 | 178.1959637 | 15 | Fecundity |
| Porras | 61 | 2018 | BYDV | Luteoviridae | yes | CPNP | yes | Poaceae | no | aphid | 18.06 | 20.72 | 80.24821493 | 15 | 41.95 | 46.57 | 180.3648344 | 15 | Fecundity |
| Porras | 61 | 2018 | BYDV | Luteoviridae | yes | CPNP | yes | Poaceae | no | aphid | 34.65 | 38.16 | 147.7930444 | 15 | 72.46 | 76.2 | 295.1213309 | 15 | Fecundity |
| Porras | 61 | 2018 | BYDV | Luteoviridae | yes | CPNP | yes | Poaceae | no | aphid | 26.63 | 28.79 | 111.5031905 | 15 | 68.83 | 74.68 | 289.2343962 | 15 | Fecundity |
| Ren | 62 | 2015 | PVY | Potyviridae | no | NCNP | no | Solanaceae | no | aphid | 2.06 | 0.03 | 0.116189500 | 15 | 1.75 | 0.02 | 0.077459666 | 15 | Size |
| Ren | 62 | 2015 | PVY | Potyviridae | no | NCNP | no | Solanaceae | no | aphid | 28.7 | 1.6 | 2.771281292 | 3 | 36.3 | 1.1 | 1.905255888 | 3 | R |
| Ren | 62 | 2015 | PVY | Potyviridae | no | NCNP | no | Solanaceae | no | aphid | 10.7 | 0.9 | 1.558845726 | 3 | 10.8 | 0.6 | 1.039230484 | 3 | T |
| Ren | 62 | 2015 | PVY | Potyviridae | no | NCNP | no | Solanaceae | no | aphid | 0.33 | 0.03 | 0.051961524 | 3 | 0.35 | 0.04 | 0.069282032 | 3 | R |
| Ren | 62 | 2015 | PVY | Potyviridae | no | NCNP | no | Solanaceae | no | aphid | 1.39 | 0.02 | 0.034641016 | 3 | 1.47 | 0.05 | 0.086602540 | 3 | Lambda |
| Salvauson | 63 | 2013 | ZYMV | Potyviridae | no | NCNP | no | Cucurbitacea | no | aphid | 65.9322 | 12.7119 | 40.19855738 | 10 | 128.136 | 12.033 | 38.05168708 | 10 | Population |
| Salvauson | 63 | 2013 | WMV | Potyviridae | no | NCNP | no | Cucurbitacea | no | aphid | 65.9322 | 12.7119 | 40.19855738 | 10 | 64.0678 | 8.1356 | 25.72702613 | 10 | Population |
| Shalileh | 64 | 2016 | TSWV | Tospoviridae | yes | CPP | yes | Solanaceae | no | thrips | 8.9639 | 0.0513 | 0.229420574 | 20 | 9.4649 | 0.5142 | 2.299572308 | 20 | Dev_time |
| Shalileh | 64 | 2016 | TSWV | Tospoviridae | yes | CPP | yes | Solanaceae | no | thrips | 13.1611 | 0.1261 | 0.690678145 | 30 | 10.8441 | 0.6936 | 3.799003658 | 30 | Longevity |
| Shalileh | 64 | 2016 | TSWV | Tospoviridae | yes | CPP | yes | Solanaceae | no | thrips | 3.8085 | 0.0052 | 0.016443843 | 10 | 2.8854 | 0.0103 | 0.032571459 | 10 | Fecundity |
| Shi | 65 | 2016 | CMV | Bromovirida | no | NCNP | no | Solanaceae | no | aphid | 60.7716 | 9.2573 | 26.18359842 | 8 | 74.6524 | 10.0257 | 28.35696182 | 8 | Population |
| Shi | 65 | 2016 | CMV | Bromovirida | no | NCNP | no | Solanaceae | no | aphid | 155.476 | 18.502 | 52.33155866 | 8 | 239.532 | 16.195 | 45.80637728 | 8 | Population |
| Shi | 65 | 2016 | CMV | Bromovirida | no | NCNP | no | Solanaceae | no | aphid | 286.424 | 16.193 | 45.80072043 | 8 | 333.473 | 16.19 | 45.79223514 | 8 | Population |
| Shi | 65 | 2016 | CMV | Bromovirida | no | NCNP | no | Solanaceae | no | aphid | 418.147 | 14.65 | 41.43645737 | 8 | 368.79 | 17.736 | 50.16498348 | 8 | Population |
| Shi | 65 | 2016 | CMV | Bromovirida | no | NCNP | no | Solanaceae | no | aphid | 489.382 | 19.691 | 55.69455851 | 8 | 403.668 | 21.429 | 60.61036485 | 8 | Population |
| Shi | 65 | 2016 | CMV | Bromovirida | no | NCNP | no | Solanaceae | no | aphid | 13.9197 | 0.3442 | 0.973544616 | 8 | 12.5048 | 0.2485 | 0.702864140 | 8 | Longevity |
| Shrestha | 66 | 2017 | SVYV | Potyviridae | no | NCSP | yes | Cucurbitacea | no | Wf | 8.59 | 0.09 | 0.569209978 | 40 | 6.85 | 0.093 | 0.588183644 | 40 | Longevity |
| Shrestha | 66 | 2017 | SVYV | Potyviridae | no | NCSP | yes | Cucurbitacea | no | Wf | 7.85 | 0.09 | 0.569209978 | 40 | 9.97 | 0.086 | 0.543911757 | 40 | Longevity |
| Shrestha | 66 | 2017 | SVYV | Potyviridae | no | NCSP | yes | Cucurbitacea | no | Wf | 50.35 | 6.91 | 43.70267726 | 40 | 76.35 | 9.34 | 59.07134669 | 40 | Fecundity |
| Shrestha | 66 | 2017 | SVYV | Potyviridae | no | NCSP | yes | Cucurbitacea | no | Wf | 23.5 | 0.2 | 0.979795897 | 24 | 20.4 | 0.2 | 0.979795897 | 24 | Dev_time |
| Shrestha | 66 | 2017 | SVYV | Potyviridae | no | NCSP | yes | Cucurbitacea | no | Wf | 864 | 3 | 14.69693845 | 24 | 862 | 4 | 19.59591794 | 24 | Size |
| Shrestha | 66 | 2017 | SVYV | Potyviridae | no | NCSP | yes | Cucurbitacea | no | Wf | 970 | 3 | 14.69693845 | 24 | 963 | 4 | 19.59591794 | 24 | Size |
| Srinivasan | 67 | 2007 | PLRV | Luteoviridae | yes | CPNP | yes | Solanaceae | no | aphid | 5.55457 | 0.99379 | 3.442589544 | 12 | 14.9423 | 0.8696 | 3.012382764 | 12 | Fecundity |
| Srinivasan | 67 | 2007 | PVY | Potyviridae | no | NCNP | no | Solanaceae | no | aphid | 5.55457 | 0.99379 | 3.442589544 | 12 | 8.35847 | 0.9583 | 3.319648577 | 12 | Fecundity |
| Srinivasan | 67 | 2007 | PLRV | Luteoviridae | yes | CPNP | yes | Solanaceae | no | aphid | 3.6597 | 0.91323 | 3.163521517 | 12 | 14.4913 | 2.6988 | 9.348917438 | 12 | Fecundity |
| Srinivasan | 67 | 2007 | PVY | Potyviridae | no | NCNP | no | Solanaceae | no | aphid | 3.6597 | 0.91323 | 3.163521517 | 12 | 4.76838 | 1.78556 | 6.185361279 | 12 | Fecundity |
| Stumpf | 68 | 2007 | TSWV | Tospoviridae | yes | CPP | yes | Asteraceae | yes | thrips | 63.0675 | 2.2951 | 21.77323037 | 90 | 71.8481 | 2.2511 | 21.35580972 | 90 | Survival |
| Stumpf | 68 | 2007 | TSWV | Tospoviridae | yes | CPP | yes | Asteraceae | yes | thrips | 63.0675 | 2.2951 | 21.77323037 | 90 | 66.4793 | 2.207 | 20.93744038 | 90 | Survival |
| Su | 69 | 2016 | TYLCV | Geminivirida | yes | CPNP | yes | Solanaceae | no | Wf | 249.842 | 20.19 | 110.5851843 | 30 | 576.236 | 78.233 | 428.4997884 | 30 | Population |
| Su | 70 | 2015 | TYLCV | Geminivirida | yes | CPNP | yes | Solanaceae | no | Wf | 30.5 | 2.6 | 11.62755348 | 20 | 51.3 | 3.1 | 13.86362146 | 20 | Weight |
| Su | 70 | 2015 | TYLCV | Geminivirida | yes | CPNP | yes | Solanaceae | no | Wf | 24.2 | 2.6 | 10.06975670 | 15 | 43.9 | 3.1 | 12.00624837 | 15 | Fecundity |
| Su | 70 | 2015 | TYLCV | Geminivirida | yes | CPNP | yes | Solanaceae | no | Wf | 70.7 | 3.6 | 13.94274004 | 15 | 83.3 | 3.6 | 13.94274004 | 15 | Survival |
| Sun | 71 | 2017 | TYLCV | Geminivirida | yes | CPNP | yes | Solanaceae | no | Wf | 25.3 | 3.3 | 12.78084504 | 15 | 25.1 | 2.4 | 10.46135746 | 19 | Fecundity |
| Sun | 71 | 2017 | TYLCV | Geminivirida | yes | CPNP | yes | Solanaceae | no | Wf | 27 | 2.1 | 8.4 | 16 | 39.4 | 3.8 | 15.2 | 16 | Fecundity |
| Sun | 71 | 2017 | TYLCV | Geminivirida | yes | CPNP | yes | Solanaceae | no | Wf | 58 | 5.8 | 22.46330340 | 15 | 73.7 | 4.5 | 19.61504524 | 19 | Survival |
| Sun | 71 | 2017 | TYLCV | Geminivirida | yes | CPNP | yes | Solanaceae | no | Wf | 76.9 | 4.6 | 18.4 | 16 | 63.8 | 6.1 | 24.4 | 16 | Survival |
| Sun | 71 | 2017 | TYLCV | Geminivirida | yes | CPNP | yes | Solanaceae | no | Wf | 60.7 | 6.4 | 24.78709341 | 15 | 69 | 4.1 | 17.87148566 | 19 | Survival |
| Sun | 71 | 2017 | TYLCV | Geminivirida | yes | CPNP | yes | Solanaceae | no | Wf | 68.1 | 5.2 | 20.8 | 16 | 51.9 | 7.1 | 28.4 | 16 | Survival |
| Sylvester | 72 | 1973 | SYVV | Tospoviridae | yes | CPP | yes | Asteraceae | yes | aphid | 39.5 | 1.6347 | 1.6347 | 112 | 40.1 | 1.8805 | 1.8805 | 30 | Fecundity |
| Sylvester | 72 | 1973 | SYVV | Tospoviridae | yes | CPP | yes | Asteraceae | yes | aphid | 39.5 | 1.6347 | 1.6347 | 112 | 28.1 | 2.1084 | 2.1084 | 46 | Fecundity |
| Sylvester | 72 | 1973 | SYVV | Tospoviridae | yes | CPP | yes | Asteraceae | yes | aphid | 18.8 | 0.4063 | 0.4063 | 112 | 14.9 | 0.5295 | 0.5295 | 30 | Longevity |
| Sylvester | 72 | 1973 | SYVV | Tospoviridae | yes | CPP | yes | Asteraceae | yes | aphid | 18.8 | 0.4063 | 0.4063 | 112 | 13.5 | 0.3539 | 0.3539 | 46 | Longevity |
| Wang | 73 | 2012 | TYLCCNV | Geminivirida | yes | CPNP | yes | Solanaceae | no | Wf | 205.9 | 58.3 | 174.9 | 9 | 396.6 | 27.3 | 81.9 | 9 | Fecundity |
| Wang | 73 | 2012 | TYLCCNV | Geminivirida | yes | CPNP | yes | Solanaceae | no | Wf | 62.9 | 11.1 | 33.3 | 9 | 91.4 | 4 | 12 | 9 | Survival |
| Wang | 73 | 2012 | TYLCCNV | Geminivirida | yes | CPNP | yes | Solanaceae | no | Wf | 22.9 | 8.1 | 24.3 | 9 | 57.1 | 8.1 | 24.3 | 9 | Survival |
| Wang | 73 | 2012 | TYLCCNV | Geminivirida | yes | CPNP | yes | Solanaceae | no | Wf | 148.2 | 30.1 | 90.3 | 9 | 347.7 | 27.7 | 83.1 | 9 | Fecundity |
| Wang | 73 | 2012 | TYLCCNV | Geminivirida | yes | CPNP | yes | Solanaceae | no | Wf | 37.8 | 8.5 | 25.5 | 9 | 84.4 | 5.6 | 16.8 | 9 | Survival |
| Wang | 73 | 2012 | TYLCCNV | Geminivirida | yes | CPNP | yes | Solanaceae | no | Wf | 17.8 | 8.5 | 25.5 | 9 | 64.4 | 7.3 | 21.9 | 9 | Survival |
| Wang | 73 | 2012 | TYLCCNV | Geminivirida | yes | CPNP | yes | Solanaceae | no | Wf | 154.9 | 18.5 | 55.5 | 9 | 218.9 | 22.6 | 67.8 | 9 | Fecundity |
| Wang | 73 | 2012 | TYLCCNV | Geminivirida | yes | CPNP | yes | Solanaceae | no | Wf | 48.8 | 6.3 | 18.9 | 9 | 76.3 | 3.3 | 9.9 | 9 | Survival |
| Wang | 73 | 2012 | TYLCCNV | Geminivirida | yes | CPNP | yes | Solanaceae | no | Wf | 22.5 | 6 | 18 | 9 | 33.8 | 5.1 | 15.3 | 9 | Survival |
| Westwood | 74 | 2013 | CMV | Bromovirida | no | NCNP | yes | Brassicaceae | no | aphid | 0.442267 | 0.017845 | 0.087422288 | 24 | 0.357593 | 0.018894 | 0.092561318 | 24 | Weight |
| Williams | 75 | 1995 | BYV | Closteroviric | no | NCSP | yes | Chenopodia | no | aphid | 0.3 | 0.15 | 0.821583836 | 30 | 6.9 | 1.73 | 9.475600244 | 30 | Fecundity |
| Williams | 75 | 1995 | BYV | Closteroviric | no | NCSP | yes | Chenopodia | no | aphid | 0.01 | 0.01 | 0.054772255 | 30 | 7.9 | 2.57 | 14.07646972 | 30 | Fecundity |
| Williams | 75 | 1995 | BYV | Closteroviric | no | NCSP | yes | Chenopodia | no | aphid | 2.1 | 0.37 | 2.026573462 | 30 | 39.3 | 6.07 | 33.24675924 | 30 | Fecundity |
| Williams | 75 | 1995 | BYV | Closteroviric | no | NCSP | yes | Chenopodia | no | aphid | 0.5 | 0.29 | 1.588395416 | 30 | 50.6 | 2.64 | 14.45987551 | 30 | Fecundity |
| Williams | 75 | 1995 | BYV | Closteroviric | no | NCSP | yes | Chenopodia | no | aphid | 1.4 | 0.93 | 5.093819784 | 30 | 21.8 | 3.15 | 17.25326056 | 30 | Fecundity |
| Williams | 75 | 1995 | BYV | Closteroviric | no | NCSP | yes | Chenopodia | no | aphid | 178.826 | 1.334 | 5.965829363 | 20 | 175.494 | 1.997 | 8.930855502 | 20 | Dev_time |
| Williams | 75 | 1995 | BYV | Closteroviric | no | NCSP | yes | Chenopodia | no | aphid | 22.6751 | 2.706 | 12.10159989 | 20 | 65.6589 | 4.4165 | 19.75118844 | 20 | Fecundity |
| Williams | 75 | 1995 | BYV | Closteroviric | no | NCSP | yes | Chenopodia | no | aphid | 1.14729 | 0.09578 | 0.428341181 | 20 | 1.43203 | 0.12313 | 0.550654100 | 20 | R |
| Williams | 75 | 1995 | BYV | Closteroviric | no | NCSP | yes | Chenopodia | no | aphid | 205.991 | 10.39 | 46.46549257 | 20 | 530.541 | 43.888 | 196.2731027 | 20 | Reprod_peri |
| Williams | 75 | 1995 | BYV | Closteroviric | no | NCSP | yes | Chenopodia | no | aphid | 216.433 | 5.997 | 26.81939932 | 20 | 206.706 | 4.797 | 21.45283617 | 20 | Dev_time |
| Williams | 75 | 1995 | BYV | Closteroviric | no | NCSP | yes | Chenopodia | no | aphid | 19.4203 | 2.9192 | 13.05505927 | 20 | 62.0473 | 4.4143 | 19.74134974 | 20 | Fecundity |
| Williams | 75 | 1995 | BYV | Closteroviric | no | NCSP | yes | Chenopodia | no | aphid | 0.9055 | 0.054776 | 0.244965719 | 20 | 1.25991 | 0.07709 | 0.344756960 | 20 | R |
| Williams | 75 | 1995 | BYV | Closteroviric | no | NCSP | yes | Chenopodia | no | aphid | 211.058 | 20.209 | 90.37739551 | 20 | 501.535 | 40.424 | 180.7816238 | 20 | Reprod_peri |
| Williams | 75 | 1995 | BYV | Closteroviric | no | NCSP | yes | Chenopodia | no | aphid | 212.249 | 5.02 | 22.45012249 | 20 | 234.337 | 7.229 | 32.32907081 | 20 | Dev_time |
| Williams | 75 | 1995 | BYV | Closteroviric | no | NCSP | yes | Chenopodia | no | aphid | 7.3362 | 0.9602 | 4.294144943 | 20 | 55.7614 | 5.5469 | 24.80649092 | 20 | Fecundity |
| Williams | 75 | 1995 | BYV | Closteroviric | no | NCSP | yes | Chenopodia | no | aphid | 0.5685 | 0.0597 | 0.266986516 | 20 | 1.4858 | 0.0765 | 0.342118400 | 20 | R |
| Williams | 75 | 1995 | BYV | Closteroviric | no | NCSP | yes | Chenopodia | no | aphid | 139.872 | 14.423 | 64.50161687 | 20 | 474.497 | 45.006 | 201.2729507 | 20 | Reprod_peri |
| Williams | 75 | 1995 | BYV | Closteroviric | no | NCSP | yes | Chenopodia | no | aphid | 268.077 | 13.721 | 61.36217743 | 20 | 244.121 | 25.603 | 114.5000968 | 20 | Dev_time |
| Williams | 75 | 1995 | BYV | Closteroviric | no | NCSP | yes | Chenopodia | no | aphid | 6.946 | 2.49343 | 11.15095795 | 20 | 66.5134 | 5.1955 | 23.23498235 | 20 | Fecundity |
| Williams | 75 | 1995 | BYV | Closteroviric | no | NCSP | yes | Chenopodia | no | aphid | 0.450829 | 0.052252 | 0.233678047 | 20 | 1.45182 | 0.09203 | 0.411570671 | 20 | R |
| Williams | 75 | 1995 | BYV | Closteroviric | no | NCSP | yes | Chenopodia | no | aphid | 139.736 | 27.14 | 121.3737698 | 20 | 589.025 | 40.997 | 183.3441577 | 20 | Reprod_peri |
| Wosula | 76 | 2013 | SPFMV | Potyviridae | no | NCNP | no | Convolvulac | no | aphid | 0.338 | 0.01 | 0.044721359 | 20 | 0.338 | 0.011 | 0.049193495 | 20 | R |
| Wosula | 76 | 2013 | SPFMV | Potyviridae | no | NCNP | no | Convolvulac | no | aphid | 8.8 | 0.3 | 1.341640786 | 20 | 9.1 | 0.2 | 0.894427190 | 20 | T |
| Wosula | 76 | 2013 | SPFMV | Potyviridae | no | NCNP | no | Convolvulac | no | aphid | 19.7 | 0.01 | 0.044721359 | 20 | 22 | 0.01 | 0.044721359 | 20 | T0 |
| Wosula | 76 | 2013 | SPFMV | Potyviridae | no | NCNP | no | Convolvulac | yes | aphid | 0.319 | 0.013 | 0.058137767 | 20 | 0.276 | 0.016 | 0.071554175 | 20 | R |
| Wosula | 76 | 2013 | SPFMV | Potyviridae | no | NCNP | no | Convolvulac | yes | aphid | 9.9 | 0.4 | 1.788854381 | 20 | 10.2 | 0.6 | 2.683281572 | 20 | T |
| Wosula | 76 | 2013 | SPFMV | Potyviridae | no | NCNP | no | Convolvulac | yes | aphid | 21.2 | 0.01 | 0.044721359 | 20 | 16.2 | 0.2 | 0.894427190 | 20 | T |
| Wosula | 76 | 2013 | SPFMV | Potyviridae | no | NCNP | no | Convolvulac | yes | aphid | 0.109 | 0.038 | 0.169941166 | 20 | #FIELD! | 0.035 | 0.156524758 | 20 | R |
| Wosula | 76 | 2013 | SPFMV | Potyviridae | no | NCNP | no | Convolvulac | yes | aphid | 11 | 4.6 | 20.57182539 | 20 | 13.5 | 8.9 | 39.80200999 | 20 | T |
| Wosula | 76 | 2013 | SPFMV | Potyviridae | no | NCNP | no | Convolvulac | yes | aphid | 3.3 | 1.2 | 5.366563145 | 20 | 0.7 | 0.1 | 0.447213595 | 20 | T |
| Wu | 77 | 2014 | PEMV | Luteoviridae | yes | CPNP | yes | Fabaceae | no | aphid | 15.37 | 0.27 | 1.909188309 | 50 | 14.64 | 0.32 | 2.262741699 | 50 | Dev_time |
| Wu | 77 | 2014 | BLRV | Luteoviridae | yes | CPNP | yes | Fabaceae | no | aphid | 15.37 | 0.27 | 1.909188309 | 50 | 14.42 | 0.32 | 2.262741699 | 50 | Dev_time |
| Wu | 77 | 2014 | PEMV | Luteoviridae | yes | CPNP | yes | Fabaceae | no | aphid | 36.39 | 0.77 | 5.444722215 | 50 | 33.76 | 0.76 | 5.374011537 | 50 | Longevity |
| Wu | 77 | 2014 | BLRV | Luteoviridae | yes | CPNP | yes | Fabaceae | no | aphid | 36.39 | 0.77 | 5.444722215 | 50 | 36.7 | 0.75 | 5.303300858 | 50 | Longevity |
| Wu | 77 | 2014 | PEMV | Luteoviridae | yes | CPNP | yes | Fabaceae | no | aphid | 7.848 | 1.392 | 9.842926394 | 50 | 8.544 | 1.416 | 10.01263202 | 50 | Fecundity |
| Wu | 77 | 2014 | BLRV | Luteoviridae | yes | CPNP | yes | Fabaceae | no | aphid | 7.848 | 1.392 | 9.842926394 | 50 | 12.024 | 1.296 | 9.164103884 | 50 | Fecundity |
| Wu | 77 | 2014 | PEMV | Luteoviridae | yes | CPNP | yes | Fabaceae | no | aphid | 26.0745 | 1.1524 | 8.148698546 | 50 | 26.2546 | 1.0088 | 7.133293208 | 50 | Fecundity |
| Wu | 77 | 2014 | BLRV | Luteoviridae | yes | CPNP | yes | Fabaceae | no | aphid | 26.0745 | 1.1524 | 8.148698546 | 50 | 31.3325 | 1.0447 | 7.387144543 | 50 | Fecundity |
| Wu | 77 | 2014 | PEMV | Luteoviridae | yes | CPNP | yes | Fabaceae | no | aphid | 25.104 | 0.984 | 6.957930726 | 50 | 23.928 | 1.056 | 7.467047609 | 50 | Fecundity |
| Wu | 77 | 2014 | BLRV | Luteoviridae | yes | CPNP | yes | Fabaceae | no | aphid | 25.104 | 0.984 | 6.957930726 | 50 | 30.096 | 1.056 | 7.467047609 | 50 | Fecundity |
| Wu | 77 | 2014 | PEMV | Luteoviridae | yes | CPNP | yes | Fabaceae | no | aphid | 18.84 | 1.176 | 8.315575746 | 50 | 16.056 | 1.32 | 9.333809511 | 50 | Fecundity |
| Wu | 77 | 2014 | BLRV | Luteoviridae | yes | CPNP | yes | Fabaceae | no | aphid | 18.84 | 1.176 | 8.315575746 | 50 | 24.864 | 1.176 | 8.315575746 | 50 | Fecundity |

### Performance database continued

| | | | | | | | | | | | | | | | | | | | |
|---|---|---|---|---|---|---|---|---|---|---|---|---|---|---|---|---|---|---|---|
| Zhang | 78 | 2014 | SRBSDV | Reoviridae | yes | CPP | yes | Poaceae | no | planthopper | 108.85 | 16.25 | 28.145825622 | 3 | 99.32 | 18.58 | 32.181504004 | 3 | Fecundity |
| Zhang | 78 | 2014 | SRBSDV | Reoviridae | yes | CPP | yes | Poaceae | no | planthopper | 122.5 | 16.44 | 28.474915276 | 3 | 107.35 | 18.42 | 31.904375875 | 3 | Fecundity |
| Zhang | 78 | 2014 | SRBSDV | Reoviridae | yes | CPP | yes | Poaceae | no | planthopper | 88.45 | 2.24 | 3.8797938089 | 3 | 85.12 | 2.82 | 4.8843832773 | 3 | Survival |
| Zhang | 78 | 2014 | SRBSDV | Reoviridae | yes | CPP | yes | Poaceae | no | planthopper | 90.84 | 1.15 | 1.9918584287 | 3 | 86.52 | 1.42 | 2.4595121467 | 3 | Survival |
| Zhang | 78 | 2014 | SRBSDV | Reoviridae | yes | CPP | yes | Poaceae | no | planthopper | 12.95 | 0.21 | 0.3637306695 | 3 | 10.12 | 0.14 | 0.2424871130 | 3 | Dev_time |
| Zhang | 79 | 2012 | TYLCCNV | Geminivirida | yes | CPNP | yes | Solanaceae | no | Wf | 267.35 | 49.167 | 163.06849106 | 11 | 877.337 | 224.323 | 743.99522284 | 11 | Population |
| Ziebel | 80 | 2011 | CMV | Bromovirida | no | NCNP | no | Solanaceae | no | aphid | 0.28537 | 0.047628 | 0.047628 | 20 | 0.218225 | 0.059775 | 0.059775 | 20 | Weight |

### Start of content for file ESM_model parameters.docx

**Electronic Supplementary Materials: “Model parameters”**

**Description of the model modified from Shaw et al. 2017**

The model explored the preference and performance of vectors before (non-viruliferous) and after acquiring a virus (viruliferous). It tracks the number of viruliferous (*V*) and non-viruliferous (*N*) vectors, and the fraction of healthy (*H*) and infected (*I*) hosts using a system of ordinary differential equations. To incorporate behavioral preferences, the model tracks the infection status of the host that each vector is on. There are four compartments for the vector population: $N_h$ number of non-viruliferous vectors on healthy hosts, $N_i$ number of non-viruliferous vectors on infected hosts, $V_h$ number of viruliferous vectors on healthy hosts, and $V_i$ number of viruliferous vectors on infected hosts. To represent virus retention mechanisms mathematically, the model includes a vector recovery rate ($\gamma$) that approximates the two retention mechanisms. Vectors retain CPNPr viruses indefinitely following acquisition, while vectors lose NCNP viruses on a daily basis. The original model (Shaw et al. 2017) assumed that ($\gamma$) is the rate at which vectors recover by cleaning their stylets while feeding, which was presumed to occur when the vector fed on a healthy plant for NPL viruses. To mimic loss of virions during phloem-feeding in infected hosts, we modified the ($\gamma$) term for NPL viruses to include recovery when the vector feeds on infected plants as well (see model equations - red text).

To account for the fact that feeding on an infected host does not always result in a vector becoming viruliferous, the model includes a component describing the rate at which vectors become viruliferous for each transmission mechanism ($\beta_v$), represented as a proportion of vectors successfully capable of transmitting virus following acquisition. Likewise, to account for the fact that probing or feeding on a susceptible host does not always result in infection, the model includes a “rate healthy hosts become infected” parameter ($\beta_i$). Values for this parameter are based on empirically derived transmission rates of viruses to highly susceptible hosts by vectors known to be able to feed, colonize, and reproduce on that host. The recovery rate ($\gamma$), successful acquisition rate ($\beta_v$), and healthy host infection rate ($\beta_i$) together represent the biological characteristics (virus traits) of aphid-transmitted pathogens with CPNPr and NCNP transmission mechanisms.

The modified model vector dynamics equations:

$$\frac{dN_h}{dt} = \underbrace{r_h[N_h + V_h]\left[1 - \frac{N_h + V_h}{K_h HF}\right]}_{\text{growth rate}} - \underbrace{\alpha_{nh} N_h [1-\mu] I^{\delta}}_{\text{move to I}} - \underbrace{\alpha_{nh} N_h \mu}_{\substack{\text{loss}\\\text{dispersal}}} + \underbrace{\alpha_{ni} N_i [1-\mu][1 - I^{\delta}]}_{\text{move to H}} - \underbrace{\beta_i \rho_{vh} N_h}_{\substack{\text{host}\\\text{infect}}} \quad (1a)$$

$$\frac{dN_i}{dt} = \underbrace{\alpha_{nh} N_h [1-\mu] I^{\delta}}_{\text{move to I}} - \underbrace{\alpha_{ni} N_i \mu}_{\substack{\text{loss}\\\text{dispersal}}} - \underbrace{\alpha_{ni} N_i [1 - I^{\delta}][1-\mu]}_{\text{move to H}} + \underbrace{\beta_i \rho_{vh} N_h}_{\substack{\text{host}\\\text{infect}}} - \underbrace{\beta_v N_i}_{\substack{\text{vector}\\\text{infect}}} + \underbrace{\gamma V_h + \gamma V_i}_{\substack{\text{vectors}\\\text{recovered}}} \quad (1b)$$

$$\frac{dV_i}{dt} = \underbrace{r_i[N_i + V_i]\left[1 - \frac{N_i + V_i}{K_i IF}\right]}_{\text{growth rate}} - \underbrace{\alpha_{vi} V_i [1-\mu] H^{\epsilon}}_{\text{move to H}} - \underbrace{\alpha_{vi} V_i \mu}_{\substack{\text{loss}\\\text{dispersal}}} + \underbrace{\alpha_{vh} V_h [1-\mu][1 - H^{\epsilon}]}_{\text{move to I}} + \underbrace{\beta_i \rho_{vh} V_h}_{\substack{\text{host}\\\text{infect}}} + \underbrace{\beta_v N_i}_{\substack{\text{vector}\\\text{infect}}} \underbrace{-\gamma V_i}_{\substack{\text{vectors}\\\text{recovered}}} \quad (1c)$$

$$\frac{dV_h}{dt} = \underbrace{\alpha_{vi} V_i [1-\mu] H^{\epsilon}}_{\text{move to H}} - \underbrace{\alpha_{vh} V_h \mu}_{\text{dispersal loss}} - \underbrace{\alpha_{vh} V_h [1-\mu][1 - H^{\epsilon}]}_{\text{move to I}} - \underbrace{\beta_i \rho_{vh} V_h}_{\substack{\text{host}\\\text{infect}}} - \underbrace{\gamma V_h}_{\substack{\text{vectors}\\\text{recovered}}} \quad (1d)$$

along with the host dynamics equations:

$$\frac{dH}{dt} = - \underbrace{\beta_i \rho_{vh}}_{\text{infect host}} \quad (2a)$$

$$\frac{dI}{dt} = \underbrace{\beta_i \rho_{vh}}_{\text{infect host}} \quad (2b)$$

where

$$\rho_h = \frac{V_h + N_h}{HF + \xi}, \qquad \rho_i = \frac{V_i + N_i}{IF + \xi}, \qquad \text{and} \qquad \rho_{vh} = \frac{V_h}{HF + \xi} \quad (3)$$

We modified several of the parameter values to better reflect our data, while keeping the model compartments intact. The original model included a vector dispersal loss term ($\mu$) with a value of 99.4%, which was derived from a study describing mortality of fall aphid migrants attempting to locate primary hosts (Ward, Leather, Pickup, & Harrington, 1998). However, this model describes local movements from one plant to neighboring hosts. Therefore, we substituted the value of 54% +/- 21 based on dispersal loss of aphids in three iterations of a mark-recapture field study (Power, 1991). The vector recovery rate ($\gamma$) was left at 0 and 1 for CPNPr and NCNP viruses, respectively. However, we did calculate new values for the rate at which vectors become viruliferous ($\beta_v$). The original model included $\beta_v$ values of 0.68 for a CPNPr virus (i.e., maximum transmission efficiency reported in Jiménez-Martínez & Bosque-Pérez (2004)) and 1 for a NCNP virus. We substituted the value of 0.54 +/- 0.14 based on transmission efficiency on three different plant genotypes for CPNPr (Jiménez-Martínez & Bosque-Pérez 2004) and 0.22 +/- 0.22 based on transmission efficiency of three non-persistent viruses reported in Wang & Ghabrial (2002). We did not change the “rate healthy hosts become infected” parameter values ($\beta_i$). The remaining parameters include those corresponding to vector responses as modified (or not) by virus infection in the host plant, including vector orientation preference for infected hosts ($\delta$) (attraction), maximum vector departure rate from infected hosts ($a_i$) (settling and feeding), and intrinsic vector growth rate on infected hosts ($r_i$). To make these estimates more representative of CPNPr and NCNP pathogens, we consulted the meta-analysis database and calculated new values using data extracted from all publications on aphid-transmitted CPNPr and NCNP pathogens (electronic supplementary material file ESM_model parameters.docx and ESM_Model_raw data.xlsx). The updated values are available in Table 1 and original parameters are available in (Shaw et al., 2017).

**Table S1.** Model parameters used in this study (adapted from Shaw et al., 2017).

| | PL-CPNPr | | | NPL-NCNP | | |
|---|---|---|---|---|---|---|
| **Parameters** | **Values** | **SD** | **(CI.lb-CI.ub)** | **Values** | **SD** | **(CI.lb-CI.ub)** |
| $r_h$ intrinsic vector growth rate on healthy hosts (n = 38) | 0.259 | 0.066 | | 0.259 | 0.066 | |
| $r_i$ intrinsic vector growth rate on infected hosts | 0.288 | | (0.272-0.305) | 0.254 | | (0.237-0.272) |
| $K_h$ vector carrying capacity on healthy hosts | 100 | | | 100 | | |
| $K_i$ vector carrying capacity on infected hosts | 100 | | | 100 | | |
| $F$ field density | 4 (2.5-5) | | | | | |
| | | | | 4 (2.5-5) | | |
| $\xi$ minimum host/ha needed to support the vectors | 0.01$F$ | | | 0.01$F$ | | |
| $a_h$ maximum vector departure rate from healthy hosts (n = 32) | 0.423 | 0.217 | | 0.423 | 0.217 | |
| $a_i$ maximum vector departure rate from infected hosts | 0.243 | | (0.163-0.324) | 0.347 | | (0.265-0.428) |
| $c_1$ same status half-departure constant | 13.76 | | | 13.76 | | |
| $c_2$ different status half-departure constant | 0.3 | | | 0.3 | | |
| $\delta$ preference of nonvirulent vectors for settling on healthy hosts | 0.782 | 0.196 | | 0.719 | 0.063 | |
| $\varepsilon$ preference of virulent vectors for settling on infected hosts (*<u>Main Model</u>*) | 1.218 | 0.196 | (1.022-1.414) | 1.281 | 0.063 | (1.218-1.344) |
| $\mathcal{E}$ preference of virulent vectors for settling on infected hosts *<u>(Model 2)</u>* | *1* | | (0.804-1.196) | *1* | | (0.937-1.063) |
| $\mathcal{E}$ preference of virulent vectors for settling on infected hosts *<u>(Model 3)</u>* | *0.782* | *0.196* | (0.586-0.978) | *0.719* | *0.063* | (0.656-0.782) |
| $\mu$ dispersal loss | 53.67 | 21.39 | | 53.67 | 21.39 | |
| $\beta_v$ rate vector on infected host becomes virulent | 0.543 | 0.14 | | 0.215 | 0.224 | (0.008-0.457) |
| $\beta_i$ rate healthy hosts become infected | 1 | | | 0.182 | | |
| $\gamma$ vector recovery rate by feeding | 0 | | | 1 | | |

To explore the possible influence of conditional vector preferences in the context of the parameter values derived from our database, we ran additional simulations by modifying the degree of viruliferous vector preference for settling on infected vs. healthy hosts. To accomplish this, we modified $\varepsilon$ to simulate loss of orientation preference after acquisition (chooses equally among infected and healthy hosts, $\varepsilon = 1$) (Model 2) and reversal of orientation preference after acquisition ($\varepsilon = \delta$) (Model 3).

$r_h$: Mean of aphid "R" parameter on all healthy plants.

$r_i$: Mean of aphid "R" parameter on infected plants, either infected by PL or NPL viruses.

$a_h$: Mean of (1 – "retention" rate) of aphid vector on healthy plants.

$a_i$: Mean of (1 – "retention" rate) of aphid vector on infected plants, either infected by PL or NPL viruses.

$\delta$: Mean of "headspace" / "Y-olfactometer" orientation rates of aphid vector towards noninfected plants (multiplied by two).

$\mathcal{E}$: 1 – $\delta$ for the main model; $\mathcal{E} = 1$ for the second model and $\mathcal{E} = \delta$ for the third model.

$\mu$: Calculated based on Power (1991); 1 – mean (recovery rates).

$\beta_v$: Calculated based on Jimenez and Bosque-Perez (2004) and Wang and Ghabrial (2007), for PL and NPL viruses, respectively. Mean (transmission rates).

$K_h$, $K_i$, $F$, $\xi$, $c_1$, $c_2$, $\beta_i$ and $\gamma$: Data from Shaw *et al.* (2017) ($c_1$: Dixon and Glen (1971); $c_2 = 0.3\ c_1$; $\beta_i$: Jimenez and Bosque-Perez (2004) for PL viruses and Mello et al. (2011) for NPL viruses)

*Original model parameters*

**Table S2:** Original model parameters (from Shaw *et al.* (2017)).

| Parameters | Units | Value range | Values (PT) | Values (NPT) |
|---|---|---|---|---|
| $r_h$ intrinsic vector growth rate on healthy hosts | 1 $d^{-1}$ | (0.01–0.6) | 0.186† | 0.27‡ |
| $r_i$ intrinsic vector growth rate on infected hosts | 1 $d^{-1}$ | (0.01–0.6) | 0.263† | 0.31‡ |
| $K_h$ vector carrying capacity on healthy hosts | vectors/host | (20–200) | 100 | 100 |
| $K_i$ vector carrying capacity on infected hosts | vectors/host | (20–200) | 100 | 100 |
| $F$ field density | 1,000,000 hosts/ha | | 4 (2.5–5) | |
| | 10,000 hosts/ha | | | 4 (2.5–5) |
| $\xi$ minimum host/ha needed to support the vectors | host/ha | 0.01($F$) | 0.01$F$ | 0.01$F$ |
| $a_h$ maximum vector departure rate from healthy hosts | 1 $d^{-1}$ | (0.014–1.4) | 0.1268†,†† | 0.1268†,†† |
| $a_i$ maximum vector departure rate from infected hosts | 1 $d^{-1}$ | (0.014–1.4) | 0.1438†,†† | 0.1438†,†† |
| $c_1$ same status half-departure constant | vectors/hosts | (1.37–137) | 13.76†† | 13.76†† |
| $c_2$ different status half-departure constant | vectors/host | (1.37–137) | 0.3$c_1$ | 0.3$c_1$ |
| $\delta$ preference of nonvirulent vectors for settling on healthy hosts | | (0.25–4) | 0.9376§ | 0.3612¶ |
| $\varepsilon$ preference of virulent vectors for settling on infected hosts | | (0.25–4) | 0.7649§ | 2.7684¶ |
| $\mu$ dispersal loss | | (0–1) | 99.4%‡‡ | 99.4%‡‡ |
| $\beta_v$ rate vector on infected host becomes virulent | 1 $d^{-1}$ | (0.7–1) | 0.68§§ | 1 |
| $\beta_i$ rate healthy hosts become infected | host $vector^{-1}$ $d^{-1}$ | (0.2–1) | 1§§ | 0.182¶¶ |
| $\gamma$ vector recovery rate by feeding | 1 $d^{-1}$ | 0, $\beta_v$ | 0 | 1 |

*Notes:* PT, persistently transmitted; NPT, nonpersistently transmitted.
Sources: †Jiménez-Martínez et al. (2004*a*), ‡Castle and Berger (1993), §Ingwell et al. (2012), ¶Castle et al. (1998), ††Dixon and Glen (1971), ‡‡Ward et al. (1998), §§Jiménez-Martínez and Bosque-Pérez (2004), ¶¶Mello et al. (2011).

**List of references:**


Castle, S.J. & Berger, P.H. (1993). Rates of growth and increase of Myzus persicae on virus-infected potatoes according to type of virus-vector relationship. Entomol. Exp. Appl., 69, 51–60.

Castle, S.J. & Mowry, T.M. (1998). Differential Settling by Myzus persicae (Homoptera: Aphididae) on Various Virus Infected Host Plants. Ann. Entomol. Soc. Am., 91, 661–667.

Dixon, A.F.G. & Glen, D.M. (1971). Morph determination in the bird cherry-oat aphid, Rhopalosiphum padi L. Ann. Appl. Biol., 68, 11–21.

Ingwell, L.L., Eigenbrode, S.D. & Bosque-Pérez, N.A. (2012). Plant viruses alter insect behavior to enhance their spread. Sci. Rep., 2, 1–6.

Jiménez-Martínez, E.S. & Bosque-Pérez, N.A. (2004). Variation in Barley Yellow Dwarf Virus Transmission Efficiency by Rhopalosiphum padi (Homoptera: Aphididae) after Acquisition from Transgenic and Nontransformed Wheat Genotypes. J. Econ. Entomol., 97, 1790–1796.

Jiménez-Martínez, E.S., Bosque-Pérez, N.A., Berger, P.H. & Zemetra, R.S. (2004a). Life history of the bird cherry-oat aphid, Rhopalosiphum padi (Homoptera: Aphididae), on transgenic and untransformed wheat challenged with Barley yellow dwarf virus. J. Econ. Entomol., 97, 203–212.

Mello, A.F.S., Olarte, R.A., Gray, S.M. & Perry, K.L. (2011). Transmission Efficiency of Potato virus Y strains PVY O and PVY N-Wi by Five Aphid Species. Plant Dis., 95, 1279–1283.

Power, A.G. (1991). Virus Spread and Vector Dynamics in Genetically Diverse Plant Populations. Ecology, 72, 232–241.

Shaw, A.K., Peace, A., Power, A.G. & Bosque-Pérez, N.A. (2017). Vector population growth and condition-dependent movement drive the spread of plant pathogens. Ecology, 98, 2145–2157.

Wang, R.Y. & Ghabrial, S.A. (2007). Effect of Aphid Behavior on Efficiency of Transmission of Soybean mosaic virus by the Soybean-Colonizing Aphid, Aphis glycines. Plant Dis., 86, 1260–1264.

Ward, S.A., Leather, S.R., Pickup, J. & Harrington, R. (1998). Mortality during dispersal and the cost of host specificity in parasites: How many aphids find hosts? J. Anim. Ecol., 67, 763–773.

Start of content for file ESM_Model outputs.xlsx

Main model: vector preference remains fixed before and after virus acquisition

Model 2: vector preference becomes neutral (no preference for infected or healthy) after virus acquisition (conditional vector preference).

Model 3: vector preference reverses to favor healthy plants after virus acquisition (conditional vector preference).

Main model outputs

| PL recovery rate / PL parameters (Main model) | | | |
|---|---|---|---|
| **Parameters** | **Preference** | **Performance** | **Departure** |
| **Graph_range** | 0.61-0.98 | 0.27-0.31 | 0.16-0.32 |
| | 40% | 13% | 50% |
| **Δ°** | 3 days | 1 day | 1 day |
| | -5.50% | -1.85% | -1.85% |
| **Δ° for 10% range** | -1.38% | -1.42% | -0.37% |
| **Interaction (not corrected by %)** | 4 days faster -7.50% | | |
| | | 1 day faster -1.85% | |
| | 5 days faster -9.26% | | |

| NPL recovery rate / NPL parameters (Main model) | | | |
|---|---|---|---|
| **Parameters** | **Preference** | **Performance** | **Departure** |
| **Graph_range** | 0,66-0,78 | 0,235-0,275 | 0,26-0,43 |
| | 15% | 15% | 40% |
| **Δ°** | 5 days | 4 days | 4 days |
| | -5.43% | -4.35% | -4.35% |
| **Δ° for 10% range** | -3.62% | -2.90% | -1.09% |
| **Interaction (not corrected by %)** | 8 days faster -8.70% | | |
| | | 8 days faster -8.70% | |
| | 8 days faster -8.70% | | |

| PL recovery rate / NPL parameters (Main model) | | | |
|---|---|---|---|
| **Parameters** | **Preference** | **Performance** | **Departure** |
| **Graph_range** | 0.61-0.98 | 0.27-0.31 | 0.16-0.32 |
| | 40% | 13% | 50% |
| **Δ°** | 3 days | 1 day | 1 day |
| | -5.50% | -1.85% | -1.85% |
| **Δ° for 10% range** | -1.38% | -1.42% | -0.37% |
| **Interaction (not corrected by %)** | 4 days faster -7.50% | | |
| | | 1 day faster -1.85% | |
| | 5 days faster -9.26% | | |

| NPL recovery rate / PL parameters (Main model) | | | |
|---|---|---|---|
| **Parameters** | **Preference** | **Performance** | **Departure** |
| **Graph_range** | 0,66-0,78 | 0,235-0,275 | 0,26-0,43 |
| | 15% | 15% | 40% |
| **Δ°** | 5 days | 5 days | 5 days |
| | -5.43% | -5.43% | -5.43% |
| **Δ° for 10% range** | -3.62% | -3.62% | -1.36% |
| **Interaction (not corrected by %)** | 8 days faster -8.70% | | |
| | | 8 days faster -8.70% | |
| | 8 days faster -8.70% | | |

Model 2 outputs

| PL recovery rate / PL parameters (Model 2) | | | |
|---|---|---|---|
| Parameters | Preference | Performance | Departure |
| Graph_range | 0.61-0.98<br>40% | 0.27-0.31<br>13% | 0.16-0.32<br>50% |
| Δ° | 3 days<br>-5.50% | 1 day<br>-1.85% | 1 day<br>-1.85% |
| Δ° for 10% range | -1.38% | -1.42% | -0.37% |
| Interaction (not corrected by %) | 4 days faster<br>-7.50% | | |
| | | 2 days faster<br>-3.70% | |
| | 5 days faster<br>-9.26% | | |

| NPL recovery rate / NPL parameters (Model 2) | | | |
|---|---|---|---|
| Parameters | Preference | Performance | Departure |
| Graph_range | 0,66-0,78<br>15% | 0,235-0,275<br>15% | 0,26-0,43<br>40% |
| Δ° | 5 days<br>-5.43% | 4 days<br>-4.35% | 4 days<br>-4.35% |
| Δ° for 10% range | -3.62% | -2.90% | -1.09% |
| Interaction (not corrected by %) | 10 days faster<br>-10.87% | | |
| | | 10 days faster<br>-10.87% | |
| | 10 days faster<br>-10.87% | | |

| PL recovery rate / NPL parameters (Model 2) | | | |
|---|---|---|---|
| Parameters | Preference | Performance | Departure |
| Graph_range | 0.61-0.98<br>40% | 0.27-0.31<br>13% | 0.16-0.32<br>50% |
| Δ° | 4 days<br>-7.41% | 1 day<br>-1.85% | 1 day<br>-1.85% |
| Δ° for 10% range | -1.85% | -1.42% | -0.37% |
| Interaction (not corrected by %) | 4 days faster<br>-7.50% | | |
| | | 2 days faster<br>-3.70% | |
| | 5 days faster<br>-9.26% | | |

| NPL recovery rate / PL parameters (Model 2) | | | |
|---|---|---|---|
| Parameters | Preference | Performance | Departure |
| Graph_range | 0,66-0,78<br>15% | 0,235-0,275<br>15% | 0,26-0,43<br>40% |
| Δ° | 4 days<br>-4.35% | 4 days<br>-4.35% | 5 days<br>-5.43% |
| Δ° for 10% range | -2.90% | -2.90% | -1.36% |
| Interaction (not corrected by %) | 8 days faster<br>-8.70% | | |
| | | 10 days faster<br>-10.87% | |
| | 8 days faster<br>-8.70% | | |

Model 3 outputs

| PL recovery rate / PL parameters (Model 3) | | | |
|---|---|---|---|
| **Parameters** | **Preference** | **Performance** | **Departure** |
| **Graph_range** | 0.61-0.98 | 0.27-0.31 | 0.16-0.32 |
| | 40% | 13% | 50% |
| **Δ°** | 4 days | 1 day | 1 day |
| | -7.41% | -1.85% | -1.85% |
| **Δ° for 10% range** | -1.85% | -1.42% | -0.37% |
| **Interaction (not corrected by %)** | 4 days faster -7.50% | | |
| | | 2 days faster -3.70% | |
| | 5 days faster -9.26% | | |

| NPL recovery rate / NPL parameters (Model 3) | | | |
|---|---|---|---|
| **Parameters** | **Preference** | **Performance** | **Departure** |
| **Graph_range** | 0,66-0,78 | 0,235-0,275 | 0,26-0,43 |
| | 15% | 15% | 40% |
| **Δ°** | 5 days | 4 days | 4 days |
| | -5.43% | -4.35% | -4.35% |
| **Δ° for 10% range** | -3.62% | -2.90% | -1.09% |
| **Interaction (not corrected by %)** | 10 days faster -10.87% | | |
| | | 10 days faster -10.87% | |
| | 10 days faster -10.87% | | |

| PL recovery rate / NPL parameters (Model 3) | | | |
|---|---|---|---|
| **Parameters** | **Preference** | **Performance** | **Departure** |
| **Graph_range** | 0.61-0.98 | 0.27-0.31 | 0.16-0.32 |
| | 40% | 13% | 50% |
| **Δ°** | 4 days | 1 day | 1 day |
| | -7.41% | -1.85% | -1.85% |
| **Δ° for 10% range** | -1.85% | -1.42% | -0.37% |
| **Interaction (not corrected by %)** | 4 days faster -7.50% | | |
| | | 2 days faster -3.70% | |
| | 5 days faster -9.26% | | |

| NPL recovery rate / PL parameters (Model 3) | | | |
|---|---|---|---|
| **Parameters** | **Preference** | **Performance** | **Departure** |
| **Graph_range** | 0,66-0,78 | 0,235-0,275 | 0,26-0,43 |
| | 15% | 15% | 40% |
| **Δ°** | 5 days | 5 days | 5 days |
| | -5.43% | -5.43% | -5.43% |
| **Δ° for 10% range** | -3.62% | -3.62% | -1.36% |
| **Interaction (not corrected by %)** | 8 days faster -8.70% | | |
| | | 8 days faster -8.70% | |
| | 8 days faster -8.70% | | |

Simulations Model 2: Time to 80% infected hosts infected for PL case, PL recovery rate (0) with PL parameters for orientation preference/performance/departure rate

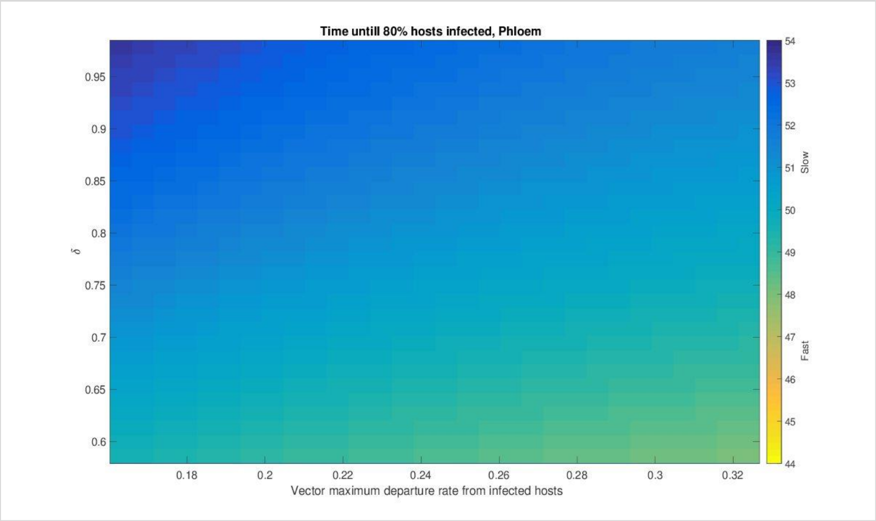


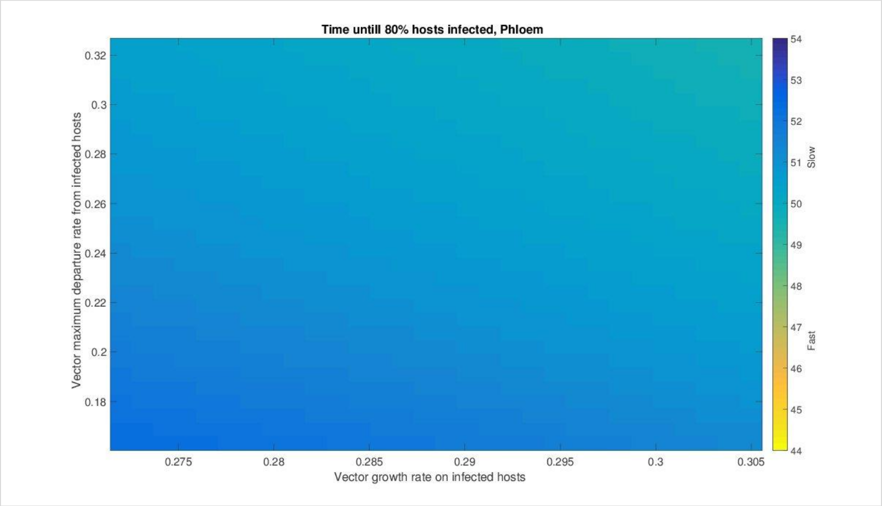


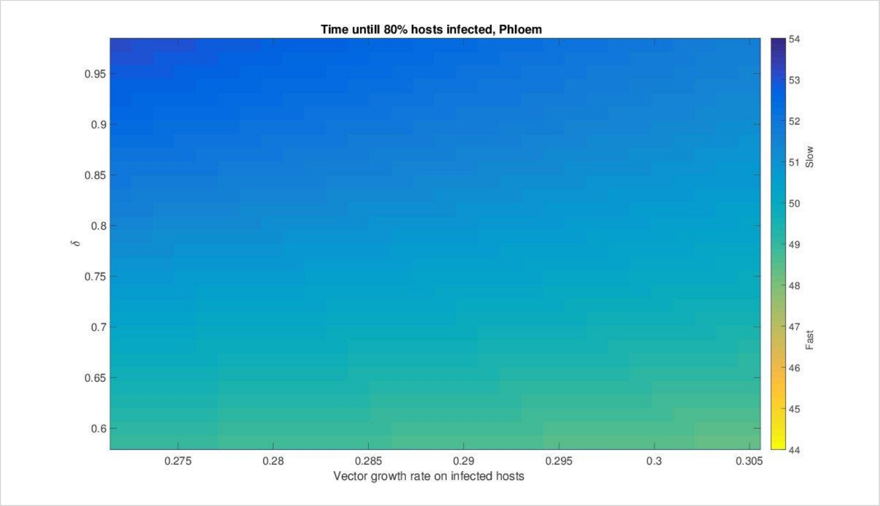

Simulations Model 2: Time to 80% infected hosts infected for PL case, PL recovery rate (0) with **NPL parameters** for orientation preference/performance/departure rate

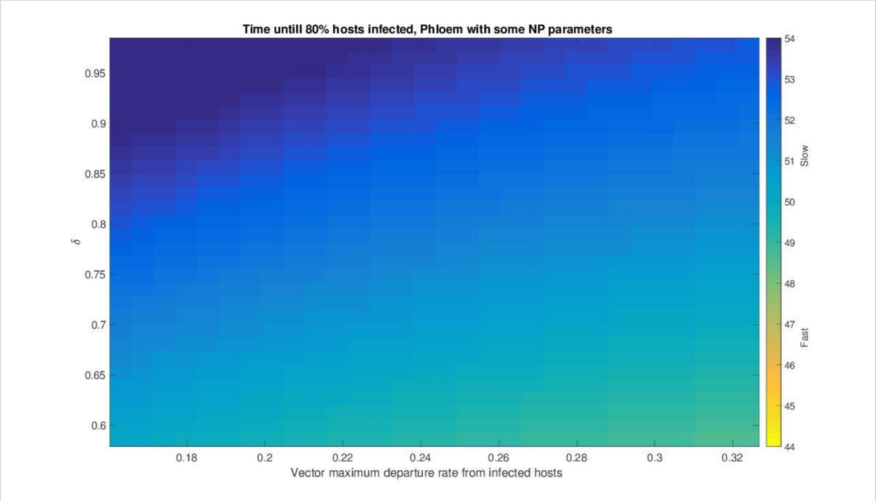


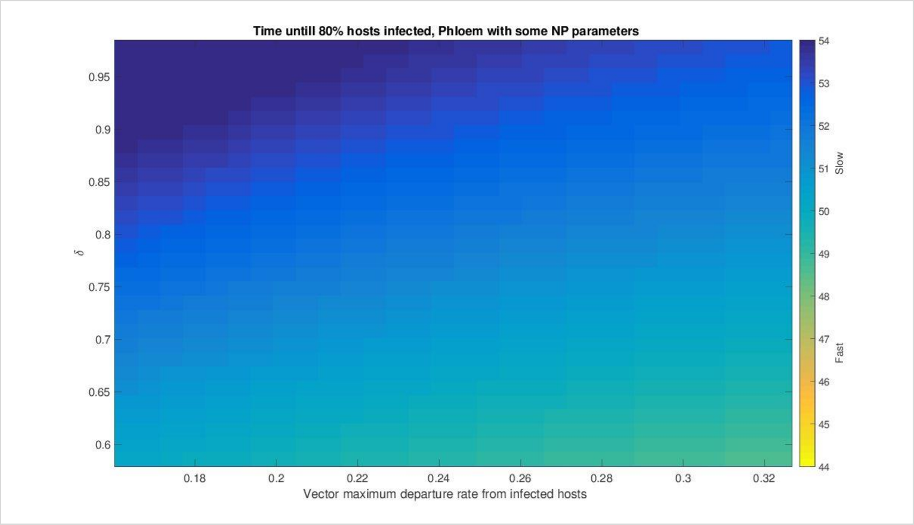


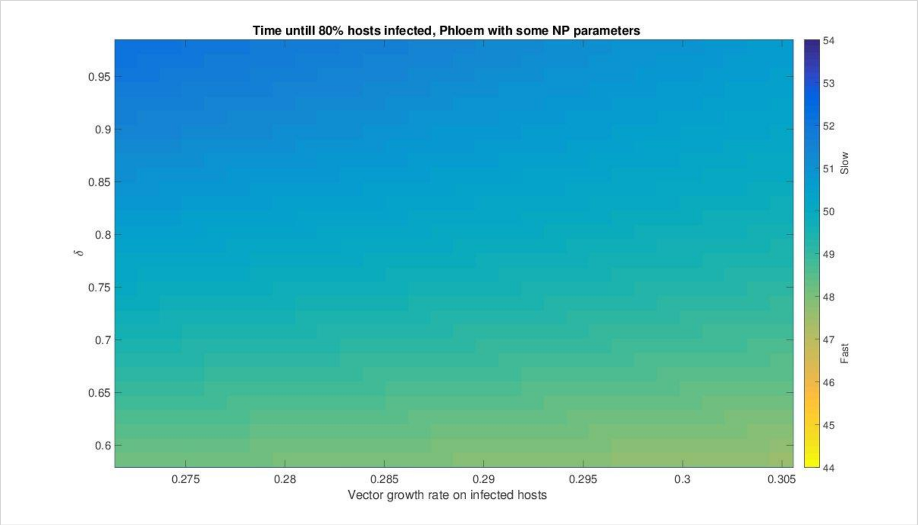

Simulations Model 2: Time to 80% infected hosts infected for NPL case, NPL recovery rate (0) with NPL parameters for orientation preference/performance/departure rate

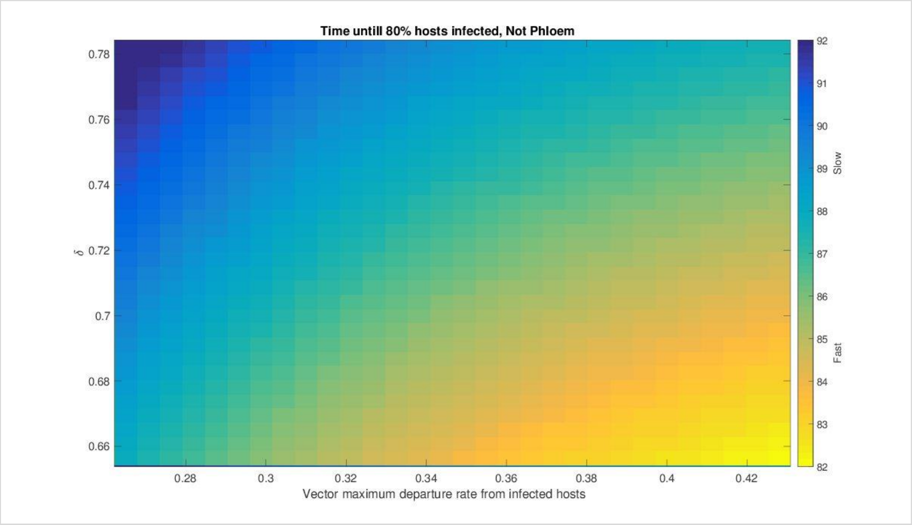


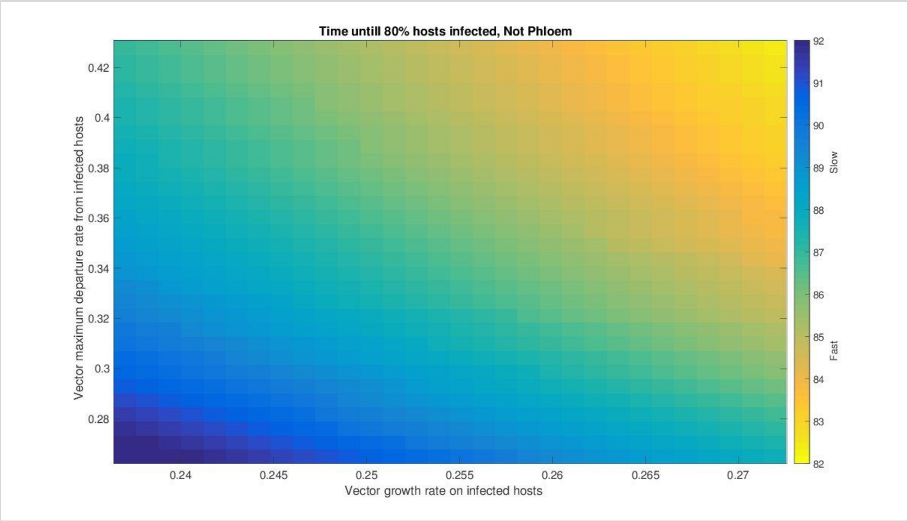


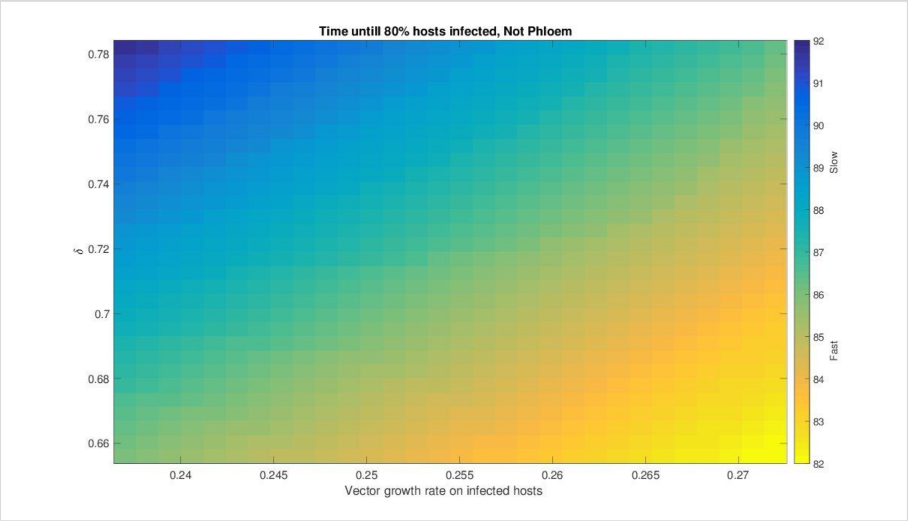

Simulations Model 2: Time to 80% infected hosts infected for NPL case, NPL recovery rate (0) with **PL parameters** for orientation preference/performance/departure rate.

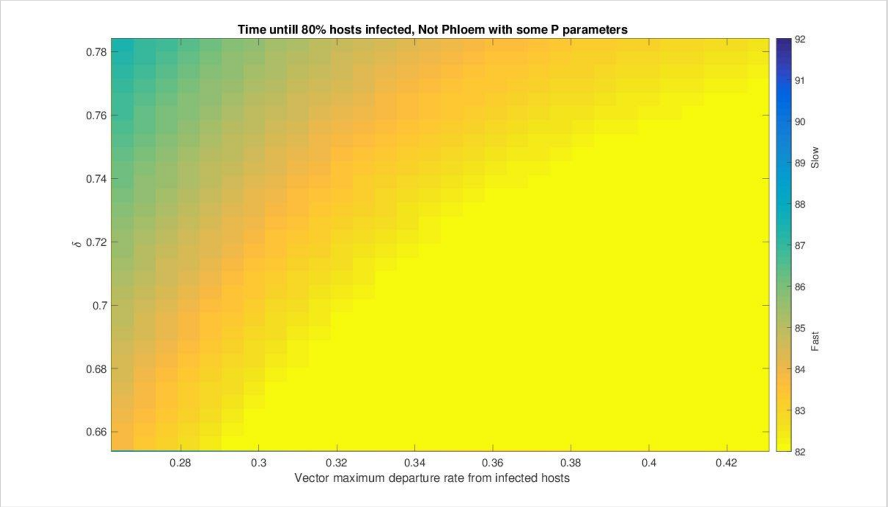


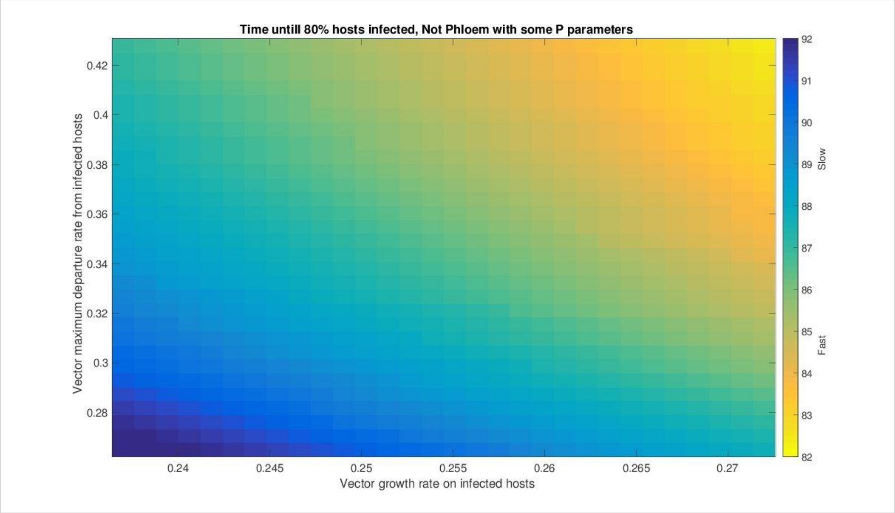


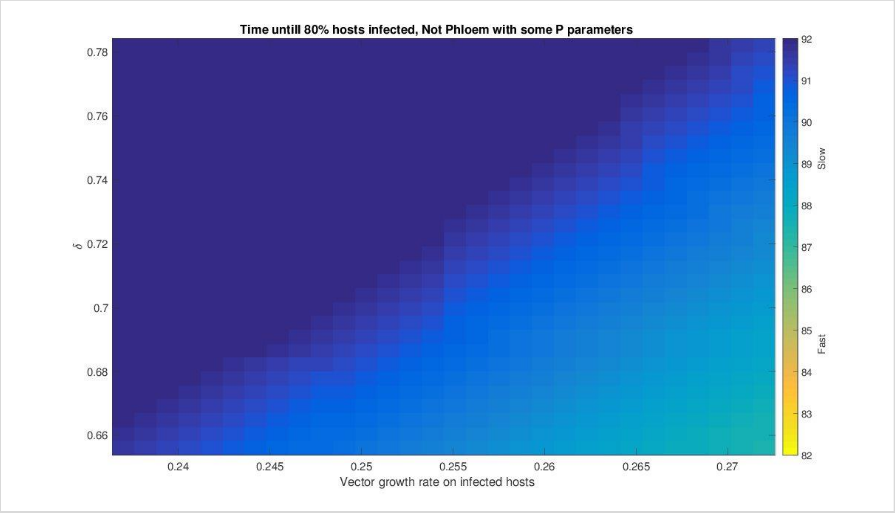

Simulations Model 3: Time to 80% infected hosts infected for PL case, PL recovery rate (0) with PL parameters for orientation preference/performance/departure rate.

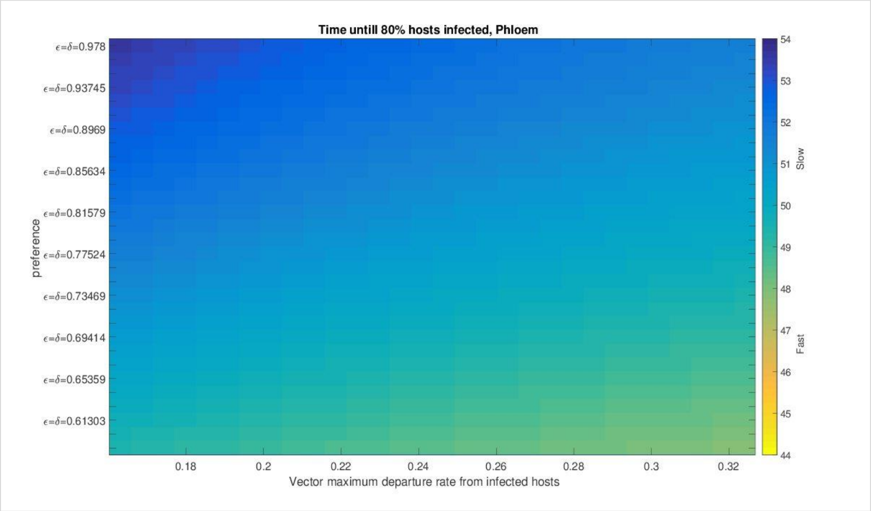


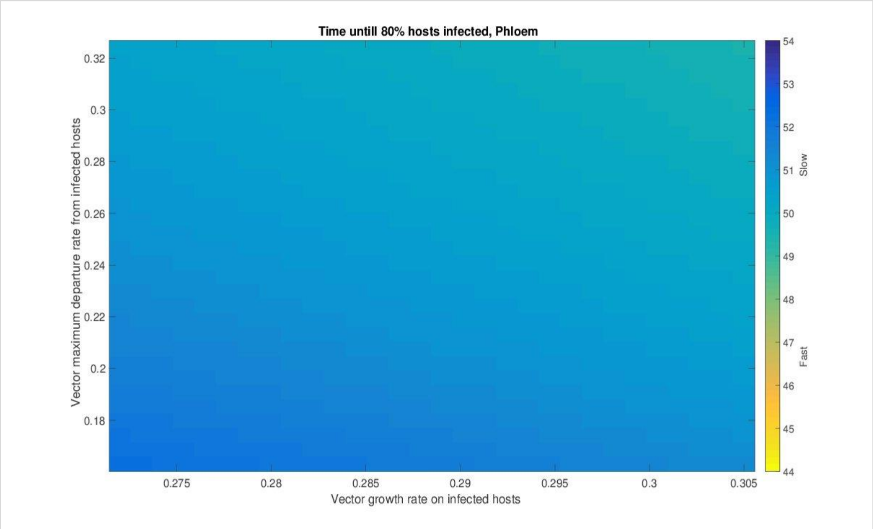


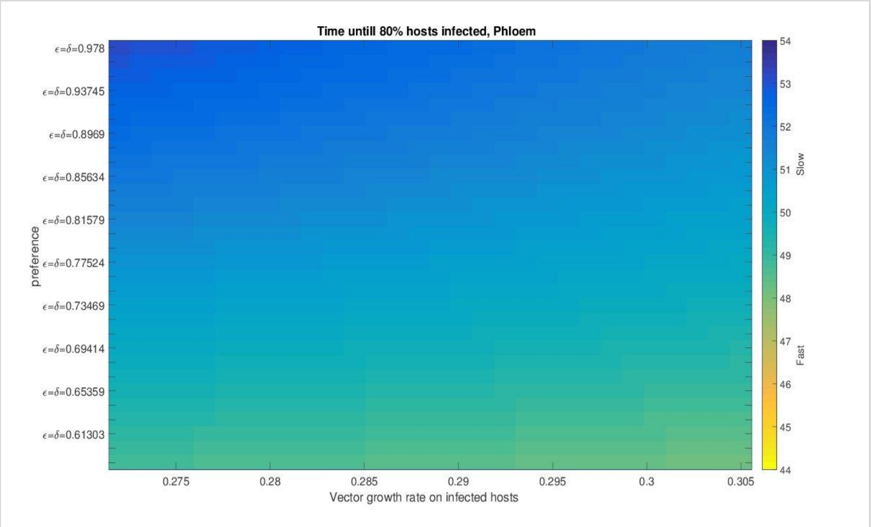

Simulations Model 3: Time to 80% infected hosts infected for PL case, PL recovery rate (0) with **NPL parameters** for orientation preference/performance/departure rate.

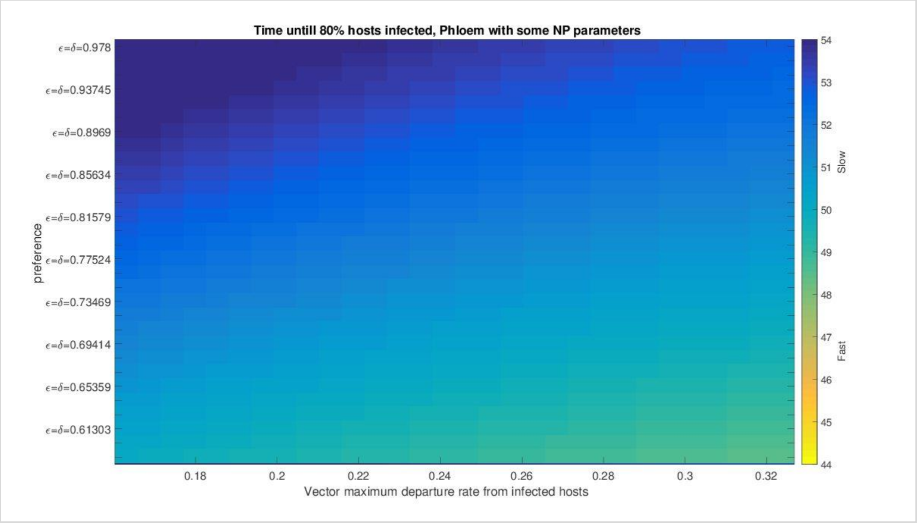


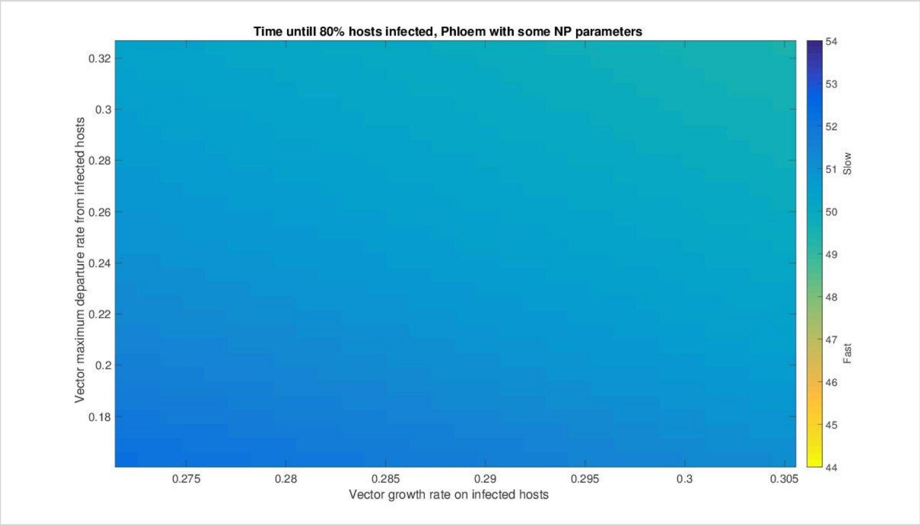


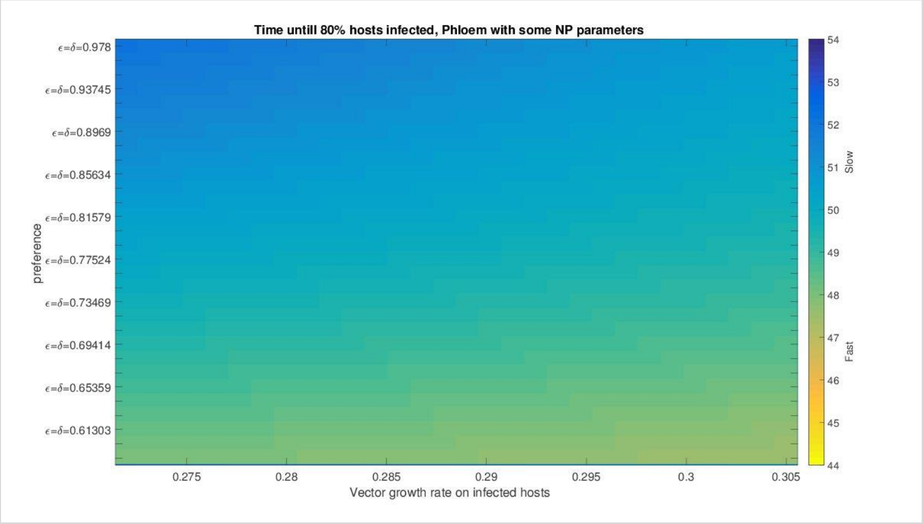

Simulations Model 3: Time to 80% infected hosts infected for NPL case, NPL recovery rate (0) with NPL parameters for orientation preference/performance/departure rate.

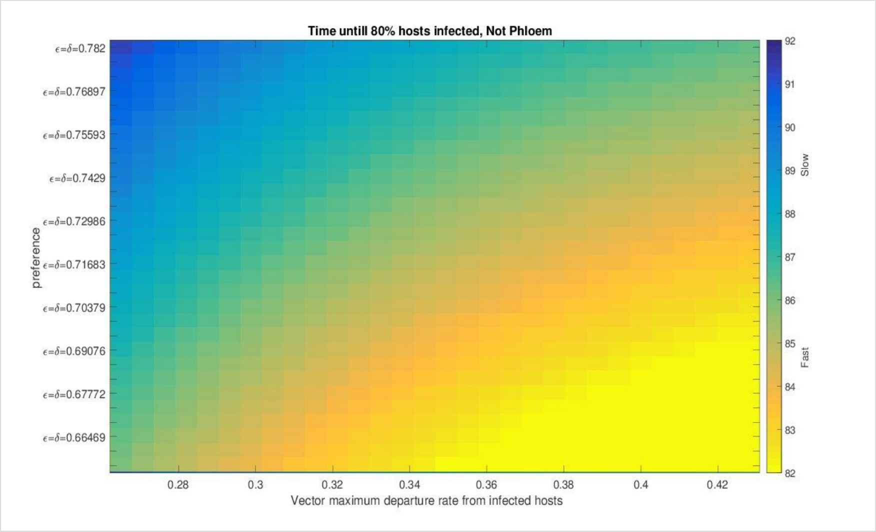


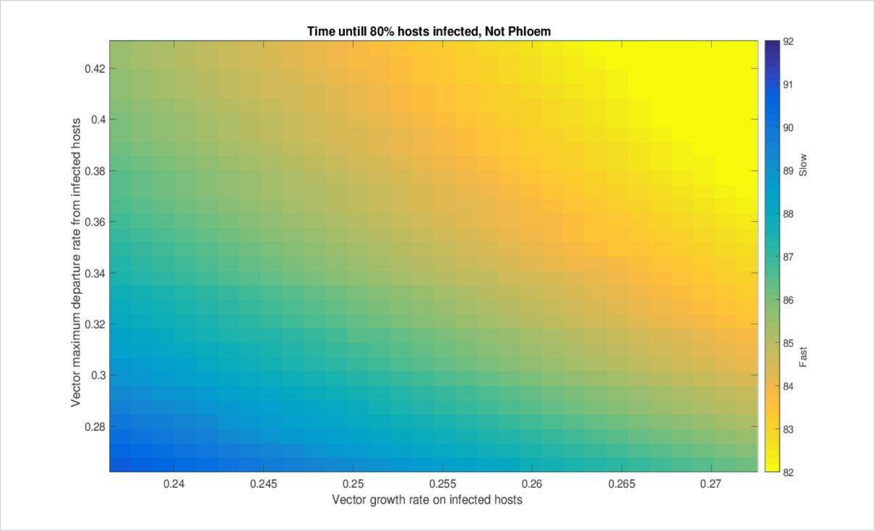


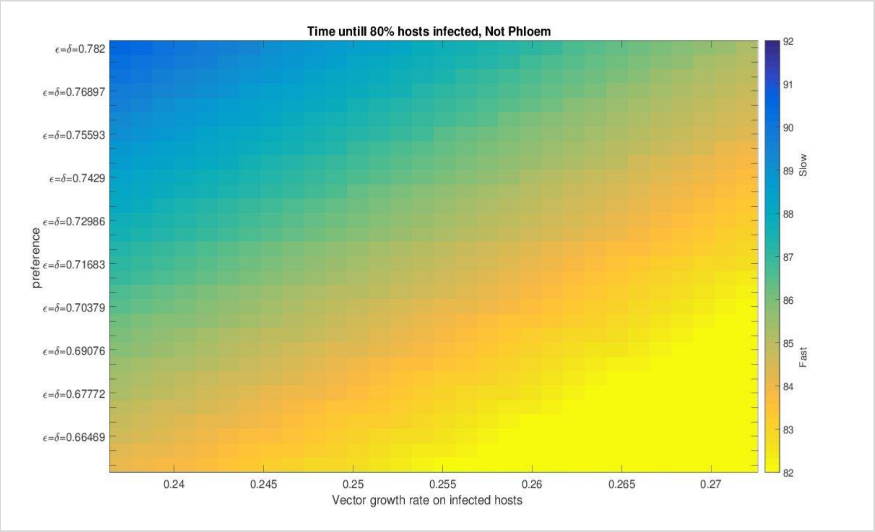

Simulations Model 3: Time to 80% infected hosts infected for NPL case, NPL recovery rate (0) with **PL parameters** for orientation preference/performance/departure rate.

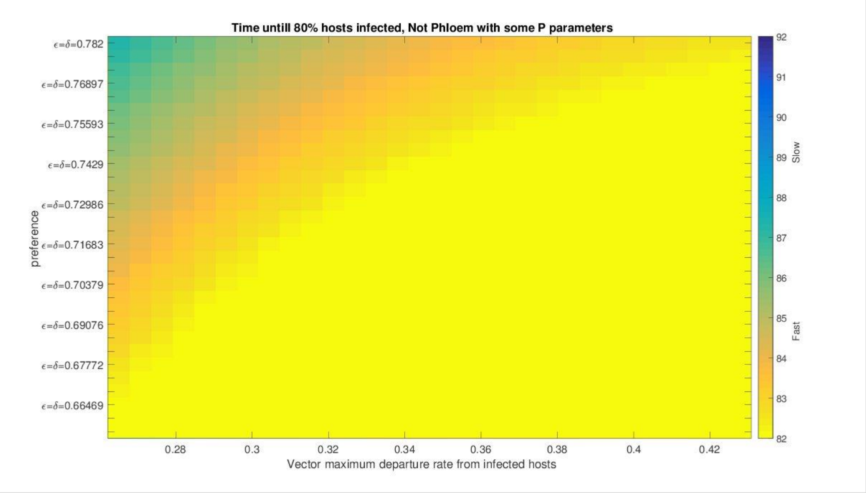


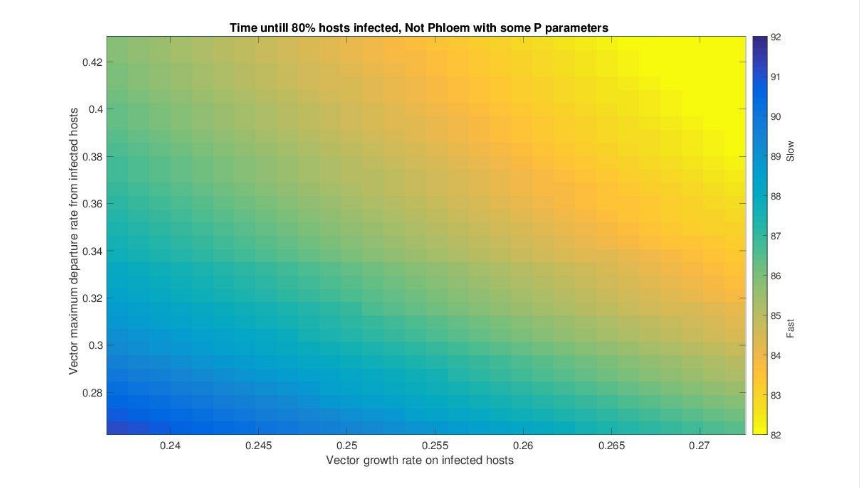


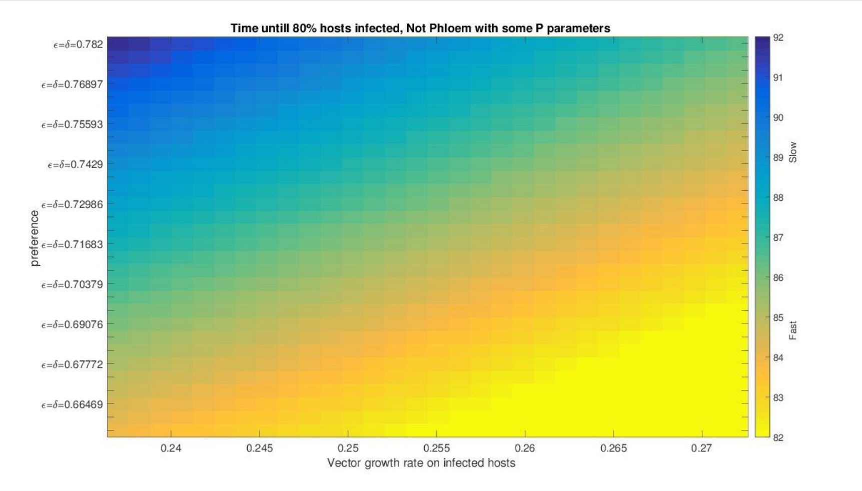

### All models: Central values

#### MAIN MODEL

| PL recovery/PL parameters | | | |
|---|---|---|---|
| Parameters | a) Depart~Rm | b) Pref~Rm | c) Pref~Depart |
| Central value | 51 | 50 | 50 |
| Mean | 50.33333333 | | |

| PL recovery/NPL parameters | | | |
|---|---|---|---|
| Parameters | a) Depart~Rm | b) Pref~Rm | c) Pref~Depart |
| Central value | 51 | 50 | 52 |
| Mean | 51 | | |

| NPL recovery/NPL parameters | | | |
|---|---|---|---|
| Parameters | a) Depart~Rm | b) Pref~Rm | c) Pref~Depart |
| Central value | 88 | 87 | 87 |
| Mean | 87.33333333 | | |

| NPL recovery/PL parameters | | | |
|---|---|---|---|
| Parameters | a) Depart~Rm | b) Pref~Rm | c) Pref~Depart |
| Central value | 88 | 92 | 84 |
| Mean | 88 | | |

| | | | Parameters | | |
|---|---|---|---|---|---|
| | | | PL | NPL | |
| | | | 1.31% | | |
| Recovery | PL | 42.37% | 50.333 | 51 | 42.05% |
| | NPL | | 87.333 | 88 | |
| | | | -0.76% | | |

#### MODEL 2 : loss of preference after virus acquisition

| PL recovery/PL parameters | | | |
|---|---|---|---|
| Parameters | a) Depart~Rm | b) Pref~Rm | c) Pref~Depart |
| Central value | 51 | 50 | 50 |
| Mean | 50.33333333 | | |

| PL recovery/NPL parameters | | | |
|---|---|---|---|
| Parameters | a) Depart~Rm | b) Pref~Rm | c) Pref~Depart |
| Central value | 51 | 50 | 51 |
| Mean | 50.66666667 | | |

| NPL recovery/NPL parameters | | | |
|---|---|---|---|
| Parameters | a) Depart~Rm | b) Pref~Rm | c) Pref~Depart |
| Central value | 86 | 86 | 86 |
| Mean | 86 | | |

| NPL recovery/PL parameters | | | |
|---|---|---|---|
| Parameters | a) Depart~Rm | b) Pref~Rm | c) Pref~Depart |
| Central value | 86 | 91 | 82 |
| Mean | 86.33333333 | | |

| | | | Parameters | | |
|---|---|---|---|---|---|
| | | | PL | NPL | |
| | | | 0.66% | | |
| Recovery | PL | 41.47% | 50.333 | 50.667 | 41.31% |
| | NPL | | 86 | 86.333 | |
| | | | -0.39% | | |

#### MODEL 3 : Preference reversion after virus acquisition

| PL recovery/PL parameters | | | |
|---|---|---|---|
| Parameters | a) Depart~Rm | b) Pref~Rm | c) Pref~Depart |
| Central value | 51 | 50 | 50 |
| Mean | 50.33333333 | | |

| PL recovery/NPL parameters | | | |
|---|---|---|---|
| Parameters | a) Depart~Rm | b) Pref~Rm | c) Pref~Depart |
| Central value | 50 | 50 | 51 |
| Mean | 50.33333333 | | |

| NPL recovery/NPL parameters | | | |
|---|---|---|---|
| Parameters | a) Depart~Rm | b) Pref~Rm | c) Pref~Depart |
| Central value | 85 | 85 | 85 |
| Mean | 85 | | |

| NPL recovery/PL parameters | | | |
|---|---|---|---|
| Parameters | a) Depart~Rm | b) Pref~Rm | c) Pref~Depart |
| Central value | 85 | 85 | 82 |
| Mean | 84 | | |

| | | | Parameters | | |
|---|---|---|---|---|---|
| | | | PL | NPL | |
| | | | 0.00% | | |
| Recovery | PL | 40.78% | 50.333 | 50.333 | 40.08% |
| | NPL | | 85 | 84 | |
| | | | 1.18% | | |

### Start of content for file ESM_Model_raw data.xlsx

Data used to derive parameter estimates for “orientation preference”

| Reference | Reference # | Virus | phloem_acq | Healthy_mea | Healthy_SD | Healthy_n | Virus_mean | Virus_SD | Virus_n | Parameter |
|---|---|---|---|---|---|---|---|---|---|---|
| Alvarez | 1 | PLRV-Wageningen | yes | 2.07 | 1.15 | 6 | 7.84 | 1.42 | 6 | headspace |
| Alvarez | 1 | PLRV-Wageningen | yes | 2.90 | 1.46 | 6 | 3.82 | 2.22 | 6 | headspace |
| Claudel | 2 | TuYV | yes | 26.53 | 9.77 | 10 | 44.89 | 15.56 | 10 | headspace |
| Claudel | 2 | TuYV | yes | 27.72 | 13.17 | 15 | 23.91 | 8.75 | 15 | headspace |
| dosSantos | 3 | BYDV-PAV | yes | 16.40 | 23.73 | 12 | 66.24 | 37.52 | 12 | headspace |
| dosSantos | 3 | BYDV-PAV | yes | 43.79 | 39.14 | 12 | 45.06 | 41.40 | 12 | headspace |
| Eigenbrode | 4 | PLRV | yes | 32.60 | 7.56 | 3 | 60.93 | 10.39 | 3 | headspace |
| Eigenbrode | 4 | PLRV | yes | 33.60 | 6.61 | 6 | 66.40 | 6.61 | 6 | headspace |
| Jimenez | 7 | BYDV | yes | 6.30 | 2.60 | 6 | 11.80 | 3.72 | 6 | headspace |
| Jimenez | 7 | BYDV | yes | 7.70 | 2.69 | 6 | 6.72 | 1.81 | 6 | headspace |
| Jimenez | 7 | BYDV | yes | 10.00 | 2.06 | 10 | 14.10 | 5.03 | 10 | headspace |
| Jimenez | 7 | BYDV | yes | 10.50 | 2.25 | 10 | 14.00 | 2.62 | 10 | headspace |
| Mauck | 9 | CMV-FNY | no | 1.62 | 2.01 | 24 | 3.21 | 1.71 | 24 | headspace |
| Mauck | 9 | CMV-FNY | no | 1.91 | 0.54 | 24 | 3.66 | 3.23 | 24 | headspace |
| Mauck | 9 | CMV-FNY | no | 1.09 | 1.52 | 24 | 1.94 | 1.52 | 24 | headspace |
| Mauck | 9 | CMV-FNY | no | 2.22 | 1.32 | 24 | 3.88 | 2.01 | 24 | headspace |
| Mauck | 10 | CMV-KVPG2 | no | 1.24 | 0.47 | 9 | 1.81 | 0.95 | 9 | headspace |
| Mauck | 10 | CMV-KVPG2 | no | 1.11 | 0.38 | 9 | 2.44 | 1.54 | 9 | headspace |
| Mauck | 10 | CMV-P1 | no | 1.16 | 0.83 | 9 | 1.70 | 0.82 | 9 | headspace |
| Medina | 11 | BYDV-PAV | yes | 4.10 | 3.07 | 10 | 7.90 | 3.76 | 10 | headspace |
| Medina | 11 | BYDV-PAV | yes | 5.10 | 2.12 | 10 | 4.90 | 3.00 | 10 | headspace |
| Rajabaskar | 15 | PLRV | yes | 36.86 | 10.69 | 15 | 63.21 | 9.30 | 15 | headspace |
| Safari | 24 | CMV | no | 36.62 | 3.36 | 12 | 64.24 | 0.94 | 12 | headspace |
| Wu | 22 | PEMV | yes | 39.73 | 18.68 | 13 | 60.43 | 19.90 | 13 | Y |
| Wu | 22 | BLRV | yes | 40.54 | 14.46 | 13 | 59.62 | 16.91 | 13 | Y |
| Wu | 23 | CMV | no | 34.07 | 11.53 | 65 | 65.93 | 11.47 | 65 | Y |

Data used to derive parameter estimates for “vector maximum departure rate” (settling/feeding preference)

| Reference | Reference # | Virus | phloem_acq | Healthy_mean | Healthy_SD | Healthy_n | Virus_mean | Virus_SD | Virus_n | Parameter |
|---|---|---|---|---|---|---|---|---|---|---|
| Blua | 5 | ZYMV | no | 2.33 | 0.80 | 12 | 2.33 | 0.83 | 12 | retention |
| Blua | 5 | ZYMV | no | 2.33 | 0.80 | 12 | 1.60 | 0.80 | 12 | retention |
| Chesnais | 15 | TuYV | yes | 62.30 | 18.98 | 15 | 60.40 | 24.79 | 15 | retention |
| Chesnais | 15 | TuYV | yes | 42.80 | 10.84 | 15 | 69.00 | 20.91 | 15 | retention |
| Chesnais | 15 | TuYV | yes | 50.00 | 23.24 | 15 | 31.50 | 25.56 | 15 | retention |
| Chesnais | 15 | TuYV | yes | 32.40 | 27.11 | 15 | 21.50 | 20.91 | 15 | retention |
| Chesnais | 15 | CaMV | no | 69.70 | 15.88 | 15 | 66.80 | 16.65 | 15 | retention |
| Chesnais | 15 | CaMV | no | 50.30 | 22.08 | 15 | 50.30 | 22.85 | 15 | retention |
| Chesnais | 15 | CaMV | no | 31.70 | 13.56 | 15 | 39.20 | 17.43 | 15 | retention |
| Chesnais | 15 | CaMV | no | 32.00 | 18.98 | 15 | 30.90 | 15.49 | 15 | retention |
| Chesnais | 65 | TuYV | yes | 53.59 | 16.88 | 15 | 50.50 | 11.29 | 15 | retention |
| Chesnais | 65 | TuYV | yes | 57.19 | 31.62 | 15 | 79.31 | 19.41 | 15 | retention |
| Chesnais | 65 | TuYV | yes | 39.67 | 18.90 | 15 | 49.30 | 17.82 | 15 | retention |
| Chesnais | 65 | TuYV | yes | 37.91 | 16.17 | 15 | 64.97 | 14.08 | 15 | retention |
| Chesnais | 65 | TuYV | yes | 48.70 | 22.03 | 15 | 57.69 | 22.92 | 15 | retention |
| Chesnais | 65 | TuYV | yes | 27.00 | 16.27 | 15 | 50.00 | 23.32 | 15 | retention |
| Fereres | 21 | SMV | no | 9.37 | 0.96 | 16 | 8.38 | 1.16 | 16 | retention |
| Fereres | 21 | SMV | no | 7.49 | 1.92 | 16 | 6.31 | 1.32 | 16 | retention |
| Mauck | 37 | CMV | no | 38.50 | 5.18 | 4 | 28.15 | 5.50 | 4 | retention |
| Mauck | 37 | CMV | no | 42.49 | 5.64 | 4 | 17.61 | 4.26 | 4 | retention |
| Mauck | 38 | CMV | no | 25.96 | 2.44 | 5 | 21.44 | 3.67 | 5 | retention |
| Mauck | 38 | CMV | no | 27.17 | 2.24 | 5 | 16.01 | 4.67 | 5 | retention |
| Mauck | 38 | CMV | no | 17.29 | 7.26 | 9 | 15.07 | 1.71 | 9 | retention |
| Mauck | 38 | CMV | no | 4.58 | 2.73 | 9 | 4.70 | 3.36 | 9 | retention |
| Mauck | 38 | CMV | no | 24.84 | 3.72 | 9 | 28.47 | 2.37 | 9 | retention |
| Mauck | 38 | CMV | no | 26.46 | 2.76 | 9 | 29.49 | 2.37 | 9 | retention |
| Mauck | 38 | CMV | no | 19.02 | 3.88 | 4 | 17.02 | 2.48 | 4 | retention |
| Mauck | 38 | CMV | no | 11.02 | 3.06 | 4 | 11.98 | 4.96 | 4 | retention |
| Mauck | 38 | CMV | no | 28.73 | 3.56 | 5 | 27.91 | 3.85 | 5 | retention |
| Mauck | 38 | CMV | no | 28.97 | 2.71 | 5 | 26.83 | 4.52 | 5 | retention |
| Rajabaskar | 48 | PLRV | yes | 28.82 | 8.69 | 15 | 78.17 | 6.68 | 15 | retention |
| Westwood | 61 | CMV | no | 10.00 | 0.00 | 3 | 5.97 | 1.99 | 3 | retention |

Data used to derive parameter estimates for “vector maximum growth rate” (performance)

| Reference | Virus | Family | phloem_acq | Healthy_mean | Healthy_SD | Healthy_n | Virus_mean | Virus_SD | Virus_n | Parameter |
|---|---|---|---|---|---|---|---|---|---|---|
| Chesnais | CaMV | Caulimoviridae | no | 0.2690 | 0.0312 | 39 | 0.2420 | 0.0292 | 34 | R |
| Chesnais | CaMV | Caulimoviridae | no | 0.2210 | 0.0493 | 30 | 0.1170 | 0.0408 | 26 | R |
| Donaldson | AMV | Bromoviridae | no | 0.3590 | 0.0069 | 12 | 0.3960 | 0.0104 | 12 | R |
| Gadhave | PRSV | Potyviridae | no | 0.3474 | 0.0827 | 20 | 0.4738 | 0.0979 | 20 | R |
| Hily | CMV | Bromoviridae | no | 0.3642 | 0.0280 | 25 | 0.3752 | 0.0230 | 25 | R |
| Hily | CMV | Bromoviridae | no | 0.3531 | 0.0275 | 25 | 0.3660 | 0.0243 | 25 | R |
| Hodge | BYMV | Potyviridae | no | 0.2800 | 0.1196 | 143 | 0.2800 | 0.0566 | 32 | R |
| Ren | PVY | Potyviridae | no | 0.3300 | 0.0520 | 3 | 0.3500 | 0.0693 | 3 | R |
| Wosula | SPFMV | Potyviridae | no | 0.3380 | 0.0447 | 20 | 0.3380 | 0.0492 | 20 | R |
| Wosula | SPFMV | Potyviridae | no | 0.3190 | 0.0581 | 20 | 0.2760 | 0.0716 | 20 | R |
| Wosula | SPFMV | Potyviridae | no | 0.1090 | 0.1699 | 20 | 0.0320 | 0.1565 | 20 | R |
| Chesnais | TuYV | Luteoviridae | yes | 0.2690 | 0.0312 | 39 | 0.2970 | 0.0375 | 39 | R |
| Chesnais | TuYV | Luteoviridae | yes | 0.2210 | 0.0493 | 30 | 0.1440 | 0.0384 | 23 | R |
| Chesnais | TuYV | Luteoviridae | yes | 0.2749 | 0.0379 | 32 | 0.2542 | 0.0281 | 33 | R |
| Chesnais | TuYV | Luteoviridae | yes | 0.2822 | 0.0509 | 35 | 0.2231 | 0.0431 | 34 | R |
| Chesnais | TuYV | Luteoviridae | yes | 0.2399 | 0.0390 | 32 | 0.2457 | 0.0445 | 31 | R |
| Davis | BLRV | Luteoviridae | yes | 0.1950 | 0.0569 | 10 | 0.2260 | 0.0411 | 10 | R |
| Davis | BLRV | Luteoviridae | yes | 0.2470 | 0.0727 | 10 | 0.2700 | 0.1075 | 10 | R |
| Davis | BLRV | Luteoviridae | yes | 0.2080 | 0.0190 | 10 | 0.2030 | 0.0791 | 10 | R |
| Davis | BLRV | Luteoviridae | yes | 0.1440 | 0.0443 | 10 | 0.2550 | 0.0854 | 10 | R |
| Davis | BLRV | Luteoviridae | yes | 0.1570 | 0.0759 | 10 | 0.1930 | 0.9487 | 10 | R |
| Davis | BLRV | Luteoviridae | yes | 0.3590 | 0.0506 | 10 | 0.3640 | 0.0348 | 10 | R |
| Fereres | BYDV_PAV | Luteoviridae | yes | 0.2900 | 0.0283 | 8 | 0.3100 | 0.0283 | 8 | R |
| Fereres | BYDV_RPV | Luteoviridae | yes | 0.2900 | 0.0283 | 8 | 0.3000 | 0.0283 | 8 | R |
| Fereres | BYDV_PAV | Luteoviridae | yes | 0.2900 | 0.0283 | 8 | 0.3100 | 0.0283 | 8 | R |
| Fereres | BYDV_RPV | Luteoviridae | yes | 0.2900 | 0.0283 | 8 | 0.3200 | 0.0283 | 8 | R |
| Fereres | BYDV_PAV | Luteoviridae | yes | 0.2900 | 0.0283 | 8 | 0.3200 | 0.0283 | 8 | R |
| Fereres | BYDV_RPV | Luteoviridae | yes | 0.2900 | 0.0283 | 8 | 0.2900 | 0.0283 | 8 | R |
| Fiebig | BYDV_MAV | Luteoviridae | yes | 0.2000 | 0.0517 | 107 | 0.1600 | 0.0474 | 90 | R |
| Fiebig | BYDV_PAV | Luteoviridae | yes | 0.2000 | 0.0517 | 107 | 0.1600 | 0.0581 | 69 | R |
| Fiebig | BYDV_MAV | Luteoviridae | yes | 0.1600 | 0.0551 | 62 | 0.1500 | 0.0421 | 71 | R |
| Fiebig | BYDV_PAV | Luteoviridae | yes | 0.1600 | 0.0551 | 62 | 0.1300 | 0.0458 | 84 | R |
| Hodge | PEMV | Luteoviridae | yes | 0.2800 | 0.1196 | 143 | 0.2800 | 0.0583 | 34 | R |
| Jimenez | BYDV_PAV | Luteoviridae | yes | 0.2390 | 0.0139 | 12 | 0.2630 | 0.0277 | 12 | R |
| Jimenez | BYDV_PAV | Luteoviridae | yes | 0.1730 | 0.0416 | 12 | 0.2440 | 0.0346 | 12 | R |
| Jimenez | BYDV_PAV | Luteoviridae | yes | 0.2630 | 0.0485 | 12 | 0.2290 | 0.0589 | 12 | R |
| Jimenez | BYDV_PAV | Luteoviridae | yes | 0.2720 | 0.0346 | 12 | 0.2340 | 0.0208 | 12 | R |
| Moiroux | TuYV | Luteoviridae | yes | 0.2700 | 0.0219 | 30 | 0.3010 | 0.0329 | 30 | R |

### Start of content for file ESM_vector database.docx

**Electronic Supplementary Materials: “Vector database”**

***Review of virus-vector relationships***

We explored the additional dimension of “transmission opportunities” to determine how the number of competent vector species (those capable of transmitting) relates to virus effects on host phenotypes, model simulations, and study limitations. For all viruses included in the meta-analysis, we surveyed the literature and recorded the number of identified competent vectors species. First, we considered online databases such as Plant Virus Online (Brunt et al., 1997), The Universal Virus Database (*ICTVdB)* (Buchen-Osmond, 2006) and DPVweb (Adams & Antoniw, 2006). When different numbers of competent vectors were indicated on these databases, we kept the higher value. To complete our database, we also surveyed Edwardson & Christie (1991) which reviewed characteristics of viruses associated with legume crops, as well as specific articles for viruses characterized more recently and not reported in databases (Ghosh, Das, Vijayanandraj, & Mandal, 2016). The assembled database, and a complete list of references, are provided in the electronic supplementary material (ESM_vector database.docx).

| Family | Virus species | Vector number | References |
|---|---|---|---|
| **Bromoviridae** | | | |
| | AMV | 22 | (Edwardson & Christie 1991) |
| | CMV | 87 | (Edwardson & Christie 1991) |
| **Caulimoviridae** | | | |
| | CaMV | 27 | DPVweb[a]: (Kennedy *et al.* 1962) |
| **Closteroviridae** | | | |
| | BYV | 22 | DPVweb: (Kennedy *et al.* 1962) |
| | LIYV | 1 | DPVweb: (Duffus *et al.* 1986) |
| | RLMV | 1 | DPVweb: (Thekke-Veetil *et al.* 2017) |
| | ToCV | 1 | DPVweb: (Wisler *et al.* 1998) |
| **Geminiviridae** | | | |
| | CdTV | 1 | DPVweb: (Brown & Nelson 1988) |
| | CLCuV | 3 | (Edwardson & Christie 1991) |
| | CLCV | 3 | (Edwardson & Christie 1991) |
| | EACMV | 1 | DPVweb: (Dubern 1994) |
| | TbCSV | 1 | DPVweb: (Varma & Malathi 2003) |
| | ToMoV | 1 | ICTVdB Management (2006) [b] |
| | ToSRV | 1 | ICTVdB Management (2006) |
| | TYLCCNV | 1 | ICTVdB Management (2006) |
| | TYLCV | 1 | ICTVdB Management (2006) |
| | WCMoV | 1 | ICTVdB Management (2006) |
| **Luteoviridae** | | | |
| | BLRV | 12 | (Edwardson & Christie 1991) |

| | | | |
|---|---|---|---|
| | BYDV | 14 | DPVweb: (Rochow & Carmichael 1979) |
| | CABYV | 2 | (Lecoq *et al.* 1992) |
| | PEMV | 13 | (Edwardson & Christie 1991) |
| | PLRV | 2 | DPVweb: (Kennedy *et al.* 1962) |
| | TuYV | 17 | (Edwardson & Christie 1991) |
| **Nanoviridae** | | | |
| | CBDV | 1 | (Ghosh *et al.* 2016) |
| **Potyviridae** | | | |
| | BYMV | 47 | (Edwardson & Christie 1991) |
| | MCMV | 6 | DPVweb: (Nault *et al.* 1978) |
| | PRSV | 21 | DPVweb: (Zettler *et al.* 1968) |
| | PVY | 26 | DPVweb: (Harrington & Gibson 1989) |
| | SMV | 32 | (Edwardson & Christie 1991) |
| | SPFMV | 4 | ICTVdB Management (2006) |
| | TuMV | 89 | (Edwardson & Christie 1991) |
| | WMV | 44 | (Edwardson & Christie 1991) |
| | ZYMV | 8 | DPVweb: (Purcifull *et al.* 1984) |
| **Reoviridae** | | | |
| | RDV | 5 | DPVweb: (Shinkai 1962; Nasu 1963) |
| | RpLV | 1 | (Lightle & Lee 2014) |
| | RRSV | 1 | DPVweb: (Shikata *et al.* 1979) |
| | SRBSDV | 1 | DPVweb: (Shinkai 1967) |
| **Secoviridae** | | | |
| | BPMV | 6 | (Edwardson & Christie 1991) |
| **Sobemovirus** | | | |
| | SBMV | 3 | (Edwardson & Christie 1991) |
| **Tospoviridae** | | | |
| | GBNV | 1 | (Lakshmi *et al.* 1995) |
| | INSV | 1 | ICTVdB Management (2006) |
| | SVNV | 2 | DPVweb: (Zhou & Tzanetakis 2013) |
| | SYVV | 1 | DPVweb: (Sylvester & Richardson 1969) |
| | TSWV | 9 | (Edwardson & Christie 1991) |
| | WSMoV | 1 | (Chen *et al.* 2014) |

[a] DPVweb: (Adams & Antoniw 2006); [b] *ICTVdB - The Universal Virus Database*, version 4, April 2006.

**Figure S1**: Estimated number of competent vector species for each virus taxon included in the meta-analysis database, delineated by site of virion acquisition in the host plant. PL viruses are transmitted by fewer documented competent vector species relative to NPL viruses. Of the PL viruses considered in our study, 80.6% (25/31) have three or fewer recorded competent vector species. In contrast, for NPL viruses acquired and inoculated via rapid probes (13 viruses, excluding SBMV, which is beetle-transmitted and semi-persistent) all of the viruses covered in our meta-analysis are transmitted by at least three or more vectors and 69.2% (9/13) are transmitted by more than 20 vector species.

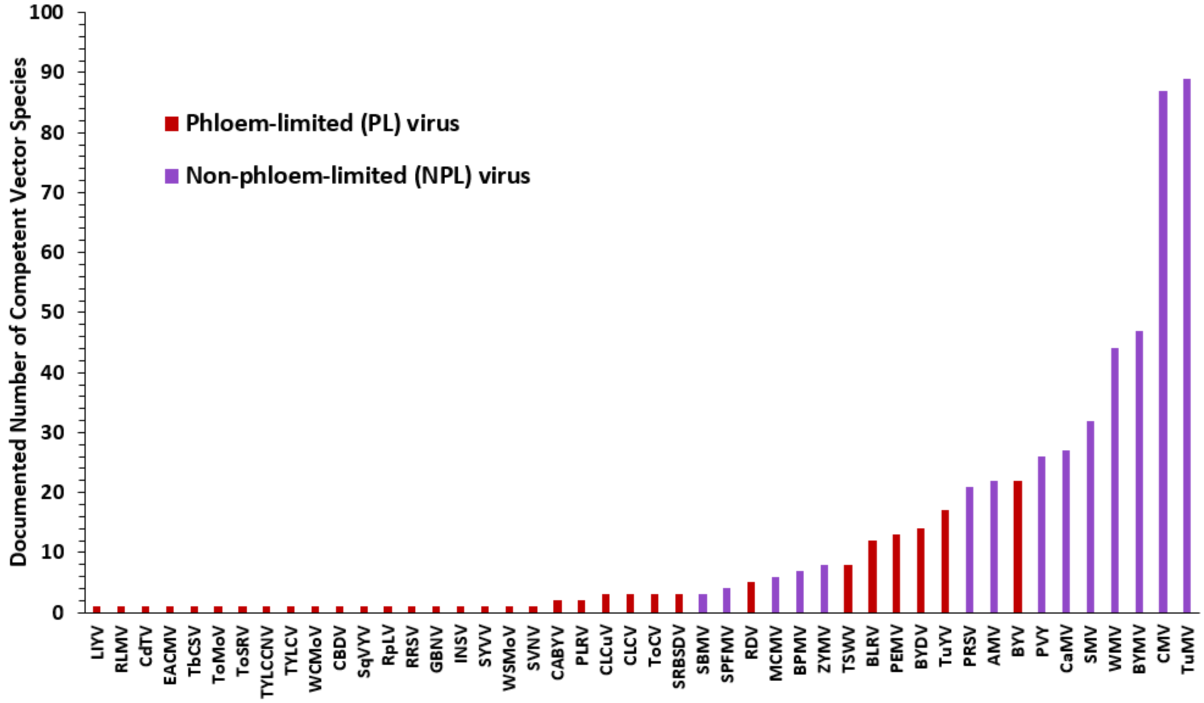


**Complete list of references:**


Adams, M.J. & Antoniw, J.F. (2006). DPVweb: a comprehensive database of plant and fungal virus genes and genomes. *Nucleic Acids Res.*, 34, D382–D385.

Brown, J.K. & Nelson, M.R. (1988). Transmission, Host Range, and Virus-Vector Relationships of Chino del Tomate Virus, a Whitefly-Transmitted Geminivirus from Sinaloa, Mexico. *Plant Dis.*, 72, 866.

Chen, W. Te, Tseng, C.H. & Tsai, C.W. (2014). Effect of Watermelon Silver Mottle Virus on the life history and feeding preference of Thrips palmi. *PLoS One*, 9.

Dubern, J. (1994). Transmission of African cassava mosaic geminivirus by the whitefly (Bemisia tabaci). *Trop. Sci.*, 34, 82–91.

Duffus, J.E., Larsen, R.C. & Liu, H.Y. (1986). Lettuce Infectious Yellows Virus - New Type of Whitefly-Transmitted Virus. *Phytopathology*, 76, 97–100.

Edwardson, J.R. & Christie, R.G. (1991). *CRC Handbook of Viruses Infecting Legumes*. *CRC Handb. Viruses Infect. Legum.*

Ghosh, A., Das, A., Vijayanandraj, S. & Mandal, B. (2016). Cardamom Bushy Dwarf Virus Infection in Large Cardamom Alters Plant Selection Preference, Life Stages, and Fecundity of Aphid Vector, *Micromyzus kalimpongensis* (Hemiptera: Aphididae). *Environ. Entomol.*, 45, 178–184.

Harrington, R. & Gibson, R.W. (1989). Transmission of potato virus Y by aphids trapped in potato crops in southern England. *Potato Res.*, 32, 167–174.

Kennedy, J.S., Day, M.F. & Eastop, V.F. (1962). *A conspectus of aphids as vectors of plant viruses.* Commonwealth Institute of Entomology, London, United Kingdom.

Lakshmi, K.V., Wightman, J.A., Reddy, D.V.R., Rao, G.V.R., Buiel, A.A.M. & Reddy, D.D.R. (1995). Transmission of Peanut Bud Necrosis Virus by Thrips palmi in India. In: *Thrips Biology and Management*. Springer US, Boston, MA, pp. 179–184.

Lecoq, H., Bourdin, D., Wipf-Scheibel, C., Bon, M., Lot, H., Lemaire, O., *et al.* (1992). A new yellowing disease of cucurbits caused by a luteovirus, cucurbit aphid-borne yellows virus. *Plant Pathol.*, 41, 749–761.

Lightle, D. & Lee, J. (2014). Raspberry viruses affect the behavior and performance of Amphorophora agathonica in single and mixed infections. *Entomol. Exp. Appl.*, 151, 57–64.

Nasu, S. (1963). Studies on some leafhoppers and planthoppers which transmit virus diseases of rice plant in Japan. *Bull. Kyushu Agric. Exp. Stn*, 8, 153–349.

Nault, L.R., Styer, W.E., Coffey, M.E., Gordon, D.T., Negi, L.S. & Niblett, C.L. (1978). Transmission of Maize Chlorotic Mottle Virus by Chrysomelid Beetles. *Phytopathology*, 68, 1071.

Purcifull, D.E., Adlerz, W.C., Simone, G.W., Hiebert, E. & Christie, S.R. (1984). Serological Relationships and Partial Characterization of Zucchini Yellow Mosaic Virus Isolated from Squash in Florida. *Plant Dis.*, 68, 230.

Rochow, W.F. & Carmichael, L.E. (1979). Specificity among barley yellow dwarf viruses in enzyme immunosorbent assays. *Virology*, 95, 415–420.

Shikata, E., Senboku, T., Kamjaipai, K., Chou, T., Tiongco, E.R. & Ling, K.C. (1979). Rice ragged stunt virus, a new member of plant reovirus group. *Ann. Phytopath. Soc. Japan*, 45, 436–443.

Shinkai, A. (1962). Studies on insect transmissions of rice virus diseases in Japan. *Bull. Natl. Inst. Agric. Sci. C*, 14, 1–112.

Shinkai, A. (1967). Transmission of rice stripe and black-streaked dwarf viruses by Ribautodelphax albifascia. *Phytopathol. Soc. Japan Ann.*, 33, 318.

Sylvester, E.S. & Richardson, J. (1969). Additional evidence of multiplication of the sowthistle yellow vein virus in an aphid vector—Serial passage. *Virology*, 37, 26–31.

Thekke-Veetil, T., Khadgi, A., Johnson, D., Burrack, H., Sabanadzovic, S. & Tzanetakis, I.E. (2017). First Report of Raspberry leaf mottle virus in Blackberry in the United States. *Plant Dis.*, 101, 265–265.

Varma, A. & Malathi, V.G. (2003). Emerging geminivirus problems: A serious threat to crop production. *Ann. Appl. Biol.*, 142, 145–164.

Wisler, G.C., Duffus, J.E., Liu, H.-Y. & Li, R.H. (1998). Ecology and Epidemiology of Whitefly-Transmitted Closteroviruses. *Plant Dis.*, 82, 270–280.

Zettler, F.W., Edwarson, J.R. & Purcifull, D.E. (1968). Ultramicroscopic differences in inclusions of papaya mosaic virus and papaya ringspot virus correlated with differential aphid transmission. *Phytopathology*, 58, 332–335.

Zhou, J. & Tzanetakis, I.E. (2013). Epidemiology of Soybean vein necrosis-associated virus. *Phytopathology*, 103, 966–971.
